# (De)composing sociality: disentangling individual-specific from dyad-specific propensities to interact

**DOI:** 10.1101/2023.08.15.549768

**Authors:** Christof Neumann, Julia Fischer

## Abstract

In group-living animals, relationships between group members are often highly differentiated. Some dyads can maintain strong and long-lasting relationships, while others are only connected by weak or fleeting ties. Evidence accumulates that aspects of social relationships are related to reproductive success and survival. Yet, few of these analyses have considered that frequent or prolonged affiliative interactions between two individuals can be principally driven by two distinct processes: namely, the overall gregariousness of individuals, and dyadic affinity, i.e., the preference the members of the dyad have to interact specifically with one another. Crucially, these two axes of sociality cannot be observed directly, although distinguishing them is essential for many research questions, for example, when estimating kin bias or when studying the link between sociality and fitness. We present a principled statistical framework to estimate the two underlying sociality axes using dyadic interaction data. We also provide a basic R package `bamoso`, which implements models based on the proposed framework and allows visual and numerical evaluation of the estimated sociality axes. We demonstrate critical features of the proposed modeling framework with simulated and empirical data: (1) the possibility of checking model fit against observed data, (2) the assessment of uncertainty in the estimated sociality parameters, and (3) the possibility to extend it to more complex models that use interaction data to estimate the relationship between individual-level social features and individual-level outcomes in a unified model. Our model provides a principled foundation to explain variation in dyadic interactions. This approach allows us to address questions about the relationship between variation in sociality characteristics and other features of interest, both within and across species.

## 1 Introduction

Individuals in many group-living species maintain differentiated dyadic relationships. Analyses of these relationships allow us to infer and describe key components of a society’s social structure (Hinde, 1976; Kappeler & van Schaik, 2002), including its social complexity (Bergman & Beehner, 2015; Fischer et al., 2017; Kappeler, 2019). Numerous studies have shown positive associations between fitness and various aspects of affiliative relationships, for example cumulative relationship strength (empirical examples: Cameron et al., 2009; Frère et al., 2010; Silk et al., 2010; reviews: Ostner & Schülke, 2018; Snyder-Mackler et al., 2020). However, other studies failed to find such evidence, and reported context-dependent or even negative associations between sociality- and fitness-related measures (Duboscq et al., 2023; Kalbitzer et al., 2017; Menz et al., 2020; Thompson & Cords, 2018; Wey & Blumstein, 2012). Dyadic relationships are also crucial in models of pathogen and knowledge transmission (e.g., Drewe & Perkins, 2015; Nightingale et al., 2015).

A crucial first step in such studies of social relationships is their quantification. Unfortunately, this is not a trivial endeavor. Relationships cannot be observed or measured directly: they are constructs inferred from interactions or spatio-temporal associations between individuals. These observed data may take the form of frequencies or durations. They are often processed to represent adjusted rates, proportions or aggregated into behavioral indices (figure 1).

**Figure 1:**
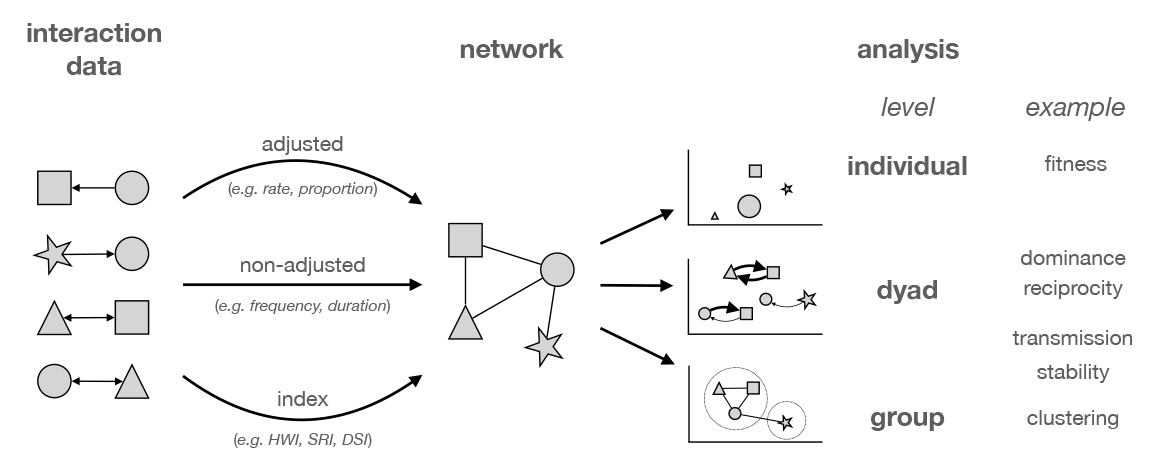
When data on dyadic interactions or associations are collected, they are often processed to represent social networks. These networks, or rather the dyadic connections, can either be generated directly from observed, non-adjusted interactions (e.g., mating or dominance networks), via adjusted observations (e.g., rates of grooming, proportion of time co-feeding), or derived indices based on the observed data (association indices (HWI, SRI), composite dyadic sociality index (DSI)). Setting up the network is usually a means to an end in this workflow; the goal is to learn about the behavior of individuals, dyads, or groups.

A crucial first step in such studies of social relationships is their quantification. Unfortunately, this is not a trivial endeavor. Relationships cannot be observed or measured directly: they are constructs inferred from interactions or spatio-temporal associations between individuals. These observed data may take the form of frequencies or durations. They are often processed to represent adjusted rates, proportions or aggregated into behavioral indices (figure 1).

While the behavior observed to quantify features of social relationships is dyadic in nature, it is essential to realize that dyadic interaction patterns are not driven by a single unobservable trait that can be assigned to each dyad. How often (or how long) two individuals interact not only depends on the specific relationship between the two individuals and their propensity to interact with each other, i.e., their dyadic affinity, but also on the individual propensities of the two individuals to interact with anyone, i.e., their individual gregariousness (Godde et al., 2013; Pepper et al., 1999; Whitehead & James, 2015) (figure 2).

**Figure 2:**
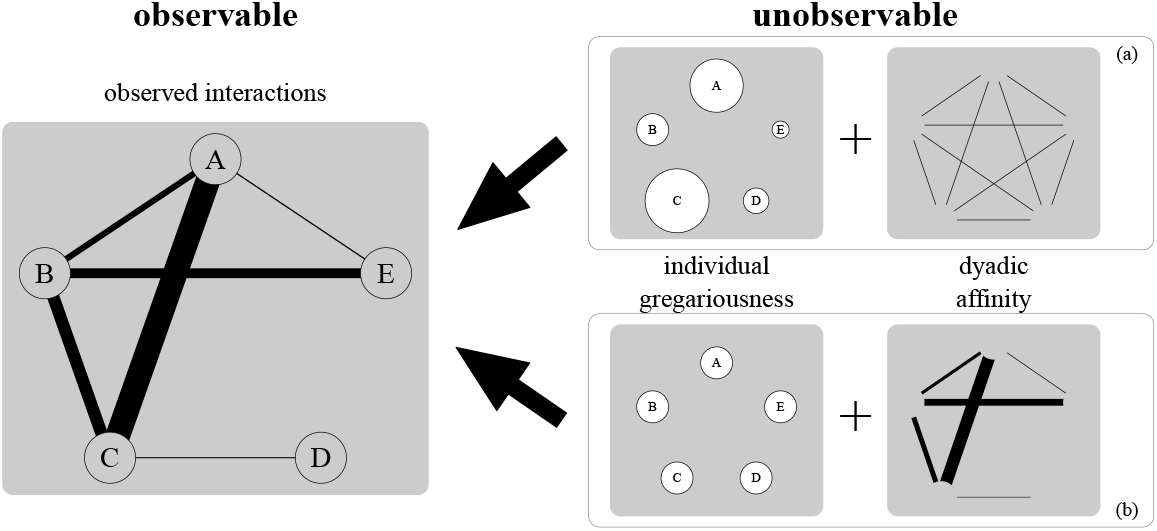
A schematic summary of our (modelling) framework. Interactions arise from a combination of individual (gregariousness) and dyadic (affinity) propensities to interact. While we observe interactions, the underlying propensities to interact are unobservable and can only be estimated from interaction data. Varying dyadic interaction durations or frequencies are represented via line width. Individual gregariousness is encoded in circle size and dyadic affinity is encoded in line width. In scenario (a), variation in interaction patterns is driven entirely by differences in individual gregariousness because all dyadic affinities are identical. In scenario (b), all individuals are equally gregarious and variation in interactions is driven by differences in dyadic affinity. Crucially, the observed interaction pattern is the same in both scenarios despite the fundamentally different underlying processes. Unlike node-level metrics in the context of network analysis (e.g., centrality), gregariousness is not an emergent feature, but is an individual feature that is estimated simultaneously with dyadic affinity in our framework.

Two individuals may interact frequently because they are both intrinsically very social (have high propensities to interact with anyone, e.g., large gregariousness values for A and C in figure 2a) and not because they have a strong affinity with each other. Likewise, two individuals may interact because they have a strong affinity with each other, even though they might both not have high propensities to interact in general (B and E in figure 2b). Simply considering observed interactions does not inform us about the relative contributions of the individual- and dyad-level propensities because the observed interactions can arise from a mixture of both processes (figure 2). In other words: the frequency or duration of dyadic interactions is due to at least the predispositions of individuals to *interact in general*, and the predispositions of dyads to *interact with one another*.

Previous attempts to disentangle individual gregariousness and dyadic affinity can be mainly found in the context of analyzing spatio-temporal associations (e.g., Godde et al., 2013; Pepper et al., 1999; Surbeck et al., 2017; Whitehead & James, 2015). The overall idea of these approaches was to arrive at dyadic affinity values while ‘controlling’ for gregariousness. These approaches have at least two drawbacks. First, individual gregariousness is not actually estimated in these procedures, but only approximated post-hoc by summing dyadic affinity values for each individual or averaging the number of individuals the focal animal associates with (Godde et al., 2013; Whitehead & James, 2015). Second, the resulting dyadic affinity estimates are hard to interpret because they are on an arbitrary scale. There is thus no straightforward way to assess how well the estimated model parameters map onto observed association data, i.e., on the actual behavior of the studied animals.

Some approaches also exist for analyzing actual dyadic interactions, as opposed to spatiotemporal associations. Most relevant for us is the social relations model (SRM). The SRM is concerned with directed interaction data and its focus is on estimating correlations between individual level effects (generalized reciprocity) and within-dyad correlations (dyadic reciprocity) (Koster et al., 2020; Ross et al., 2024). Indeed, parameters in the SRM can be interpreted similarly as gregariousness and affinity when applied to directed data, i.e., individual level propensities to give, individual level propensities to receive and dyadic propensities to interact. What is missing from these approaches (and their implementations, e.g., in the STRAND and bisonR packages, Ross et al. (2024), Hart et al. (2023)) is the ability to separate individual and dyad level propensities when analyzing undirected data and when data come from more than one behavior (multilayer networks).

We aim to offer an addition to the toolbox of methods for analyzing undirected interaction data to inform our assessment of social relationships. Our approach can be seen as a modification of the social relations model. We outline a general, flexible, and extendable modeling framework that allows differentiating the contributions of individual and dyadic propensities to observed interaction and association patterns. We use *observed* dyadic interaction patterns to estimate the underlying but unobservable axes of individual-level gregariousness and dyad-level affinity.

We do this by fitting a model that reflects an assumed generative process based on contributions from individuals and dyads to explain variation in the observed interactions between dyads in a Bayesian framework (Koster et al., 2020; McElreath, 2020; Redhead et al., 2024; Ross et al., 2024). In contrast to previous approaches such as those implemented in `STRAND` and `bisonR` (Hart et al., 2023; Ross et al., 2024), our approach allows for the simultaneous consideration of multiple behaviors.

Our approach has at least five advantageous features. First, we can obtain estimates for dyadic *and* individual propensities from a single model fitted to a given data set (including the potential to assess uncertainty in these estimates directly, see Lusseau et al. (2008) and Farine & Strandburg-Peshkin (2015), see also supplement S1). Second, we can make predictions from this model, which allows for objectively assessing model fit, i.e., we can directly compare predicted interaction patterns to observed interaction patterns (posterior predictive checks, Gabry et al. (2019)). Third, we can model multiple interaction types simultaneously, even with different error distributions (for example, the frequency of grooming bouts and durations of proximity). Fourth, the model can be extended such that the sociality features can be estimated and simultaneously used as predictors in downstream models of other quantities of interest, carrying over uncertainty from the sociality estimates if desired. Fifth, it is also possible to extend the model so that we can use the estimated sociality features as a response variable (Rathke & Fischer, 2021).

## 2 Outline of the modelling approach

In line with the notion that dyadic interactions are the outcome of individual and dyadic propensities to engage in interactions, we can summarize our approach as follows:

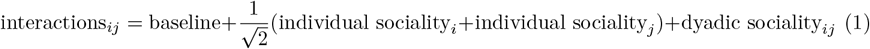

where interactions_*ij*_ reflects the observed interactions between individuals *i* and *j*. The terms individual gregariousness_*i*_ and individual gregariousness_*j*_ correspond to the individual gregariousness values of *i* and *j*, and dyadic affinity_*ij*_ is the pairwise affinity for dyad *ij*. We multiply the sum of the two individual gregariousness values by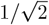in order to bring the two underlying distributions of gregariousness and affinity to the same scale. The *baseline* term reflects the expected interactions for two individuals of average individual gregariousness that have an average dyadic affinity. Because it is an intercept term, its primary purpose is to calibrate the average to a level that makes the model applicable and the parameters comparable even if the absolute rates or interactions of interactions differ substantially across behavior types (or group identity, species, season, etc.). In other words, we are primarily interested in within-group processes and as such the baseline parameter is of no direct conceptual interest to us in the context of the present work.

The model we described extends to more than one observed behavior in a straightforward way:

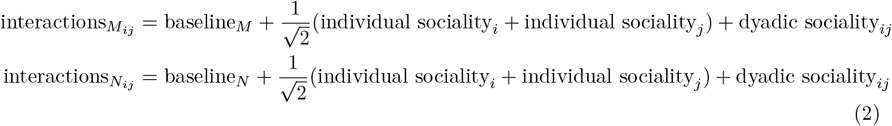

where the only addition is that we incorporate a separate baseline parameter for each additional behavior (here behavior *M* and behavior *N*).

More formally, we can express the model as follows (see supplement S2 for Stan code). We are assuming for the sake of illustration that we observed two behaviors: a count variable, *y*_*ij*_ (for example, the frequency with which two individuals groomed or the number of times two individuals approached each other) and a proportion, *z*_*ij*_ (for example the proportion of time two individuals spent in proximity).

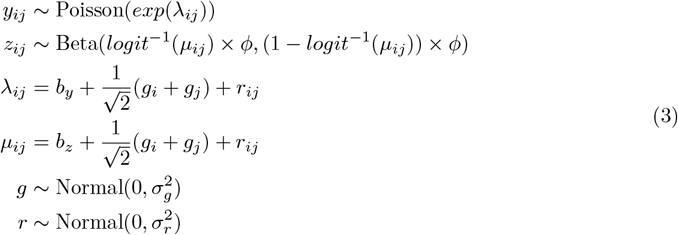

In this implementation, we estimate four key parameters from the two observed interaction networks. First, we estimate the variance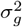for individual gregariousness, i.e., how much do individuals vary in their propensities to interact. Second, we estimate the variance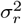 that reflects variation in dyadic affinities, i.e., dyad-specific propensities to interact. The third and fourth components, *b*_*y*_ and *b*_*z*_, are the baseline parameters for the two modeled behaviors. For example, *b*_*y*_ reflects the expected interactions for behavior *y* of two individuals of average individual gregariousness with an average pairwise affinity. Depending on which error distribution is used, we need to estimate additional parameters (in the example above we need to estimate a dispersion parameter, *ϕ*, for the Beta proportion).

In essence, the model we outlined is a multi-membership model with an additional varying intercept for dyad ID. The model is closely related to the social relations model (Koster et al., 2015, 2019, 2020), albeit with a focus on undirected data and the ability to fit multiple behaviors simultaneously.

After fitting the model to data, two main pieces of information can be obtained. First, we can extract the actual values for each individual’s gregariousness and each dyad’s affinity. These individual and dyadic values reflect how much a given individual or dyad differs from an average individual or dyad. In statistical terms, we estimate varying intercepts for individuals and dyads. The obtained individual and dyadic values might then be used downstream to construct networks based on dyadic connections that are not confounded by individual propensities.

Second, we can assess the relative magnitude of individual and dyadic contributions to variation in interactions. Note that in this context, ‘interactions’ is used in a generic sense, i.e., it could mean *duration, proportion, count, frequency*, etc. If the individual gregariousness variation is large relative to the variation in dyadic affinity, then one might conclude that individual propensities are more important than dyadic relationships in determining who interacts with whom (or how long) (scenario (a) in figure 2). In the extreme case of the dyadic component being near 0, one might even conclude that interaction patterns are determined by individual propensities alone. If, on the other hand, the variation in the individual component is near 0, while the variance in the dyadic component is large, we could conclude that individuals are practically identical with respect to their general propensity to interact and that any variation in observed interactions is largely due to dyadic, i.e., relationship, effects (scenario (b) in figure 2).

The model as outlined above makes three key assumptions. First, we assume that only one individual-level axis and one dyad-level axis explain interaction patterns, regardless of how many interaction types we observed. Second, both individual gregariousness and dyadic affinities are normally distributed. We use this as a convenient statistical shorthand to indicate that most individual and dyadic propensities are ‘average’, while fewer individuals/dyads have smaller and stronger propensities. Third, we also assume that the model with only gregariousness and affinity as predictors is correctly and sufficiently specified, i.e., there are no confounding variables. For example, an essential driver for dyadic interaction patterns and/or dyadic propensities to interact is relatedness, which in this implementation is ignored but the model could be adopted to incorporate such potential confounds (Kawam et al. (2024), see also supplement S3). Similarly, spatio-temporal associations may be driven by external factors, such as food resources (Whitehead & James, 2015). At the same time, the model does not require thresholding in the sense that there is a minimum number of observations or observation effort required (Farine & Whitehead, 2015; Schülke et al., 2022), which also removes researcher degrees of freedom since there are no universally accepted rules for thresholding.

### 2.1 bamoso R package

We provide a rudimentary R package to facilitate the application of the model to empirical data. The functions in the package fit the model in a Bayesian framework using the Stan engine for probabilistic modeling (Stan Development Team, 2024). A major advantage of using a Bayesian approach is that the model produces posterior distributions for all the quantities of interest, which allow for assessment of certainty in individual quantities or differences between quantities (e.g., has individual A a stronger connection to B than to C?, see Kajokaite et al. (2022) and Neumann & Fischer (2023) for examples).

In our package implementation, we used weakly informative priors for the individual gregariousness and dyadic affinity variation (50% of probability density for the two standard deviations falls between approximately 0.3 and 1.5, which still allows for smaller and larger estimates). For the baseline values for each behavior, we also used priors that were weakly informative. Here, however, the actual observed data inform how these priors were set exactly. This step is justified because we are estimating an intercept, and there is good reason to assume that the empirical median is a good approximation of the quantity we want to estimate (Bürkner, 2017).

## 3 Illustrative examples with simulated data

In the following, we provide two examples using simulated data to illustrate the intuition behind our model. In the first example, our goal is predominantly descriptive, i.e., we only estimate the latent sociality components, i.e., individual gregariousness and dyadic affinity. The second example illustrates an application where the ultimate goal is to estimate the relationship between individual sociality and an individual-level outcome, representing a template for an empirical study.

### 3.1 Example 1: two observed behaviors

We simulated interaction data for two interaction types (reflecting measuring a discrete and a continuous behavior) for a group of 15 individuals (see supplement S4 for more details). We created two scenarios. In the first, individual gregariousness variation was large compared to dyadic affinity variation (*σ*_*g*_ = 1.7, *σ*_*a*_ = 0.3). In the second scenario we set the variation in dyadic affinity large compared to gregariousness variation (*σ*_*a*_ = 1.7, *σ*_*g*_ = 0.3). Behavioral baselines were set to −1.7 for both behaviors in both scenarios.

For each scenario, we fitted our model to these interaction data (equation 2) using our `bamoso` R package. We then performed posterior predictive checks (Gabry et al., 2019), i.e., we looked at our model’s ability to predict interaction patterns that are similar to what we ‘observed’ (i.e., the interaction network that we simulated, see supplement S5 for illustrations). In both scenarios, the model was able to make predictions that matched the observed data very well (top two rows in figure 3). These plots show histograms and density plots of the observed interactions and overlays interactions predicted by the model. The model also recovered the input values for variation in individual gregariousness and dyadic affinity well (posterior medians, scenario 1: *σ*_*g*_ = 1.44, *σ*_*a*_ = 0.21; scenario 2: *σ*_*a*_ = 1.70, *σ*_*g*_ = 0.21).

**Figure 3:**
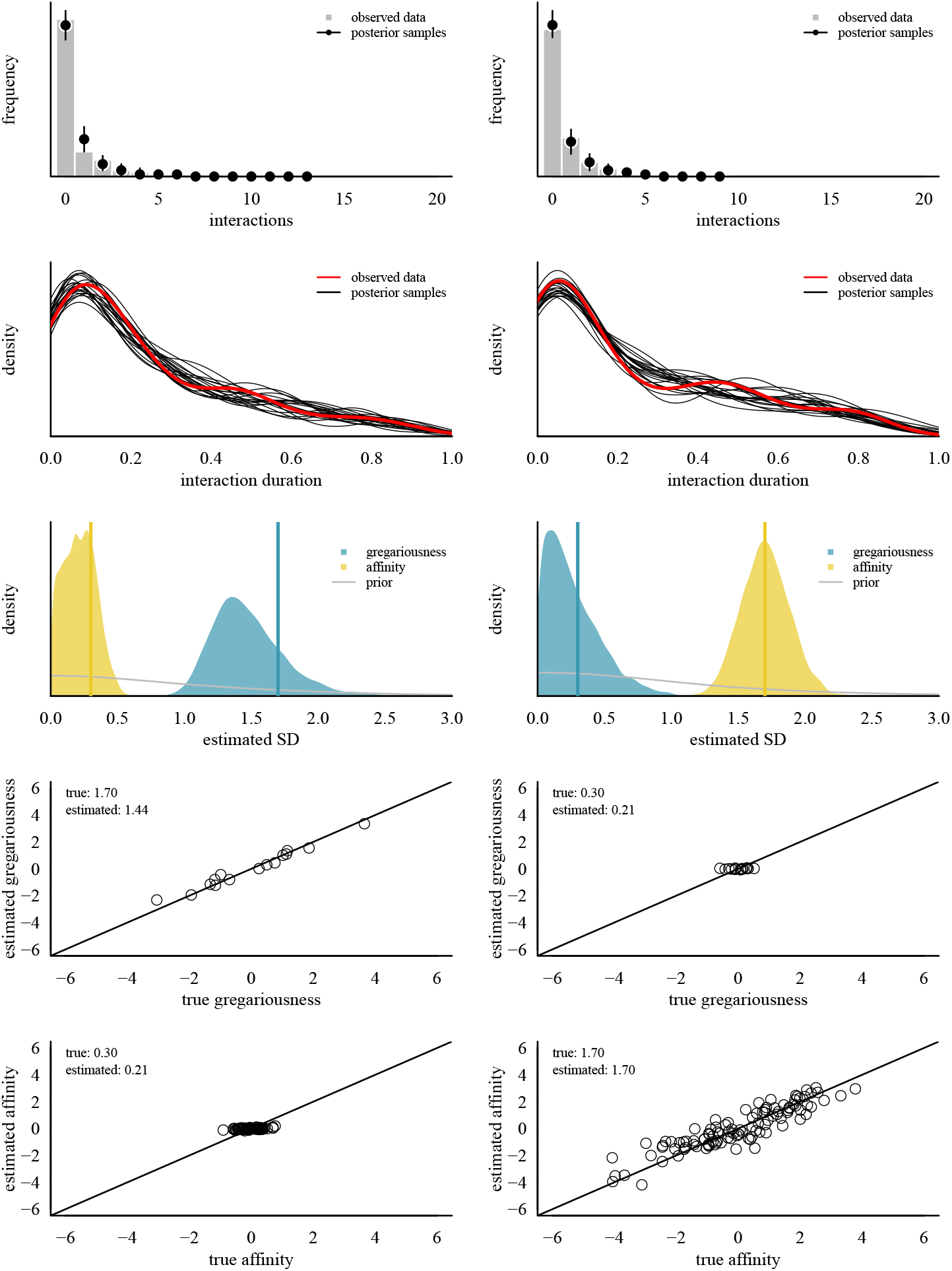
Model results of the two scenarios, each using two different interaction types (a frequency and a continuous proportion) to infer the underlying individual and dyadic sociality axes gregariousness and affinity. The two top rows show posterior predictive checks for both behaviors. The third row represents posterior distributions for the estimated variation in gregariousness and affinity where the vertical lines represent the true values. The two bottom rows show the relationship between true values and estimated values, i.e., posterior medians, for gregariousness and affinity.

As for recovering individual and dyadic values (two bottom rows in figure 3), we observed that if the underlying true variation was large, the model faithfully recovered the true individual and dyadic values. When the underlying variation was small, the model recovered the fact that the variation was small (small estimated SDs).

In sum, this small example serves as a proof of concept. It illustrates the underlying idea of our model: starting from two interaction networks, we decompose the underlying drivers of these interactions to arrive at estimates of individual, dyadic and group features. More systematic evaluations are presented below (section 4).

### 3.2 Example 2: from interactions to fitness

Imagine we want to study the relationship between individual gregariousness and an outcome related to fitness, like the number of offspring produced, longevity or survival (e.g., Dal Pesco et al., 2022; McFarland et al., 2017; Schülke et al., 2010; Wey & Blumstein, 2012). Typically, such a study requires at least two separate steps. First, the individual sociality variable is quantified from raw dyadic interaction data (in the cited examples, these were degree from a sociality index network, cumulative strength of the three strongest edges from a sociality index network, and principal component scores from network metrics). The second step then takes these values and uses them as a predictor variable in the statistical model, in which the response variable is the (fitness-related) outcome of interest. Although a common practice, this workflow has at least two related downsides. First, it assumes that step 1 produces values that genuinely reflect the construct of interest. Second, it completely ignores any uncertainty in the values reflecting sociality.

Here we provide a simulated example that simultaneously estimates gregariousness and feeds it directly into a single actual model of interest, carrying over uncertainty from estimating gregariousness into modeling the outcome of interest. In other words, we explicitly treated the quantification of sociality as integral part of our primary statistical model. In this context, it might be helpful to think of gregariousness as a personality feature and oppose it with the independent tendencies of individuals to form relationships of varying strength, i.e., dyadic affinity. To reiterate, gregariousness and dyadic affinity represent two distinct, and not necessarily correlated, mechanisms, which manifest in observable interactions (figure 2).

To achieve this, we set up a group of 25 individuals, in which each individual was assigned an individual value of sociality. These values represented each individual’s true gregariousness and were drawn from a normal distribution with mean = 0 and *σ*_*g*_ = 0.9. Then we assigned each dyad an affinity value (normally distributed with mean = 0 and *σ*_*r*_ = 2.3, *n* = 300 dyads). Biologically, this reflects a situation in which dyadic affinity is much more important than gregariousness in explaining interaction frequencies. With these values, we then generated interactions for each dyad (following the count component in equation 3), with a baseline rate of − 1.5, i.e., a dyad of two average individuals with an average affinity would be expected to interact approximately 0.2 times (*exp*(−1.5 + 0.5 *(0 + 0) + 0) = 0.22). Next, we generated a count variable representing our outcome for each individual (e.g., the number of offspring produced). These numbers were drawn from a Poisson distribution with a mean of *exp*(*c*_0_ + *c*_1_× *g*), where *g* is the individuals’ gregariousness values generated above and *c*_0_ and *c*_1_ are intercept and slope set to *c*_0_ = − 0.5 and *c*_1_ = 1.2, respectively. This resulted in a positive relationship between gregariousness and outcome, as illustrated in figure 4a. We were ultimately interested in this relationship, but crucially, we did not know the gregariousness predictor and needed to estimate it from the ‘observed’ interaction data.

**Figure 4:**
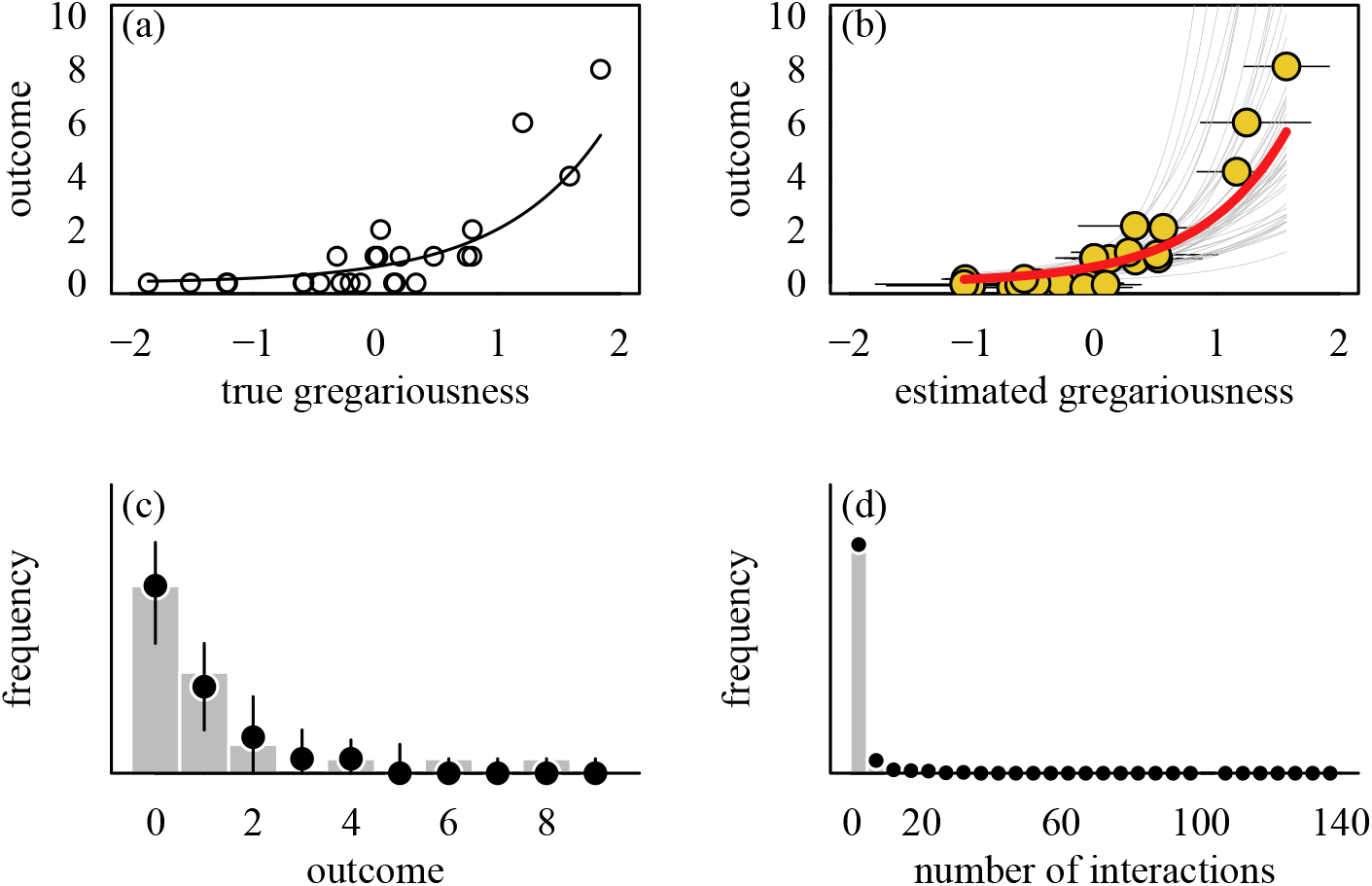
Estimating the relationship between gregariousness and an outcome using simulated data. Panel (a) shows the true underlying association between gregariousness and outcome, which in our case, is a relatively strong positive relationship. Panel (b) shows the results of our model as thick red line, which *estimated* gregariousness from interaction data. The panel shows as circles the posterior medians of estimated gregariousness and the horizontal lines represent the 50% credible intervals for each estimate. In red is the estimated model of the outcome (using the median posterior intercept and slope). The thin grey lines correspond to model predictions using 50 random draws. Panels (c) and (d) depict posterior predictive checks for the outcome variable and the interaction data (using 50 random draws). The filled bars represent histograms of the observed counts, while the circles and vertical bars represent the median of the 50 draws and their 89% quantiles.

To this aim, we set up the following generative model (equation 4, code is in supplement S2). In brief, we estimated the individual sociality component (gregariousness) and the dyadic sociality component (affinity) from the interaction data and in parallel used the estimated gregariousness (but not the dyadic affinity) as the predictor for modeling the outcome. To iterate, we used the estimated gregariousness because we were interested in understanding how an *individual* feature of sociality maps onto an *individual* feature of interest (our outcome). We assumed that we sampled the behavior with equal observation effort per individual and dyad.^1^

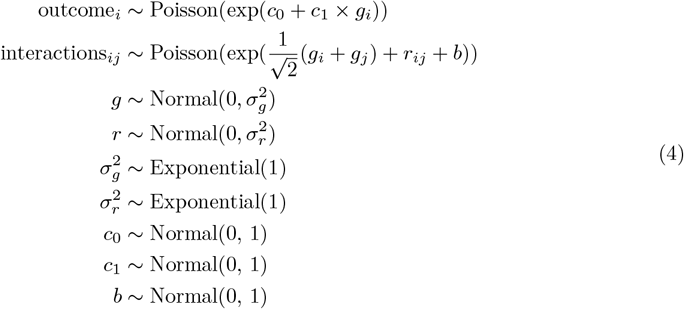

Fitting this model to our simulated interaction and outcome data produced estimates for all our quantities of interest. Foremost, the model recovered the true relationship between gregariousness and outcome fairly well (posterior medians; *c*_0_ = − 0.49, *c*_1_ = − 1.42, figure 4b) and was able to predict the observed outcomes faithfully overall (figure 4c). Similarly, also the parameters corresponding to modeling the interactions closely resembled their true values (posterior medians; *b* = −1.60, *σ*_*g*_ = 0.90, *σ*_*r*_ = 2.39). Thus, we were able to predict interaction patterns from the model that resemble the input interactions (figure 4d).

This simulated example illustrates how a model could be designed to answer questions about the association between individual sociality and an outcome variable of interest. It also demonstrates some of the key advantages of our approach: allowing for model checking on the level of the observed interactions (and also at the level of the outcome of interest), and it illustrates how uncertainty in the estimation of the sociality component is carried over into the actual model of interest.

Nevertheless, the model we fitted here could be improved. Importantly, we did not run prior predictive checks (Gabry et al., 2019), which likely would have led to setting more reasonable priors and hence more solid inference given that our model, while overall recovering the true parameters reasonably well, also resulted in draws that made seemingly implausible predictions.^2^

## 4 Evaluation

We created artificial data sets to evaluate the model’s performance across a range of biologically plausible scenarios. This analysis extends the illustrative example we presented above (figure 3). In particular, we varied individual gregariousness variation (SD ranging continuously between 0.1 and 3), dyadic affinity variation (SD ranging continuously between 0.1 and 3), and behavioral baseline (ranging continuously between -3 and 1). We also varied group size (number of individuals: 5 through 30), the number of behaviors (one or two), and the distribution type of the behavior (frequency, discrete proportion, continuous proportion, and duration). We created a total of 2,000 data sets.

We first visually evaluated the relationship across data sets in how well the model recovered the variation in the underlying process (i.e., the relationship between estimated variation and ground truth). We also evaluated for each simulated data set the correlation between true and estimated individual gregariousness values and the correlation between true and estimated dyadic affinity values.

Our model recovered ground truth well for both components: when the simulated variation (i.e., ground truth) was small, the estimated variation was also small, and vice versa (supplement S6). Notably, for the gregariousness component, our model sometimes underestimated the true variation which was primarily prevalent in smaller groups (supplement S6). This underestimation is likely a consequence of having not enough information in the data to pull the estimate away from the prior and the prior having most of its density near 0 (see figure 3). Finally, we also found that the correlations between the true and estimated individual and dyadic propensities were generally positive and approached 1 when the true values became larger (supplement S6).

## 5 Applied examples

In the following section, we illustrate the application of the framework with two real-world examples. The first example descriptively compares the variation in the two sociality axes in groups of four primate species. The second example illustrates a ‘complete’ workflow where sociality is used as a predictor in a model of reproductive success in kangaroos.

### 5.1 Grooming in macaques

To illustrate a potential application, we analyzed grooming networks of four social groups belonging to four macaque species (*Macaca* sp.). The genus is widely used in comparative research in behavioral ecology (e.g., Balasubramaniam et al., 2012; De Moor et al., 2025; Neumann & Fischer, 2023; Thierry et al., 2008). Here, we chose four species representing the diversity of macaque social styles (Thierry, 2007) to describe how much gregariousness and dyadic affinity vary across the different species and possibly in relation to their social style. For each species, we obtained data for one group. All but one of the groups (*sylvanus*) were observed in the wild, using focal animal sampling. The interaction data were undirected, and grooming was recorded as frequencies (duration for *nigra*). Group sizes ranged between 15 and 20 individuals, all of which were adult females.

We found substantial variation between the groups in the absolute and relative magnitudes of estimated individual gregariousness and dyadic affinity variation (figures 5a-d and supplement S7). In two groups (*assamensis* and *nigra*), affinity variation was smaller than gregariousness variation (figure 5a-d). In the *nigra* group, the estimate for the dyadic affinity component was so small that it could be argued that it appears to play only a minor role, if any, in explaining interaction patterns. In other words, the results suggest that for this group, grooming patterns were primarily driven by individual differences in grooming propensities. Across species, we found no obvious pattern. However, our results are tentatively more compatible with the notion that variation in both gregariousness and affinity is larger in more despotic species (*fuscata* and *assamensis*, supplement S7).

**Figure 5:**
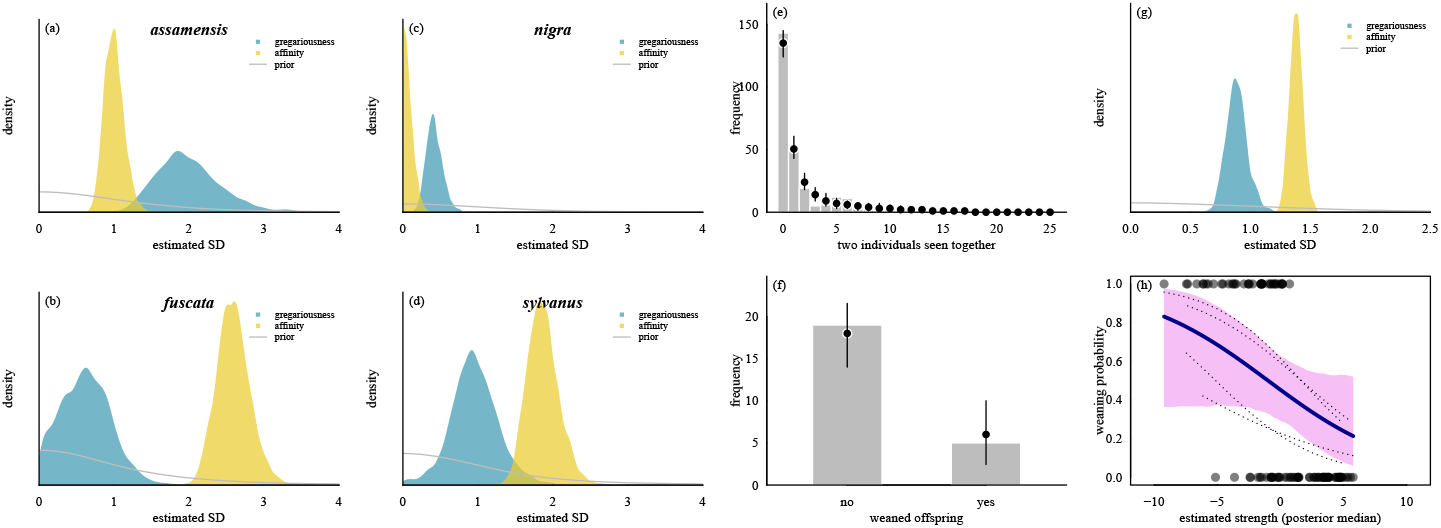
Illustrating empirical use cases of our suggested modeling framework. (a)-(d): Variation in individual and dyadic propensities to groom in four groups of macaques. Displayed are posterior distributions of individual gregariousness and dyadic affinity parameters. The four underlying interaction networks were undirected grooming networks (frequencies or durations) in groups of 15 to 20 adult females. (e)-(h): Reanalyzing Menz et al. 2020. Panels (e) and (f) show posterior predictive checks for association patterns (how often dyads were seen together) and the outcome for the reproductive success measure (number of females that weaned successfully) from 50 draws. The bars in each panel correspond to the observed frequencies. The circles and the vertical lines represent the median value predicted by the model alongside the range of predicted values from the 50 draws. These checks only reflect year 1. Panel (g) shows the posterior distributions of the estimated variation in the two sociality components. Panel (h) shows the model predictions for weaning using estimated strength from association data (plus 89% credible interval). Dotted lines are predictions for the four different years.

In the future, it would be desirable to incorporate a more diverse species data set, with possibly more than one group per species, to understand how much sociality components vary both *within* and *between* species (De Moor et al., 2025).

### 5.2 Reproductive success in kangaroos

In this example, we reanalyzed a data set from Carter and colleagues (Carter et al., 2020; Menz et al., 2020) from a study that found negative relationships between social features and several measures of reproductive success in grey kangaroos (*Macropus giganteus*). The primary goal here is to illustrate how our framework can be extended using observed interaction data to quantify gregariousness and dyadic affinity. Specifically, we aimed for an analysis that uses raw behavioral data (data on spatio-temporal associations in this example) to estimate the actual model of interest, i.e., the relationship between individual sociality and reproductive success. In this way, we avoid doing this analysis in two separate steps, allowing us to carry over the uncertainty from estimating the two sociality components into our model of interest. This example can be considered an extension of the simulated example in section 3.2 above (figure 4) with empirical, as opposed to simulated, data.

The authors of the original study constructed networks from association data using the Half-Weight Index (HWI: Cairns & Schwager, 1987). The authors extracted two metrics from these networks that represented individual sociality: strength and the local cluster coefficient. These two metrics then served as predictor variables in models of three measures of reproductive success (see Menz et al. (2020) for details).

Implementing this approach in our framework came with two key challenges. The authors calculated derived metrics from HWI networks: strength and clustering. To follow this aspect of the original analysis, we had to code the algorithms to obtain strength and clustering in Stan (see supplement S8). The second challenge was to code up the entire remaining model in Stan.

In addition, we needed to account for the fact that data came from individuals across multiple sampling periods (years) that only partly overlapped in composition, i.e., some individuals were represented in more than one period, and some only occurred in one period. Since this is not a direct replication attempt and only serves as an illustration for a real-world application, we made additional changes to the model design. First, we used a subset of 47 individuals (around one-third of the original sample size) to speed up model fitting. We also allowed the slopes for the two network measures (strength and clustering) to vary between the four years. Finally, we analyzed only one outcome variable (weaning success), instead of all three. Code for this analysis is in the supplementary files (repl_06_kangaroos.R).

The results show that our model successfully predicted the association patterns and the eventual outcome variable (did the female successfully wean an infant in a given year) (figure 5e and f). Furthermore, looking at the magnitude of variation in the two sociality components, we found that variation in both was substantially larger than 0 and that the dyadic affinity component was slightly larger than the individual sociality component (affinity: median = 1.38, 89%CI: 1.29 − 1.47; individual: median = 0.88, 89%CI: 0.74 - 1.04, figure 5g). Finally, the effect of individual network strength on weaning probability our model recovered was strongly negative and therefore agreed with the original findings of Menz et al. (2020): Females with higher strengths were less likely to wean successfully (figure 5h).

## 6 Discussion

In this study, we provided a novel framework for using dyadic interaction data to obtain estimates for individual-specific propensities (gregariousness) and dyad-specific propensities (affinity) to interact. We argued that observed interactions are outcomes driven by the combination of these two processes. Estimating them both simultaneously from a generative model has distinct advantages. First, the parameters the model estimates have clear interpretations and allow checking whether we can predict our observed interaction patterns. Such a technique can serve as a sanity check on model validity, a step that should be incorporated in current approaches. Second, since we actually *estimate* the two sociality components, we can grasp the uncertainty of the estimated sociality values for individuals and dyads.

Our proposed framework fits nicely with recent advances and the current advent of Bayesian methods in the analysis of social interaction networks (e.g., Ross et al. (2024), Hart et al. (2023), Farine & Strandburg-Peshkin (2015), Neumann & Fischer (2023)). Table 1 provides a summary of key features and differences among current frameworks.

**Table 1:** Comparison of three frameworks and packages to study interaction networks.

|  | bamoso | BISoN | STRAND |
| --- | --- | --- | --- |
| predict/simulate interactions on the level of behavior | yes | no | not documented |
| support for more than one behavior (multilayer networks) | yes | no | not documented |
| estimate edge weights free of individual component | yes | no | yes |
| generative formulation | yes | no | yes |
| framework for downstream models | manual | via **brms** | yes |
| focus on interaction networks that are | undirected | undirected | directed |
| goal | descriptive | inference | inference |
| goal is quantification of dyadic relationships | yes | yes via network metrics | no |
| allow missing values in networks | yes | no | not documented |
| impute missing values in networks | no | no | no |
| additional predictors | rudimentary | yes | yes |
| has hex sticker | no | yes | yes |

However, estimating the sociality components is rarely the ultimate goal in empirical questions (but see our comparative example above). Instead, we typically use sociality estimates as predictors in further downstream analyses to study variation in biologically relevant traits, such as reproductive performance, survival, and disease or knowledge transmission. We illustrated how this goal could be achieved while propagating the uncertainty in the sociality estimation to the actual model of interest (i.e., the model that links some particular sociality axis to an outcome of interest) instead of treating sociality values as point estimates.

We have applied our framework in one empirical study thus far in which we addressed questions about meat transfers in a population of Guinea baboons, *Papio papio* (O’Hearn et al., 2024). Here we used estimated gregariousness to predict the audience size of subjects during meat eating events and estimated dyadic affinity to predict audience composition. Conditional on the composition of the audience, we used the dyadic affinities to predict who in the audience obtained meat from the possessor. The results in O’Hearn et al. (2024) showed that gregariousness was positively associated with audience size and the higher the dyadic affinity the higher the probability of an individual to be in the audience and, conditional on being in the audience, to obtain meat. Overall these results demonstrate that our framework can be applied to empirical questions in a meaningful way.

The study of sociality, how and why individuals interact and associate, has a long history (Hinde, 1976; Ostner & Schülke, 2018). In this paper, we were not directly concerned with the theoretical framework of social relationships per se, i.e., we ignored critical dimensions of relationships like predictability, stability, symmetry, diversity, and tenor (Cords & Aureli, 2000; Fischer et al., 2017; Hinde, 1976; Ostner & Schülke, 2018; Silk et al., 2013). Our primary focus was on understanding the strength of relationships in an affiliative context. We argued that we can use observed dyadic interactions to estimate contributions from the group (baseline), the dyad, and the individual to relationship strength. This decomposition is essential because when studies describe the fitness consequences of sociality, this is necessarily done on an individual level: *individuals* survive longer or produce more offspring (Snyder-Mackler et al., 2020). But it remains unclear what exactly is the underlying mechanism if quantities like ‘stronger relationships’ or ‘higher centrality’ are used to predict fitness outcomes when it is unknown how differentiated dyadic affinities are in the first place. A more parsimonious explanation is that if interaction patterns are entirely driven by gregariousness, such correlations *could be* misleading. Honing in on the mechanisms thus requires quantifying the underlying individual and dyadic components. Only after that we can look meaningfully into individual-level summaries of sociality (e.g., ‘stronger relationships’ with the gregariousness component removed).

An issue that exacerbates the potential for generating misleading results is the nature of how dyad-level data are broken down to the individual level (often termed ‘social integration’, Snyder-Mackler et al. (2020)). How social integration has been quantified from dyadic interaction or association data varies substantially between studies, undermining meaningful comparisons across studies (Ellis et al., 2019; Ostner & Schülke, 2018; Schülke et al., 2022; Snyder-Mackler et al., 2020). One of the essential critical advantages of our approach is that we provide a principled workflow that eliminates at least some of the researcher degrees of freedom when estimating social integration. To be explicit, we are not saying that previous approaches necessarily led to wrong conclusions. However, the model we suggest is tractable and interpretable because of its generative nature. It should allow us to arrive at more readily comparable results across studies.

One crucial issue to keep in mind is the overall question when analyzing interaction patterns. There are scenarios where the actual interaction frequency or duration may be more informative than estimating the underlying gregariousness and affinity. If the goal is to study, for example, disease transmission or transmission of cultural techniques or knowledge, then it might not be relevant to partial out the individual component of network connections (Farine & Whitehead, 2015; Godde et al., 2013). If a dyad interacts very often, then this might facilitate *transmission* irrespective of whether the interaction frequency is due to individual or dyadic processes. But even then, it might be relevant to quantify the underlying processes because more gregarious individuals might be more susceptible to carry the disease or possess information in the first place. And more generally, it seems advisable to actually *model* interaction frequencies instead of using point estimates as truth.

One drawback of our framework is that it likely requires coding up the models in Stan when going beyond relatively simple descriptions of the two sociality axes. It seems unlikely that it is possible to develop a software package that simplifies the implementation and does justice to all the possible combinations and peculiarities of all the different study systems and designs. For example, our package provides only functions to estimate the sociality components but does not allow fitting more complex models in a formula-syntax approach. We envision our R package as a descriptive and pedagogical tool that helps researchers describe and understand interaction patterns, before moving on to arguably more complex models fitted either by hand or some other package. However, we are convinced the effort to code up models like the ones we presented is well worth it because of the advantages we can gain from the insights such models can generate (e.g., O’Hearn et al., 2024).

In this light, we see a number of ways to extend our modeling framework, which we partly already incorporated in the applied examples above. First, the model should allow incorporating covariates at the level of the individual gregariousness and dyadic affinity levels (see Kawam et al. (2024), Koster (2020), supplement S3). Second, future extensions could handle directed behavior (given versus received, Ross et al. (2024), Koster (2020)). Third, if data were recorded over multiple sampling periods, this could be accounted for with varying estimates for period (see the kangaroo example). Fourth, actual network metrics could be included, if required, like clustering, centrality, or betweenness (kangaroo example, Hart et al. (2023)). Fifth, more than one gregariousness or affinity axis may exist, and if they do, might correlate and it is also possible that the individual and dyadic components themselves correlate; for example, individuals with higher individual gregariousness also form stronger relationships. All these suggestions seem worthwhile to implement because of their potential to further extend our understanding of the building blocks of social relationships.

## Conclusion

Understanding interaction patterns is vital to understand relationships. This study presents a principled modeling framework to estimate the underlying latent sociality components from interaction data on the individual and the dyad level. We provided simulated and empirical examples to illustrate and evaluate its application. This workflow provides a promising extension of our toolbox to understand how and why individuals interact and to investigate the consequences of variation in the different sociality components on other phenomena of interest.

## Supporting information

bamoso R package

readme

replication R code files

package tutorial

## Acknowledgments

We thank Roger Mundry and Julie Duboscq for discussions and comments on earlier drafts. Two reviewers and the editorial staff provided very helpful and constructive comments. We thank Julia Ostner and Oliver Schülke for providing data on Assamese macaques from Phu Khieo Wildlife Sanctuary, and we thank Andrew MacIntosh and Julie Duboscq for providing data on Japanese and crested macaques, respectively. This publication was funded by the Deutsche Forschungsgemeinschaft (DFG, German Research Foundation) - Project-ID 454648639 - SFB 1528 “Cognition of Interaction” - Project B05, and Grant/Award Number: 254142454 / GRK 2070 (“Understanding Social Relationships”).

## Supplements

### S1 Uncertainty

In this section, we illustrate how varying sampling effort results in varying degrees of uncertainty in assessing the different components in our model. Imagine two researchers observing the same group of animals during different periods. For the sake of the example, we assume that the underlying behavior of the subjects is the same, as is group composition. The only difference between the two observation periods is the time spent observing the animals. Intuitively, the more time we spend observing, the more precise our estimates of behavioral rates should become. We illustrate this here for a simulated example in which we observed a countable behavior. The only difference in the data generation was the time we allowed individuals to interact (statistically speaking, we varied the offset term). In one case, we observed the group for, say, two hours (which represents low observation effort in our example). In the second case, 100 hours were spent observing the animals (high observation effort). We set the baseline rate of the behavior to −0.7. Since we simulated a count variable, we expected an average pair of two average individuals to interact *exp*(−0.7) = 0.5 times per hour, which amounts to approximately 1 expected interaction (*exp*(−0.7 + *log*(2))) with low observation effort and approximately 50 interactions (*exp*(−0.7 + *log*(100))) for high observation effort. The difference is that observing 50 interactions in 100 hours should leave us much more confident that the true rate is about 0.5 interactions per hour than observing 1 interaction in 2 hours.

Figure S1 visualizes this intuition. We fitted our model to data generated with the above-mentioned settings. Across all parameters shown in figure S1, the posteriors are narrower in the scenario with higher observation effort.

**Figure S1:**
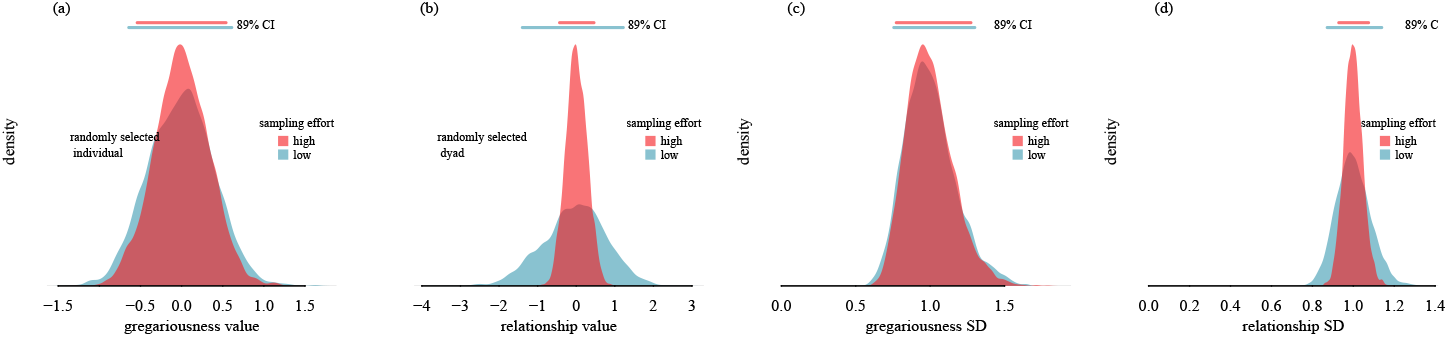
Uncertainty in gregariousness and relationship estimation. Observed behavior was a count variable. (a) and (b) show posteriors for one randomly selected individual and dyad, respectively. (c) and (d) show the posteriors for the estimated group-level variation for gregariousness and relationships. Within each panel, the posteriors were centered to aid visual comparison ((a) and (b) were centered at 0, (c) and (d) were centered at 1).

### S2 Stan code

There are three ways to obtain the raw Stan code for the models presented in this manuscript. The easiest way is to browse to [https://github.com/username/anonymous/inst/extdata], where all model code is accessible.

Raw Stan files can also be found in the source code of the accompanying R package `anonymous`. Unzip the file `anonymous_0.5.1.tar.gz` (which is part of the supplements). The Stan code files are in the directory `inst/extdata`.

The Stan code is also accessible via the `anonymous` package itself. This requires that the package has been successfully installed (see supplementary file `repl_00_readme.md` for instructions).

- general model (used in figures 3 and 5a-d)

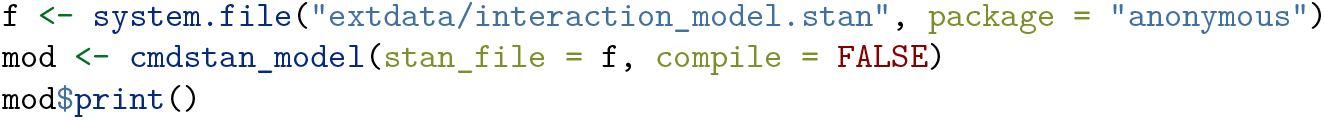
- workflow example (used in figure 4)

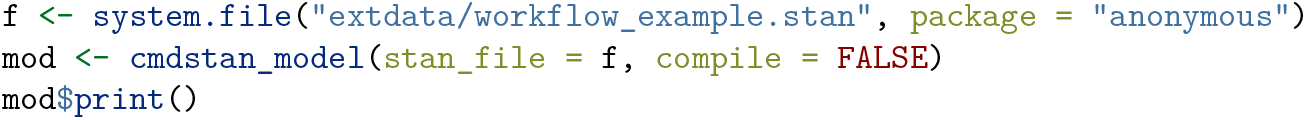
- kangaroo example (used in figure 5e-h)

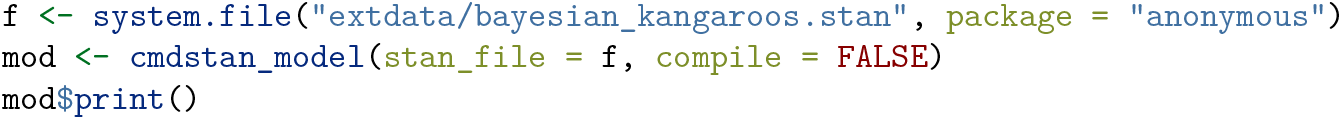

### S3 Models with covariates

This document is a slightly edited version of a vignette that is part of the `anonymous` package (https://github.com/username/anonymous/).

Here we illustrate how the basic sociality model can be extended. We start with the same premise as before: Interactions arise because of variation in the individual gregariousness and dyadic affinity. But now let’s consider the possibility that dyadic affinities are stronger between related individuals. Further, let’s consider the possibility that there are sex differences in individual gregariousness. Both seem like reasonable and plausible features of a social system. The slight twist is that we do not talk about potential sex differences in *interaction rates*. Instead, we talk about potential sex differences in *propensities* to interact. Of course, the same applies to relatedness and affinity: it’s about propensities to interact, not actual interactions. Figure S2 visualizes the idea.

**Figure S2:**
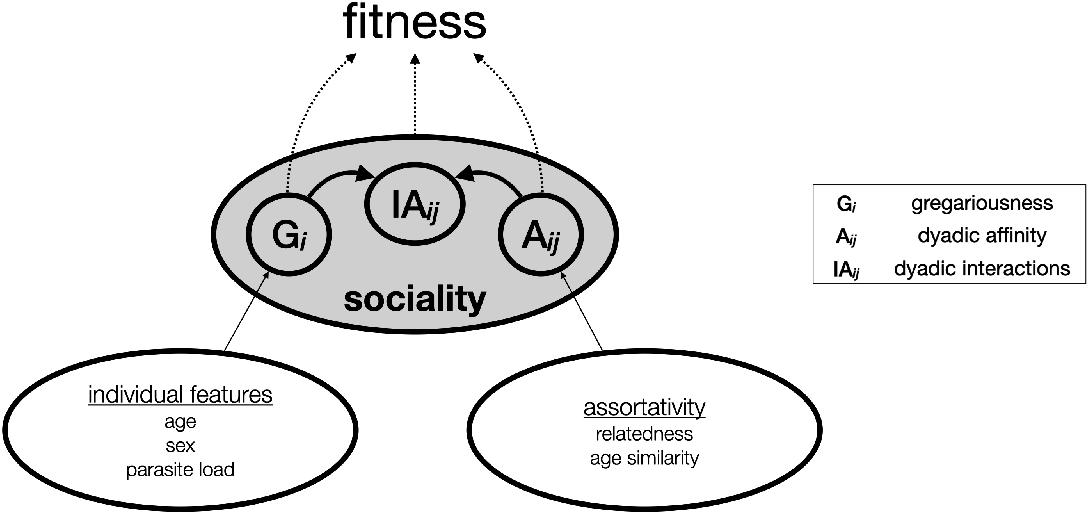
A simple flow chart illustrating the idea how covariates might affect individual and dyadic propensities to interact.

We can also formalize this. And that’s the advantage of taking the generative approach. In the framework of simple network metrics (I’m thinking of SRI or HWI or DSI) this is just not possible and we rely on ad-hoc solutions with potentially sever shortcomings (for some background see Hart et al. (2022)): SRI, DSI and friends are non-parametric indices and therefore can’t be incorporated in generative simulations. We can do better, I like to think (McElreath, 2020). So here is an attempt to a formal generative model:

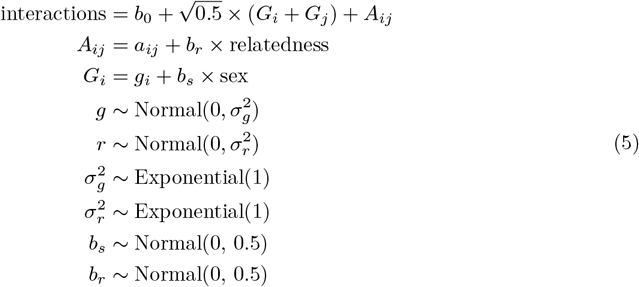

In this equation we just add slope parameters (*b*_*r*_ and *b*_*s*_) for the effects of relatedness and sex. Note how *b*_*r*_ only affects affinity and *b*_*s*_ only affects gregariousness (figure S2). Everything else is the same as before. In the future I might also translate this model into a DAG, which might also prove helpful, but not just now (and please don’t consider figure S2 as a formal DAG).

Before we get to a worked example: this is still a fairly simple model, and as such it can be fitted with the `anonymous` package. Naturally, we can envision models much more complex than this.

At this point it becomes hard to provide a universal syntax to fit such models without actually doing some hand-coding in Stan. But for now, let’s stay with this scenario where we have one dyad-level predictor and one individual-level predictor.

### S3.1 An example with sharks

The example data set that we use concerns reef sharks (*Carcharhinus melanopterus*) and the data come from (Mourier, 2020; Mourier & Planes; 2021). The only reason for using this data set is that it is the first that I found with the following features:

- raw interaction or association data
- a dyad-level predictor (here relatedness)
- an individual-level predictor (here sex)

The purpose of this example is primarily to illustrate the feasibility of integrating covariates. I won’t discuss the biological relevance of any results. So, let’s load the data:

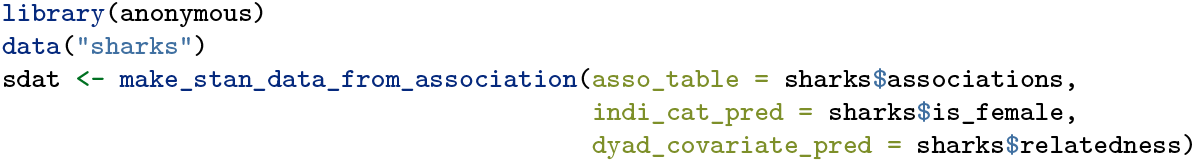

And fit the model (which takes about two minutes on my laptop):

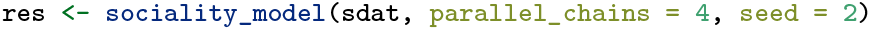

Let’s start by looking at some diagnostics plots (figure S3).

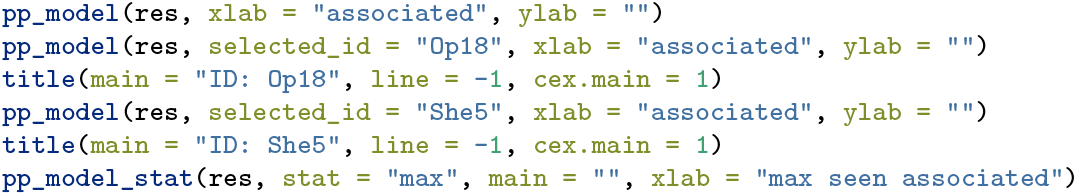

**Figure S3:**
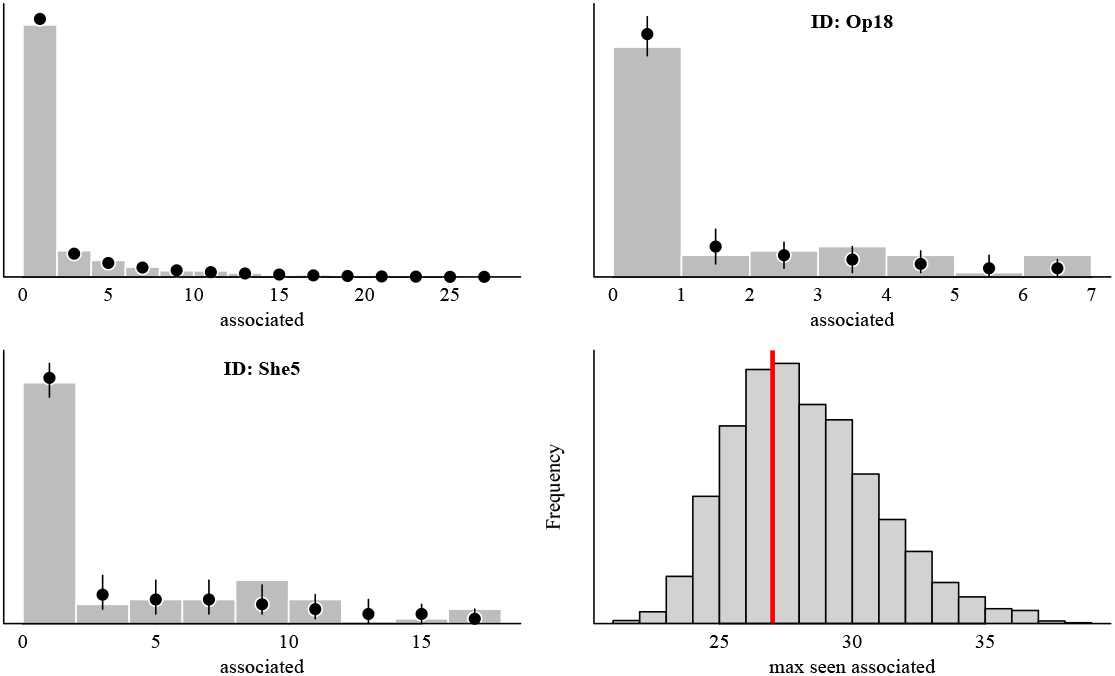
Posterior predictive checks for the shark model.

Now let’s extract the posteriors of the *b*_*s*_ and *b*_*r*_ parameters. This is done by accessing the results object’s internal components but will in the future be possible via `extract_samples()`.

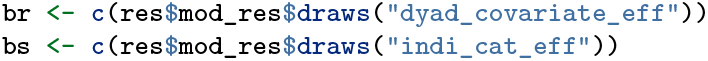

Now let’s plot the posteriors:

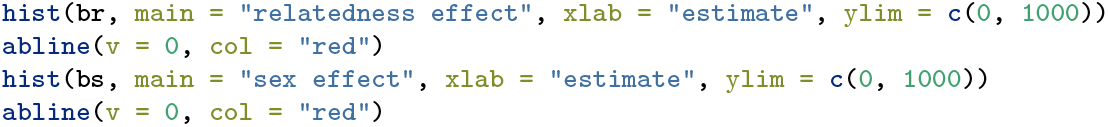

**Figure S4:**
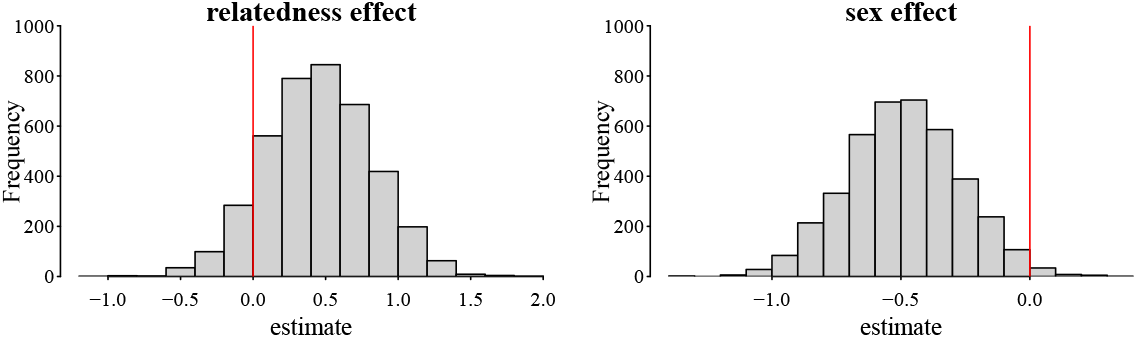
Association between relatedness and propensities to interact with specific partners (dyadic affinity) and association between sex and propensity to interact with anyone (gregariousness). Negative values for the sex parameters indicate the females have lower values than males.

It appears that most posterior mass for the relatedness falls in the positive range (posterior median: 0.45, proportion of samples larger than 0: 0.89, figure S4). It also appears that the sex effect is pretty reliably negative, i.e., females have lower gregariousness values than males (posterior median: −0.49, proportion of samples smaller than 0: 0.99).

### S4 Data generation and simulated examples

#### One behavior observed

We created four scenarios that differed with respect to how individual propensities vary relative to dyadic propensities while keeping the group-level baseline constant.

- **scenario 1)** Both individual propensities and dyadic propensities varied little, which reflects a system in which interaction patterns vary stochastically around the group baseline. In other words, all individuals had practically the same propensities to interact (i.e., gregariousness), and all pairwise affinities were practically the same.
- **scenario 2)** Individual propensities varied substantially relative to dyadic propensities. This scenario reflects a system in which differentiated interaction patterns emerge, which are predominantly driven by individual differences and where dyad-specific propensities are practically absent and do not contribute to interaction patterns (although looking at interaction patterns only, this might not become apparent, see figure 2a).
- **scenario 3)** Individual propensities varied little relative to dyadic propensities. This situation reflects a system in which individuals do not differ from each other, while across dyads, pairwise affinity varies from having very weak to very strong affinities (see figure 2b).
- **scenario 4)** Both components varied substantially, which reflects a system in which both individual and dyadic propensities drive interaction patterns.

In each scenario, we simulated a group of 20 individuals (190 dyads) and set up a vector of individual propensities and a vector of dyadic propensities. Each of these vectors was drawn from a normal distribution with a mean of 0 and a standard deviation of 0.15 or 1.7, depending on whether the component varied a little or substantially in a given scenario. We then generated a count outcome from a Poisson distribution for each dyad using the equation above with a baseline of 2.5, which was the same across all scenarios. This means that if two individuals each had an individual propensity of 0 and their dyadic affinity was 0 as well, we expected them to interact about 12 times^3^ 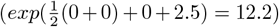. As in all our examples, these data are undirected.

Before applying our model, we first show the emerging interaction patterns under the different relative magnitudes of individual to dyadic contributions to interactions (figure S5). In the first scenario (with small individual and dyad variation), the emerging dyadic interaction numbers were all fairly similar and clustered around the intercept. The second and third scenarios led to fairly similar interaction patterns even though the underlying mechanisms were different. In scenario 2, interactions were driven predominantly by individual differences, while in scenario 3, interactions were driven by differences between dyads (see also figure 2). Scenario 4 also showed a differentiated interaction pattern although here both individual and dyadic components contributed to its emergence.

Next, we turned the question around, i.e., if we only know the observed interaction patterns, can our model recover and separate the underlying individual and dyadic contributions? To this end, we fitted our model to each of the four data sets created by our four scenarios (figure S6). First, we looked at our model’s ability to predict interaction patterns that are similar to what we generated above, i.e., we performed posterior predictive checks (Gabry et al., 2019). In all four scenarios, the model was able to make predictions that matched the observed data very well (top row in figure S6). These plots show the densities of the observed interactions of 190 dyads and overlays 20 densities of interactions predicted by the model. As for recovering individual and dyadic values, we observed that in case the underlying true variation was large, the model recovered the true individual and dyadic values faithfully. When the underlying variation was small, the model recovered the fact that the variation was small (small estimated SDs).

**Figure S5:**
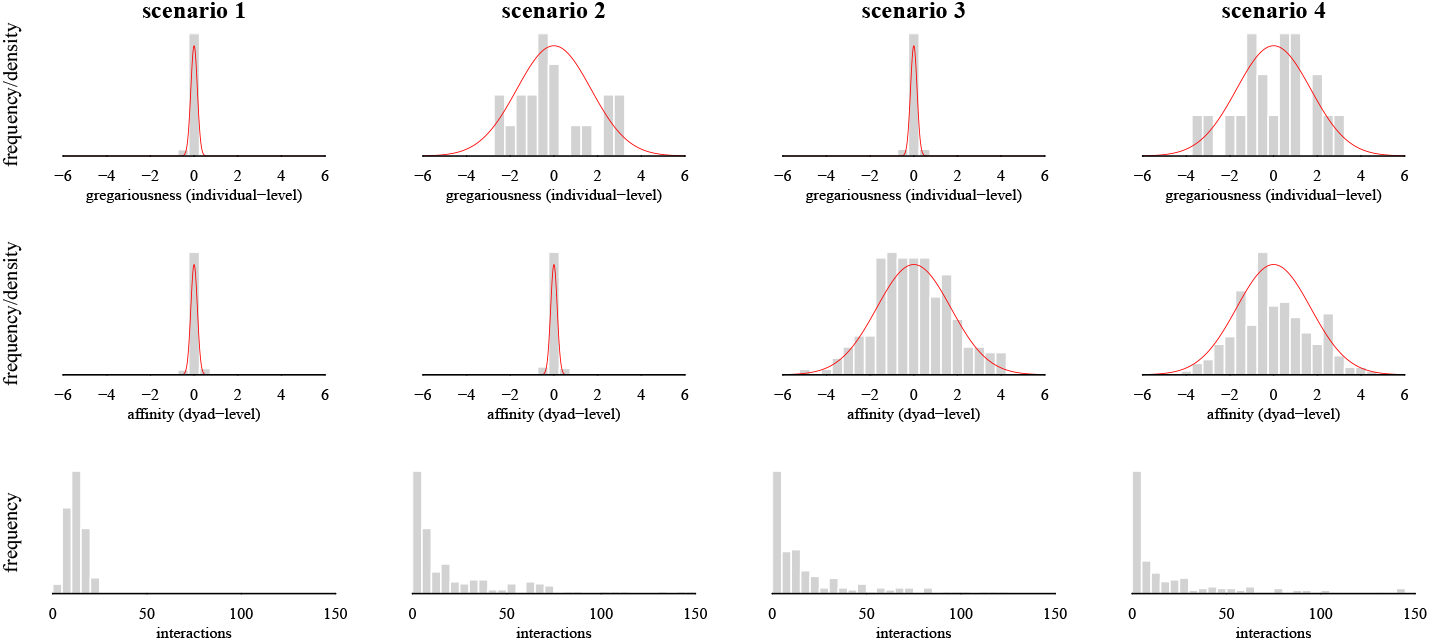
Emerging dyadic interaction patterns in groups of 20 individuals forming 190 dyads. Scenarios differed by combining low and high variability for individual and dyadic interaction propensities. The top row shows as histogram the generated individual gregariousness values. The red density comes from a normal distribution with the same parameters as the one the values in the histogram were drawn from. The middle row shows the corresponding data for dyadic affinity values across the four scenarios. The bottom row shows as histogram the emergent interactions given the simulated individual gregariousness and dyadic affinity values.

#### Two behaviors observed

The second example illustrates a case when data from more than one behavior, for example, grooming and co-feeding, should be used to estimate the underlying individual and dyadic sociality components. A widely used index to quantify dyadic sociality, the composite dyadic sociality index (Silk et al., 2006, 2013), attempts precisely that but does not account for individual sociality. The core idea here is that we simultaneously fit our model to two behavioral response distributions and estimate a single (shared) individual and a single (shared) dyadic component while estimating two separate baselines for the two different behaviors. We created two scenarios where we first generated two behaviors resembling a count and a duration (figure S7). In the first scenario, dyads varied a lot while individuals varied little (similar to scenario 2 above). In the second scenario, individuals varied little, but dyads varied greatly (similar to scenario 3 above).

One additional central assumption that the model makes when applied in this way (i.e., with more than one modeled behavior) is that there is indeed a single underlying individual component and a single underlying dyadic component. This assumption is made implicitly, for example, by the widely used composite sociality index (Silk et al., 2006, 2013). Whether this assumption holds can be investigated by visually examining the model’s predictions for the two behaviors. If the underlying processes differ substantially, the predictions will deviate from the observed data for either or both behaviors. In the case of our example, the predictions match the observed data well for both behaviors in both scenarios (figure S7, top two rows). Likewise, the estimated individual and dyadic values were recovered well for the cases where the variation in the given component was considerable. It must be noted that poor posterior predictive checks might also be due to other reasons, and conversely good posterior predictive checks alone are not unequivocal evidence for the validity of the assumption.

**Figure S6:**
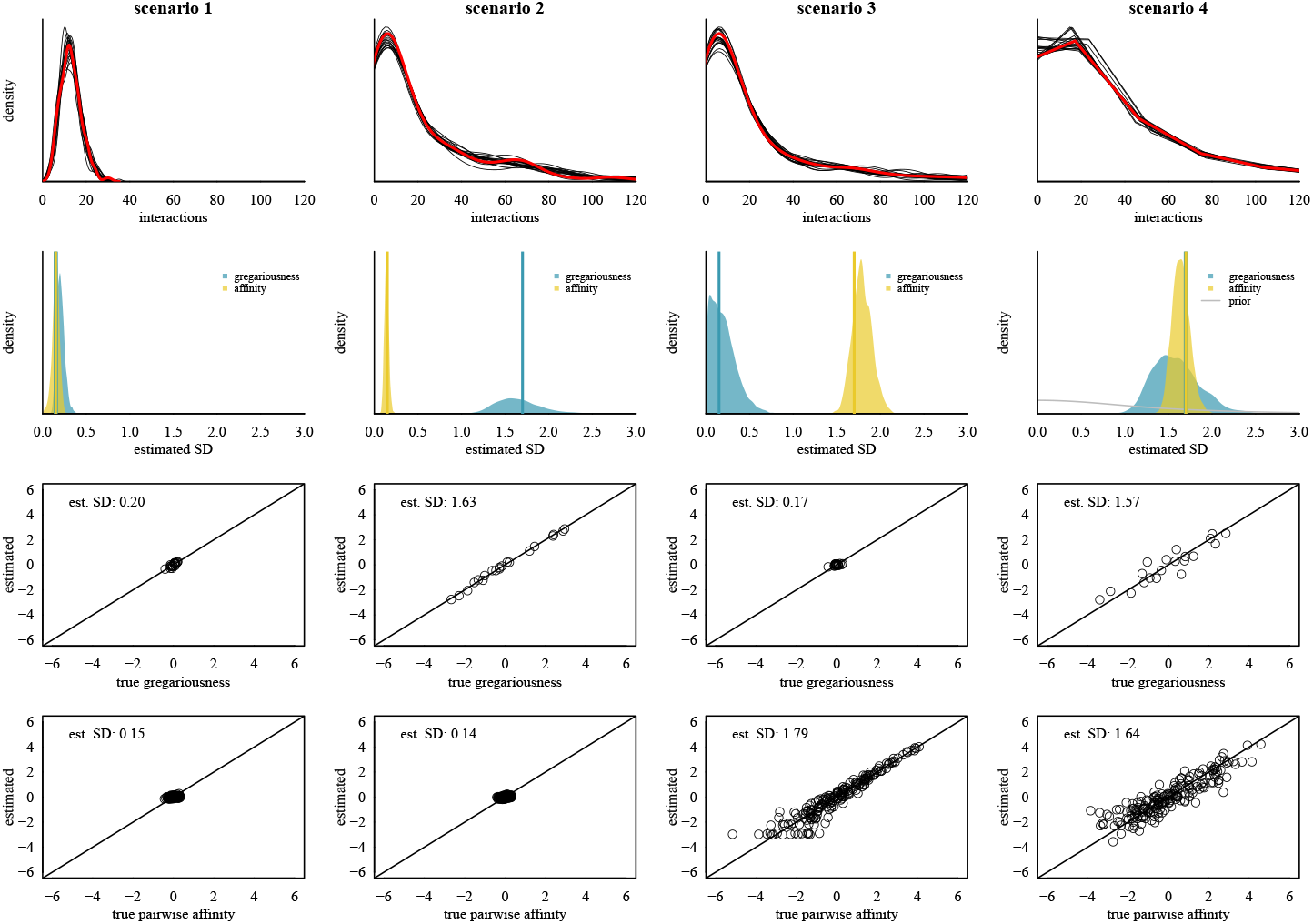
Model results of the four scenarios. In the top row are predicted patterns from the model (red is density of the observed data and black are predicted densities from 20 draws). In the second row are posterior distributions for the SDs of individual gregariousness and dyadic affinity. Below in the two bottom rows are relationships between true (as per simulation) and estimated (posterior mean) individual gregariousness and dyadic affinity values. The inset numbers are the posterior medians of the estimated sociality values (originally set to 0.15 and 1.7). The estimated intercepts for the four scenarios were 2.53, 2.51, 2.47 and 2.52 (originally set to 2.5).

**Figure S7:**
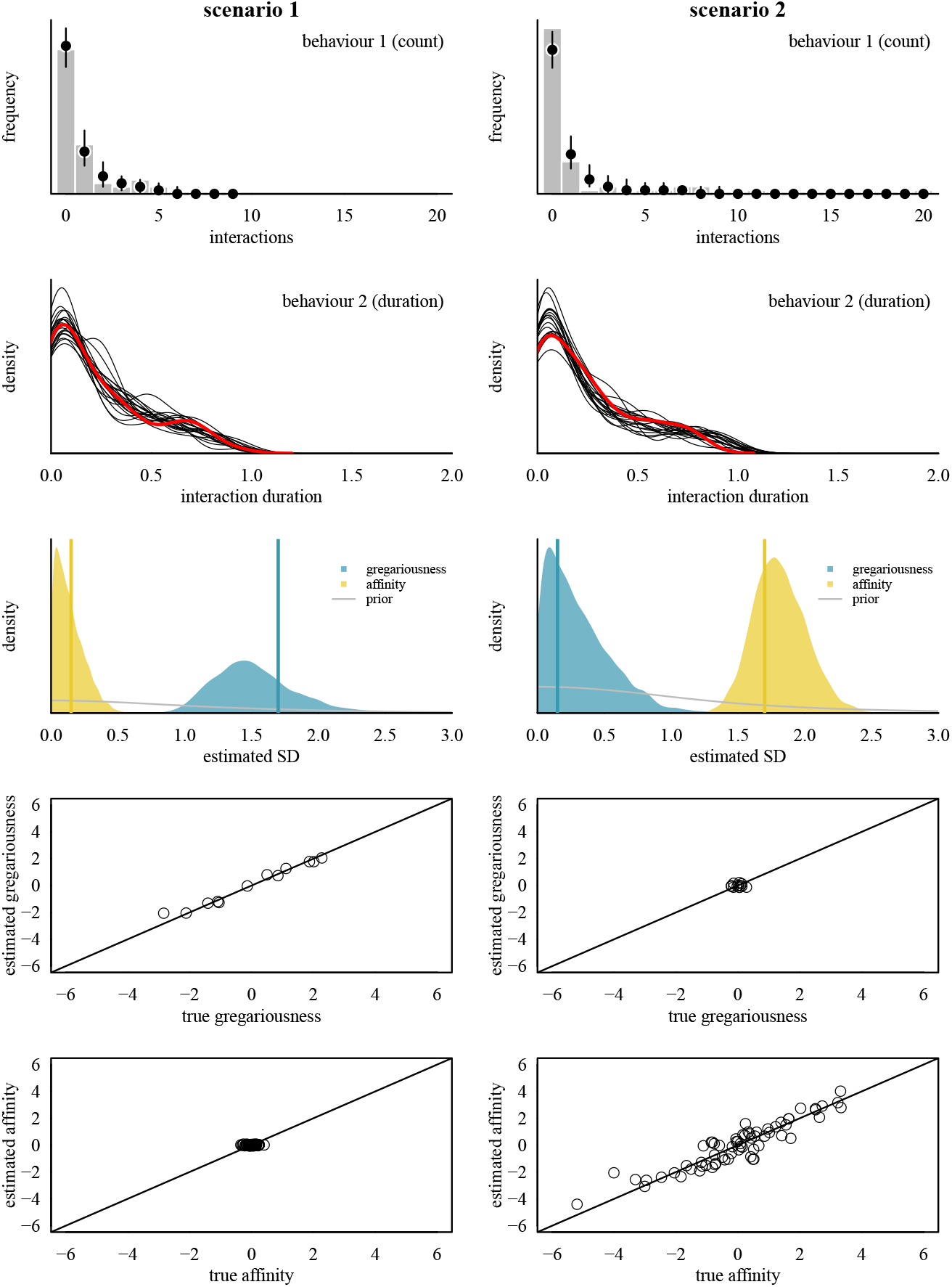
Model results of the two scenarios, each using two different interaction types (a frequency and a duration) to infer the underlying individual and dyadic sociality axes. The two top rows show posterior predictive checks for both behaviors. The third row summarizes variation in gregariousness and affinity. The two bottom rows show the relationship between true and estimated values for gregariousness and affinity.

### S5 Posterior predictive checks

In this section we illustrate with two examples how posterior predictive checks can be indicative of poor model fit.

#### S5.1 Example 1

We generate an example data set for 21 individuals with one behavior. Then we fit two models to those interaction data. The first model is the correct one (it reflects the data generation process). The second model is wrong from the outset. It is a model that omits estimating dyadic affinity altogether^4^. Since the second model is wrong, we would hope to see some indication of this model being wrong in the posterior checks. And if we can’t see it’s wrong, it would be nice to see at least that it is *worse* than the first model.

**Figure S8:**
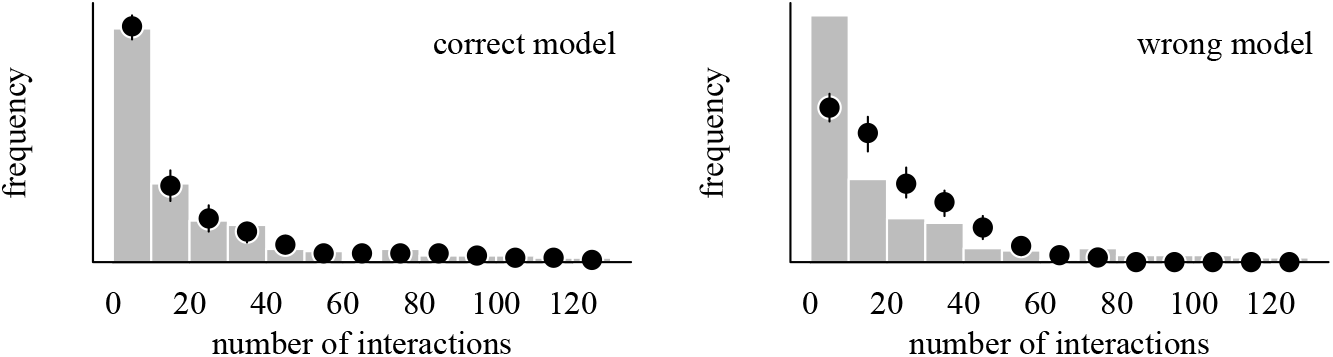
Posterior predictive check for simulated data and fitted to a correct and a wrong model. The histograms in both panels are identical and show the distribution of *observed* interactions. The overlaid circles and lines are from posterior simulations, i.e., we generated new interaction patterns from our estimated model parameters. We did this for all draws. In a ‘good’ model, the simulated data align closely with the observed data. Here, the second model produces arguably worse predictions than the first model.

**Figure S9:**
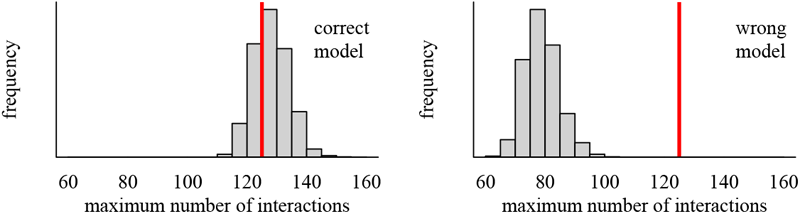
Another posterior predictive check indicating that one of the fitted models is worse than the other. Here we show the maximum value of interactions from each draw. This is done as a histogram of the 4000 values (corresponding to 4000 draws). The red line indicates the maximum value in the *observed* data. In the first case, the observed value falls nicely in the middle of the posterior samples. In the second case, the observed value is much larger than what the model we applied generated. In other words, in none of the posterior samples did we predict a maximum value as high as in the observed data.

It is important to reiterate that this procedure does not allow us to say that we may have found the *true* model. Rather, we can use this framework to assess how well our model at hand can predict new data that is similar to what we have observed in the first place. If our model cannot do that, we learn that we probably need a better model.

If we have two competing models (as in the example above), we can at least get to a point where we can assess that one model is *better* than the other.

### S5.2 Code for example 1

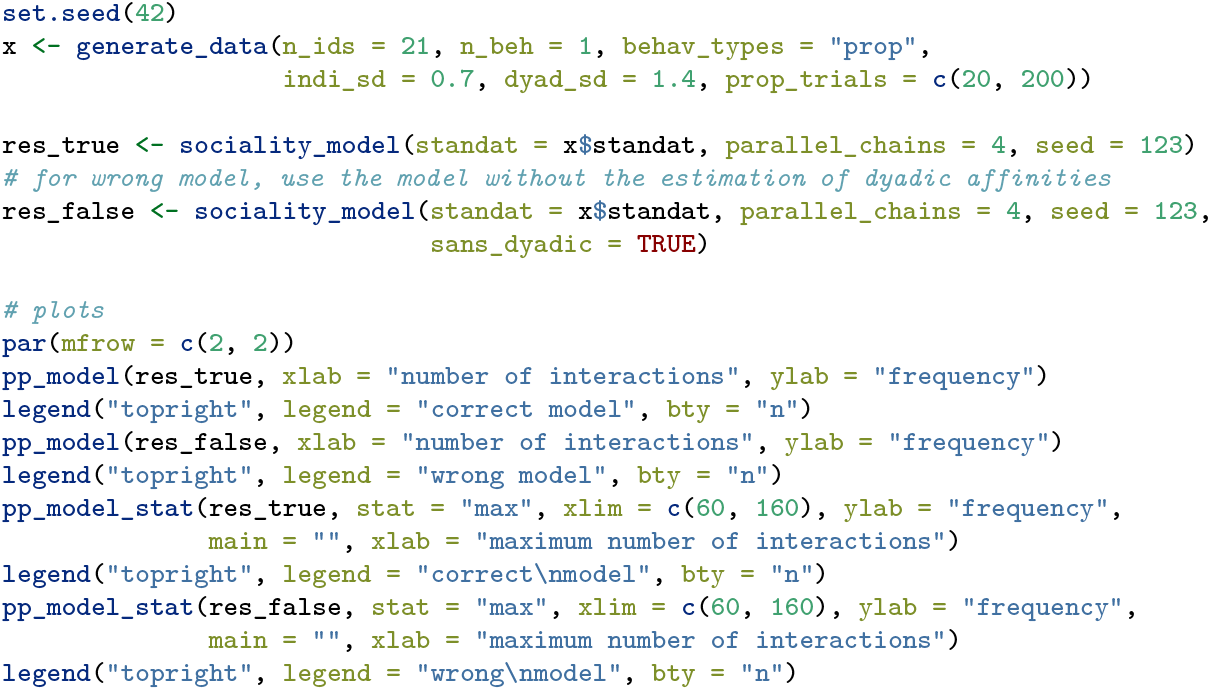

### S5.3 Example 2

The second example is a bit more complex. Here we simulate two kinds of interactions (say grooming and proximity). We then add the caveat the we now assume (and simulate) that there are two distinct affinity axes (not only one) and that we can set the degree of correlation among them! We also have two gregariousness axes (not just one, and all that applies to affinity below also applies to gregariousness but we’ll omit that here for brevity). The key is that we set the correlation to a very small value (0.1 in the example). That effectively means that the two affinity axes do *not* correlate. Another way of looking at that is that we now have two distinct (“uncorrelated”) processes that drive behavior 1 and 2. If our model assumes that there is exactly one such axis (which it does), then it should have trouble with data that were generated like this.^5^

Below we simulate such a data set and then we fit our original model (which is the wrong model in the context here). We also fit a second: a model that actually estimates the correlation (that’s the correct model here). The prediction being that the first model will have trouble while the second model will be much better.

The results of this posterior check do not necessarily indicate that the first model is good and second model is bad in an absolute sense. Instead, the comparison is relative: model one is better than model two, with respect to predict the observed interaction patterns. Although, trying to speak objectively: arguably the correct model does make a pretty good job of predicting the interactions (even in an absolute sense).

### S5.4 Code for example 2

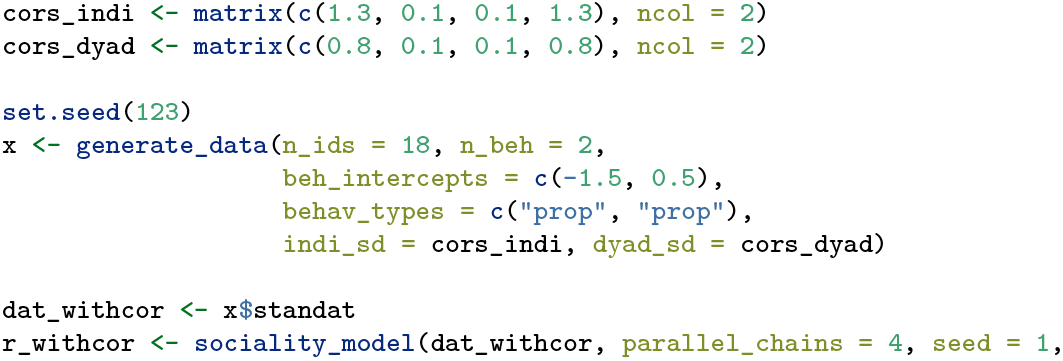

**Figure S10:**
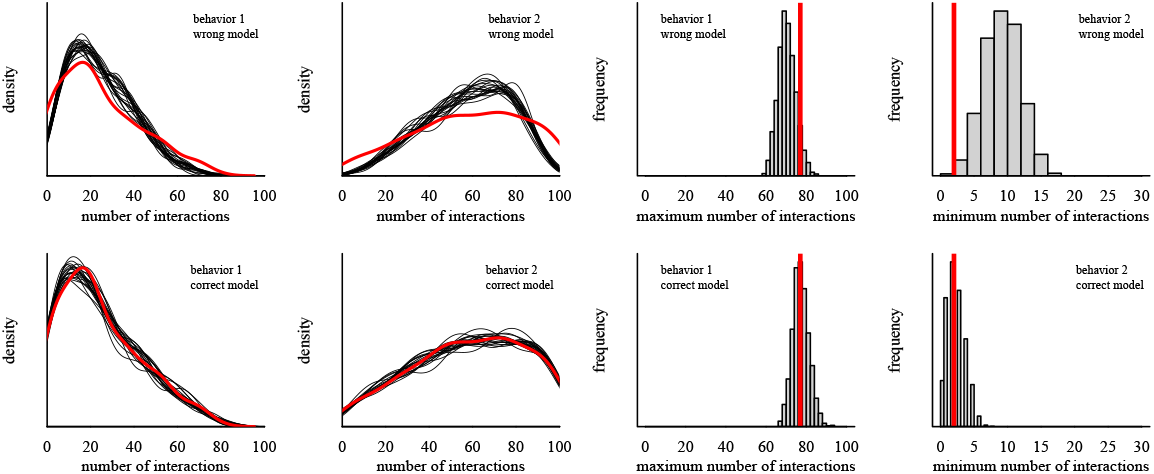
Posterior predictive checks can help identifying problems with model fits. Top row: incorrect model. Here, there are deviations between predictions and observed data, indicating that this model is substantially worse than the model in the top row. These deviations range from fairly obvious (interactions in behavior 2) to less severe (maximum values in behavior 1). Bottom row: correct model. Both behaviors are well predicted by the model (and quite obviously better than in the top row), i.e., the posterior samples are very similar to the observed data. Both models were fitted to the same data set.

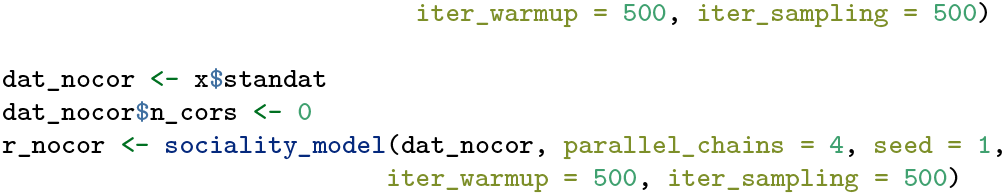

### S6 Evaluation results

**Figure S11:**
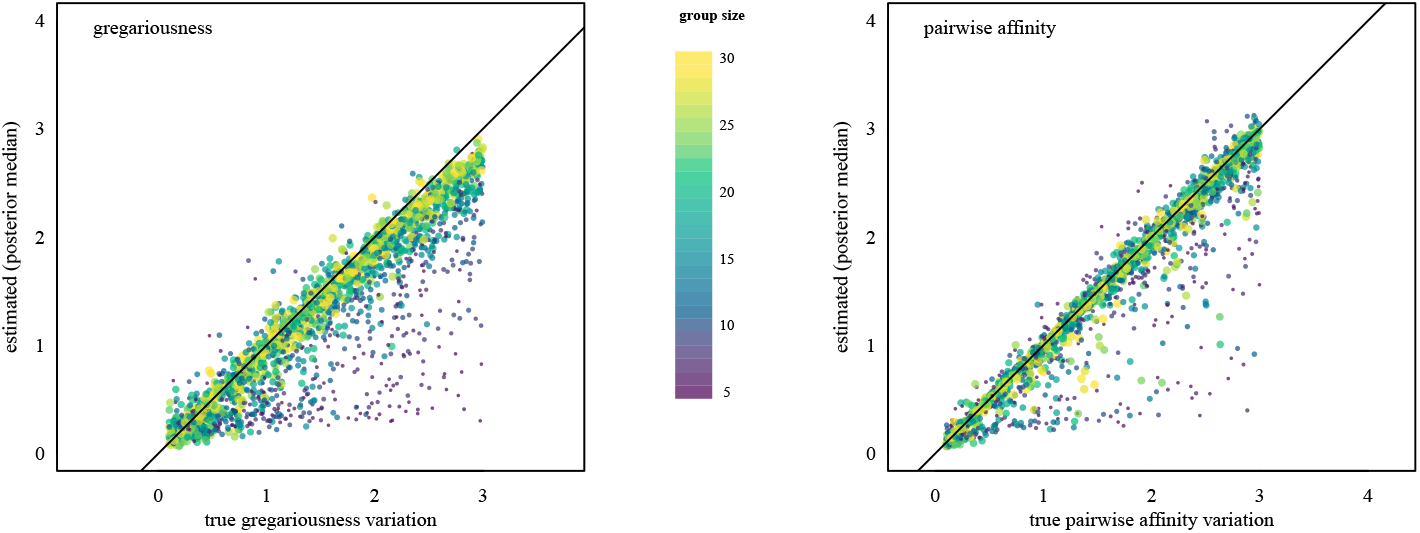
Recovering individual and dyadic sociality variation. Each circle represents one simulated interaction network. Group size is encoded by color and area (smaller circles correspond to smaller groups).

**Figure S12:**
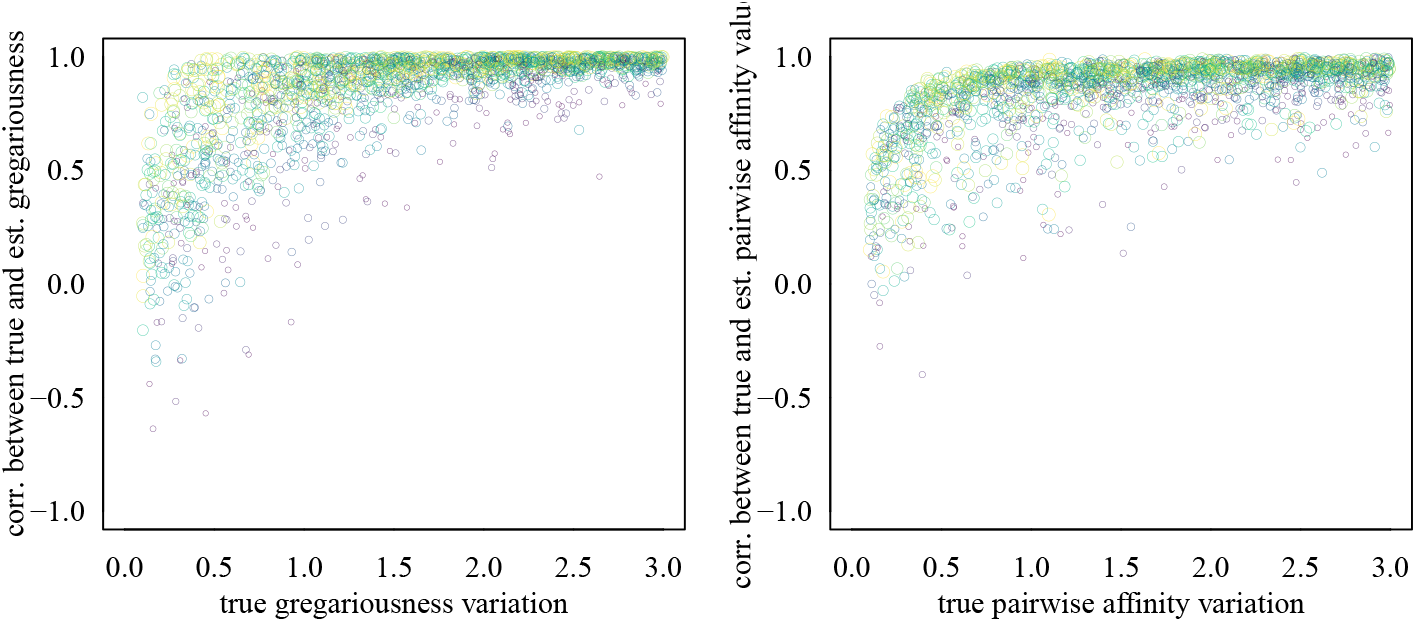
Correlations between true and estimated individual gregariousness values and between true and estimated pairwise affinity values within simulated networks. Each circle represents one simulated interaction network

**Figure S13:**
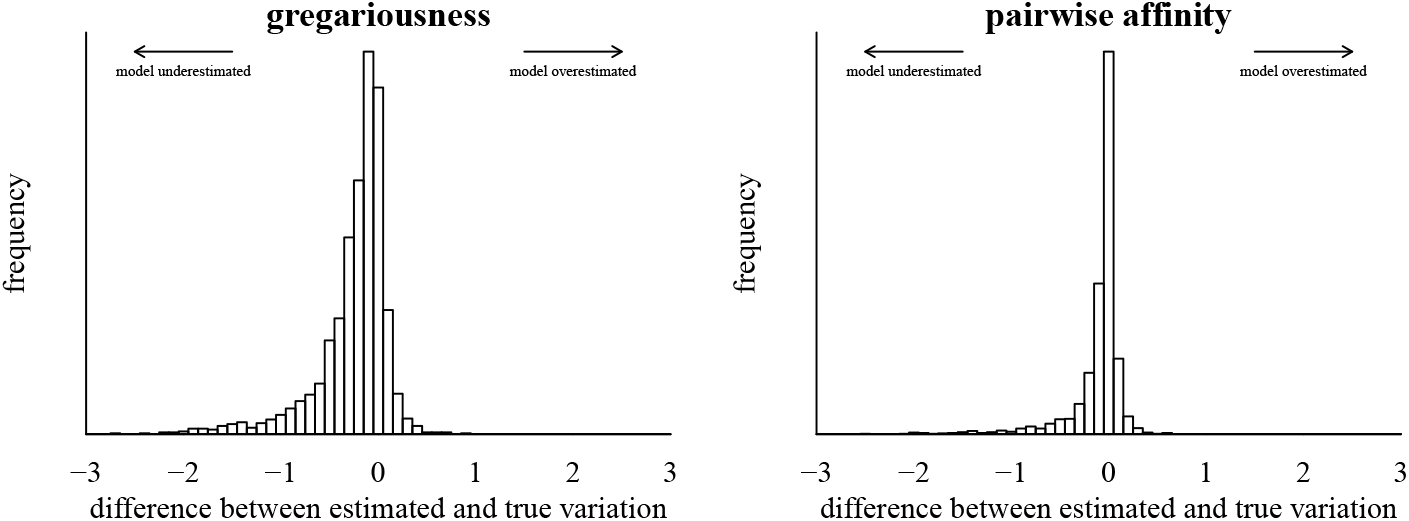
Differences between true and estimated gregariousness and affinity variation in simulated data.

**Figure S14:**
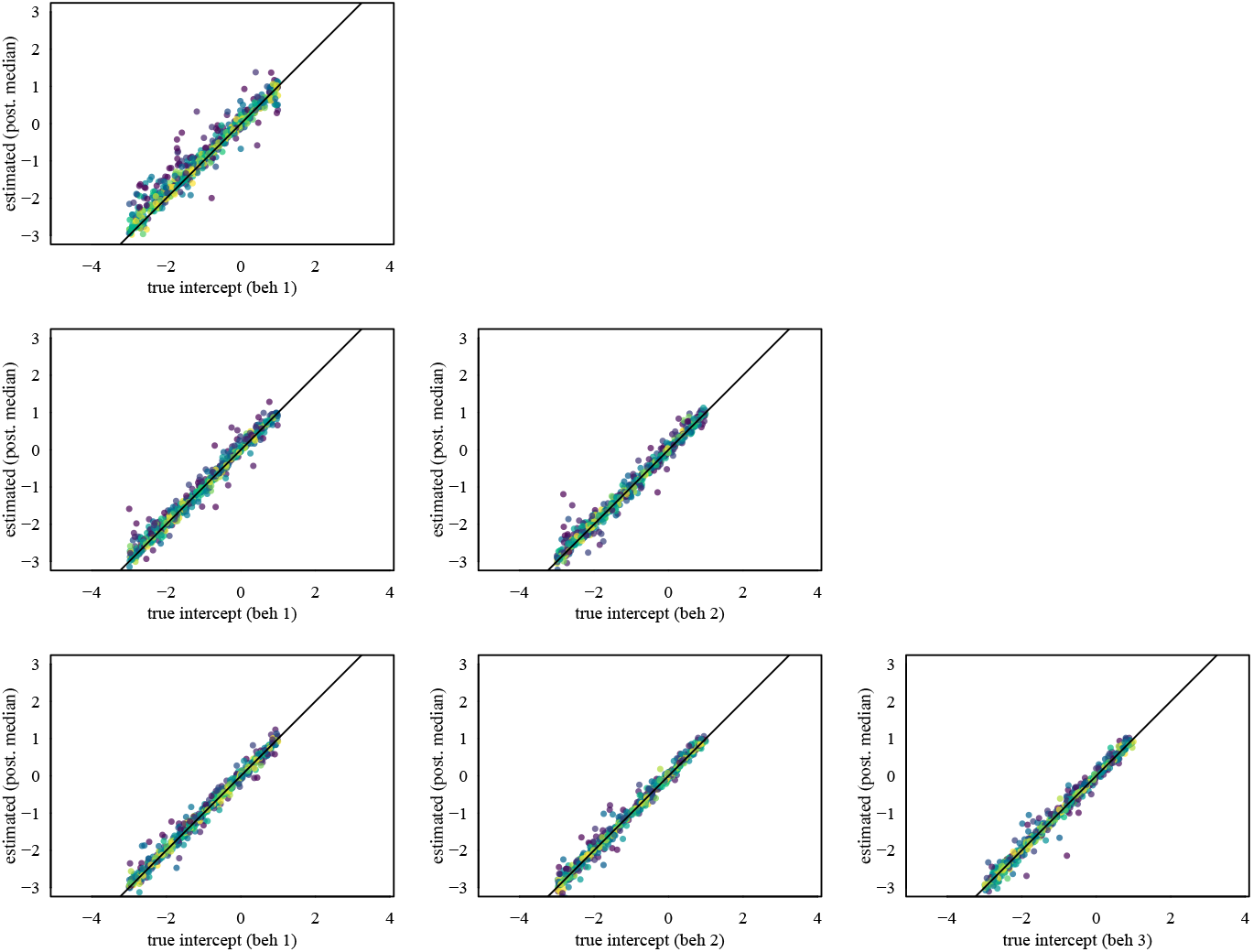
Recovery of intercepts (group level baseline parameters), stratified by the number of behaviors that were included in the model. Top row: data sets with one behavior generated. Middle row: data sets with two behaviors generated. Bottom row: data sets with three behaviors generated.

**Figure S15:**
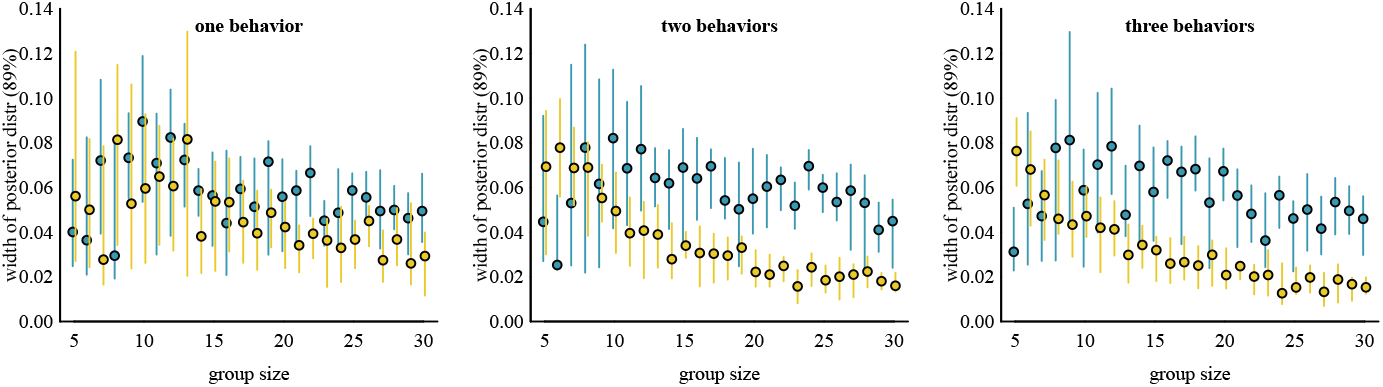
Width of posteriors in relation to group sizes. Generally, posteriors become narrower with increasing group size.

### S7 Macaque grooming

**Figure S16:**
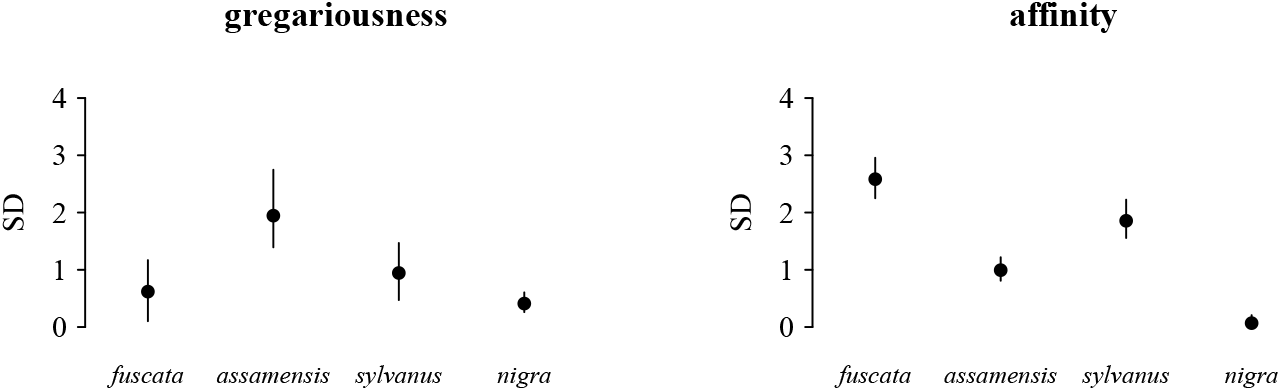
Variation in gregariousness and relationships in four groups from four species (tentatively ordered by their social style, from despotic (*fuscata*) to tolerant (*nigra*)). Shown are posterior medians with 89% credible intervals. This figure presents essentially the same information as the figure above, displayed in a different way.

**Figure S17:**
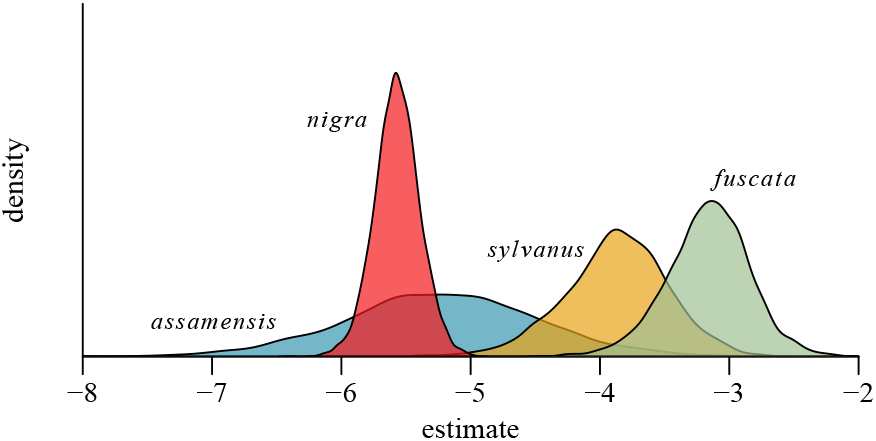
Posterior distributions of baseline parameters for the sociality models for the four macaque species.

### S8 Network strength and clustering in Stan

In order to implement the analysis of the kangaroo data (figure 5), we had to code functions that calculate strength and the local clustering coefficient from adjacency matrices in Stan. As reference we used the `igraph` package (Csardi & Nepusz, 2006), which the original authors also used. Figure S18 shows for a simulated data set that our Stan functions generated results identical to `igraph`’s `transitivity(graph, type = “weighted”)` (for clustering) and `strength(graph)` (for strength). For definitions of strength and clustering see Croft et al. (2008).

**Figure S18:**
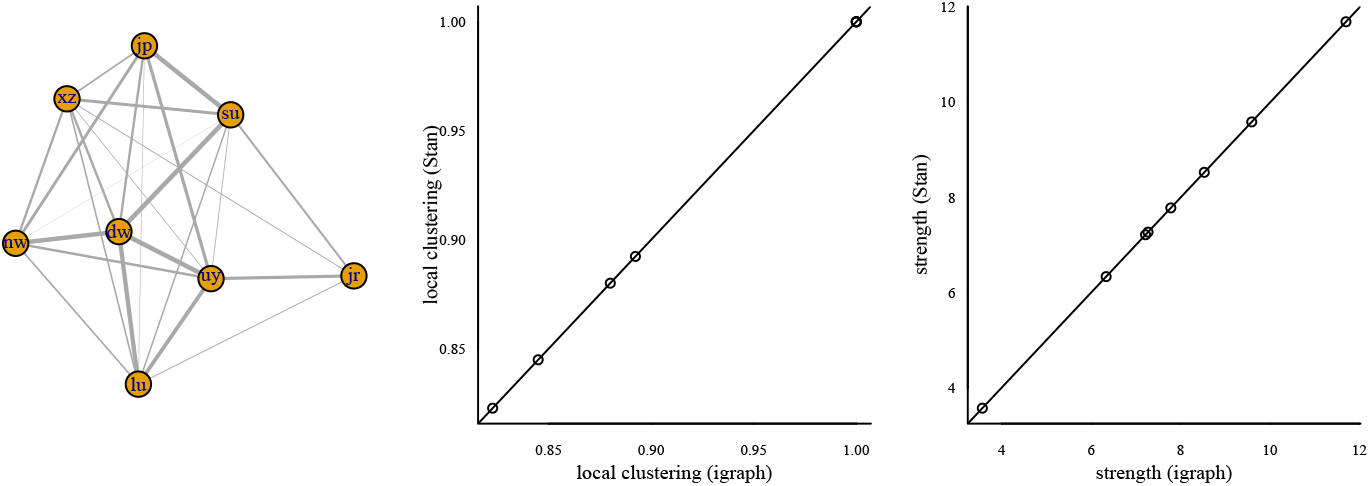
Functions for obtaining local clustering and strength written in Stan produce identical results as the `igraph` package.

## Footnotes

1 Accounting for varying observation effort via offset terms can easily be integrated.

2 In figure 4b, some of the predictions for the outcome for the individual with the highest gregariousness were larger than 50 (not visible in the figure), which seems not very likely given the range of the *observed* outcomes.

3 We need to exponentiate (i.e., use a link function) because we use a Poisson process with a log-link to generate the data.

4 For now, it’s included in the package for illustrative purposes only. In the future, we might add formal (i.e., numerical) model comparisons.

5 Essentially, if we assume only one axis to exist we can also see this as having multiple axes that are all perfectly correlated.

