## Supplementary material for "(De)composing sociality: disentangling individual-specific from dyad-specific propensities to interact": bamoso R package: bayesian_kangaroos.stan

```
functions {
  // convert dyadic values ('HWI', or 'strength', or 'dyad_soc_vals') to individual summed strength value per period
vector dyads2indistrength(int n_model,
                          int n_dyads_over_periods,
                          vector dyad_soc_vals,
                          array[] int model_id,
                          array[] int model_period,
                          array[] int navi_time_period,
                          array[] int navi_id1,
                          array[] int navi_id2,
                          array[] int navi_vector_loc) {
  vector[n_model] res = rep_vector(0.0, n_model);
  for (m in 1:n_model) {
    int temp_id = model_id[m];
    int temp_period = model_period[m];
    for (n in 1:n_dyads_over_periods) {
      if (navi_time_period[n] == temp_period) {
        if (navi_id1[n] == temp_id) {
          res[m] = res[m] + dyad_soc_vals[navi_vector_loc[n]];
        }
        if (navi_id2[n] == temp_id) {
          res[m] = res[m] + dyad_soc_vals[navi_vector_loc[n]];
        }
      }
    }
  }
  return(res);
}

// take individual sociality values and match them to individual values needed for model
vector match_indisoc(vector indi_soc_vals, int n_model, int n_ids_over_periods,
                     array[] int model_id, array[] int model_period, array[] int navi2_1_id_num, array[] int navi2_2_period) {
  vector[n_model] res = rep_vector(0.0, n_model);
  for (m in 1:n_model) {
    int temp_id = model_id[m];
    int temp_period = model_period[m];
    for (o in 1:n_ids_over_periods) {
      if (navi2_1_id_num[o] == temp_id) {
        if (navi2_2_period[o] == temp_period) {
          res[m] = indi_soc_vals[o];
        }
      }
    }
  }
  return(res);
}

// convert individual values to sums of two individuals in a dyad
// returns a vector that corresponds to all period-dyads
// in R (for one period): outer(rowSums(m), rowSums(m), "+")[lower.tri(m)]
vector indi2indisum(int n_dyads_over_periods,
                    int n_ids_over_periods,
                    array[] int navi_time_period,
                    array[] int navi_id1,
                    array[] int navi_id2,
                    vector indi_soc_vals,
                    array[] int navi2_id,
                    array[] int navi2_time_period) {
  vector[n_dyads_over_periods] res = rep_vector(0.0, n_dyads_over_periods);
  for (i in 1:n_dyads_over_periods) {
    int temp_id1 = navi_id1[i];
    int temp_id2 = navi_id2[i];
    int temp_period = navi_time_period[i];
    for (o in 1:n_ids_over_periods) {
      if (navi2_time_period[o] == temp_period) {
        if (navi2_id[o] == temp_id1) {
          res[i] = res[i] + indi_soc_vals[o];
        }
        if (navi2_id[o] == temp_id2) {
          res[i] = res[i] + indi_soc_vals[o];
        }
      }
    }
  }
  return(res);
}

// obtain local weighted or unweighted transitivity (individual cluster coefficient)
// modelled after igraph::transitivity(net, "weighted") and igraph::transitivity(net, "local")
// objects that are handed over as matrices to stan need to be hardcoded as
//   integer vectors (eg. navi_triads -> navi_triads_1)
// function returns one value per individual and period (i.e. if A was present in periods 2 and 3 it gets 2 values, if B was present only in period 2 it gets only one value)
//   in other words: this returns a vector that may have a different size from what is necessary for any downstream model

vector cluster_local_weighted(int n_periods,
                   int n_dyads,
                   int n_ids_over_periods,
                   int n_dyads_over_periods,
                   array[] int per_period_ids,
                   array[] int per_period_triads_per_id,
                   // array[] int navi_triads,
                   array[] int navi_triads_1_id1,
                   array[] int navi_triads_2_id2,
                   array[] int navi_triads_3_id3,
                   array[] int navi_triads_4_period,
                   array[] int navi_triads_5_f1_2,
                   array[] int navi_triads_6_f1_3,
                   array[] int navi_triads_7_f2_1,
                   array[] int navi_triads_8_f2_3,
                   array[] int navi_triads_9_f3_1,
                   array[] int navi_triads_10_f3_2,
                   // array[] int navi_triads_sel,
                   array[] int navi_triads_sel_2_loc_across,
                   array[] int navi_triads_sel_3_id,
                   array[] int navi_triads_sel_4_period,
                   // array[] int navi2,
                   array[] int navi2_1_id_num,
                   array[] int navi2_2_period,
                   // array[] int navi_dyads,
                   array[] int navi_dyads_1_id1,
                   array[] int navi_dyads_2_id2,
                   array[] int navi_dyads_4_period,
                   array[] int navi_dyads_5_vectorloc,
                   vector edgeweights, // typically: dyadic sociality
                   // vector strengths,
                   // vector degrees,
                   real cut_off,
                   int keep_nan,
                   int do_weighted) // use edge *weights* (or not)
                   {
  vector[n_ids_over_periods] res;
  vector[size(edgeweights)] edgeweights_binary;
  vector[n_ids_over_periods] degrees = rep_vector(0.0, n_ids_over_periods);
  vector[n_ids_over_periods] strengths = rep_vector(0.0, n_ids_over_periods);

  // calculate degree and strength for each individual in each period
  for (m in 1:n_ids_over_periods) {
    int temp_id = navi2_1_id_num[m];
    int temp_period = navi2_2_period[m];
    for (n in 1:n_dyads_over_periods) {
      if (navi_dyads_4_period[n] == temp_period) {
        if (navi_dyads_1_id1[n] == temp_id) {
          if (edgeweights[navi_dyads_5_vectorloc[n]] > cut_off) {
            degrees[m] = degrees[m] + 1.0;
            strengths[m] = strengths[m] + edgeweights[navi_dyads_5_vectorloc[n]];
          }
        }
        if (navi_dyads_2_id2[n] == temp_id) {
          if (edgeweights[navi_dyads_5_vectorloc[n]] > cut_off) {
            degrees[m] = degrees[m] + 1;
            strengths[m] = strengths[m] + edgeweights[navi_dyads_5_vectorloc[n]];
          }
        }
      }
    }
  }


  // make each edge binary given a cut-off
  for (i in 1:size(edgeweights)) {
    if (edgeweights[i] > cut_off) {
      edgeweights_binary[i] = 1.0;
    } else {
      edgeweights_binary[i] = 0.0;
    }
  }
  for (p in 1:n_periods) {
    for (i in 1:per_period_ids[p]) {
      // which id (numeric code...)
      int temp_id = 0;
      int cnt = 1;
      for (k in 1:n_ids_over_periods) {
        if (navi2_2_period[k] == p) {
          if (cnt == i) {
            temp_id = navi2_1_id_num[k];
          }
          cnt = cnt + 1;
        }
      }

      // get indices of triad data
      array[per_period_triads_per_id[p]] int temp_index;
      cnt = 1;
      for (n in 1:size(navi_triads_sel_3_id)) {
        if (navi_triads_sel_4_period[n] == p) {
          if (navi_triads_sel_3_id[n] == temp_id) {
            temp_index[cnt] = navi_triads_sel_2_loc_across[n];
            cnt = cnt + 1;
          }
        }
      }
      // get sum of wij and wih (the two dyads with i)
      // and binary versions
      vector[size(temp_index)] dyadic_sums;
      vector[size(temp_index)] dyadic_prod;
      cnt = 1;
      for (k in 1:size(navi_triads_1_id1)) {
        if (navi_triads_4_period[k] == p) {
          if (navi_triads_1_id1[k] == temp_id) {
            dyadic_sums[cnt] = edgeweights[navi_triads_5_f1_2[k]] + edgeweights[navi_triads_6_f1_3[k]]; // f1_2 and f1_3
            dyadic_prod[cnt] = edgeweights_binary[navi_triads_5_f1_2[k]] * edgeweights_binary[navi_triads_6_f1_3[k]] * edgeweights_binary[navi_triads_8_f2_3[k]];
            // extra is f2_3
            cnt = cnt + 1;
          }
          if (navi_triads_2_id2[k] == temp_id) {
            dyadic_sums[cnt] = edgeweights[navi_triads_7_f2_1[k]] + edgeweights[navi_triads_8_f2_3[k]]; // f2_1 and f2_3
            dyadic_prod[cnt] = edgeweights_binary[navi_triads_7_f2_1[k]] * edgeweights_binary[navi_triads_8_f2_3[k]] * edgeweights_binary[navi_triads_6_f1_3[k]];
            // extra is f1_3
            cnt = cnt + 1;
          }
          if (navi_triads_3_id3[k] == temp_id) {
            dyadic_sums[cnt] = edgeweights[navi_triads_9_f3_1[k]] + edgeweights[navi_triads_10_f3_2[k]]; // f3_1 and f3_2
            dyadic_prod[cnt] = edgeweights_binary[navi_triads_9_f3_1[k]] * edgeweights_binary[navi_triads_10_f3_2[k]] * edgeweights_binary[navi_triads_5_f1_2[k]];
            // extra is f1_2
            cnt = cnt + 1;
          }

        }
      }
      // part 2 of equation combined
      real part2 = 0.0;
      if (do_weighted) {
        for (k in 1:size(dyadic_sums)) {
          part2 = part2 + dyadic_sums[k] * 0.5 * dyadic_prod[k];
        }
      } else {
        for (k in 1:size(dyadic_sums)) {
          part2 = part2 + dyadic_prod[k];
        }
      }


      // first part (individual strength and degree)
      real s = 0.0;
      real d = 0.0;
      real part1 = 0.0;
      if (do_weighted) {
        for (k in 1:size(navi2_2_period)) {
          if (navi2_2_period[k] == p) {
            if (navi2_1_id_num[k] == temp_id) {
              s = strengths[k];
              d = degrees[k];
            }
          }
        }
        part1 = 1 / (s * (d - 1));
      } else {
        for (k in 1:size(navi2_2_period)) {
          if (navi2_2_period[k] == p) {
            if (navi2_1_id_num[k] == temp_id) {
              s = degrees[k]; // just need degrees again for unweighted version...
              d = degrees[k];
            }
          }
        }
        part1 = 1 / (s * (d - 1));
      }


      // calculate final value and add to results
      real final_val = part1 * part2;
      for (k in 1:size(navi2_2_period)) {
        if (navi2_2_period[k] == p) {
          if (navi2_1_id_num[k] == temp_id) {
            res[k] = final_val * 2;
          }
        }
      }


    }
  }

  // replace nan with 0
  if (keep_nan == 0) {
    for (i in 1:size(res)) {
      if (is_nan(res[i])) {
        res[i] = 0.0;
      }
    }
  }
  return(res);
}


}

data {

  int<lower=0> n_ids; // total number of individuals
  int<lower=0> n_dyads;
  int<lower=0> n_periods;
  int<lower=0> n_dyads_over_periods; // total number of observations (dyads in p1 + dyads in p2 + ...) for association data
  int<lower=0> n_ids_over_periods; // number of id/period combinations
  int<lower=0> n_model; // total number of observations for the actual individual-level model

  // parameter for cut-off for degree (a degree is only counted if larger than this value)
  // and also counts towards clustering only if larger than this value
  real edge_cutoff;

  // navigation through ids and dyads for associations
  array[n_dyads_over_periods, 5] int<lower=0> navi_dyads; // first two cols for ids, third: dyad, fourth: period, fifth: location in a vector with dyadic values
  array[n_ids_over_periods, 2] int<lower=0> navi2; // first col is id, second col is numeric period

  // navigation through association data for clustering calculations
  int<lower=0> navi_triad_nrow;
  int<lower=0> navi_triad_sel_nrow;
  array[navi_triad_nrow, 10] int<lower=0> navi_triads;
  array[navi_triad_sel_nrow, 4] int<lower=0> navi_triads_sel;
  array[n_periods] int<lower=0> per_period_ids;
  array[n_periods] int<lower=0> per_period_triads_per_id;

  // navigation through ids and period for model
  array[n_model] int<lower=0> model_id;
  array[n_model] int<lower=0> model_period;

  // data declarations for association data
  array[n_dyads_over_periods] int<lower=0> seen_together; // numerator
  array[n_dyads_over_periods] int<lower=0> seen_either; // 'observation effort' (denominator)

  // response variable
  array[n_model] int<lower=0> model_y;

  // additional predictors for the actual model
  vector[n_model] predictor_primip;
  vector[n_model] predictor_resid_gs;
  vector[n_model] predictor_num_sightings;

}
parameters {
  // for association part
  real<lower=0> indi_soc_sd; // SD parameter (sd_1 in brms)
  vector[n_ids_over_periods] indi_soc_vals_z; // blups on z-scale (z_1 in brms)
  real<lower=0> dyad_soc_sd; // SD parameter (sd_1 in brms)
  vector[n_dyads_over_periods] dyad_soc_vals_z; // blups on z-scale (z_1 in brms)
  vector[n_periods] period_intercepts; // intercepts for periods (as 'fixed effect', i.e. no hyperprior)

  // for actual model
  // global parameters
  real slope_strength;
  // real slope_indisoc;
  real slope_cluster;
  real intercept;

  real slope_primip;
  real slope_resid_gs;
  real slope_num_sightings;

  // group-level parameters ('random effects')
  real<lower=0> model_indi_intercept_sd;
  vector[n_ids] model_indi_intercepts_z; // standardized BLUPS
  real<lower=0> model_period_intercept_sd;
  vector[n_periods] model_period_intercepts_z;
  // random slopes for strength in period
  real<lower=0> model_period_strength_slope_sd;
  vector[n_periods] model_period_strength_slope_z; //
  // random slopes for cluster in period
  real<lower=0> model_period_cluster_slope_sd;
  vector[n_periods] model_period_cluster_slope_z; //
  // random slopes for indisoc in period
  // real<lower=0> model_period_indisoc_slope_sd;
  // vector[n_periods] model_period_indisoc_slope_z; //

}
transformed parameters {
  vector[n_ids_over_periods] indi_soc_vals; // actual BLUPS for individual sociality
  vector[n_dyads_over_periods] dyad_soc_vals; // actual BLUPS for dyadic sociality
  vector[n_ids] model_indi_intercepts;
  vector[n_periods] model_period_intercepts;
  vector[n_periods] model_period_strength_slope;
  // vector[n_periods] model_period_indisoc_slope;
  vector[n_periods] model_period_cluster_slope;

  // for association part
  indi_soc_vals = (indi_soc_sd * indi_soc_vals_z);
  dyad_soc_vals = (dyad_soc_sd * dyad_soc_vals_z);

  // for actual model
  model_indi_intercepts = (model_indi_intercept_sd * model_indi_intercepts_z);
  model_period_intercepts = (model_period_intercept_sd * model_period_intercepts_z);

  model_period_strength_slope = (model_period_strength_slope_sd * model_period_strength_slope_z);
  // model_period_indisoc_slope = (model_period_indisoc_slope_sd * model_period_indisoc_slope_z);
  model_period_cluster_slope = (model_period_cluster_slope_sd * model_period_cluster_slope_z);
}

model {
  // needed helper objects
  // model 1: association indices (network stuff)
  // vector[n_dyads_over_periods] indi_sums = rep_vector(0.0, n_dyads_over_periods);
  // model 2: actual regression model
  vector[n_model] strength_predictor = rep_vector(0.0, n_model);
  // vector[n_model] indisoc_predictor = rep_vector(0.0, n_model);
  vector[n_model] cluster_predictor = rep_vector(0.0, n_model);
  // vector[n_model] model_mu = rep_vector(0.0, n_model);


  // get to associations: individual and dyadic sociality contributions
  seen_together ~ binomial_logit(seen_either, period_intercepts[navi_dyads[, 4]] +
                                                sqrt(0.5) * indi2indisum(n_dyads_over_periods, n_ids_over_periods, navi_dyads[, 4], navi_dyads[, 1], navi_dyads[, 2],
                                                                   indi_soc_vals, navi2[, 1], navi2[, 2]) +
                                                dyad_soc_vals);

  // actual model of binary response
  // compute individual strength values for the downstream model
  strength_predictor = dyads2indistrength(n_model, n_dyads_over_periods, dyad_soc_vals, model_id, model_period, navi_dyads[, 4], navi_dyads[, 1], navi_dyads[, 2], navi_dyads[, 5]);
  // compute (rather: extract) individual clustering values for downstream model
  cluster_predictor = cluster_local_weighted(n_periods, n_dyads, n_ids_over_periods, n_dyads_over_periods,
                                          per_period_ids, per_period_triads_per_id,
                                          navi_triads[, 1], navi_triads[, 2], navi_triads[, 3], navi_triads[, 4], navi_triads[, 5],
                                          navi_triads[, 6], navi_triads[, 7], navi_triads[, 8], navi_triads[, 9], navi_triads[, 10],
                                          navi_triads_sel[, 2], navi_triads_sel[, 3], navi_triads_sel[, 4],
                                          navi2[, 1], navi2[, 2],
                                          navi_dyads[, 1], navi_dyads[, 2], navi_dyads[, 4], navi_dyads[, 5],
                                          dyad_soc_vals, edge_cutoff, 0, 0) - 0.5; // centered
  // we ignore the individual-level predictor because this is captured by strength
  // indisoc_predictor = match_indisoc(indi_soc_vals, n_model, n_ids_over_periods, model_id, model_period, navi2[, 1], navi2[, 2]);

  model_y ~ bernoulli_logit(intercept + model_indi_intercepts[model_id] + model_period_intercepts[model_period]
                                + (slope_strength + model_period_strength_slope[model_period]) .* strength_predictor
                                // + (slope_strength) .* strength_predictor // without random slopes
                                // + (slope_indisoc + model_period_indisoc_slope[model_period]) .* indisoc_predictor
                                // + (slope_indisoc) .* indisoc_predictor // without random slope
                                + (slope_cluster + model_period_cluster_slope[model_period]) .* cluster_predictor
                                // + (slope_cluster) .* cluster_predictor // without random slopes
                                + slope_primip * predictor_primip
                                + slope_resid_gs * predictor_resid_gs
                                + slope_num_sightings * predictor_num_sightings
                                // + slope_indisoc * indisoc_predictor
                                );


  // priors for model parameters
  slope_strength ~ normal(0, 0.2);
  // slope_indisoc ~ normal(0, 0.2);
  slope_cluster ~ normal(0, 0.2);
  intercept ~ normal(0, 0.5);

  slope_primip ~ normal(0, 1);
  slope_resid_gs ~ normal(0, 1);
  slope_num_sightings ~ normal(0, 1);

  model_indi_intercept_sd ~ student_t(3, 0, 1);
  model_indi_intercepts_z ~ normal(0, 1);

  model_period_intercept_sd ~ student_t(3, 0, 1);
  model_period_intercepts_z ~ normal(0, 1); // model_period_strength_slope_sd ~ student_t(3, 0, 1);
  model_period_strength_slope_sd ~ normal(0, 0.2);
  model_period_strength_slope_z ~ normal(0, 1);
  model_period_cluster_slope_sd ~ normal(0, 0.2);
  model_period_cluster_slope_z ~ normal(0, 1);
  // model_period_indisoc_slope_sd ~ normal(0, 0.2);
  // model_period_indisoc_slope_z ~ normal(0, 1);


  // priors for association
  indi_soc_sd ~ student_t(3, 0, 1);
  indi_soc_vals_z ~ normal(0, 1);
  dyad_soc_sd ~ student_t(3, 0, 1);
  dyad_soc_vals_z ~ normal(0, 1);

  // prior for period intercepts
  period_intercepts ~ student_t(3, -5, 1);

}

generated quantities {
  array[n_dyads_over_periods] int<lower=0> seen_together_rep;
  vector[n_dyads_over_periods] indi_sums_for_pp = rep_vector(0.0, n_dyads_over_periods);
  vector[n_model] strength_for_pp = rep_vector(0.0, n_model);
  // vector[n_model] indisoc_for_pp = rep_vector(0.0, n_model);
  vector[n_model] cluster_for_pp = rep_vector(0.0, n_model);

  array[n_model] int<lower=0> model_y_rep;


  // individual summed strength values from dyadic sociality values
  strength_for_pp = dyads2indistrength(n_model, n_dyads_over_periods, dyad_soc_vals, model_id, model_period, navi_dyads[, 4], navi_dyads[, 1], navi_dyads[, 2], navi_dyads[, 5]);
  // individual sociality
  // indisoc_for_pp = match_indisoc(indi_soc_vals, n_model, n_ids_over_periods, model_id, model_period, navi2[, 1], navi2[, 2]);
  // clustering values
  cluster_for_pp = cluster_local_weighted(n_periods, n_dyads, n_ids_over_periods, n_dyads_over_periods,
                                          per_period_ids, per_period_triads_per_id,
                                          navi_triads[, 1], navi_triads[, 2], navi_triads[, 3], navi_triads[, 4], navi_triads[, 5],
                                          navi_triads[, 6], navi_triads[, 7], navi_triads[, 8], navi_triads[, 9], navi_triads[, 10],
                                          navi_triads_sel[, 2], navi_triads_sel[, 3], navi_triads_sel[, 4],
                                          navi2[, 1], navi2[, 2],
                                          navi_dyads[, 1], navi_dyads[, 2], navi_dyads[, 4], navi_dyads[, 5],
                                          dyad_soc_vals, edge_cutoff, 0, 0) - 0.5;


  model_y_rep = bernoulli_rng(inv_logit(intercept + model_indi_intercepts[model_id] + model_period_intercepts[model_period]
                                        + (slope_strength + model_period_strength_slope[model_period]) .* strength_for_pp
                                        // + (slope_strength) .* strength_for_pp
                                        // + (slope_indisoc + model_period_indisoc_slope[model_period]) .* indisoc_for_pp
                                        + (slope_cluster + model_period_cluster_slope[model_period]) .* cluster_for_pp
                                        // + (slope_cluster) .* cluster_for_pp
                                        + slope_primip * predictor_primip
                                        + slope_resid_gs * predictor_resid_gs
                                        + slope_num_sightings * predictor_num_sightings
                                        ));


  // for association data
  // regenerate output of how often is each pair seen together
  // 1) dyads as sums of two individuals
  indi_sums_for_pp = indi2indisum(n_dyads_over_periods, n_ids_over_periods,
                                  navi_dyads[, 4], navi_dyads[, 1], navi_dyads[, 2],
                                  indi_soc_vals,
                                  navi2[, 1], navi2[, 2]);
  // regenerate output for association data
  seen_together_rep = binomial_rng(seen_either,
      inv_logit(dyad_soc_vals + (sqrt(0.5) * indi_sums_for_pp) + period_intercepts[navi_dyads[, 4]]));

}
```
