## Supplementary material for "(De)composing sociality: disentangling individual-specific from dyad-specific propensities to interact": bamoso R package: interaction_model.stan

```
// interactions can be any combination of counts,
//   discrete proportions (count and trials), continuous proportions (beta),
//   positive continuous (gamma)

// we require actually a second observation effort matrix that is
//   of type 'integer', otherwise stan will complain

// similarly, there are two matrices for the actual interactions,
//   one with integers (for pois and binomial) and
//   one with continuous numbers (for beta and gamma models)

functions {
  real lin2prob(real x, real obseff) {
    return(1 - exp(-exp(x) * obseff));
  }
  vector lin2prob_vec(vector x, vector obseff) {
    return(1 - exp(-exp(x) .* obseff));
  }

  real prob2lin(real x, real obseff) {
    return(log((-log(1 - x))/obseff));
  }
  vector prob2lin_vec(vector x, vector obseff) {
    return(log((-log(1 - x))./obseff));
  }

  real mybern_lpdf(real y, real mu, real obseff) {
    real bern_mu = lin2prob(mu, obseff);
    if (y == 0) {
      return bernoulli_lpmf(0 | bern_mu);
    } else {
      return bernoulli_lpmf(1 | bern_mu);
    }
  }
}

data {
  int<lower=0> n_ids;
  int<lower=0> n_dyads;
  int<lower=0> n_beh;
  array[n_beh + 1] int<lower=0,upper=6> behav_types;
  // 1: counts (poiss); 2: counts (proportion, binomial); 3: cont. duration (gamma);
  // 4: cont. proportion (beta); 5: cont. duration with 0 (mixture gamma/bernoulli)
  // 6: binary (bernoulli)
  array[n_dyads, n_beh] int<lower=0> interactions;
  matrix[n_dyads, n_beh] interactions_cont;
  array[n_dyads, 2] int<lower=0> dyads_navi;
  matrix[n_dyads, n_beh] obseff;
  array[n_dyads, n_beh] int<lower=0> obseff_int;

  // flag for prior predictive check
  int<lower=0,upper=1> prior_only;

  // priors for intercepts
  matrix[n_beh, 2] prior_matrix;
  matrix[n_beh, 2] prior_matrix2;

  // info about indexing and number of optional shape parameter(s)
  int<lower=0> gamma_shape_n;
  array[n_beh] int<lower=0> gamma_shape_pos;
  int<lower=0> beta_shape_n;
  array[n_beh] int<lower=0> beta_shape_pos;
}
parameters {
  real<lower=0> indi_soc_sd; // SD parameter
  vector[n_ids] indi_soc_vals_z; // blups on z-scale
  real<lower=0> dyad_soc_sd; // SD parameter
  vector[n_dyads] dyad_soc_vals_z; // blups on z-scale
  vector[n_beh] beh_intercepts; // intercepts for behaviours
  vector[gamma_shape_n] beh_intercepts_add; // second set of intercepts for behaviours
  // vector[gamma_shape_n] aux_baserates_unconstrained; // baserates

  vector[gamma_shape_n] shapes_gamma_raw; // unconstrained (<lower=0>)
  vector<lower=0>[beta_shape_n] shapes_beta; // unconstrained (<lower=0>)
  // vector[beta_shape_n] shapes_beta_raw; // unconstrained (<lower=0>)
}
transformed parameters {
  vector[n_ids] indi_soc_vals;  // actual blups
  vector[n_dyads] dyad_soc_vals;  // actual blups
  vector[n_dyads] scaled_indi_sums;  // sums of dyadic values, scaled
  // vector[gamma_shape_n] aux_baserates = inv_logit(aux_baserates_unconstrained); // baserates
  vector<lower=0>[gamma_shape_n] shapes_gamma = exp(shapes_gamma_raw);
  // vector<lower=0>[beta_shape_n] shapes_beta = exp(shapes_beta_raw);
  indi_soc_vals = (indi_soc_sd * indi_soc_vals_z);
  dyad_soc_vals = (dyad_soc_sd * dyad_soc_vals_z);
  scaled_indi_sums = sqrt(0.5) * (indi_soc_vals[dyads_navi[, 1]] + indi_soc_vals[dyads_navi[, 2]]);
}
model {
  vector[n_dyads] lp = rep_vector(0.0, n_dyads);
  // linear predictor
  lp = lp + scaled_indi_sums + dyad_soc_vals;

  if (prior_only == 0) {
    for (i in 1:n_beh) {
      if (behav_types[i] == 5) {
        // bernoulli/gamma mixture
        for (n in 1:n_dyads) {
          if (interactions_cont[n, i] == 0) {
            target += mybern_lpdf(interactions_cont[n, i] | beh_intercepts[i] + lp[n], obseff[n, i]);
          } else {
            target += mybern_lpdf(interactions_cont[n, i] | beh_intercepts[i] + lp[n], obseff[n, i]);
            target += gamma_lpdf(interactions_cont[n, i] |
              shapes_gamma[gamma_shape_pos[i]],
              shapes_gamma[gamma_shape_pos[i]] /
              exp(lp[n] + beh_intercepts_add[gamma_shape_pos[i]] + log(obseff[n, i]))
              );
            // y[i] ~ gamma(shape_gamma, shape_gamma/exp(mu_gamma));
          }
        }
      }
      if (behav_types[i] == 1) {
        interactions[, i] ~ poisson(exp(lp + beh_intercepts[i] + log(obseff[, i])));
      }
      if (behav_types[i] == 2) {
        interactions[, i] ~ binomial_logit(obseff_int[, i], lp + beh_intercepts[i]);
      }
      if (behav_types[i] == 3) {
        interactions_cont[, i] ~ gamma(shapes_gamma[gamma_shape_pos[i]],
                                       shapes_gamma[gamma_shape_pos[i]] / exp(lp + beh_intercepts[i] + log(obseff[, i]))); //
      }
      if (behav_types[i] == 4) {
        interactions_cont[, i] ~ beta(inv_logit(lp + beh_intercepts[i]) * shapes_beta[beta_shape_pos[i]],
                                     (1 - inv_logit(lp + beh_intercepts[i])) *  shapes_beta[beta_shape_pos[i]] ); //
      }
      if (behav_types[i] == 6) {
        for (n in 1:n_dyads) {
          interactions[n, i] ~ bernoulli(lin2prob(lp[n] + beh_intercepts[i], obseff[n, i]));
        }
      }
    }
  }


  for (i in 1:n_beh) {
    // declare priors for intercepts based on observed data
    // has been done outside Stan (and is passed as data)
    beh_intercepts[i] ~ student_t(3, prior_matrix[i, 1], prior_matrix[i, 2]);

    // extra priors for shape/dispersion parameters
    if (behav_types[i] == 3) {
      // shapes[gamma_shape_pos[i]] ~ gamma(0.01, 0.01);
      // shapes_gamma_raw[gamma_shape_pos[i]] ~ normal(1, 1);
      shapes_gamma_raw[gamma_shape_pos[i]] ~ student_t(4, 1, 1);
    }
    if (behav_types[i] == 4) {
      shapes_beta[beta_shape_pos[i]] ~ gamma(0.1, 0.1);
      // shapes_beta_raw[beta_shape_pos[i]] ~ student_t(4, 1, 1);
    }
    if (behav_types[i] == 5) {
      // gamma prior comes from prior_matrix2
      beh_intercepts_add[gamma_shape_pos[i]] ~ student_t(3, prior_matrix2[i, 1], prior_matrix2[i, 2]); // gamma in mixture
      shapes_gamma_raw[gamma_shape_pos[i]] ~ normal(1, 1);
    }
  }
  indi_soc_vals_z ~ normal(0, 1);
  dyad_soc_vals_z ~ normal(0, 1);
  // indi_soc_sd ~ student_t(3, 0, 1);
  // dyad_soc_sd ~ student_t(3, 0, 1);
  indi_soc_sd ~ exponential(2);
  dyad_soc_sd ~ exponential(2);
}

generated quantities {
  array[n_dyads, n_beh] int interactions_pred;
  matrix[n_dyads, n_beh] interactions_pred_cont;
  matrix[n_dyads, n_beh] log_lik; // pointwise log likelihoods

  {
    vector[n_dyads] LP = scaled_indi_sums + dyad_soc_vals; // linear predictor without behaviour-specific intercept
    for (i in 1:n_beh) {
      if (behav_types[i] == 1) {
        interactions_pred[, i] = poisson_log_rng(LP + beh_intercepts[i] + log(obseff[, i]));
        // interactions_pred_cont[, i] = to_vector(poisson_log_rng(LP + beh_intercepts[i] + log(obseff[, i])));
        for (jj in 1:n_dyads) { // loglik
          log_lik[jj, i] = poisson_log_lpmf(interactions[jj, i] | LP[jj] + beh_intercepts[i] + log(obseff[jj, i]));
        }
      }
      if (behav_types[i] == 2) {
        interactions_pred[, i] = binomial_rng(obseff_int[, i], inv_logit(LP + beh_intercepts[i]));
        // interactions_pred_cont[, i] = to_vector(binomial_rng(obseff_int[, i], inv_logit(LP + beh_intercepts[i])));
        for (jj in 1:n_dyads) { // loglik
          log_lik[jj, i] = binomial_logit_lpmf(interactions[jj, i] | obseff_int[jj, i], LP[jj] + beh_intercepts[i]);
        }
      }
      if (behav_types[i] == 3) {
        interactions_pred_cont[, i] = to_vector(gamma_rng(shapes_gamma[gamma_shape_pos[i]],
        shapes_gamma[gamma_shape_pos[i]] *
        exp(-(LP + beh_intercepts[i] + log(obseff[, i])))));
        for (jj in 1:n_dyads) { // loglik
          log_lik[jj, i] = gamma_lpdf(interactions_cont[jj, i] | shapes_gamma[gamma_shape_pos[i]],
          shapes_gamma[gamma_shape_pos[i]] * exp(-(LP[jj] + beh_intercepts[i] + log(obseff[jj, i]))));
        }
      }
      if (behav_types[i] == 4) {
        interactions_pred_cont[, i] = to_vector(beta_rng(
                                      shapes_beta[beta_shape_pos[i]] * (    inv_logit(LP + beh_intercepts[i])),
                                      shapes_beta[beta_shape_pos[i]] * (1 - inv_logit(LP + beh_intercepts[i]))
                                      ));
        for (jj in 1:n_dyads) { // loglik
          log_lik[jj, i] = beta_lpdf(interactions_cont[jj, i] |
                                     inv_logit(LP[jj] + beh_intercepts[i])  * shapes_beta[beta_shape_pos[i]],
                                (1 - inv_logit(LP[jj] + beh_intercepts[i])) * shapes_beta[beta_shape_pos[i]]);

        }
      }
      if (behav_types[i] == 5) {
        for (n in 1:n_dyads) {
          int aux = bernoulli_rng(lin2prob(beh_intercepts[i] + LP[n], obseff[n, i]));
          if (aux == 0) {
            interactions_pred_cont[n, i] = 0.0;
          } else {
            interactions_pred_cont[n, i] = gamma_rng(shapes_gamma[gamma_shape_pos[i]] , shapes_gamma[gamma_shape_pos[i]] / exp(LP[n] + beh_intercepts_add[gamma_shape_pos[i]] + log(obseff[n, i])));
          }

          if (interactions_cont[n, i] == 0) {
            log_lik[n, i] = mybern_lpdf(interactions_cont[n, i] | beh_intercepts[i] + LP[n], obseff[n, i]);
          } else {
            log_lik[n, i] = mybern_lpdf(interactions_cont[n, i] | beh_intercepts[i] + LP[n], obseff[n, i]) +
              gamma_lpdf(interactions_cont[n, i] | shapes_gamma[gamma_shape_pos[i]],
              shapes_gamma[gamma_shape_pos[i]] / exp(LP[n] + beh_intercepts_add[gamma_shape_pos[i]] + log(obseff[n, i])));
          }

        }
      }
      if (behav_types[i] == 6) {
        for (n in 1:n_dyads) {
          interactions_pred[n, i] = bernoulli_rng(lin2prob(beh_intercepts[i] + LP[n], obseff[n, i]));
          log_lik[n, i] = bernoulli_lpmf(interactions[n, i] | lin2prob(LP[n] + beh_intercepts[i], obseff[n, i]));
        }
      }
    }

  }

}
```
