## Supplementary material for "(De)composing sociality: disentangling individual-specific from dyad-specific propensities to interact": bamoso R package: interaction_model_corr.stan

```
// model axes separately for different behaviors
//   allows assessment of correlation

functions {
  vector chol2corvec(matrix inputobj, int n_axes, int n_cor) {
    matrix[n_axes, n_axes] cor_mat = multiply_lower_tri_self_transpose(inputobj);
    vector[n_cor] out;
    int idx = 1;
    for (k in 1:rows(cor_mat)) {
      for (j in 1:(k - 1)) {
        out[idx] = cor_mat[j, k];
        idx = idx + 1;
      }
    }
  return out;
  }
}

data {
  int<lower=0> n_ids;
  int<lower=0> n_dyads;
  int<lower=2> n_beh;
  array[n_beh + 1] int<lower=0> behav_types; // 1: counts (poiss), 2: counts (proportion, binomial)
  array[n_dyads, n_beh] int<lower=0> interactions;
  matrix[n_dyads, n_beh] interactions_cont;
  array[n_dyads, 2] int<lower=0> dyads_navi;
  matrix[n_dyads, n_beh] obseff;
  array[n_dyads, n_beh] int<lower=0> obseff_int;

  // priors for intercepts
  matrix[n_beh, 2] prior_matrix;

  // info about indexing and number of optional shape parameter(s)
  int<lower=0> gamma_shape_n;
  array[n_beh] int<lower=0> gamma_shape_pos;
  int<lower=0> beta_shape_n;
  array[n_beh] int<lower=0> beta_shape_pos;

  int<lower=0> n_cors;
}
parameters {
  vector<lower=0>[n_beh] indi_soc_sd; // SD parameter
  matrix[n_beh, n_ids] indi_soc_vals_z; // blups on z-scale
  vector<lower=0>[n_beh] dyad_soc_sd; // SD parameter
  matrix[n_beh, n_dyads] dyad_soc_vals_z; // blups on z-scale
  vector[n_beh] beh_intercepts; // intercepts for behaviours

  vector<lower=0>[gamma_shape_n] shapes;
  vector<lower=0>[beta_shape_n] shapes_beta;

  // correlation
  cholesky_factor_corr[n_beh] chol_indi_dims; // cholesky factor of correlation matrix for individual-level dims
  cholesky_factor_corr[n_beh] chol_dyad_dims; // cholesky factor of correlation matrix for dyad-level dims

}

transformed parameters {
  matrix[n_ids, n_beh] indi_soc_vals;  // actual blups
  matrix[n_dyads, n_beh] dyad_soc_vals;  // actual blups
  matrix[n_dyads, n_beh] scaled_indi_sums;  // sums of dyadic values, scaled
  indi_soc_vals = transpose(diag_pre_multiply(indi_soc_sd, chol_indi_dims) * indi_soc_vals_z);
  dyad_soc_vals = transpose(diag_pre_multiply(dyad_soc_sd, chol_dyad_dims) * dyad_soc_vals_z);
  for (i in 1:n_beh) {
    scaled_indi_sums[, i] = sqrt(0.5) * (indi_soc_vals[dyads_navi[, 1], i] + indi_soc_vals[dyads_navi[, 2], i]);
  }

}
model {
  vector[n_dyads] lp = rep_vector(0.0, n_dyads);
  vector[n_dyads] indi_sums = rep_vector(0.0, n_dyads);

  for (i in 1:n_beh) {
    lp = rep_vector(0.0, n_dyads);
    // indi_sums = sqrt(0.5) * (indi_soc_vals[dyads_navi[, 1], i] + indi_soc_vals[dyads_navi[, 2], i]);
    // linear predictor
    lp = lp + scaled_indi_sums[, i] + dyad_soc_vals[, i];

    if (behav_types[i] == 1) {
      interactions[, i] ~ poisson(exp(lp + beh_intercepts[i] + log(obseff[, i])));
    }
    if (behav_types[i] == 2) {
      interactions[, i] ~ binomial_logit(obseff_int[, i], lp + beh_intercepts[i]);
    }
    if (behav_types[i] == 3) {
      interactions_cont[, i] ~ gamma(shapes[gamma_shape_pos[i]],
                                       shapes[gamma_shape_pos[i]] * exp(-(lp + beh_intercepts[i] + log(obseff[, i])))); //
    }
    if (behav_types[i] == 4) {
      interactions_cont[, i] ~ beta(inv_logit(lp + beh_intercepts[i]) * shapes_beta[beta_shape_pos[i]],
                                     (1 - inv_logit(lp + beh_intercepts[i])) *  shapes_beta[beta_shape_pos[i]] ); //
    }
  }

  for (i in 1:n_beh) {
    // declare priors for intercepts based on observed data
    // has been done outside Stan (and is passed as data)
    beh_intercepts[i] ~ student_t(3, prior_matrix[i, 1], prior_matrix[i, 2]);

    // extra priors for shape/dispersion parameters
    if (behav_types[i] == 3) {
      shapes[gamma_shape_pos[i]] ~ gamma(0.01, 0.01);
    }
    if (behav_types[i] == 4) {
      shapes_beta[beta_shape_pos[i]] ~ gamma(0.1, 0.1);
    }
    indi_soc_vals_z[i, ] ~ normal(0, 1);
    dyad_soc_vals_z[i, ] ~ normal(0, 1);
  }
  // indi_soc_sd ~ student_t(3, 0, 1);
  // dyad_soc_sd ~ student_t(3, 0, 1);
  indi_soc_sd ~ exponential(2);
  dyad_soc_sd ~ exponential(2);
  chol_indi_dims ~ lkj_corr_cholesky(2);
  chol_dyad_dims ~ lkj_corr_cholesky(2);
}

generated quantities {
  array[n_dyads, n_beh] int interactions_pred;
  matrix[n_dyads, n_beh] interactions_pred_cont;
  matrix[n_dyads, n_beh] log_lik; // pointwise log likelihoods
  vector<lower=-1,upper=1>[n_cors] cors_indi = chol2corvec(chol_indi_dims, n_beh, n_cors);
  vector<lower=-1,upper=1>[n_cors] cors_dyad = chol2corvec(chol_dyad_dims, n_beh, n_cors);

  //

  {
    for (i in 1:n_beh) {
      vector[n_dyads] LP = scaled_indi_sums[, i] + dyad_soc_vals[, i]; // linear predictor without behaviour-specific intercept
      if (behav_types[i] == 1) {
        interactions_pred[, i] = poisson_log_rng(LP + beh_intercepts[i] + log(obseff[, i]));
        for (jj in 1:n_dyads) { // loglik
          log_lik[jj, i] = poisson_log_lpmf(interactions[jj, i] | LP[jj] + beh_intercepts[i] + log(obseff[jj, i]));
        }
      }
      if (behav_types[i] == 2) {
        interactions_pred[, i] = binomial_rng(obseff_int[, i], inv_logit(LP + beh_intercepts[i]));
        for (jj in 1:n_dyads) { // loglik
          log_lik[jj, i] = binomial_logit_lpmf(interactions[jj, i] | obseff_int[jj, i], LP[jj] + beh_intercepts[i]);
        }
      }
      if (behav_types[i] == 3) {
        interactions_pred_cont[, i] = to_vector(gamma_rng(shapes[gamma_shape_pos[i]],
                                      shapes[gamma_shape_pos[i]] *
                                        exp(-(LP + beh_intercepts[i] + log(obseff[, i])))));
        for (jj in 1:n_dyads) { // loglik
          log_lik[jj, i] = gamma_lpdf(interactions_cont[jj, i] | shapes[gamma_shape_pos[i]],
                                       shapes[gamma_shape_pos[i]] * exp(-(LP[jj] + beh_intercepts[i] + log(obseff[jj, i]))));
        }
      }
      if (behav_types[i] == 4) {
          interactions_pred_cont[, i] = to_vector(beta_rng(
                                          shapes_beta[beta_shape_pos[i]] * (    inv_logit(LP + beh_intercepts[i])),
                                          shapes_beta[beta_shape_pos[i]] * (1 - inv_logit(LP + beh_intercepts[i]))
                                          ));
        for (jj in 1:n_dyads) { // loglik
          log_lik[jj, i] = beta_lpdf(interactions_cont[jj, i] |
                                     inv_logit(LP[jj] + beh_intercepts[i])  * shapes_beta[beta_shape_pos[i]],
                                (1 - inv_logit(LP[jj] + beh_intercepts[i])) * shapes_beta[beta_shape_pos[i]]);

        }

      }

    }

  }
}
```
