## Supplementary material for "(De)composing sociality: disentangling individual-specific from dyad-specific propensities to interact": bamoso R package: interaction_model_indicat.stan

```
// interactions can be any combination of counts,
//   discrete proportions (count and trials), continuous proportions (beta),
//   positive continuous (gamma)

// we require actually a second observation effort matrix that is
//   of type 'integer', otherwise stan will complain

// similarly, there are two matrices for the actual interactions,
//   one with integers (for pois and binomial) and
//   one with continuous numbers (for beta and gamma models)

// this a first extension with a categorical individual-level predictor (e.g., sex)


data {
  int<lower=0> n_ids;
  int<lower=0> n_dyads;
  int<lower=0> n_beh;
  // int<lower=0> behav_types[n_beh + 1]; // 1: counts (poiss), 2: counts (proportion, binomial) // old way
  array[n_beh + 1] int<lower=0> behav_types; // 1: counts (poiss), 2: counts (proportion, binomial)
  // int<lower=0> interactions[n_dyads, n_beh]; // old way
  array[n_dyads, n_beh] int<lower=0> interactions;
  matrix[n_dyads, n_beh] interactions_cont;
  // int<lower=0> dyads_navi[n_dyads, 2]; // old way
  array[n_dyads, 2] int<lower=0> dyads_navi;
  matrix[n_dyads, n_beh] obseff;
  // int<lower=0> obseff_int[n_dyads, n_beh]; // old way
  array[n_dyads, n_beh] int<lower=0> obseff_int;

  // priors for intercepts
  matrix[n_beh, 2] prior_matrix;

  // info about indexing and number of optional shape parameter(s)
  int<lower=0> gamma_shape_n;
  // int<lower=0> gamma_shape_pos[n_beh]; // old way
  array[n_beh] int<lower=0> gamma_shape_pos;
  int<lower=0> beta_shape_n;
  // int<lower=0> beta_shape_pos[n_beh]; // old way
  array[n_beh] int<lower=0> beta_shape_pos;

  // additional predictors
  vector<lower=0,upper=1>[n_ids] indi_cat_pred;
}
parameters {
  real<lower=0> indi_soc_sd; // SD parameter
  vector[n_ids] indi_soc_vals_z; // blups on z-scale
  real<lower=0> dyad_soc_sd; // SD parameter
  vector[n_dyads] dyad_soc_vals_z; // blups on z-scale
  vector[n_beh] beh_intercepts; // intercepts for behaviours

  vector<lower=0>[gamma_shape_n] shapes;
  vector<lower=0>[beta_shape_n] shapes_beta;

  real indi_cat_eff;
}

transformed parameters {
  vector[n_ids] indi_soc_vals;  // actual blups
  vector[n_dyads] dyad_soc_vals;  // actual blups
  vector[n_ids] indi_cat_aux;
  vector[n_dyads] scaled_indi_sums;  // sums of dyadic values, scaled
  indi_cat_aux = indi_cat_eff * indi_cat_pred;
  indi_soc_vals = (indi_soc_sd * indi_soc_vals_z) + indi_cat_aux;
  dyad_soc_vals = (dyad_soc_sd * dyad_soc_vals_z);
  scaled_indi_sums = sqrt(0.5) * (indi_soc_vals[dyads_navi[, 1]] + indi_soc_vals[dyads_navi[, 2]]);
}

model {
  // vector[n_dyads] lp = rep_vector(0.0, n_dyads);
  // vector[n_dyads] indi_sums = 0.5 * (indi_soc_vals[dyads_navi[, 1]] + indi_soc_vals[dyads_navi[, 2]]);

  // linear predictor
  vector[n_dyads] lp = scaled_indi_sums + dyad_soc_vals;


  for (i in 1:n_beh) {
    if (behav_types[i] == 1) {
      interactions[, i] ~ poisson(exp(lp + beh_intercepts[i] + log(obseff[, i])));
    }
    if (behav_types[i] == 2) {
      interactions[, i] ~ binomial_logit(obseff_int[, i], lp + beh_intercepts[i]);
    }
    if (behav_types[i] == 3) {
      interactions_cont[, i] ~ gamma(shapes[gamma_shape_pos[i]],
                                       shapes[gamma_shape_pos[i]]  * exp(-(lp + beh_intercepts[i] + log(obseff[, i]))))  ; //
    }
    if (behav_types[i] == 4) {
      interactions_cont[, i] ~ beta(inv_logit(lp + beh_intercepts[i]) * shapes_beta[beta_shape_pos[i]],
                                     (1 - inv_logit(lp + beh_intercepts[i])) *  shapes_beta[beta_shape_pos[i]] )  ; //
    }
  }

  for (i in 1:n_beh) {
    // declare priors for intercepts based on observed data
    // has been done outside Stan (and is passed as data)
    beh_intercepts[i] ~ student_t(3, prior_matrix[i, 1], prior_matrix[i, 2]);

    // extra priors for shape/dispersion parameters
    if (behav_types[i] == 3) {
      shapes[gamma_shape_pos[i]] ~ gamma(0.01, 0.01);
    }
    if (behav_types[i] == 4) {
      shapes_beta[beta_shape_pos[i]] ~ gamma(0.1, 0.1);
    }
  }
  indi_soc_vals_z ~ normal(0, 1);
  dyad_soc_vals_z ~ normal(0, 1);
  indi_soc_sd ~ student_t(3, 0, 1);
  dyad_soc_sd ~ student_t(3, 0, 1);
  indi_cat_eff ~ normal(0, 1);
}

generated quantities {
  array[n_dyads, n_beh] int interactions_pred;
  matrix[n_dyads, n_beh] interactions_pred_cont;
  matrix[n_dyads, n_beh] log_lik; // pointwise log likelihoods

  {
    vector[n_dyads] LP = scaled_indi_sums + dyad_soc_vals; // linear predictor without behaviour-specific intercept
    for (i in 1:n_beh) {
      if (behav_types[i] == 1) {
        interactions_pred[, i] = poisson_log_rng(LP + beh_intercepts[i] + log(obseff[, i]));
        for (jj in 1:n_dyads) { // loglik
          log_lik[jj, i] = poisson_log_lpmf(interactions[jj, i] | LP[jj] + beh_intercepts[i] + log(obseff[jj, i]));
        }
      }
      if (behav_types[i] == 2) {
        interactions_pred[, i] = binomial_rng(obseff_int[, i], inv_logit(LP + beh_intercepts[i]));
        for (jj in 1:n_dyads) { // loglik
          log_lik[jj, i] = binomial_logit_lpmf(interactions[jj, i] | obseff_int[jj, i], LP[jj] + beh_intercepts[i]);
        }
      }
      if (behav_types[i] == 3) {
        interactions_pred_cont[, i] = to_vector(gamma_rng(shapes[gamma_shape_pos[i]],
                                      shapes[gamma_shape_pos[i]] *
                                        exp(-(LP + beh_intercepts[i] + log(obseff[, i])))));
        for (jj in 1:n_dyads) { // loglik
          log_lik[jj, i] = gamma_lpdf(interactions_cont[jj, i] | shapes[gamma_shape_pos[i]],
                                       shapes[gamma_shape_pos[i]] * exp(-(LP[jj] + beh_intercepts[i] + log(obseff[jj, i]))));
        }
      }
      if (behav_types[i] == 4) {
          interactions_pred_cont[, i] = to_vector(beta_rng(
                                          shapes_beta[beta_shape_pos[i]] * (    inv_logit(LP + beh_intercepts[i])),
                                          shapes_beta[beta_shape_pos[i]] * (1 - inv_logit(LP + beh_intercepts[i]))
                                          ));
        for (jj in 1:n_dyads) { // loglik
          log_lik[jj, i] = beta_lpdf(interactions_cont[jj, i] |
                                     inv_logit(LP[jj] + beh_intercepts[i])  * shapes_beta[beta_shape_pos[i]],
                                (1 - inv_logit(LP[jj] + beh_intercepts[i])) * shapes_beta[beta_shape_pos[i]]);

        }

      }

    }


  }
}
```
