## Supplementary material for "(De)composing sociality: disentangling individual-specific from dyad-specific propensities to interact": bamoso R package: workflow_example.stan

```
// this is the model for the workflow example in the manuscript
// coded for cmdstanr

data {
  int<lower=0> n_ids;
  int<lower=0> n_dyads;
  array[n_dyads, 1] int<lower=0> interactions;
  array[n_dyads, 2] int<lower=0> dyads_navi;

  matrix[1, 2] prior_matrix;
  array[n_ids] int<lower=0> y;
}

parameters {
  real<lower=0> indi_soc_sd;
  vector[n_ids] indi_soc_vals_z;
  real<lower=0> dyad_soc_sd;
  vector[n_dyads] dyad_soc_vals_z;
  vector[1] beh_intercepts;

  real intercept;
  real slope;
}

transformed parameters {
  vector[n_ids] indi_soc_vals; // individual blups
  vector[n_dyads] dyad_soc_vals;  // dyad blups
  vector[n_ids] lp_mod;  // linear predictor for the model
  indi_soc_vals = (indi_soc_sd * indi_soc_vals_z);
  dyad_soc_vals = (dyad_soc_sd * dyad_soc_vals_z);
  lp_mod = intercept + slope * indi_soc_vals;
}

model {
  vector[n_dyads] lp = rep_vector(0.0, n_dyads);
  vector[n_dyads] indi_sums = 0.5 * (indi_soc_vals[dyads_navi[, 1]] + indi_soc_vals[dyads_navi[, 2]]);
  lp = lp + indi_sums + dyad_soc_vals; // +

  interactions[, 1] ~ poisson(exp(lp + beh_intercepts[1]));
  y ~ poisson_log(lp_mod);

  // priors
  beh_intercepts[1] ~ student_t(3, prior_matrix[1, 1], prior_matrix[1, 2]);
  indi_soc_vals_z ~ normal(0, 1);
  dyad_soc_vals_z ~ normal(0, 1);
  indi_soc_sd ~ exponential(0.1);
  dyad_soc_sd ~ exponential(0.1);
  intercept ~ normal(0, 1);
  slope ~ normal(0, 1);
}

generated quantities {
   array[n_dyads, 1] int interactions_pred;
   array[n_ids] int y_rep;
   interactions_pred[, 1] = poisson_log_rng( 0.5 * (indi_soc_vals[dyads_navi[, 1]] + indi_soc_vals[dyads_navi[, 2]]) + dyad_soc_vals + beh_intercepts[1]);
   y_rep = poisson_log_rng(lp_mod);
}
```
