## Supplementary material for "(De)composing sociality: disentangling individual-specific from dyad-specific propensities to interact": bamoso R package: REFERENCES.bib

@article{hart2022,
  title = {Common permutation methods in animal social network analysis do not control for non-independence},
  author = {Hart, Jordan D A and Weiss, Michael N and Brent, Lauren J N and Franks, Daniel W},
  year = {2022},
  journal = {Behavioral Ecology and Sociobiology},
  volume = {76},
  pages = {151},
  doi = {10.1007/s00265-022-03254-x}
}

@book{mcelreath2020,
  title = {Statistical rethinking: a {{Bayesian}} course with examples in {{R}} and {{Stan}}},
  shorttitle = {Statistical rethinking},
  author = {McElreath, Richard},
  year = {2020},
  edition = {2},
  publisher = {{Chapman and Hall/CRC}},
  doi = {10.1201/9780429029608},
  isbn = {978-0-429-02960-8}
}


@book{bolker2008,
  title = {Ecological models and data in {{R}}},
  author = {Bolker, Benjamin M},
  year = {2008},
  publisher = {Princeton University Press}
}

@article{buerkner2017,
  title = {{\texttt{brms}}: An {R} package for {Bayesian} multilevel models using {Stan}},
  author = {Paul-Christian Bürkner},
  journal = {Journal of Statistical Software},
  year = {2017},
  volume = {80},
  number = {1},
  pages = {1--28},
  doi = {10.18637/jss.v080.i01}
}

@misc{mourier2020,
  title = {Kinship does not predict the structure of a shark social network},
  author = {Mourier, Johann},
  year = {2020},
  publisher = {{Dryad}},
  doi = {10.5061/DRYAD.WM37PVMKX}
}

@article{mourier2021,
  title = {Kinship does not predict the structure of a shark social network},
  author = {Mourier, Johann and Planes, Serge},
  year = {2021},
  journal = {Behavioral Ecology},
  volume = {32},
  number = {2},
  pages = {211--222},
  doi = {10.1093/beheco/araa119}
}


@article{menz2020,
  title = {Higher sociability leads to lower reproductive success in female kangaroos},
  author = {Menz, Clementine S and Carter, Alicia J and Best, Emily C and Freeman, Natalie J and Dwyer, Ross G and Blomberg, Simon P and Goldizen, Anne W},
  year = {2020},
  journal = {Royal Society Open Science},
  volume = {7},
  pages = {200950},
  doi = {10.1098/rsos.200950}
}

@misc{menz2020data,
  title = {Higher sociability leads to lower reproductive success in female kangaroos},
  author = {Carter, Alecia J and Menz, Clementine S and Best, Emily C and Freeman, Natalie J and Dwyer, Ross G and Blomberg, Simon P and Goldizen, Anne W},
  year = {2020},
  publisher = {Dryad},
  doi = {10.5061/dryad.79cnp5hsb}
}

@article{silk2006,
  title = {Social relationships among adult female baboons (\emph{{P}apio cynocephalus}) I. Variation in the strength of social bonds},
  author = {Silk, Joan B and Altmann, Jeanne and Alberts, Susan C},
  year = {2006},
  journal = {Behavioral Ecology and Sociobiology},
  volume = {61},
  pages = {183--195},
  doi = {10.1007/s00265-006-0249-2}
}

@article{silk2013a,
  title = {A practical guide to the study of social relationships},
  author = {Silk, Joan B and Cheney, Dorothy L and Seyfarth, Robert M},
  year = {2013},
  journal = {Evolutionary Anthropology},
  volume = {22},
  pages = {213--225},
  doi = {10.1002/evan.21367}
}

@article{cairns1987,
  title = {A comparison of association indices},
  journal = {Animal Behaviour},
  volume = {35},
  number = {5},
  pages = {1454--1469},
  year = {1987},
  doi = {10.1016/S0003-3472(87)80018-0},
  author = {Sara J. Cairns and Steven J. Schwager}
}

@book{whitehead2008,
  title = {Analyzing animal societies: quantitative methods for vertebrate social analysis},
  author = {Whitehead, Hal},
  year = {2008},
  publisher = {University of Chicago Press},
  address = {Chicago}
}

@article{whitehead2015,
  title = {Generalized affiliation indices extract affiliations from social network data},
  author = {Whitehead, Hal and James, Richard},
  year = {2015},
  journal = {Methods in Ecology and Evolution},
  volume = {6},
  pages = {836--844},
  doi = {10.1111/2041-210X.12383}
}

@article{godde2013,
  title = {Correcting for the impact of gregariousness in social network analyses},
  author = {Godde, Sophie and Humbert, Lionel and C{\^o}t{\'e}, Steeve D and R{\'e}ale, Denis and Whitehead, Hal},
  year = {2013},
  journal = {Animal Behaviour},
  volume = {85},
  pages = {553--558},
  doi = {10.1016/j.anbehav.2012.12.010}
}

@article{rathke2021,
  title = {Social aging in male and female {B}arbary macaques},
  author = {Rathke, Eva-Maria and Fischer, Julia},
  year = {2021},
  journal = {American Journal of Primatology},
  volume = {83},
  pages = {e23272},
  doi = {10.1002/ajp.23272}
}

@article{gabry2019,
  author = {Gabry, Jonah and Simpson, Daniel and Vehtari, Aki and Betancourt, Michael and Gelman, Andrew},
  title = {Visualization in {B}ayesian workflow},
  journal = {Journal of the Royal Statistical Society: Series A},
  volume = {182},
  number = {2},
  pages = {389--402},
  doi = {10.1111/rssa.12378},
  year = {2019}
}
