## Supplementary material for "(De)composing sociality: disentangling individual-specific from dyad-specific propensities to interact": bamoso R package: asso_indices.R

```
#' various association indices
#'
#' @param xdata a group/observation \eqn{\times} individual matrix. Needs to be
#'              without additional columns (like 'date' or 'period' etc).
#' @param loop logical, use a for-loop internally, or
#'             an \code{apply()}. Default is \code{TRUE}.
#'             Does not have consequences for the results (just speed).
#' @param out_net character of length 1. Can be one of \code{"hwi"},
#'                \code{"sri"}, \code{"hwig"}, or \code{"hwig_ori"}.
#'                If supplied a symmetric square matrix (network) is returned
#'                as output, with the supplied index. If \code{NULL}
#'                (the default), a data.frame is returned with all indices.
#' @references
#' \insertRef{cairns1987}{bamoso}
#'
#' \insertRef{whitehead2008}{bamoso}
#'
#' \insertRef{whitehead2015}{bamoso}
#'
#' @return a data frame (or matrix)
#' @export
#'
#' @examples
#' data("socprogexample")
#' # results obtained from SOCPROG 2.9
#' socres <- socprogexample$dyadicmetrics
#'
#' # raw data
#' xdata <- socprogexample$raw_asso
#'
#' # calculate indices with package function
#' ares <- asso_indices(xdata)
#'
#' # compare to original
#' # dyadic indices
#' identical(round(ares$sri, 10), round(socres$sri, 10))
#' identical(round(ares$hwi, 10), round(socres$hwi, 10))
#' identical(round(ares$hwig, 10), round(socres$hwig, 10))
#'
#' # gregariousness predictor (for generalized affiliation index)
#' identical(round(ares$greg_pred_hwi, 10),
#'           round(socres$greg_predictor_hwi, 10))
#' identical(round(ares$greg_pred_sri, 10),
#'           round(socres$greg_predictor_sri, 10))
#'


asso_indices <- function(xdata, loop = TRUE, out_net = NULL) {
  dyads <- t(combn(seq_len(ncol(xdata)), 2))
  if (!is.null(colnames(xdata))) {
    id_codes <- colnames(xdata)
  } else {
    id_codes <- seq_len(ncol(xdata))
  }

  colnames(dyads) <- c("i1", "i2")

  seen_together <- apply(dyads, 1, function(x) (sum(rowSums(xdata[, x]) == 2)))
  seen_either <- apply(dyads, 1, function(x) (sum(rowSums(xdata[, x]) >= 1)))
  seen1 <- colSums(xdata)[dyads[, 1]]
  seen2 <- colSums(xdata)[dyads[, 2]]

  hwi <- seen_together / (0.5 * (seen1 + seen2))

  hwig_out <- numeric(length(hwi))

  # loop doesn't even slow it down much...
  if (loop) {
    for (i in seq_len(nrow(dyads))) {
      hwi_a <- sum(hwi[dyads[, 1] == dyads[i, 1] | dyads[, 2] == dyads[i, 1]])
      hwi_b <- sum(hwi[dyads[, 1] == dyads[i, 2] | dyads[, 2] == dyads[i, 2]])
      hwig_out[i] <- hwi[i] * (sum(hwi) / (hwi_a * hwi_b))
    }

  } else {
    hwi_a <- apply(dyads, 1, function(x) {
      sum(hwi[dyads[, 1] == x[1] | dyads[, 2] == x[1]])
    })
    hwi_b <- apply(dyads, 1, function(x) {
      sum(hwi[dyads[, 1] == x[2] | dyads[, 2] == x[2]])
    })
    hwig_out <- hwi * (sum(hwi) / (hwi_a * hwi_b))
  }

  sri <- seen_together / seen_either

  # gregariousness predictor (Whitehead and James 2015, eq 4)
  # requires backtransform into square matrix
  # Matlab source code: genaffilcalc.m (lines 309ff)

  # assm=assocmg-diag(diag(assocmg));%zeros on diagonal
  # nuu=length(assm(1,:));
  # greg=log(sum(assm,2)*sum(assm,1)-(ones(nuu,1)*sum(assm,1)+
  #          sum(assm,2)*ones(1,nuu)-assm).*assm);
  # assocm=greg-diag(diag(greg)); %full(sum(assm,2)*sum(assm,1)/sum(assm(:)));

  # for SRI:
  m <- matrix(0, ncol = ncol(xdata), nrow = ncol(xdata))
  m[lower.tri(m)] <- sri
  m <- m + t(m)
  nvec <- rep(1, ncol(m))
  x1 <- outer(colSums(m), rowSums(m)) # matlab: sum(assm,2)*sum(assm,1)
  x2 <- outer(nvec, rowSums(m)) # matlab: ones(nuu,1)*sum(assm,1)
  x3 <- outer(colSums(m), nvec) # matlab: sum(assm,2)*ones(1,nuu)
  y <- log(x1 - (x2 + x3 - m) * m)
  greg_pred_sri <- y[lower.tri(y)]
  # fix occasional problematic cases (cf. matlab code, genaffil.m (l 313))
  if (any(is.infinite(greg_pred_sri))) {
    greg_pred_sri[is.infinite(greg_pred_sri)] <- 0
  }

  # for HWI:
  m <- matrix(0, ncol = ncol(xdata), nrow = ncol(xdata))
  m[lower.tri(m)] <- hwi
  m <- m + t(m)
  nvec <- rep(1, ncol(m))
  x1 <- outer(colSums(m), rowSums(m)) # matlab: sum(assm,2)*sum(assm,1)
  x2 <- outer(nvec, rowSums(m)) # matlab: ones(nuu,1)*sum(assm,1)
  x3 <- outer(colSums(m), nvec) # matlab: sum(assm,2)*ones(1,nuu)
  y <- log(x1 - (x2 + x3 - m) * m)
  greg_pred_hwi <- y[lower.tri(y)]
  # fix occasional problematic cases (cf. matlab code, genaffil.m (l 313))
  if (any(is.infinite(greg_pred_hwi))) {
    greg_pred_hwi[is.infinite(greg_pred_hwi)] <- 0
  }

  # create a column that allows ordering in an alternative way that reflect
  #   how dyad indices were created (either with 'combn' or with
  #   sth. like: 'which(lower.tri(m), arr.ind = TRUE)'
  alt_dyads <- which(upper.tri(matrix(0, ncol(xdata), ncol(xdata))),
                     arr.ind = TRUE)
  alt_order <- apply(dyads, 1, function(x) {
    which(alt_dyads[, 1] == x[1] & alt_dyads[, 2] == x[2])
  })


  res <- data.frame(i1 = dyads[, "i1"],
                    i2 = dyads[, "i2"],
                    i1_name = id_codes[dyads[, "i1"]],
                    i2_name = id_codes[dyads[, "i2"]],
                    alt_order = alt_order,
                    # not sure why need to multiply to replicate SOCPROG!
                    hwig = hwig_out * 2,
                    hwig_ori = hwig_out,
                    hwi = hwi,
                    sri = sri,
                    greg_pred_hwi = greg_pred_hwi,
                    greg_pred_sri = greg_pred_sri,
                    seen1 = seen1,
                    seen2 = seen2,
                    seen_together = seen_together,
                    seen_either = seen_either)
  if (is.null(out_net)) {
    return(res)
  }

  if (out_net %in% c("hwig", "hwig_ori", "sri", "hwi")) {
    outmat <- matrix(ncol = ncol(xdata), nrow = ncol(xdata), 0)
    colnames(outmat) <- id_codes
    rownames(outmat) <- id_codes
    outmat[as.matrix(res[, c("i1", "i2")])] <- res[, out_net]
    return(outmat)
  }

}
```
