## Supplementary material for "(De)composing sociality: disentangling individual-specific from dyad-specific propensities to interact": bamoso R package: check_data.R

```
#' some basic data checks
#'
#' mostly concerning dimensions and naming of input data
#'
#' @details
#' This function is very rudimentary! It only performs some very basic
#' checks for now.
#'
#' * do dimensions match within and between interaction matrices?
#'
#' * do dimensions match across interaction matrices and observation
#'   effort data?
#'
#' * if column and row names are provided: do they align across
#'   interaction matrices?
#'
#' * are there any individuals for which all interaction data values are \code{NA}
#'
#' @inheritParams make_stan_data_from_matrices
#' @importFrom stats na.omit
#' @return just some textual output
#' @export
#' @examples
#' data("grooming")
#' mats <- list(grooming$ass$groom[1:5, 1:5])
#' obseff <- outer(grooming$ass$obseff[1:5], grooming$ass$obseff[1:5], "+")
#' check_data(mats = mats, obseff = obseff)
#'
#' \dontrun{
#' data("grooming")
#' mats <- list(grooming$ass$groom[1:5, 1:5])
#' # set one value to NA
#' mats[[1]][1, 4] <- NA
#' obseff <- outer(grooming$ass$obseff[1:5], grooming$ass$obseff[1:5], "+")
#' check_data(mats = mats, obseff = obseff)
#' }
#'
#' \dontrun{
#' data("grooming")
#' mats <- list(grooming$ass$groom[1:5, 1:5])
#' # set individual to NA
#' mats[[1]][2, ] <- NA
#' mats[[1]][, 2] <- NA
#' obseff <- outer(grooming$ass$obseff[1:5], grooming$ass$obseff[1:5], "+")
#' check_data(mats = mats, obseff = obseff)
#' }


check_data <- function(mats,
                       behav_types = NULL,
                       obseff = NULL) {

  # check dimensions
  if (isFALSE(length(unique(unlist(unique(lapply(mats, dim))))) == 1)) {
    stop("mismatch in dimensions found in interaction matrices", call. = FALSE)
  } else {
    message("dimensions of interaction matrices match (good)")
  }
  # in obseff data
  if (!is.null(obseff)) {
    if (!is.list(obseff)) obseff <- list(obseff)
    do <- unique(unlist(lapply(obseff, dim)))
    if (isFALSE(length(do) == 1)) {
      stop("mismatch in dimensions found in observation effort data", call. = FALSE)
    } else {
      message("dimension of observation effort data match (good)")
    }
  }

  # check for individuals for which all values are NA
  aux <- lapply(mats, function(x) {
    diag(x) <- NA
    (rowSums(is.na(x)) + colSums(is.na(x))) == ncol(x) * 2
  })
  aux <- do.call("rbind", aux)
  if (any(aux)) {
    message("found *individuals* in interaction data for which *all* dyadic behavior values are NA.\n",
            "Those individuals will be removed in make_standata_from_matrices().")
  }


  # names of individuals in observation effort ------------
  n_obseff <- NULL
  if (!is.null(obseff)) {
    # is a list at this point
    onames <- list()
    for (i in seq_len(length(obseff))) {
      if (is.vector(obseff[[i]])) {
        if (is.null(names(obseff))) {
          onames[[length(onames) + 1]] <- rep(NA, length(obseff[[i]]))
          cat("at least one observation effort is missing names\n")
        } else {
          onames[[length(onames) + 1]] <- names(obseff[[i]])
        }
      } else {
        rn <- rownames(obseff[[i]])
        cn <- colnames(obseff[[i]])
        if (is.null(rn)) {
          onames[[length(onames) + 1]] <- rep(NA, nrow(obseff[[i]]))
          cat("at least one observation effort is missing row names\n")
        } else {
          onames[[length(onames) + 1]] <- rownames(obseff[[i]])
        }
        if (is.null(cn)) {
          onames[[length(onames) + 1]] <- rep(NA, ncol(obseff[[i]]))
          cat("at least one observation effort is missing column names\n")
        } else {
          onames[[length(onames) + 1]] <- colnames(obseff[[i]])
        }
      }
    }

    ndat <- do.call("cbind", onames)
    # compare names across the matrices/vectors
    xtest2 <- apply(ndat, 1, function(x) length(unique(na.omit(x))))
    if (isTRUE(any(xtest2 != 1))) {
      message("name mismatch found between or within observation effort data (not good)")
    } else {
      cat("all observation effort data have matching column and row names (good)\n")
    }
    n_obseff <- nrow(ndat)
  }

  # names of individuals in interaction matrices ------------
  n_ints <- NULL
  cn <- lapply(mats, colnames)
  rn <- lapply(mats, rownames)

  cn <- lapply(seq_len(length(cn)), function(x) {
    if (is.null(cn[[x]])) return(rep(NA, ncol(mats[[x]])))
    cn[[x]]
  })
  rn <- lapply(seq_len(length(rn)), function(x) {
    if (is.null(rn[[x]])) return(rep(NA, nrow(mats[[x]])))
    rn[[x]]
  })

  ndat <- do.call("cbind", c(cn, rn))
  n_ints <- nrow(ndat)
  # any dim without names?
  xtest <- apply(ndat, 2, function(x) any(is.na(x)))

  if (all(xtest)) {
    cat("no column and row names provided\n")
  } else {
    if (any(xtest)) {
      cat("at least one interaction matrix is missing column and/or row names\n")
    } else {
      # compare names across the matrices
      xtest2 <- apply(ndat, 1, function(x) length(unique(na.omit(x))))
      if (isTRUE(any(xtest2 != 1))) {
        cat("name mismatch found between matrices or between rows and columns",
            "of the same matrix\n")
      } else {
        cat("all interaction matrices have matching column and",
            "row names (good)\n")
      }
    }
  }

  tot <- unique(c(n_obseff, n_ints))
  if (length(tot) == 1) {
    message("found ", tot, " individuals in all data sources (good)")
  } else {
    stop("found different numbers of individuals across the provided ",
         "data sources", call. = FALSE)
  }


  # if it ran so far, we can check a few things about data integrity
  if (any(unlist(lapply(mats, function(x) any(is.na(x)))))) {
    message("found NA values in interaction matrices (not good)")
  }
  if (any(unlist(lapply(mats, function(x) any(x < 0, na.rm = TRUE))))) {
    message("found negative values in interaction matrices (not good)")
  }
  if (!is.null(obseff)) {
    if (any(unlist(lapply(obseff, function(x) any(is.na(x)))))) {
      message("found NA values in observation effort data (not good)")
    }
    if (any(unlist(lapply(obseff, function(x) any(x < 0, na.rm = TRUE))))) {
      message("found negative values in observation effort data (not good)")
    }
  }

  # check for proportion data (more successes than trials)
  # check for beta values outside 0-1 range
}
```
