## Supplementary material for "(De)composing sociality: disentangling individual-specific from dyad-specific propensities to interact": bamoso R package: cor_plot.R

```
#' correlation plot
#'
#' visualizes posterior samples of gregariousness or affinity values
#'
#' @param model result of \code{\link{sociality_model}}
#' @param greg logical (default is \code{TRUE}). Plot gregariousness or
#'          affinity.
#' @param axis1,axis2 name of behaviors for which the correlation should be
#'          processed
#' @param n_draws numeric, number of draws to be plotted
#' @param ... additional argument to \code{\link{plot}}
#'
#' @importFrom grDevices contourLines
#' @export
#' @return a plot
#'
#' @details
#' Two randomly selected individuals/dyads are highlighted.
#'
#'
#' @examples
#' model <- fit_example_model(num = 1)
#' summary(model)
#' cor_plot(model, asp = 1, xlab = "A", ylab = "B")
#' cor_plot(model, greg = FALSE, xlim = c(-3, 3), asp = 1)

# requires MASS package...
# @importFrom MASS kde2d


cor_plot <- function(model,
                     greg = TRUE,
                     axis1 = NULL,
                     axis2 = NULL,
                     n_draws = 200,
                     ...
                     ) {

  if (!requireNamespace("MASS", quietly = TRUE)) {
    stop("Package 'MASS' is required for this function but is not installed. ",
         "Please install it using install.packages('MASS').")
  }

  if (is.null(axis1)) {
    axis1_num <- 1
    axis1 <- names(model$standat$beh_names)[axis1_num]
  } else {
    axis1_num <- which(names(model$standat$beh_names) == axis1)
    axis1 <- names(model$standat$beh_names)[axis1_num]
  }
  if (is.null(axis2)) {
    axis2_num <- 2
    axis2 <- names(model$standat$beh_names)[axis2_num]
  } else {
    axis2_num <- which(names(model$standat$beh_names) == axis2)
    axis2 <- names(model$standat$beh_names)[axis2_num]
  }


  if (greg) {
    s1 <- extract_samples(model, "indi_vals", axis = axis1_num)
    s2 <- extract_samples(model, "indi_vals", axis = axis2_num)
  } else {
    s1 <- extract_samples(model, "dyad_vals", axis = axis1_num)
    s2 <- extract_samples(model, "dyad_vals", axis = axis2_num)
  }

  sel <- seq_len(nrow(s1))
  if (nrow(s1) > n_draws) {
    sel <- sample(nrow(s1), n_draws)
  }

  ix <- sample(colnames(s1))

  make_color <- rep(FALSE, ncol(s1))
  # make_color[sample(seq_along(make_color), 2)] <- TRUE
  make_color[c(length(make_color) - 1, length(make_color))] <- TRUE
  names(make_color) <- ix

  colors <- rep(grey(0.1, 0.2), ncol(s1))
  names(colors) <- ix
  colors[make_color] <- c(adjustcolor(c("red", "blue")))

  plot(0, 0, type = "n", ...)

  for (i in ix) {
    points(s1[sel, i], s2[sel, i], pch = ".", cex = 0.5, col = colors[i])
  }
  for (i in ix) {
    zz <- MASS::kde2d(s1[, i], s2[, i], n = 51)
    pdata <- contourLines(zz, levels = quantile(zz$z, 0.95))
    if (make_color[i]) {
      polygon(pdata[[1]]$x, pdata[[1]]$y, border = "black", lwd = 4)
    }
    polygon(pdata[[1]]$x, pdata[[1]]$y,
            border = adjustcolor(colors[i], 1), lwd = 2)
  }

  if (interactive()) {
    message(paste0("the red individual/dyad is ", shQuote(ix[make_color][1])))
    message(paste0("the blue individual/dyad is ", shQuote(ix[make_color][2])))
  }

  NULL
}
```
