## Supplementary material for "(De)composing sociality: disentangling individual-specific from dyad-specific propensities to interact": bamoso R package: extract_samples.R

```
#' posterior draws
#'
#' @param mod_res model results from
#'                \code{\link{sociality_model}})
#' @param what character, one of \code{"dyad_sd"}, \code{"indi_sd"},
#'             \code{"dyad_vals"}, \code{"indi_vals"} or
#'             \code{"beh_intercepts"}. See details
#' @param axis integer, indicates which axis is to be returned. Only relevant
#'             if the model was fitted with correlations (and at least 2
#'              behaviors).
#'
#' @return a (named) matrix with variable number of columns representing the
#'         posterior draws for the desired quantity.
#' @export
#' @details
#' The \code{what = } argument can take on one the following values:
#' \describe{
#'  \item{\code{"indi_sd"}:}{The gregariousness variation}
#'  \item{\code{"dyad_sd"}:}{The affinity variation}
#'  \item{\code{"indi_vals"}}{The gregariousness values for each individual}
#'  \item{\code{"dyad_vals"}}{The affinity values for each dyad}
#'  \item{\code{"beh_intercepts"}}{The intercepts for each behavior}
#' }
#'
#'
#' @examples
#' \dontrun{
#' mat <- generate_data(n_ids = 6, n_beh = 2,
#'                      behav_types = c("dur_gamma", "dur_beta"),
#'                      indi_sd = 2, dyad_sd = 0.5)
#' mats <- mat$processed$interaction_matrices
#' colnames(mats[[1]]) <- colnames(mats[[2]]) <- letters[1:ncol(mats[[1]])]
#' matlist <- list(groom = mats[[1]], prox = mats[[2]])
#' sdat <- make_stan_data_from_matrices(mats = matlist,
#'                                      behav_types = c("dur_gamma", "dur_beta"),
#'                                      obseff = NULL)
#' res <- sociality_model(standat = sdat, parallel_chains = 4,
#'                        iter_sampling = 1000, iter_warmup = 1000,
#'                        refresh = 0, adapt_delta = 0.9)
#' head(extract_samples(res, what = "beh_intercepts"))
#' head(extract_samples(res, what = "indi_vals")[, 1:4])
#' head(extract_samples(res, what = "dyad_vals")[, 1:4])
#'
#' # with correlations
#' data(nigra2)
#' s <- make_stan_data_from_matrices(mats = list(groom = nigra2$groom,
#'                                               prox = nigra2$prox),
#'                                   behav_types = c("count", "prop"),
#'                                   obseff = list(nigra2$obseff_groom,
#'                                                 nigra2$obseff_prox),
#'                                   correlations = TRUE)
#'
#' r <- sociality_model(s, parallel_chains = 4, seed = 17, adapt_delta = 0.9)
#' head(extract_samples(r, "indi_sd", axis = 1)) # grooming axis
#' head(extract_samples(r, "indi_sd", axis = 2)) # proximity axis
#' head(extract_samples(r, what = "indi_vals")[, 1:4])
#' head(extract_samples(r, "indi_cors"))
#' }

extract_samples <- function(mod_res,
                            what = c("indi_sd",
                                     "dyad_sd",
                                     "beh_intercepts",
                                     "indi_vals",
                                     "dyad_vals",
                                     "indi_cors",
                                     "dyad_cors"
                            ),
                            axis = 1) {

  standat <- mod_res$standat
  mod_res <- mod_res$mod_res

  if (standat$n_cors == 0) {
    if ("indi_cors" %in% what || "dyad_cors" %in% what) {
      message("model doesn't contain correlations (removing 'indi_cors' ",
              "and/or 'dyad_cors' from argument 'what = ')\n")
      what <- what[what != "indi_cors"]
      what <- what[what != "dyad_cors"]
    }
  }

  # number of draws
  nd <- mod_res$num_chains() * mod_res$metadata()$iter_sampling
  outres <- matrix(ncol = 0, nrow = nd)


  if ("indi_cors" %in% what) {
    bnames <- names(standat$beh_names)
    xindex <- which(upper.tri(matrix(ncol = standat$n_beh,
                                     nrow = standat$n_beh)),
                    arr.ind = TRUE)
    xnames <- apply(xindex, 1, function(x) {
      paste(bnames[x[1]], bnames[x[2]], sep = "_X_")
    })

    x <- mod_res$draws(variables = "cors_indi", format = "draws_matrix")
    res <- matrix(as.numeric(x), ncol = standat$n_cors)
    colnames(res) <- paste0("cor_", xnames)
    outres <- cbind(outres, res)
  }

  if ("dyad_cors" %in% what) {
    bnames <- names(standat$beh_names)
    xindex <- which(upper.tri(matrix(ncol = standat$n_beh,
                                     nrow = standat$n_beh)),
                    arr.ind = TRUE)
    xnames <- apply(xindex, 1, function(x) {
      paste(bnames[x[1]], bnames[x[2]], sep = "_X_")
    })

    x <- mod_res$draws(variables = "cors_dyad", format = "draws_matrix")
    res <- matrix(as.numeric(x), ncol = standat$n_cors)
    colnames(res) <- paste0("cor_", xnames)
    outres <- cbind(outres, res)
  }

  # for indi SD and dyad SD: default is first axis!!!

  if ("indi_sd" %in% what) {
    x <- mod_res$draws(variables = "indi_soc_sd", format = "draws_matrix")
    res <- matrix(as.numeric(x), ncol = ncol(x))
    res <- res[, axis, drop = FALSE]
    colnames(res) <- "greg_sd"
    outres <- cbind(outres, res)
  }

  if ("dyad_sd" %in% what) {
    x <- mod_res$draws(variables = "dyad_soc_sd", format = "draws_matrix")
    res <- matrix(as.numeric(x), ncol = ncol(x))
    res <- res[, axis, drop = FALSE]
    colnames(res) <- "affi_sd"
    outres <- cbind(outres, res)
  }

  if ("beh_intercepts" %in% what) {
    x <- mod_res$draws(variables = "beh_intercepts", format = "draws_matrix")
    res <- matrix(as.numeric(x), ncol = ncol(x))
    if (!is.null(standat)) {
      colnames(res) <- names(standat$beh_names)
    }
    outres <- cbind(outres, res)
  }

  if ("indi_vals" %in% what) {
    x <- mod_res$draws(variables = "indi_soc_vals", format = "draws_matrix")
    res <- matrix(as.numeric(x), ncol = ncol(x))
    if (standat$n_cors > 0) {
      res <- res[, grepl(paste0(",", axis), colnames(x))]
    }

    if (!is.null(standat)) {
      colnames(res) <- names(standat$id_codes)
    }
    outres <- cbind(outres, res)
  }

  if ("dyad_vals" %in% what) {
    x <- mod_res$draws(variables = "dyad_soc_vals", format = "draws_matrix")
    res <- matrix(as.numeric(x), ncol = ncol(x))
    if (standat$n_cors > 0) {
      res <- res[, grepl(paste0(",", axis), colnames(x))]
    }
    if (!is.null(standat)) {
      dyads <- paste(names(standat$id_codes)[standat$dyads_navi[, 1]],
                     names(standat$id_codes)[standat$dyads_navi[, 2]],
                     sep = "_@_")
      colnames(res) <- dyads
    }
    outres <- cbind(outres, res)
  }

  outres
}
```
