## Supplementary material for "(De)composing sociality: disentangling individual-specific from dyad-specific propensities to interact": bamoso R package: fit_example_model.R

```
#' fit simple example models with simple example data
#'
#' @param num an integer
#'
#' @return a model object
#' @export
#'

fit_example_model <- function(num = 1) {
  if (num == 1) {
    if (interactive()) {
      cat("12 indis, 2 behaviors, cors generated, and fitted\n")
    }
    cors_indi <- matrix(c(0.7, 0.8, 0.8, 0.8), ncol = 2)
    cors_dyad <- matrix(c(0.7, 0.3, 0.3, 1.0), ncol = 2)

    x <- generate_data(n_ids = 12, n_beh = 2, beh_intercepts = c(0.5, 0.5), behav_types = c("count", "count"), indi_sd = cors_indi, dyad_sd = cors_dyad)

    s <- x$standat

    r <- sociality_model(s, parallel_chains = 4, seed = 1, adapt_delta = 0.85, show_exceptions = FALSE, refresh = 0,
                         iter_warmup = 500, iter_sampling = 100)
    return(r)
  }


  if (num == 2) {
    if (interactive()) {
      cat("12 indis, 2 behaviors, cors generated,",
          "but *not used* during data generation\n")
    }

    cors_indi <- matrix(c(0.7, 0.8, 0.8, 0.8), ncol = 2)
    cors_dyad <- matrix(c(0.7, 0.3, 0.3, 1.0), ncol = 2)

    x <- generate_data(n_ids = 12, n_beh = 2, beh_intercepts = c(0.5, 0.5), behav_types = c("count", "count"), indi_sd = cors_indi, dyad_sd = cors_dyad)

    s <- x$standat
    s$n_cors <- 0

    r <- sociality_model(s, parallel_chains = 4, seed = 1, adapt_delta = 0.85, show_exceptions = FALSE, refresh = 0,
                         iter_warmup = 500, iter_sampling = 100)
    return(r)
  }

  if (num == 3) {
    if (interactive()) {
      cat("12 indis, 3 behaviors, cors generated, and fitted\n")
    }

    cors_indi <- matrix(c(0.7, 0.8, 0.2, 0.8, 0.9, 0.4, 0.2, 0.4, 1.2), ncol = 3)
    cors_dyad <- matrix(c(0.3, 0.8, -0.2, 0.8, 0.4, 0.4, -0.2, 0.4, 1.0), ncol = 3)

    x <- generate_data(n_ids = 12, n_beh = 3, beh_intercepts = c(0.5, 0.5, 0.5), behav_types = c("count", "count", "count"),
                       indi_sd = cors_indi, dyad_sd = cors_dyad)

    s <- x$standat
    r <- sociality_model(s, parallel_chains = 4, seed = 1, adapt_delta = 0.85, show_exceptions = FALSE, refresh = 0,
                         iter_warmup = 500, iter_sampling = 100)
    return(r)
  }

  if (num == 4) {
    if (interactive()) {
      cat("12 indis, 4 behaviors, cors generated, and fitted\n")
    }

    cors_indi <- matrix(c(1, 0.5, 0.5, -0.2, 0.5, 1, 0.5, 0.5, 0.5, 0.5, 1, 0.5, -0.2, 0.5, 0.5, 1), ncol = 4)
    cors_dyad <- matrix(c(1, 0.5, 0.5, -0.2, 0.5, 1, 0.5, 0.5, 0.5, 0.5, 1, 0.5, -0.2, 0.5, 0.5, 1), ncol = 4)

    x <- generate_data(n_ids = 12, n_beh = 4, beh_intercepts = c(0.5, 0.5, 0.5, 0.5), behav_types = c("count", "count", "count", "count"),
                       indi_sd = cors_indi, dyad_sd = cors_dyad)

    s <- x$standat
    r <- sociality_model(s, parallel_chains = 4, seed = 1, adapt_delta = 0.85, show_exceptions = FALSE, refresh = 0,
                         iter_warmup = 500, iter_sampling = 100)
    return(r)
  }

  if (num == 5) {
    if (interactive()) {
      cat("12 indis, 2 behaviors, no cors\n")
    }

    x <- generate_data(n_ids = 12, n_beh = 2,
                       beh_intercepts = c(0.5, 0.5),
                       behav_types = c("count", "prop"),
                       indi_sd = 1.2, dyad_sd = 0.8)

    s <- x$standat
    r <- sociality_model(s, parallel_chains = 4, seed = 1, adapt_delta = 0.85, show_exceptions = FALSE, refresh = 0,
                         iter_warmup = 500, iter_sampling = 100)
    return(r)
  }

  if (num == 6) {
    if (interactive()) {
      cat("12 indis, 1 behavior\n")
    }

    x <- generate_data(n_ids = 12, n_beh = 1,
                       beh_intercepts = c(0.5),
                       behav_types = c("count"),
                       indi_sd = 1.2, dyad_sd = 0.8)

    s <- x$standat
    r <- sociality_model(s, parallel_chains = 4, seed = 1, adapt_delta = 0.85, show_exceptions = FALSE, refresh = 0,
                         iter_warmup = 500, iter_sampling = 100)
    return(r)
  }


}
```
