## Supplementary material for "(De)composing sociality: disentangling individual-specific from dyad-specific propensities to interact": bamoso R package: generate_data.R

```
#' generate/simulate affiliation data
#'
#' @param n_ids numeric, number of individuals
#' @param n_beh numeric, number of behaviors
#' @param behav_types character vector of length \code{n_beh} that describes
#'        the kind of data to be generated. Possible values are \code{"prop"},
#'        \code{"count"}, \code{"dur_gamma"} and \code{"dur_beta"}.
#'        There are also experimental \code{"dur_gamma0"} and \code{"binary"}.
#' @param indi_sd numeric, the SD for the individual component. Must be
#'                positive. Default is a random value
#'                (\code{runif(1, 0.1, 2)}). Can also be a correlation/SD matrix.
#'                See details.
#' @param dyad_sd numeric, the SD for the dyadic component. Must be
#'                positive. Default is a random value
#'                (\code{runif(1, 0.1, 2)}). Can also be a correlation/SD matrix.
#'                See details.
#' @param beh_intercepts numeric of the same length as \code{n_beh}:
#'          the intercepts on the linear scale for each behavior
#'          (the group-level average). Default is random
#'          (\code{rnorm(n_beh, -2, 2)}).
#' @param beh_intercepts_add numeric of the same length as \code{n_beh}:
#'          the additional intercepts on the linear scale for those
#'          behaviors that require more than one intercept (currently
#'          only: \code{"dur_gamma0"}). Values are ignored for all
#'          other behaviors. Default \code{rnorm(n_beh, -2, 2)}.
#' @param indi_covariate_slope numeric, default is \code{NULL}. Fits an
#'          additional covariate effect on the gregariousness vector.
#'          (think: age)
#' @param indi_cat_slope numeric, default is \code{NULL}. Fits an additional
#'          categorical (binary!) effect on the gregariousness vector.
#'          (think: sex) Note: this is coded as dummy-coded z-transformed
#'          variable.
#' @param dyad_covariate_slope numeric, default is \code{NULL}. Fits an
#'          additional covariate effect on the affinity vector.
#'          (think: relatedness)
#' @param dyad_cat_slope numeric, default is \code{NULL}. Fits an additional
#'          categorical (binary!) effect on the affinity vector.
#'          (think: same-sex? yes/no) Note: this is coded as dummy-coded
#'          z-transformed variable.
#' @param prop_trials numeric of length 1 or 2, default is \code{100}, i.e.
#'          per dyad there are 100 'trials' observed. If numeric of length 2,
#'          each dyad gets its own number of trials ranging between the two
#'          values supplied. Only relevant for data generated as proportions.
#' @param disp_pars_gamma numeric of the same length as \code{n_beh}.
#'          Dispersion parameter(s) for any behavior with
#'          \code{behav_types = "dur_gamma"} (or \code{"dur_gamma0"}).
#'          See details.
#' @param disp_pars_beta numeric of the same length as \code{n_beh}.
#'          Dispersion parameter(s) for any behavior with
#'          \code{behav_types = "dur_beta"}. See details.
#' @param count_obseff numeric of length 1 or 2, default is \code{1}, i.e.
#'          per dyad there is 1 unit of observation effort (think 'hours').
#'          If numeric of length 2, each dyad get its own observation effort,
#'          sampled as uniform real between the two values. This is
#'          relevant for data generated as counts (and also \code{"dur_gamma"},
#'          \code{"dur_gamma0"} and \code{"binary"}).
#' @param exact logical, default is \code{TRUE}: should the varying intercepts
#'          for \code{indi_sd} and \code{dyad_sd} be re-scaled so that they
#'          have means of 0 and exact SD as supplied.
#' @param force_z_predictors logical, force all (if there are any) covariates
#'          (individual or dyad level features) to be z-standardized.
#'          Default is TRUE.
#'
#' @details
#' Currently, four data types/distributions via \code{behav_types}
#' are supported are \code{"count"}, \code{"prop"},
#' \code{"dur_gamma"} and \code{"dur_beta"}.
#'
#' Experimentally, there is also \code{"dur_gamma0"} which is a mixture
#' of gamma durations and Bernoulli: "duration of behavior if it occurred" and
#' a 'pure' Bernoulli ("interacted or not").
#'
#' For the dispersion parameters the input vector must be of the same
#' length and the indexing must match \code{behav_types}. For example,
#' \code{behav_types = c("count", "dur_gamma")} requires a vector of
#' length 2, e.g. \code{disp_pars_gamma = c(0, 0.6)}, where the
#' first item will be ignored, i.e. only the second entry is relevant.
#'
#' Sometimes the data generation leads to extreme interaction values.
#' For \code{behav_types = "dur_gamma"} sometimes 0's occur in
#' the final data (I suspect due to machine precision). If such cases
#' occur, the data generation will add the smallest non-zero value
#' to such dyads. For the same reason, \code{behav_types = "dur_beta"}
#' sometimes generates 0 and 1 values, which are also replaced by
#' adding (for 0s) or subtracting (for 1s) tiny random numbers. For
#' \code{behav_types = c("count", "prop")}, sometimes completely
#' empty matrices are produced (no dyad ever 'interacted'). While
#' such matrices are an interesting edge case, they represent
#' a challenge for the model with its default settings. Therefore,
#' if such cases occur, the function will return a warning.
#'
#' When the effects specified by \code{indi_covariate_slope=} and friends
#' are at their default \code{NULL}, the covariate vectors won't show
#' up in the \code{res$standata}. When the slopes are set to \code{0} (or
#' any other non-\code{NULL} value), the predictors show up in the stan data.
#'
#'
#'
#' @importFrom stats cor density rnorm rpois runif rbinom rgamma
#' @importFrom stats plogis rbeta
#' @importFrom utils combn
#' @importFrom Rdpack reprompt
#'
#' @return a list with three named items, which are lists too:
#'  \describe{
#'    \item{\code{$processed}:}{misc output generated, most importantly
#'                              interactions in matrix format}
#'    \item{\code{$standat}:}{standata to be passed to
#'                            \code{\link{sociality_model}} (see
#'                            \code{\link{make_stan_data_from_matrices}})}
#'    \item{\code{$input_data}:}{the input data supplied via arguments or
#'                               generated randomly in the function if using
#'                               the arguments' defaults}
#'  }
#'
#' @export
#' @examples
#' ex <- generate_data(n_ids = 7, n_beh = 2,
#'                     behav_types = c("count", "dur_gamma"),
#'                     indi_sd = 1.2, dyad_sd = 0.8,
#'                     disp_pars_gamma = c(0, 0.6),
#'                     beh_intercepts = c(1.4, -0.7))
#' # two interaction matrices:
#' ex$processed$interaction_matrices[[1]]
#' ex$processed$interaction_matrices[[2]]
#'
#' # with correlated behavior-specific axes
#' cors_indi <- matrix(c(1.3, 0.7, 0.7, 1.3), ncol = 2)
#' cors_dyad <- matrix(c(0.3, 0.3, 0.3, 0.3), ncol = 2)
#' ex <- generate_data(n_ids = 7, n_beh = 2,
#'                     behav_types = c("count", "dur_gamma"),
#'                     indi_sd = cors_indi,
#'                     dyad_sd = cors_dyad,
#'                     disp_pars_gamma = c(0, 0.6),
#'                     beh_intercepts = c(1.4, -0.7))
#' # two interaction matrices:
#' ex$processed$interaction_matrices[[1]]
#' round(ex$processed$interaction_matrices[[2]], 3)


# n_ids = 5
# n_beh = 3
# behav_types = c("count", "dur_gamma0", "prop")
# indi_sd = 1.2
# dyad_sd = 0.8
# indi_covariate_slope = indi_cat_slope = dyad_covariate_slope = dyad_cat_slope = NULL
# disp_pars_gamma = c(NA, 0.6, NA)
# disp_pars_beta = NULL
# beh_intercepts = c(1.4, -0.5, -0.7)
# beh_intercepts_add = c(-0.5, -0.5, -0.7)
# prop_trials = 100
# count_obseff = 1
# exact = TRUE
# force_z_predictors=TRUE


generate_data <- function(n_ids = NULL,
                          n_beh = 1,
                          behav_types = "count",
                          indi_sd = NULL,
                          dyad_sd = NULL,
                          indi_covariate_slope = NULL,
                          indi_cat_slope = NULL,
                          dyad_covariate_slope = NULL,
                          dyad_cat_slope = NULL,
                          disp_pars_gamma = NULL, # for durations as gamma
                          disp_pars_beta = NULL, # for durations as beta
                          beh_intercepts = NULL,
                          beh_intercepts_add = NULL,
                          prop_trials = 100, # number of trials for binomial
                          count_obseff = 1, # observation effort for count data
                          exact = TRUE,
                          force_z_predictors = TRUE) {

  if (isTRUE(n_ids < 4)) stop("we need at least four individuals", call. = FALSE)

  # those used to be function arguments, but were removed
  indi_intercept <- 0
  dyad_intercept <- 0

  # add predictors into data item?
  do_indi_covariate_slope <- TRUE
  do_indi_cat_slope <- TRUE
  do_dyad_covariate_slope <- TRUE
  do_dyad_cat_slope <- TRUE

  if (is.null(indi_covariate_slope)) {
    indi_covariate_slope <- 0
    do_indi_covariate_slope <- FALSE
  }
  if (is.null(indi_cat_slope)) {
    indi_cat_slope <- 0
    do_indi_cat_slope <- FALSE
  }
  if (is.null(dyad_covariate_slope)) {
    dyad_covariate_slope <- 0
    do_dyad_covariate_slope <- FALSE
  }
  if (is.null(dyad_cat_slope)) {
    dyad_cat_slope <- 0
    do_dyad_cat_slope <- FALSE
  }

  if (is.null(n_ids)) n_ids <- sample(5:30, 1)
  if (is.null(n_beh)) n_beh <- sample(1:4, 1)
  if (is.null(indi_sd)) indi_sd <- runif(1, 0.1, 2)
  if (is.null(dyad_sd)) dyad_sd <- runif(1, 0.1, 2)
  if (is.null(disp_pars_gamma)) disp_pars_gamma <- runif(n_beh, 1, 10)
  if (is.null(disp_pars_beta)) disp_pars_beta <- runif(n_beh, 5, 30)
  if (is.null(beh_intercepts)) beh_intercepts <- rnorm(n_beh, -2, 2)
  if (is.null(beh_intercepts_add)) beh_intercepts_add <- rnorm(n_beh, -2, 2)

  # prelims
  n_dyads <- n_ids * (n_ids - 1) / 2
  dyads <- which(upper.tri(diag(n_ids)), arr.ind = TRUE)
  # beh_intercepts <- as.list(beh_intercepts)

  # some checks
  # dealing with correlated sociality scales
  mulength <- 1
  if (length(indi_sd) != 1 && length(dyad_sd) != 1) {
    if (length(indi_sd) != length(dyad_sd)) stop("indi SD and dyad SD need to have the same dimensions", call. = FALSE)
    mulength <- ncol(indi_sd)
  }
  # index for columns in BLUP matrices
  colindex <- rep(1, n_beh)
  if (mulength > 1) {
    colindex <- seq_len(mulength)
  }
  if (n_beh != length(beh_intercepts)) {
    stop("require one intercept for each behavior (there is a mismatch ",
         "between n_beh and length(beh_intercepts)", call. = FALSE)
  }
  if (length(indi_sd) != length(dyad_sd)) {
    stop("indi SD and dyad SD need to have the same dimensions", call. = FALSE)
  }

  # how many columns should the SD matrices have?
  if (length(indi_sd) == 1) {
    if (n_beh > 1) {
      indi_sd_mat <- matrix(ncol = length(n_beh), nrow = length(n_beh), 1)
      diag(indi_sd_mat) <- indi_sd
      dyad_sd_mat <- matrix(ncol = length(n_beh), nrow = length(n_beh), 1)
      diag(dyad_sd_mat) <- dyad_sd
    }
    if (n_beh == 1) {
      indi_sd_mat <- matrix(indi_sd)
      dyad_sd_mat <- matrix(dyad_sd)
    }
  } else {
    # then it's matrix...
    indi_sd_mat <- indi_sd
    dyad_sd_mat <- dyad_sd
  }
  # rm(indi_sd)
  # rm(dyad_sd)

  # we can't deal generically with correlations with dur_gamma0
  # (i.e. behavior where we need to estimate more than one intercept)
  if ("dur_gamma0" %in% behav_types && length(indi_sd_mat) != 1) {
    stop("can't handle correlations with dur_gamma0")
  }

  if (min(unlist(count_obseff)) < 0) {
    stop("can't handle negative observation effort!")
  }


  # create sociality values (individual-level and dyad-level) ----

  # individual-level data
  indi_data <- data.frame(id = seq_len(n_ids),
                          feature_cat = sample(c(0, 1), n_ids, replace = TRUE),
                          feature_cont = rnorm(n_ids)
  )
  # make sure that there are at least two for each category
  indi_data$feature_cat[sample(seq_len(n_ids), 4)] <- c(1, 1, 0, 0)
  if (force_z_predictors) {
    indi_data$feature_cat <- as.numeric(scale(indi_data$feature_cat))
    indi_data$feature_cont <- as.numeric(scale(indi_data$feature_cont))
  }

  # individual sociality
  indi_soc_vals_ini <- rnorm_multi(n = n_ids,
                                   mu = rep(0, mulength),
                                   Sigma = indi_sd_mat,
                                   empirical = exact)
  indi_soc_vals <- indi_soc_vals_ini +
                   indi_cat_slope * indi_data$feature_cat +
                   indi_covariate_slope * indi_data$feature_cont
  indi_data$indi_soc_vals_ini <- indi_soc_vals_ini
  indi_data$indi_soc_vals <- indi_soc_vals
  # just checking:
  # summary(lm(indi_soc_vals ~ 0 + feature_cat + feature_cont, indi_data))
  # summary(lm(I(indi_soc_vals[, 1]) ~ 0 + feature_cat + feature_cont, indi_data))


  # dyad-level data
  dyad_data <- data.frame(dyad = seq_len(n_dyads),
                          id1 = dyads[, 1],
                          id2 = dyads[, 2],
                          feature_cat = sample(c(0, 1), n_dyads, replace = TRUE),
                          feature_cont = rnorm(n_dyads)
  )
  # make sure that there are at least two for each category
  dyad_data$feature_cat[sample(seq_len(n_dyads), 4)] <- c(1, 1, 0, 0)
  if (force_z_predictors) {
    dyad_data$feature_cat <- as.numeric(scale(dyad_data$feature_cat))
    dyad_data$feature_cont <- as.numeric(scale(dyad_data$feature_cont))
  }

  # dyadic sociality
  dyad_soc_vals_ini <- rnorm_multi(n = n_dyads,
                                   mu = rep(0, mulength),
                                   Sigma = dyad_sd_mat,
                                   empirical = exact)
  # dyad_soc_vals <- dyad_soc_vals_ini
  dyad_soc_vals <- dyad_soc_vals_ini +
                   dyad_cat_slope * dyad_data$feature_cat +
                   dyad_covariate_slope * dyad_data$feature_cont
  dyad_data$dyad_soc_vals_ini <- dyad_soc_vals_ini
  dyad_data$dyad_soc_vals <- dyad_soc_vals

  # just checking:
  # summary(lm(dyad_soc_vals ~ 0 + feature_cat + feature_cont, dyad_data))

  # select behavior scale types, currently:
  #  - count/frequency (as in poisson)
  #  - proportion (as in proportion): still technically a count (out of 100 'trials')
  #  - dur_gamma (continuous duration)
  #  - dur_beta (continuous proportion)

  if (length(behav_types) == 1) {
    if (behav_types == "random") {
      btypes <- sample(c("count", "prop", "dur_gamma"), n_beh, replace = TRUE)
    } else {
      btypes <- rep(behav_types, n_beh)
    }
  }


  if (length(behav_types) > 1) {
    if (length(behav_types) != n_beh) stop("number of behavior types does not match number of behaviors")
    if (any(behav_types == "random")) stop("can't have random behavior type with more than one behavior (need to specify for each behavior individually)")
    btypes <- behav_types
  }
  behav_types <- btypes

  # and prep for output (convert to numeric)
  # default is count/frequency
  behav_types_num <- rep(1, n_beh) # counts
  behav_types_num[btypes == "prop"] <- 2
  behav_types_num[btypes == "dur_gamma"] <- 3
  behav_types_num[btypes == "dur_beta"] <- 4
  behav_types_num[btypes == "dur_gamma0"] <- 5
  behav_types_num[btypes == "binary"] <- 6

  # indexing for optional shape/dispersion parameters
  gamma_shape <- logical(n_beh)
  gamma_shape[btypes == "dur_gamma"] <- TRUE
  gamma_shape[btypes == "dur_gamma0"] <- TRUE
  gamma_shape_pos <- numeric(n_beh)

  beta_shape <- logical(n_beh)
  beta_shape[btypes == "dur_beta"] <- TRUE
  beta_shape_pos <- numeric(n_beh)


  # temporary observation effort
  obseff <- matrix(ncol = n_beh, nrow = n_dyads, 1)
  for (i in seq_len(n_beh)) {
    if (btypes[i] == "prop") {
      if (length(prop_trials) == 1) {
        obseff[, i] <- prop_trials
      } else {
        obseff[, i] <- sample(as.integer(prop_trials[1]):as.integer(prop_trials[2]),
                              size = n_dyads, replace = TRUE)
      }
    }
    if (btypes[i] %in% c("count", "dur_gamma", "dur_gamma0", "binary")) {
      if (length(count_obseff) == 1) {
        obseff[, i] <- count_obseff
      } else {
        obseff[, i] <- runif(n = n_dyads,
                             min = count_obseff[1],
                             max = count_obseff[2])
      }
    }
  }

  interactions <- matrix(ncol = n_beh, nrow = n_dyads)
  for (i in seq_len(n_beh)) {
    lp <- indi_intercept + dyad_intercept +
      sqrt(0.5) * (indi_soc_vals[dyads[, 1], colindex[i]] + indi_soc_vals[dyads[, 2], colindex[i]]) +
      dyad_soc_vals[, colindex[i]]

    if (btypes[i] == "dur_gamma0") {
      lp_bern <- lp + beh_intercepts[i] # first is for bern
      # probvec <- 1 - exp(-(exp(lp_bern) * obseff[, i]))
      lb_gamma <- lp + beh_intercepts_add[i] # second/additional intercept is for gamma
      lb_gamma <- lb_gamma + log(obseff[, i])

      aux_y1 <- rbinom(n = n_dyads, size = 1, prob = lin2prob(lp_bern, obseff[, i]))
      # aux_y1 <- rbinom(n = n_dyads, size = 1, prob = plogis(lp_bern))
      aux_y2 <- rgamma(n = n_dyads, shape = disp_pars_gamma[i], rate = disp_pars_gamma[i]/exp(lb_gamma))
      interactions[, i] <- aux_y1 * aux_y2
      gamma_shape_pos[i] <- i

      if (all(aux_y1 == 1)) {
        warning("'dur_gamma0' with only non-zero values generated (this might not be a problem)")
      }
      if (all(aux_y1 == 0)) {
        warning("'dur_gamma0' with only zero-values generated (this is gonna be a problem for fitting the default model)")
      }
    }
    if (btypes[i] == "binary") {
      lp_bern <- lp + beh_intercepts[i]
      interactions[, i] <- rbinom(n = n_dyads, size = 1, prob = lin2prob(lp_bern, obseff[, i]))
    }


    lp_b <- lp + unlist(beh_intercepts[i])[1]
    if (btypes[i] == "count") {
      interactions[, i] <- rpois(n = n_dyads,
                                 lambda = exp(lp_b + log(obseff[, i])))
    }
    if (btypes[i] == "prop") {
      interactions[, i] <- rbinom(n = n_dyads,
                                  size = obseff[, i],
                                  prob = exp(lp_b) / (1 + exp(lp_b))) # inverse logit...
    }
    if (btypes[i] == "dur_gamma") {
      # use same parameterization as in dur_gamma0
      lb_gamma <- lp + beh_intercepts[i]
      lb_gamma <- lb_gamma + log(obseff[, i])
      interactions[, i] <- rgamma(n = n_dyads, shape = disp_pars_gamma[i], rate = disp_pars_gamma[i]/exp(lb_gamma))
      gamma_shape_pos[i] <- i
    }

    if (btypes[i] == "dur_beta") {
      # first translate mean (and variance) into shape1 and shape2 (for rbeta())
      lp_offs <- lp_b
      # mean needs to be on logit (positive [0-1]) scale
      mu <- plogis(lp_offs)
      phi <- disp_pars_beta[i]

      interactions[, i] <- rbeta(n = n_dyads,
                                 shape1 = mu * phi,
                                 shape2 = (1 - mu) * phi)
      # make sure no 0 and 1 in the results
      if (any(interactions[, i] == 1)) {
        sel <- which(interactions[, i] == 1)
        interactions[sel, i] <- interactions[sel, i] - runif(length(sel), 0, 0.00001)
      }
      if (any(interactions[, i] == 0)) {
        sel <- which(interactions[, i] == 0)
        interactions[sel, i] <- interactions[sel, i] + runif(length(sel), 0, 0.00001)
      }
      beta_shape_pos[i] <- i
    }

  }

  # create default priors for behavior intercepts
  # at this point 'interactions' is not yet split into discrete vs continuous
  # we need a second set for behaviors with more than one mean parameter
  prior_matrix <- matrix(ncol = 2, nrow = n_beh)
  prior_matrix2 <- matrix(ncol = 2, nrow = n_beh, 0)
  for (i in 1:n_beh) {
    response <- interactions[, i]
    prior_matrix[i, ] <- make_prior(response = response,
                                    type = behav_types[i],
                                    obseff = obseff[, i])
    # currently only relevant for dur_gamma0
    prior_matrix2[i, ] <- make_prior(response = response,
                                     type = behav_types[i],
                                     obseff = obseff[, i],
                                     second = TRUE)
  }

  # dsi
  aux <- interactions / obseff
  aux <- apply(aux, 2, function(y) y / mean(y))
  dsi <- rowMeans(aux)

  # sum of top 3
  sum_top3 <- numeric(n_ids)
  for (i in seq_len(n_ids)) {
    x <- dsi[apply(dyads, 1, function(x) any(x == i))]
    sum_top3[i] <- sum(sort(x, decreasing = TRUE)[1:3])
  }

  # interaction and effort matrices
  imats <- vector("list", n_beh)
  omats <- vector("list", n_beh)
  for (i in seq_len(n_beh)) {
    m <- matrix(ncol = n_ids, nrow = n_ids, 0)
    o <- matrix(ncol = n_ids, nrow = n_ids, 0)
    for (k in seq_len(nrow(dyads))) {
      m[dyads[k, 1], dyads[k, 2]] <- interactions[k, i]
      o[dyads[k, 1], dyads[k, 2]] <- obseff[k, i]
    }
    imats[[i]] <- m
    omats[[i]] <- o
  }

  # flag for when there is an empty interaction matrix generated
  empty <- any(unlist(lapply(imats, sum)) == 0)
  if (empty) {
    if (interactive()) {
      warning("at least one of the matrices is empty",
              call. = FALSE)
    } else {
      message("at least one of the matrices is empty")
    }
  }
  #
  gamma_shape_pos_mod <- numeric(n_beh)
  gamma_shape_pos_mod[gamma_shape_pos > 0] <- seq_len(sum(gamma_shape_pos > 0))
  beta_shape_pos_mod <- numeric(n_beh)
  beta_shape_pos_mod[beta_shape_pos > 0] <- seq_len(sum(beta_shape_pos > 0))

  # two tweaks:
  #   behav_types needs to be of at least length 2 (so it can be declared
  #               a non-scalar for stan): hence a 0 is appended
  #   obseff_int needs to be declared as an integer matrix so that it
  #              can account for binomial trials

  # create placeholder id names
  id_codes <- as.character(seq_len(n_ids))
  vec <- rep(0, n_ids)
  names(vec) <- id_codes

  # code behave_types as named vector
  bdata <- c(behav_types_num, 0)
  names(bdata) <- c(behav_types, "0")

  # and dyads which were removed (doesn't apply here, but for consistency)
  # index <- which(upper.tri(imats[[1]]), arr.ind = TRUE)
  # removed_dyads <- index[is.na(interactions[, 1]), , drop = FALSE]
  removed_dyads <- which(upper.tri(matrix(1)), arr.ind = TRUE)


  standat <- list(id1 = dyads[, 1],
                  id2 = dyads[, 2],
                  behav_types = bdata,
                  interactions = apply(interactions, 2, as.integer),
                  interactions_cont = apply(interactions, 2, as.numeric),
                  n_beh = as.integer(n_beh),
                  n_ids = as.integer(n_ids),
                  n_dyads = as.integer(n_dyads),
                  dyads_navi = as.matrix(dyads[, 1:2]),
                  obseff = obseff,
                  prior_matrix = prior_matrix,
                  prior_matrix2 = prior_matrix2, # second set of priors (for now only relevant for dur_gamma0)
                  obseff_int = apply(obseff, 2, as.integer),
                  gamma_shape_pos = gamma_shape_pos_mod,
                  gamma_shape_n = sum(gamma_shape_pos > 0),
                  beta_shape_pos = beta_shape_pos_mod,
                  beta_shape_n = sum(beta_shape_pos > 0),
                  id_codes = vec,
                  beh_names = rep(0, n_beh),
                  removed_dyads = removed_dyads,
                  n_cors = 0,
                  prior_only = 0
                  )

  # standat$indi_cat_pred <- 0
  # standat$indi_covariate_pred <- 0
  # standat$dyad_cat_pred <- 0
  # standat$dyad_covariate_pred <- 0

  # if slopes are not specified, don't append predictors to stan data
  if (do_indi_cat_slope) {
    standat$indi_cat_pred       <- indi_data$feature_cat
  }
  if (do_indi_covariate_slope) {
    standat$indi_covariate_pred <- indi_data$feature_cont
  }
  if (do_dyad_cat_slope) {
    standat$dyad_cat_pred       <- dyad_data$feature_cat
  }
  if (do_dyad_covariate_slope) {
    standat$dyad_covariate_pred <- dyad_data$feature_cont
  }


  if (is.null(names(behav_types))) {
    names(standat$beh_names) <- paste0("behav_", LETTERS[seq_len(n_beh)])
  } else {
    names(standat$beh_names) <- names(behav_types)
  }


  if (length(indi_sd) > 1) {
    standat$n_cors <- sum(upper.tri(indi_sd))
  }

  list(standat = standat,
       input_data = list(behav_type = btypes,
                         empty = empty,
                         indi_data = indi_data,
                         # indi_soc_vals = indi_soc_vals,
                         indi_sums = sqrt(0.5) * (indi_soc_vals[dyads[, 1]] +
                                                  indi_soc_vals[dyads[, 2]]),
                         # dyad_soc_vals = dyad_soc_vals,
                         dyads = dyads,
                         indi_sd = indi_sd,
                         # indi_intercept = indi_intercept,
                         dyad_sd = dyad_sd,
                         # dyad_intercept = dyad_intercept,
                         # dyadic_covariate_predictor = dyadic_covariate_predictor,
                         dyad_data = dyad_data,
                         indi_cat_slope = indi_cat_slope,
                         indi_covariate_slope = indi_covariate_slope,
                         # indi_pred_intercept = indi_pred_intercept,
                         dyad_cat_slope = dyad_cat_slope,
                         dyad_covariate_slope = dyad_covariate_slope,
                         # dyad_pred_intercept = dyad_pred_intercept,
                         disp_pars_gamma = disp_pars_gamma,
                         disp_pars_beta = disp_pars_beta,
                         beh_intercepts = beh_intercepts,
                         beh_intercepts_add = beh_intercepts_add,
                         obseff = obseff),
       processed = list(interaction_cor = ifelse(all(colSums(interactions) > 0),
                                                 cor(interactions, method = "s"),
                                                 NA),
                        dsi = dsi,
                        sum_top3 = sum_top3,
                        interaction_matrices = imats,
                        obseff_matrices = omats))

}
```
