## Supplementary material for "(De)composing sociality: disentangling individual-specific from dyad-specific propensities to interact": bamoso R package: get_model.R

```
#' wrapper to provide model executable
#'
#' @param type character, specify which model
#' @return a cmdstan model object
#' @export
#'
#' @details
#' \code{type} can be one of
#'
#' \itemize{
#'   \item \code{"simple"}: the basic model as described in the manuscript
#'   \item \code{"sans_dyadic"}: the basic model without the dyadic component
#'   \item \code{"cor_mod"}: the model fitting separate axes for each behavior
#'                           and also the correlations among them
#'   \item \code{"with_preds"}: the model with an individual- and dyad-level
#'                             predictor
#' }
#'
#'
#' @importFrom cmdstanr cmdstan_model
#' @importFrom Rdpack reprompt

get_model <- function(type = c("simple",
                               "with_preds",
                               "cor_mod",
                               "sans_dyadic")) {

  if (missing(type)) type <- "simple"
  if (!type %in% c("simple", "with_preds", "cor_mod", "sans_dyadic")) {
    stop ("couldn't determine which model to get...", call. = FALSE)
  }

  if (type == "simple") {
    f <- system.file("extdata/interaction_model.stan", package = "bamoso")
    mod <- cmdstan_model(stan_file = f, compile = TRUE, dir = NULL)
  }
  if (type == "with_preds") {
    f <- system.file("extdata/interaction_model_withpreds.stan", package = "bamoso")
    mod <- cmdstan_model(stan_file = f, compile = TRUE, dir = NULL)
  }
  if (type == "cor_mod") {
    f <- system.file("extdata/interaction_model_corr.stan", package = "bamoso")
    mod <- cmdstan_model(stan_file = f, compile = TRUE, dir = NULL)
  }
  if (type == "sans_dyadic") {
    f <- system.file("extdata/interaction_model_sans_dyadic.stan", package = "bamoso")
    mod <- cmdstan_model(stan_file = f, compile = TRUE, dir = NULL)
  }


  mod
}
```
