## Supplementary material for "(De)composing sociality: disentangling individual-specific from dyad-specific propensities to interact": bamoso R package: make_prior.R

```
#' set prior distributions for intercept parameters
#'
#' @param response numeric vector with the response data
#' @param type character. Possible values are \code{"prop"},
#'        \code{"count"}, \code{"dur_gamma"} and \code{"dur_beta"}.
#' @param obseff numeric vector with offset values for \code{"count"}
#'        and \code{"dur_gamma"} data, or with number of 'trials' for
#'        \code{"prop"}
#' @param second logical (default is \code{FALSE}) for flagging generating
#'          second set of priors (currently only relevant for
#'          \code{"dur_gamma0"})
#'
#' @details The resulting values can be used to set priors for the intercept
#'          values in the form of \code{student_t(3, location, scale)}.
#'          The values produced here are fairly similar to the default ones
#'          produced by \code{brms}.
#'
#' @return two values for mean (location) and spread (scale).
#' @importFrom stats mad median qlogis
#' @importFrom brms get_prior rstudent_t
#' @export
#'
#' @examples
#' s <- generate_data(n_ids = 10, n_beh = 1, behav_types = "count",
#'                    beh_intercepts = -3, indi_sd = 0.1, dyad_sd = 0.1)
#' xdata <- data.frame(response = s$standat$interactions[, 1],
#'                     obseff = s$standat$obseff[, 1])
#' make_prior(xdata$response, type = "count", obseff = xdata$obseff)
#' brms::get_prior(response ~ 1, data = xdata, family = poisson())
#'
#' s <- generate_data(n_ids = 10, n_beh = 1, behav_types = "count",
#'                    beh_intercepts = 4, indi_sd = 2, dyad_sd = 2)
#' xdata <- data.frame(response = s$standat$interactions[, 1],
#'                     obseff = s$standat$obseff[, 1])
#' make_prior(xdata$response, type = "count", obseff = xdata$obseff)
#' brms::get_prior(response ~ 1, data = xdata, family = poisson())


make_prior <- function(response, type, obseff = NULL, second = FALSE) {

  # secondary intercepts:
    # - dur_gamma0: gamma part
  if (second) {
    location <- 0
    scale <- 0
    if (type == "dur_gamma0") {
      if (!is.null(obseff)) obseff <- obseff[response > 0]
      response <- response[response > 0]
      if (is.null(obseff)) obseff <- rep(1, length(response))
      if (any(obseff == 0, na.rm = TRUE)) {
        stop("can't have any 0s in the observation effort for a 'dur_gamma0' behaviour", call. = FALSE)
      }
      lresp <- log(response / obseff)
      location <- round(median(lresp), 1)
      if (is.infinite(location)) {
        stop("couldn't set a prior (prior mean is infinite)", call. = FALSE)
      }
      scale <- round(mad(lresp), 1)
      if (isTRUE(scale < 2.5)) scale <- 2.5
    }
    return(c(location = location, scale = scale))
  }

  if (type == "dur_gamma0" || type == "binary") {
    scale <- 2.5
    binresp <- mean(as.numeric(response > 0))
    meanobseff <- 1
    if (!is.null(obseff)) meanobseff <- mean(obseff)
    location <- round(prob2lin(binresp, meanobseff), 1)
  }

  if (type == "dur_gamma") {
    if (any(obseff == 0, na.rm = TRUE)) {
      stop("can't have any 0s in the observation effort for 'gamma' behaviour",
           call. = FALSE)
    }
    if (is.null(obseff)) obseff <- rep(1, length(response))
    if (any(response == 0)) response <- response + 0.1
    lresp <- log(response / obseff)
    location <- round(median(lresp), 1)
    scale <- round(mad(lresp), 1)
    if (scale < 2.5) scale <- 2.5
  }


  if (type == "count") {
    if (any(obseff == 0, na.rm = TRUE)) {
      stop("can't have any 0s in the observation effort for 'count' behaviour",
           call. = FALSE)
    }
    if (is.null(obseff)) obseff <- rep(1, length(response))
    if (any(response == 0)) response <- response + 0.1
    lresp <- log(response / obseff)
    location <- round(median(lresp), 1)
    scale <- round(mad(lresp), 1)
    if (scale < 2.5) scale <- 2.5
  }

  if (type == "prop" || type == "dur_beta") {
    if (is.null(obseff) && type == "prop") {
      stop("require number of 'trials' supplied as observation effort")
    }

    scale <- 2.5
    if (type == "prop") {
      if (any(obseff == 0, na.rm = TRUE)) {
        stop("can't have any 0s in the observation effort for a 'prop' behaviour", call. = FALSE)
      }
      if (any(response / obseff > 1, na.rm = TRUE)) {
        stop("can't have any values in 'prop' behaviour where the count is larger than the observation effort", call. = FALSE)
      }
      lresp <- round(qlogis(response / obseff), 1)
    }
    if (type == "dur_beta") {
      lresp <- round(qlogis(response), 1)
    }
    if (any(is.infinite(lresp))) lresp[is.infinite(lresp)] <- 2.5
    location <- round(median(lresp), 1)
    if (location < -2.5) location <- -2.5
    # if (is.infinite(location)) location <- 2.5
  }


  c(location = location, scale = scale)
}
```
