## Supplementary material for "(De)composing sociality: disentangling individual-specific from dyad-specific propensities to interact": bamoso R package: make_stan_data_from_association.R

```
#' convert group-by-individual association data into data list ready for Stan
#'
#' @param asso_table a 0/1 matrix where columns correspond to individuals and
#'        rows to individual samples
#' @param indi_cat_pred vector with binary individual-
#'                      level predictor. Must contain one value per individual
#'                      and can only reflect two categories (as 0's and 1's)!
#'                      Examples are sex (female or not female), age (old or
#'                      not old), residence status (resident or migrant).
#'                      Default is \code{NULL} and if so, will occur as
#'                      \code{NULL} in the output object.
#' @param indi_covariate_pred vector with a continuous predictor
#'          on individual level. Default is \code{NULL} and if so,
#'          will occur as \code{NULL} in the output object.
#' @param dyad_cat_pred,dyad_covariate_pred vector with dyad-level predictors.
#'          Either binary, or continuous. Default is \code{NULL} and if so,
#'          will occur as \code{NULL} in the output object.
#'
#' @details For now differences in sampling periods are ignored: each
#'          individual is assumed to be present (theoretically observable)
#'          in each sample.
#'
#'          If supplied via column names in \code{asso_table}, id codes are
#'          present in the output as names of vector of zeros.
#'
#' @return a list
#' @export
#'
#' @examples
#' # generate some data
#' x <- sim_data_whitehead2015(n_ind = 5, n_periods = 9, ignore_temporal = TRUE)
#' assos <- x$assolist
#' head(assos)
#'
#' make_stan_data_from_association(assos)
#'


make_stan_data_from_association <- function(asso_table,
                                            indi_cat_pred = NULL,
                                            indi_covariate_pred = NULL,
                                            dyad_cat_pred = NULL,
                                            dyad_covariate_pred = NULL
) {
  # get dyadic counts
  ares <- asso_indices(asso_table)

  n_ids <- length(unique(c(ares[, "i1"], ares[, "i2"])))
  n_dyads <- n_ids * (n_ids - 1) * 0.5

  if (!is.null(indi_cat_pred)) {
    if (length(indi_cat_pred) != n_ids) {
      stop("individual-level predictor doesn't have correct length", call. = FALSE)
    }
  } else {
    indi_cat_pred <- rep(0, n_ids)
  }
  if (!is.null(indi_covariate_pred)) {
    if (length(indi_covariate_pred) != n_ids) {
      stop("individual-level predictor doesn't have correct length", call. = FALSE)
    }
  } else {
    indi_covariate_pred <- rep(0, n_ids)
  }

  if (!is.null(dyad_cat_pred)) {
    if (length(dyad_cat_pred) != n_dyads) {
      if (length(as.numeric(as.matrix(dyad_cat_pred))) != (n_ids ^ 2)) {
        stop("dyad-level predictor doesn't have correct dimensions", call. = FALSE)
      }
    }
  } else {
    dyad_cat_pred <- rep(0, n_dyads)
  }
  if (!is.null(dyad_covariate_pred)) {
    if (length(dyad_covariate_pred) != n_dyads) {
      if (length(as.numeric(as.matrix(dyad_covariate_pred))) != (n_ids ^ 2)) {
        stop("dyad-level predictor doesn't have correct dimensions", call. = FALSE)
      }
    }
  } else {
    dyad_covariate_pred <- rep(0, n_dyads)
  }


  interactions <- matrix(ncol = 1, nrow = n_dyads)
  obseff_dat <- matrix(ncol = 1, nrow = n_dyads)
  m <- matrix(ncol = n_ids, nrow = n_ids)
  index <- which(upper.tri(m), arr.ind = TRUE)

  for (k in seq_len(n_dyads)) {
    xline <- index[k, 1] == ares[, "i1"] & index[k, 2] == ares[, "i2"]
    interactions[k, 1] <- ares[xline, "seen_together"]
    obseff_dat[k, 1] <- ares[xline, "seen_either"]
  }

  if (!is.null(colnames(asso_table))) {
    id_codes <- as.character(colnames(asso_table))
  } else {
    id_codes <- as.character(seq_len(n_ids))
  }
  if (all(id_codes == "")) {
    id_codes <- as.character(seq_len(n_ids))
  }
  vec <- rep(0, n_ids)
  names(vec) <- id_codes

  # make prior
  prior_matrix <- matrix(ncol = 2, nrow = 1)
  prior_matrix[1, ] <- make_prior(response = interactions[, 1],
                                  obseff = obseff_dat[, 1],
                                  type = "prop")
  vec_behaviors <- c(assoc = 0)

  # code behave_types as named vector
  bdata <- c(2, 0) # prop
  names(bdata) <- c("prop", "0")

  # and dyads which were removed (doesn't apply here, but for consistency)
  removed_dyads <- which(upper.tri(matrix(1)), arr.ind = TRUE)

  out <- list(id1 = index[, 1],
              id2 = index[, 2],
              interactions = apply(interactions, 2, as.integer),
              interactions_cont = apply(interactions, 2, as.numeric),
              n_beh = as.integer(1),
              n_ids = as.integer(n_ids),
              n_dyads = as.integer(n_dyads),
              dyads_navi = as.matrix(index[, 1:2]),
              gamma_shape_pos = 0,
              gamma_shape_n = 0,
              beta_shape_pos = 0,
              beta_shape_n = 0,
              indi_cat_pred = 0, # binary
              indi_covariate_pred = 0, # continuous
              dyad_cat_pred = 0,
              dyad_covariate_pred = 0,
              obseff = obseff_dat,
              obseff_int = apply(obseff_dat, 2, as.integer),
              prior_matrix = prior_matrix,
              prior_matrix2 = prior_matrix - prior_matrix, # just placeholder to make stan data 'complete'
              id_codes = vec,
              beh_names = vec_behaviors,
              removed_dyads = removed_dyads,
              behav_types = bdata,
              n_cors = 0,
              prior_only = 0
  )

  if (!is.null(indi_cat_pred)) {
    out$indi_cat_pred <- indi_cat_pred
  }
  if (!is.null(indi_covariate_pred)) {
    out$indi_covariate_pred <- indi_covariate_pred
  }
  if (!is.null(dyad_cat_pred)) {
    if (is.matrix(dyad_cat_pred)) {
      if (ncol(dyad_cat_pred) != nrow(dyad_cat_pred)) stop("dyad_cat_pred is not square!", call. = FALSE)
      temp <- numeric(n_dyads)
      for (k in seq_len(n_dyads)) {
        temp[k] <- dyad_cat_pred[index[k, 1], index[k, 2]]
      }
      dyad_cat_pred <- temp
    }
    out$dyad_cat_pred <- dyad_cat_pred
  }
  if (!is.null(dyad_covariate_pred)) {
    if (is.matrix(dyad_covariate_pred)) {
      if (ncol(dyad_covariate_pred) != nrow(dyad_covariate_pred)) stop("dyad_covariate_pred is not square!", call. = FALSE)
      temp <- numeric(n_dyads)
      for (k in seq_len(n_dyads)) {
        temp[k] <- dyad_covariate_pred[index[k, 1], index[k, 2]]
      }
      dyad_covariate_pred <- temp
    }

    out$dyad_covariate_pred <- dyad_covariate_pred
  }

  out
}
```
