## Supplementary material for "(De)composing sociality: disentangling individual-specific from dyad-specific propensities to interact": bamoso R package: make_stan_data_from_matrices.R

```
#' convert interaction matrices into data list ready for Stan
#'
#' @param mats list with (named) square interaction matrices
#' @param obseff numeric vector with observation effort. Must be positive
#'               and one value for each individual, e.g. focal observation
#'               time or number of samples. Can also be a square matrix in which
#'               dyadic observation time is already coded for each dyad.
#'               Can also be a list of such vectors or matrices, which then
#'               is mapped to each behavior (in that case this list needs to be
#'               of the same length as \code{mats}).
#' @param behav_types character vector of length \code{n_beh} that describes
#'                    the kind of data. Possible values are \code{"prop"},
#'                    \code{"count"}, \code{"dur_gamma"} and \code{"dur_beta"}.
#'                    At its default all behaviors are considered
#'                    to be \code{"count"}. There is also experimental support
#'                    for \code{"dur_gamma0"} (zero augmented durations
#'                    ("mixture")) and \code{"binary"}, i.e. simple binary
#'                    'associations'/interactions.
#' @param indi_cat_pred vector with binary individual-
#'                      level predictor. Must contain one value per individual
#'                      and can only reflect two categories (as 0's and 1's)!
#'                      Examples are sex (female or not female), age (old or
#'                      not old), residence status (resident or migrant).
#'                      Default is \code{NULL} and if so, will occur as
#'                      \code{NULL} in the output object.
#' @param indi_covariate_pred vector with a continuous predictor
#'          on individual level. Default is \code{NULL} and if so,
#'          will occur as \code{NULL} in the output object.
#' @param dyad_cat_pred,dyad_covariate_pred vector with dyad-level predictors.
#'          Either binary, or continuous. Default is \code{NULL} and if so,
#'          will occur as \code{NULL} in the output object.
#'
#' @param correlations logical, default is \code{FALSE}. Flags whether
#'          subsequent model should estimate correlations between individuals
#'          and dyadic axes. This only makes sense if there are at least
#'          two sampled behaviors. If one or more behaviors are
#'          \code{"dur_gamma0"} correlations are not supported yet and this
#'          results in an error if set to \code{TRUE}.
#'
#' @details If supplied via column names in \code{mats[[1]]}, id codes are
#'          present in the output as names of vector of zeros.
#'
#'
#'
#' @importFrom cmdstanr cmdstan_model
#' @return a named list that can be handed over to Stan as \code{data} via
#'         \code{\link{sociality_model}}:
#'  \itemize{
#'     \item \code{$id1} integer index of individual 1
#'     \item \code{$id2} integer index of individual 2
#'     \item \code{$interactions} matrix of interactions (integer)
#'     \item \code{$interactions_cont} matrix of interactions (continuous)
#'     \item \code{$n_beh} integer number of behaviors
#'     \item \code{$n_ids} integer number of individuals
#'     \item \code{$n_dyads} integer number of dyads
#'     \item \code{$dyads_navi} integer index dyads
#'     \item \code{$obseff} matrix with continuous observation effort
#'     \item \code{$obseff_int} matrix with integer observation effort
#'     \item \code{$gamma_shape_n} number of response vectors with gamma
#'                                       likelihood
#'     \item \code{$gamma_shape_pos} integer index for responses with gamma
#'                                         likelihood
#'     \item \code{$beta_shape_n} number of response vectors with beta
#'                                      likelihood
#'     \item \code{$beta_shape_pos} integer index for responses with
#'                                        beta likelihood
#'     \item \code{$prior_matrix} matrix with priors for intercepts
#'     \item \code{$prior_matrix2} matrix with priors for additional intercepts
#'     \item \code{$generate_predictions} currently unused
#'     \item \code{$id_codes} vector of 0's with names corresponding to
#'                                  individuals' codes
#'     \item \code{$beh_names} vector of 0's with names corresponding to
#'                                   behavior codes
#'     \item \code{$indi_cat_pred} vector of 0's and 1's reflecting the
#'                                   individual-level categorical predictor
#'  }
#' @export
#'
#' @examples
#' mats <- vector(mode = "list", length = 2)
#' m1 <- matrix(rpois(64, 1), ncol = 8)
#' diag(m1) <- 0
#' m2 <- matrix(rpois(64, 5), ncol = 8)
#' diag(m2) <- 0
#' colnames(m1) <- colnames(m2) <- letters[1:8]
#' rownames(m1) <- rownames(m2) <- letters[1:8]
#'
#' matlist <- list(beh1 = m1, beh2 = m2)
#' make_stan_data_from_matrices(mats = matlist)
#'
#' # adding a predictor for individual-level axis
#' make_stan_data_from_matrices(mats = matlist,
#'                              indi_cat_pred = sample(c(0, 1), 8, TRUE))
#'
#' o <- rep(0.5, 8)
#' names(o) <- letters[1:8]
#' o[1:2] <- c(1.5, 10.5)
#' make_stan_data_from_matrices(mats = matlist, obseff = o)$obseff
#'
#' # dealing with dyads that were not coresident (NA values)
#' m1 <- matrix(rpois(64, 1), ncol = 8)
#' diag(m1) <- 0
#' m1[2, 5] <- NA
#' m1[5, 2] <- NA
#' res <- make_stan_data_from_matrices(list(m1))
#' res$n_dyads # should be 28 if all dyads were coresident, but is 27 (removing dyad [2,5])
#'

make_stan_data_from_matrices <- function(mats,
                                         behav_types = NULL,
                                         obseff = NULL,
                                         indi_cat_pred = NULL,
                                         indi_covariate_pred = NULL,
                                         dyad_cat_pred = NULL,
                                         dyad_covariate_pred = NULL,
                                         correlations = FALSE
                                         ) {

  # some easy checks:
  if ("dur_gamma0" %in% behav_types && correlations == TRUE) {
    stop("can't handle correlations with dur_gamma0", call. = FALSE)
  }


  # convert obseff to list of matrices
  # if NULL: convert to matrices filled with 1's in upper triangle
  # if vectors: sum individual values and put them in upper triangle
  obseff <- obseff_to_matrix(obseff = obseff, mats = mats)

  # set diagonals of matrices to NA
  mats <- lapply(mats, function(x) {
    diag(x) <- NA
    x
  })
  obseff <- lapply(obseff, function(x) {
    diag(x) <- NA
    x
  })


  # check whether matrices are identical with respect to NA values
  testobj <- c(mats, obseff)
  test1 <- unique(lapply(testobj, is.na))
  if (length(test1) != 1) {
    if (is.null(obseff)) stop("behavior matrices differ with respect to NA values (or column and row names)", call. = FALSE)
    if (!is.null(obseff)) stop("behavior and observation effort matrices differ with respect to NA values (or column and row names)", call. = FALSE)
  }

  # at this point we have matrices that are identical with respect to row and column names and where there are NA values

  # need to remove any ids for which *all* dyadic values are NA
  # but keep individuals for which only a subset of dyadic values is NA
  aux <- which((rowSums(is.na(mats[[1]])) + colSums(is.na(mats[[1]]))) != ncol(mats[[1]]) * 2)
  for (i in seq_len(length(mats))) {
    mats[[i]] <- mats[[i]][aux, aux]
    obseff[[i]] <- obseff[[i]][aux, aux]
  }


  n_ids <- ncol(mats[[1]])
  n_dyads <- n_ids * (n_ids - 1) * 0.5
  n_beh <- length(mats)

  if (!is.null(indi_cat_pred)) {
    indi_cat_pred <- na.omit(indi_cat_pred)
    if (length(indi_cat_pred) != n_ids) {
      stop("individual-level predictor doesn't have correct length",
           call. = FALSE)
    }
  }

  # deal with behavior types
  if (is.null(behav_types)) {
    behav_types <- rep("count", n_beh)
  }
  if (length(behav_types) != n_beh) {
    stop("require exactly ", n_beh, " 'behave_types' specified")
  }
  if (!all(behav_types %in% c("count", "prop", "dur_gamma", "dur_gamma0", "dur_beta", "binary"))) {
    stop("unknown behavior type supplied")
  }

  # and prep for output (convert to numeric)
  behav_types_num <- rep(1, n_beh) # counts (pois as default)
  behav_types_num[behav_types == "prop"] <- 2
  behav_types_num[behav_types == "dur_gamma"] <- 3
  behav_types_num[behav_types == "dur_beta"] <- 4
  behav_types_num[behav_types == "binary"] <- 6

  # indexing for optional shape/dispersion parameters
  gamma_shape <- logical(n_beh)
  gamma_shape[behav_types == "dur_gamma"] <- TRUE
  gamma_shape[behav_types == "dur_gamma0"] <- TRUE
  gamma_shape_pos <- numeric(n_beh)

  beta_shape <- logical(n_beh)
  beta_shape[behav_types == "dur_beta"] <- TRUE
  beta_shape_pos <- numeric(n_beh)

  for (i in seq_len(n_beh)) {
    if (behav_types[i] %in% c("dur_gamma", "dur_gamma0")) {
      gamma_shape_pos[i] <- i
    }
    if (behav_types[i] == "dur_beta") {
      beta_shape_pos[i] <- i
    }
  }

  gamma_shape_pos_mod <- numeric(n_beh)
  gamma_shape_pos_mod[gamma_shape_pos > 0] <- seq_len(sum(gamma_shape_pos > 0))
  beta_shape_pos_mod <- numeric(n_beh)
  beta_shape_pos_mod[beta_shape_pos > 0] <- seq_len(sum(beta_shape_pos > 0))


  interactions <- matrix(ncol = length(mats), nrow = n_dyads, NA)
  obseff_dat <- matrix(ncol = length(mats), nrow = n_dyads, NA)
  index <- which(upper.tri(mats[[1]]), arr.ind = TRUE)
  na_dyads <- which(apply(index, 1, function(x) is.na(mats[[1]][x[1], x[2]])))
  # if (is.null(obseff)) {
  #   sdata <- rep(0.5, n_ids)
  # } else {
  #   sdata <- obseff
  # }
  sdata <- obseff

  for (i in seq_along(mats)) {
    for (k in seq_len(n_dyads)) {
      val1 <- mats[[i]][index[k, 1], index[k, 2]]
      val2 <- mats[[i]][index[k, 2], index[k, 1]]
      # old way
      # interactions[k, i] <- val1
      # interactions[k, i] <- interactions[k, i] + val2
      # new way
      # if both vals are NA: the dyad is NA
      if (is.na(val1) && is.na(val2)) next
      # val <- sum(val1, val2, na.rm = TRUE) # ignore NA if they only appear in one matrix triangle
      val <- sum(val1, val2) # ignore NA if they only appear in one matrix triangle
      interactions[k, i] <- val
    }
  }

  ## check for pot. problems wrt all(0) or all(!0)
  for (i in seq_len(n_beh)) {
    if (behav_types[i] == "dur_gamma0") {
      if (all(interactions[, i] > 0)) {
        warning("'dur_gamma0' without zero values detected ",
                "(this might not be a problem)")
      }
      if (all(interactions[, i] == 0)) {
        warning("'dur_gamma0' with only zero-values detected ",
                "(this is likely going be a problem for fitting ",
                "the default model)")
      }
    }
    if (behav_types[i] == "binary") {
      if (all(interactions[, i] == 0)) {
        warning("'binary' with only 0 values detected")
      }
      if (all(interactions[, i] == 1)) {
        warning("'binary' with all values being 1 detected")
      }
    }
  }


  # deal with observation effort -------------
  # 1) supplied as single vector or matrix (applies to all behaviors equally)
  if (!is.list(sdata)) {
    if (is.vector(sdata)) {
      # make sure order is the same in vector and interaction matrix columns
      if (!is.null(colnames(mats[[1]])) && !is.null(names(sdata))) {
        sdata <- sdata[colnames(mats[[1]])]
      }
      for (k in seq_len(n_dyads)) {
        obseff_dat[k, ] <- sdata[index[k, 1]] + sdata[index[k, 2]]
      }
    }
    if (is.matrix(sdata)) {
      obseff_dat[, ] <- sdata[index]
    }
  }

  # 2) supplied as list of vectors or matrices (mapped onto each behavior)
  if (is.list(sdata)) {
    if (length(sdata) != length(mats)) {
      stop("behaviour matrices and observation effort can't be matched")
    }
    for (i in seq_along(mats)) {
      if (is.vector(sdata[[i]])) {
        for (k in seq_len(n_dyads)) {
          # old way
          obseff_dat[k, i] <- sdata[[i]][index[k, 1]] + sdata[[i]][index[k, 2]]
        }
      }
      if (is.matrix(sdata[[i]])) {
        obseff_dat[, i] <- sdata[[i]][index]
      }
    }
  }

  # set obseff for props to 100 if not supplied...
  # if (is.null(obseff)) {
    for (i in seq_along(mats)) {
      if (behav_types[i] == "prop" & all(obseff_dat[, i] == 1 ,na.rm = TRUE)) {
        obseff_dat[, i] <- 100
      }
    }
  # }

  # filter for non-NA dyads
  # do it at the beginning of the function
  # this an index for those *dyads* that have non-NA values
  sel <- which(!is.na(interactions[, 1]))
  # and dyads which were removed
  removed_dyads <- index[is.na(interactions[, 1]), , drop = FALSE]


  # create default priors for behaviors
  # at this point 'interactions' is not yet split into discrete vs continuous
  prior_matrix <- matrix(ncol = 2, nrow = length(mats))
  for (i in seq_along(mats)) {
    response <- interactions[sel, i]
    if (behav_types[i] == "dur_gamma0") {
      # for bernoulli part of mixture...
      prior_matrix[i, ] <- c(0, 2.5)
      next
    }
    prior_matrix[i, ] <- make_prior(response = response,
                                    type = behav_types[i],
                                    obseff = obseff_dat[sel, i])
  }
  # need a second set in case we have behaviors with more than one mean parameter
  prior_matrix2 <- matrix(ncol = 2, nrow = n_beh, 0)
  for (i in 1:n_beh) {
    # for gamma part in the mixture
    if (behav_types[i] == "dur_gamma0") {
      response <- interactions[, i]
      prior_matrix2[i, ] <- make_prior(response = response,
                                       type = "dur_gamma0",
                                       obseff = obseff_dat[, i]
                                       # obseff = obseff[, i]
                                       )
    }
  }

  if (!is.null(colnames(mats[[1]]))) {
    id_codes <- as.character(colnames(mats[[1]]))
  } else {
    id_codes <- as.character(seq_len(n_ids))
  }
  vec <- rep(0, n_ids)
  names(vec) <- id_codes

  if (!is.null(names(mats))) {
    beh_codes <- as.character(names(mats))
  } else {
    beh_codes <- paste0("behav_", LETTERS[seq_len(length(mats))])
  }
  vec_behaviors <- rep(0, length(beh_codes))
  names(vec_behaviors) <- beh_codes

  # code behave_types as named vector
  bdata <- c(behav_types_num, 0)
  names(bdata) <- c(behav_types, "0")

  out <- list(id1 = index[sel, 1],
              id2 = index[sel, 2],
              behav_types = bdata,
              interactions = apply(interactions[sel, , drop = FALSE], 2, as.integer),
              interactions_cont = apply(interactions[sel, , drop = FALSE], 2, as.numeric),
              n_beh = length(mats),
              n_ids = n_ids,
              # n_dyads = as.integer(n_dyads),
              n_dyads = length(sel),
              dyads_navi = as.matrix(index[sel, 1:2]),
              obseff = obseff_dat[sel, , drop = FALSE],
              obseff_int = apply(obseff_dat[sel, , drop = FALSE], 2, as.integer),
              gamma_shape_pos = gamma_shape_pos_mod,
              gamma_shape_n = sum(gamma_shape_pos > 0),
              beta_shape_pos = beta_shape_pos_mod,
              beta_shape_n = sum(beta_shape_pos > 0),
              # individual/dyad level predictors:
              # indi_cat_pred = 0, # binary
              # indi_covariate_pred = 0, # continuous
              # dyad_cat_pred = 0,
              # dyad_covariate_pred = 0,
              prior_matrix = prior_matrix,
              prior_matrix2 = prior_matrix2,
              id_codes = vec,
              beh_names = vec_behaviors,
              removed_dyads = removed_dyads,
              n_cors = 0,
              prior_only = 0)

  if (!is.null(indi_cat_pred)) {
    out$indi_cat_pred <- indi_cat_pred
    standardization_check_predictors(indi_cat_pred, "indi_cat_pred")
  }
  if (!is.null(indi_covariate_pred)) {
    out$indi_covariate_pred <- indi_covariate_pred
    standardization_check_predictors(indi_covariate_pred, "indi_covariate_pred")
  }
  if (!is.null(dyad_cat_pred)) {
    out$dyad_cat_pred <- dyad_cat_pred
    standardization_check_predictors(dyad_cat_pred, "dyad_cat_pred")
  }
  if (!is.null(dyad_covariate_pred)) {
    out$dyad_covariate_pred <- dyad_covariate_pred
    standardization_check_predictors(dyad_covariate_pred, "dyad_covariate_pred")
  }

  if (correlations) {
    out$n_cors <- out$n_beh * (out$n_beh - 1) * 0.5
  }


  out
}


standardization_check_predictors <- function(x, kind) {
  x_mean <- abs(mean(x)) > 0.0001
  x_sd <- abs(1 - sd(x)) > 0.0001
  if (x_mean || x_sd) {
    warning("the variable ", shQuote(kind), " is not standardized (mean != 0 and/or sd != 1)", call. = FALSE)
  }
  NULL
}
```
