## Supplementary material for "(De)composing sociality: disentangling individual-specific from dyad-specific propensities to interact": bamoso R package: model_comparison.R

```
#' model cross validation / comparison
#'
#' @param x a dyadic model or a (named) list of such models
#'
#' @return output of \code{\link[loo]{loo}} or \code{\link[loo]{loo_compare}}
#' @export
#' @examples
#' \dontrun{
#' library(loo)
#' model1 <- fit_example_model(num = 1)
#' model_comparison(model1)
#'
#' model2 <- fit_example_model(num = 2)
#' model_comparison(list(mod1 = model1, mod2 = model2))
#' }
#'

# x=list(r_correct = r_correct, r_incorrect = r_incorrect)
# x=r_correct
model_comparison <- function(x) {
  if (!requireNamespace("loo", quietly = TRUE)) {
    stop("Package 'loo' is required for this function but is not installed. ",
         "Please install it using install.packages('loo').")
  }

  if (inherits(x, "dyadicmodel")) {
    res <- loo::loo(x$mod_res$draws("log_lik", format = "draws_matrix"))
    return(res)
  }

  if (inherits(x, "list")) {
    if (!all(unlist(lapply(x, inherits, "dyadicmodel")))) {
      stop("not a list of dyadic models...")
    }
    zz <- lapply(x, function(y) loo::loo(y$mod_res$draws("log_lik", format = "draws_matrix")))
    return(loo::loo_compare(zz))
  }
}
```
