## Supplementary material for "(De)composing sociality: disentangling individual-specific from dyad-specific propensities to interact": bamoso R package: model_summary.R

```
#' summary of model
#'
#' @param mod_res model results from
#'                \code{\link{sociality_model}})
#' @param probs numeric, the desired quantiles (default is
#'              \code{c(0.055, 0.945)}, i.e. a 89% interval)
#' @param raw logical, default is \code{FALSE}. If \code{TRUE}, the original
#'            summary is returned (which can also be obtained via
#'            \code{mod_res$summary()})
#'
#' @return a data frame
#' @export
#' @importFrom posterior rhat ess_bulk ess_tail summarise_draws
#' @importFrom stats sd
#' @examples
#' \dontrun{
#' data("grooming")
#' mat <- grooming$ass$groom[1:9, 1:9]
#' obseff <- grooming$ass$obseff[1:9]
#' standat <- make_stan_data_from_matrices(mats = list(groom = mat),
#'                                         behav_types = "count",
#'                                         obseff = list(obseff))
#' res <- sociality_model(standat = standat, parallel_chains = 4,
#'                        adapt_delta = 0.9, seed = 17)
#' model_summary(res)
#' }
#'
#'
#'
#'


model_summary <- function(mod_res,
                          probs = c(0.055, 0.945),
                          raw = FALSE
) {

  standat <- mod_res$standat
  mod_res <- mod_res$mod_res

  vars <- c("indi_soc_sd",
            "dyad_soc_sd",
            "beh_intercepts")
  if (standat$n_cors > 0) {
    vars <- c(vars, "cors_indi", "cors_dyad")
  }
  vars <- c(vars, "indi_soc_vals", "dyad_soc_vals")

  if (raw) {
    res <- as.data.frame(mod_res$summary(variables = vars))

    return(res)
  }

  xdraws <- mod_res$draws(variables = vars,
                          format = "draws_matrix")

  out <- as.data.frame(summarise_draws(xdraws,
                                       mean,
                                       median,
                                       sd,
                                       ~quantile(.x, probs = probs),
                                       rhat,
                                       ess_bulk = ~round(ess_bulk(.x)),
                                       ess_tail = ~round(ess_tail(.x))))

  out <- data.frame(label = out$variable,
                    categ = NA,
                    beh = NA,
                    stanvarname = out$variable,
                    out[, -1], check.names = FALSE)

  out$categ[grepl("dyad_soc_vals", out$label)] <- "dyad_vals"
  out$categ[grepl("indi_soc_vals", out$label)] <- "indi_vals"
  out$categ[grepl("beh_intercepts", out$label)] <- "intercept"
  out$categ[grepl("indi_soc_sd", out$label)] <- "sd"
  out$categ[grepl("dyad_soc_sd", out$label)] <- "sd"

  out$label[grepl("indi_soc_sd", out$label)] <- "greg_sd"
  out$label[grepl("dyad_soc_sd", out$label)] <- "affi_sd"


  if (standat$n_cors > 0) {
    bnames <- names(standat$beh_names)
    xindex <- which(upper.tri(matrix(ncol = standat$n_beh, nrow = standat$n_beh)), arr.ind = TRUE)
    xnames <- apply(xindex, 1, function(x) paste(bnames[x[1]], bnames[x[2]], sep = " * "))

    out$label[grepl("cors_indi", out$label)] <- "greg_cor"
    out$label[grepl("cors_dyad", out$label)] <- "affi_cor"
    out$categ[grepl("greg_cor", out$label)] <- "cor"
    out$categ[grepl("affi_cor", out$label)] <- "cor"
    out$beh[grepl("greg_cor", out$label)] <- xnames
    out$beh[grepl("affi_cor", out$label)] <- xnames

    out$beh[grepl("intercept", out$categ)] <- bnames

    out$beh[out$label == "greg_sd"] <- bnames
    out$beh[out$label == "affi_sd"] <- bnames
    out$beh[which(out$categ == "indi_vals")] <- rep(bnames, each = standat$n_ids)
    out$beh[which(out$categ == "dyad_vals")] <- rep(bnames, each = standat$n_dyads)
  }


  if (!is.null(standat)) {
    out$label[grepl("indi_soc_vals", out$label)] <- names(standat$id_codes)

    dyads <- paste(names(standat$id_codes)[standat$dyads_navi[, 1]],
                   names(standat$id_codes)[standat$dyads_navi[, 2]],
                   sep = "_@_")
    out$label[grepl("dyad_soc_vals", out$label)] <- dyads
    behs <- names(standat$beh_names)
    out$label[grepl("beh_intercepts", out$label)] <- behs
  }
  out
}
```
