## Supplementary material for "(De)composing sociality: disentangling individual-specific from dyad-specific propensities to interact": bamoso R package: pp_binary.R

```
# xx <- generate_data(n_ids = 18,  behav_types = "binary",
#                     indi_sd = 1, dyad_sd = 1.2,
#                     beh_intercepts = c(-1.4),
#                     exact = TRUE)
#
# xx <- generate_data(n_ids = 18, n_beh = 2, behav_types = c("binary", "binary"),
#                     indi_sd = 1, dyad_sd = 1.2,
#                     beh_intercepts = c(-1.4, 1),
#                     exact = TRUE)
#
# xx <- generate_data(n_ids = 15, n_beh = 2, behav_types = c("dur_gamma0", "binary"),
#                     indi_sd = 1, dyad_sd = 1.5,
#                     beh_intercepts = c(-1.4, -0.5),
#                     beh_intercepts_add = c(2, 0), count_obseff = c(0.1, 11),
#                     exact = TRUE, disp_pars_gamma = c(11))
# xres <- sociality_model(standat = xx$standat, parallel_chains = 4, chains = 4, iter_sampling = 350, show_exceptions = F, refresh = 0, adapt_delta = 0.95, silent = TRUE)


#' Posterior predictive checks for binary data
#'
#' @param mod_res model result from \code{\link{sociality_model}}
#' @param xvar numeric or character, the index or name of the behavior to
#'             be plotted. Default is \code{1}, i.e. the first behavior.
#' @param n_draws numeric of length 1. Number of posterior draws.
#' @param selected_id character of length one. Choose a single id.
#' @param all_ids logical, plot all individuals next to each other (default
#'          is \code{FALSE}). This overrides \code{selected_id}.
#' @param xlab,ylab character, axis labels
#'
#' @importFrom graphics title
#' @returns a plot
#' @export
#'
#' @examples
#' \dontrun{
#' # two behaviours with binary component (dur_gamma0, and binary)
#' set.seed(27)
#' xx <- generate_data(n_ids = 15, n_beh = 2,
#'                     behav_types = c("dur_gamma0", "binary"),
#'                     indi_sd = 1, dyad_sd = 2,
#'                     beh_intercepts = c(-1.4, -0.5),
#'                     beh_intercepts_add = c(2, 0))
#' xres <- sociality_model(standat = xx$standat, parallel_chains = 4,
#'                         chains = 4, iter_sampling = 350, refresh = 0,
#'                         show_exceptions = F, adapt_delta = 0.95,
#'                         silent = TRUE, seed = 47)
#'
#' pp_binary(mod_res = xres, xvar = 1, all_ids = TRUE)
#' pp_binary(mod_res = xres, xvar = 2, all_ids = TRUE)
#' # and the gamma component for the mixture behavior
#' pp_model_dens(mod_res = xres, xvar = 1, xlim = c(0, 1000))
#'
#' # 'global' summary (pooled across individuals)
#' pp_binary(mod_res = xres, xvar = 1, all_ids = FALSE)
#' pp_binary(mod_res = xres, xvar = 2, all_ids = FALSE)
#' }
pp_binary <- function(mod_res,
                      xvar = 1,
                      n_draws = 20,
                      selected_id = NULL,
                      all_ids = FALSE,
                      xlab = "sucess/ocurred/non-zero values(?)",
                      ylab = ""
                      ) {
  standat <- mod_res$standat
  mod_res <- mod_res$mod_res
  btype <- names(standat$behav_types)[xvar]

  if (!btype %in% c("dur_gamma0", "binary")) {
    stop("binary checks require behavior type to be dur_gamma0 or binary")
  }
  if (length(selected_id) > 1) stop("please select only one id")

  if (btype == "dur_gamma0") {
    var_pattern <- "interactions_pred_cont"
  }
  if (btype == "binary") {
    var_pattern <- "interactions_pred"
  }
  p <- as.matrix(mod_res$draws(variables = var_pattern, format = "matrix"))
  p <- p[sample(nrow(p), n_draws), ]
  p <- p[, grepl(paste0(",", xvar, "\\]", collapse = ""), colnames(p))]
  p <- t(apply(p > 0, 1, as.integer))

  if (all_ids) {
    navi <- standat$dyads_navi
    id_codes <- names(standat$id_codes)
    obsval <- numeric(length(id_codes))
    pp_vals <- matrix(ncol = length(id_codes), nrow = n_draws)

    for (i in seq_along(id_codes)) {
      selected_id <- id_codes[i]
      id_index <- which(names(standat$id_codes) == selected_id)
      dyad_index <- which(apply(navi, 1, function(xx) id_index %in% xx))
      # p <- p[, dyad_index, drop = FALSE]
      pp_vals[, i] <- rowSums(p[, dyad_index, drop = FALSE])
      obsval[i] <- sum(standat$interactions_cont[dyad_index, xvar] > 0)
    }
    colnames(pp_vals) <- id_codes
    names(obsval) <- id_codes
    id_labs <- id_codes
    xlocs <- seq_along(obsval)
  } else {
    # selected_id <- "2"
    if (!is.null(selected_id)) {
      if (!selected_id %in% names(standat$id_codes)) {
        stop("didn't find", shQuote(selected_id), "in stan data")
      }
      id_index <- which(names(standat$id_codes) == selected_id)
      navi <- standat$dyads_navi
      dyad_index <- which(apply(navi, 1, function(xx) id_index %in% xx))
      p <- p[, dyad_index, drop = FALSE]
      pp_vals <- rowSums(p)
      obsval <- sum(standat$interactions_cont[dyad_index, xvar] > 0)
    } else {
      obsval <- sum(standat$interactions_cont[, xvar] > 0)
      pp_vals <- rowSums(p)
    }
    pp_vals <- t(t(pp_vals))
    id_labs <- NA
    xlocs <- NULL
  }

  plot(0, 0, xlim = c(0.5, length(obsval) + 0.5),
       ylim = c(0, max(pp_vals, obsval) * 1.05),
       type = "n", yaxs = "i", axes = FALSE, ann = FALSE)
  title(xlab = xlab, ylab = ylab)
  rect(seq_along(obsval) - 0.3, 0, seq_along(obsval) + 0.3, obsval)
  axis(1, tcl = 0, at = xlocs, labels = id_labs)
  axis(2)

  sapply(seq_along(obsval), function(x) {
    points(runif(n_draws, x - 0.1, x + 0.1), pp_vals[, x], pch = 16,
           col = grey(0.3, alpha = 0.6), xpd = TRUE)
  })

  points(seq_along(obsval), colMeans(pp_vals),
         col = 'red', pch = 4, lwd = 2, cex = 1.5)

  box(bty = "l")

  invisible(list(observed = obsval, fromdraws = pp_vals))
}
```
