## Supplementary material for "(De)composing sociality: disentangling individual-specific from dyad-specific propensities to interact": bamoso R package: pp_model.R

```
#' Posterior predictive checks for discrete data (via histogram)
#'
#' @param mod_res model result from \code{\link{sociality_model}}
#' @param xvar numeric or character, the index or name of the behavior to
#'             be plotted. Default is \code{1}, i.e. the first behavior.
#' @param xlim numeric
#' @param xbreaks numeric
#' @param selected_id character of length 1, if provided: plot only data for
#'                    that individual. If left at its default (\code{NULL}),
#'                    plot aggregate over all individuals.
#' @param print_info logical, default is \code{FALSE}. If \code{TRUE}, some
#'                   summaries are printed to screen for the selected
#'                   individual. (This argument is ignored if
#'                    \code{selected_id = NULL}).
#' @param ... further arguments for \code{\link{hist}}
#'
#' @importFrom graphics box hist points rect segments axis
#' @importFrom stats quantile
#' @return a plot
#' @export
#'

pp_model <- function(mod_res,
                     xvar = 1,
                     xlim = NULL,
                     xbreaks = NULL,
                     selected_id = NULL,
                     print_info = FALSE,
                     ...) {

  standat <- mod_res$standat
  mod_res <- mod_res$mod_res

  if (is.character(xvar)) {
    xvar <- which(names(standat$beh_names) == xvar)
  }

  if (is.null(xbreaks)) xbreaks <- "Sturges" # default of 'hist()'
  if (length(selected_id) > 1) stop("please select only one id", call. = FALSE)

  # suppress warnings due to handing over graphical parameters via ...
  #   that are not needed in this first call to hist
  x <- suppressWarnings(hist(standat$interactions[, xvar],
                             plot = FALSE,
                             breaks = xbreaks, ...))
  p <- as.matrix(mod_res$draws(variables = "interactions_pred",
                               format = "matrix"))
  p <- p[, grepl(paste0(",", xvar, "\\]", collapse = ""), colnames(p))]
  # remove overflow cases if any
  xtest <- is.na(rowSums(p))
  if (any(xtest)) {
    if (interactive()) message("removed ", sum(!xtest), " draws because of overflow")
    p <- p[!xtest, , drop = FALSE]
  }

  # deal with individual selection
  if (!is.null(selected_id)) {
    if (!selected_id %in% names(standat$id_codes)) {
      stop("didn't find ", shQuote(selected_id), " in stan data")
    }
    id_index <- which(names(standat$id_codes) == selected_id)
    navi <- standat$dyads_navi
    dyad_index <- which(apply(navi, 1, function(xx) id_index %in% xx))
    p <- p[, dyad_index, drop = FALSE]
    x <- suppressWarnings(hist(standat$interactions[dyad_index, xvar],
                               plot = FALSE,
                               breaks = xbreaks,
                               ...))

    if (print_info) {
      print_data <- standat$interactions[dyad_index, xvar]
      cat("observed total:", sum(print_data), "\n")
      cat("---------------\n")
      cat("lower10% predicted total:", quantile(rowSums(p), 0.1), "\n")
      cat("median predicted total:", median(rowSums(p)), "\n")
      cat("upper10% predicted total:", quantile(rowSums(p), 0.9), "\n")
    }

  }


  # transform everything into bins...
  pd <- as.matrix(t(apply(p, 1, function(z) {
    table(cut(z, breaks = x$breaks, include.lowest = TRUE))
  }
  )))
  dimnames(pd) <- NULL

  pd <- apply(pd, 2, quantile, probs = c(0.055, 0.5, 0.945))
  # remove 'empty' cells
  sel <- which(colSums(rbind(x$counts, pd)) > 0)

  maxval <- max(x$counts, pd)
  if (is.null(xlim)) xlim <- range(x$breaks)

  suppressWarnings(plot(0, 0, type = "n", xlim = xlim,
                        ylim = c(0, maxval * 1.05),
                        yaxs = "i", axes = FALSE, las = 1, bty = "n", ...))
  axis(1)

  # for (i in seq_along(x$counts)) {
  for (i in sel) {
    rect(xleft = x$breaks[i], ybottom = 0, xright = x$breaks[i + 1],
         ytop = x$counts[i], col = "grey", border = "white")
    points(x$mids[i], pd[2, i], pch = 21, cex = 1.5, xpd = TRUE,
           col = "white", bg = "black")
    segments(x0 = x$mids[i], y0 = pd[1, i], x1 = x$mids[i], y1 = pd[3, i])
  }
  box(bty = "l")
  invisible(rbind(observed = x$counts, pd, bin_mids = x$mids))
}
```
