## Supplementary material for "(De)composing sociality: disentangling individual-specific from dyad-specific propensities to interact": bamoso R package: pp_model_dens.R

```
#' Posterior predictive checks for continuous or discrete data (via density)
#'
#' @param mod_res model result from \code{\link{sociality_model}}
#' @param xvar numeric or character, the index or name of the behavior to
#'             be plotted. Default is \code{1}, i.e. the first behavior.
#' @param n_draws numeric
#' @param xlim,ylim numeric, each of length two. Overrides axis limits gleaned
#'                  from the data. Default is \code{NULL} (take values from the
#'                  model and data)
#' @param xlab,ylab character, axis labels
#' @param xadjust numeric, \code{adjust} parameter for \code{\link{density}}
#' @param plot_draws logical, should the random draws actually be plotted
#'                   (default is \code{TRUE})
#' @param selected_id character of length 1, if provided: plot only data for
#'                    that individual. If left at its default (\code{NULL}),
#'                    plot aggregate over all individuals.
#' @param print_info logical, default is \code{FALSE}. If \code{TRUE}, some
#'                   summaries are printed to screen for the selected
#'                   individual. (This argument is ignored if
#'                    \code{selected_id = NULL}).
#' @param ... further arguments for \code{\link{density}}
#'
#' @importFrom graphics box hist points rect segments
#' @importFrom stats quantile
#' @return a plot
#' @export
#'

pp_model_dens <- function(mod_res,
                          xvar = 1,
                          n_draws = 20,
                          plot_draws = TRUE,
                          xlim = NULL,
                          ylim = NULL,
                          xlab = "behaviour prop/duration/freq(?)",
                          ylab = "",
                          xadjust = NULL,
                          selected_id = NULL,
                          print_info = FALSE,
                          ...) {

  standat <- mod_res$standat
  mod_res <- mod_res$mod_res

  if (is.character(xvar)) {
    xvar <- which(names(standat$beh_names) == xvar)
  }

  if (is.null(xadjust)) xadjust <- 1 # density()'s default
  if (length(selected_id) > 1) stop("please select only one id")

  if (standat$behav_types[xvar] %in% c(3, 4, 5)) {
    sel <- seq_len(nrow(standat$interactions_cont))
    if (standat$behav_types[xvar] %in% c(5)) {
      # only use non-zero data for gamma mixture
      sel <- which(standat$interactions_cont[, xvar] > 0)
    }

    x <- suppressWarnings(density(standat$interactions_cont[sel, xvar],
                                  adjust = xadjust, ...))
    p <- as.matrix(mod_res$draws(variables = "interactions_pred_cont",
                                 format = "matrix"))
  } else {
    x <- suppressWarnings(density(standat$interactions[, xvar],
                                  adjust = xadjust, ...))
    p <- as.matrix(mod_res$draws(variables = "interactions_pred",
                                 format = "matrix"))
  }

  p <- p[, grepl(paste0(",", xvar, "\\]", collapse = ""), colnames(p))]
  if (standat$behav_types[xvar] %in% c(5)) {
    # only use non-zero data for gamma mixture
    p <- p[, sel]
  }


  # remove overflow cases if any
  xtest <- is.na(rowSums(p))
  if (any(xtest)) {
    if (interactive()) message("removed ", sum(!xtest), " draws because of overflow")
    p <- p[!xtest, , drop = FALSE]
  }

  p <- p[sample(nrow(p), n_draws), ]

  # deal with individual selection
  if (!is.null(selected_id)) {
    if (!selected_id %in% names(standat$id_codes)) {
      stop("didn't find", shQuote(selected_id), "in stan data")
    }
    id_index <- which(names(standat$id_codes) == selected_id)
    navi <- standat$dyads_navi
    dyad_index <- which(apply(navi, 1, function(xx) id_index %in% xx))
    p <- p[, dyad_index, drop = FALSE]

    if (standat$behav_types[xvar] %in% c(3, 4)) {
      x <- suppressWarnings(density(standat$interactions_cont[dyad_index, xvar],
                                    adjust = xadjust, ...))
    } else {
      x <- suppressWarnings(density(standat$interactions[dyad_index, xvar],
                                    adjust = xadjust, ...))
    }

    if (print_info) {
      if (standat$behav_types[xvar] %in% c(3, 4)) {
        print_data <- standat$interactions_cont[dyad_index, xvar]
      } else {
        print_data <- standat$interactions[dyad_index, xvar]
      }

      cat("observed total:", sum(print_data), "\n")
      cat("---------------\n")
      cat("lower10% predicted total:", quantile(rowSums(p), 0.1), "\n")
      cat("median predicted total:", median(rowSums(p)), "\n")
      cat("upper10% predicted total:", quantile(rowSums(p), 0.9), "\n")
    }

  }


  suppressWarnings(pd <- apply(p, 1, density, adjust = xadjust, ...))
  maxx <- max(max(x$x, unlist(lapply(pd, function(y) max(y$x)))))
  maxy <- max(max(x$y, unlist(lapply(pd, function(y) max(y$y)))))
  miny <- 0
  minx <- 0

  if (is.null(xlim)) xlim <- c(minx, maxx)
  if (is.null(ylim)) ylim <- c(miny, maxy * 1.05)

  suppressWarnings(plot(0, 0, type = "n", axes = FALSE,
                        xlim = xlim, ylim = ylim,
                        xlab = xlab, ylab = ylab,
                        yaxs = "i", las = 1, bty = "n", xaxs = "i", ...))
  axis(1)
  box(bty = "l")

  if (plot_draws) {
    for (i in seq_len(n_draws)) {
      points(pd[[i]]$x, pd[[i]]$y, type = "l", lwd = 0.5)
    }
  }

  points(x$x, x$y, type = "l", lwd = 2, col = "red")

  invisible(list(observed = x, fromdraws = pd))

}
```
