## Supplementary material for "(De)composing sociality: disentangling individual-specific from dyad-specific propensities to interact": bamoso R package: pp_model_stat.R

```
#' Posterior predictive checks with summary statistics
#'
#' @param mod_res model result from \code{\link{sociality_model}}
#' @param xvar numeric or character, the index or name of the behavior to
#'             be plotted. Default is \code{1}, i.e. the first behavior.
#' @param stat statistic
#' @param xlim numeric
#' @param ... further arguments to hist
#'
#' @importFrom graphics abline
#'
#' @return a plot
#' @export
#'

pp_model_stat <- function(mod_res,
                          xvar = 1,
                          stat = c("mean", "median", "min", "max", "iqr", "range_width"),
                          xlim = NULL,
                          ...) {

  standat <- mod_res$standat
  mod_res <- mod_res$mod_res

  if (is.character(xvar)) {
    xvar <- which(names(standat$beh_names) == xvar)
  }

  if (standat$behav_types[xvar] %in% c(3, 4)) {
    p <- as.matrix(mod_res$draws(variables = "interactions_pred_cont",
                                 format = "matrix"))
    x <- standat$interactions_cont[, xvar]
  } else {
    p <- as.matrix(mod_res$draws(variables = "interactions_pred",
                                 format = "matrix"))
    x <- standat$interactions[, xvar]
  }


  p <- p[, grepl(paste0(",", xvar, "\\]", collapse = ""), colnames(p))]

  # remove overflow cases if any
  xtest <- !is.na(rowSums(p))
  if (any(xtest)) {
    if (interactive()) message("removed ", sum(!xtest), " draws because of overflow")
    p <- p[xtest, , drop = FALSE]
  }

  if (stat == "mean") {
    p <- rowMeans(p)
    x <- mean(x)
  }
  if (stat == "min") {
    p <- apply(p, 1, min)
    x <- min(x)
  }
  if (stat == "max") {
    p <- apply(p, 1, max)
    x <- max(x)
  }
  if (stat == "median") {
    p <- apply(p, 1, median)
    x <- median(x)
  }
  if (stat == "range_width") {
    p <- apply(p, 1, max) - apply(p, 1, min)
    x <- max(x) - min(x)
  }
  if (stat == "iqr") {
    p <- apply(p, 1, function(yy) diff(quantile(yy, probs = c(0.25, 0.75))))
    x <- diff(quantile(x, probs = c(0.25, 0.75)))
  }

  h <- hist(p, plot = FALSE)
  if (is.null(xlim)) xlim <- range(h$breaks)
  ylim <- c(0, max(h$counts) * 1.05)

  suppressWarnings(hist(p, xlim = xlim, ylim = ylim, yaxs = "i",
                        axes = FALSE, las = 1, bty = "n", ...))
  axis(1)

  abline(v = x, lwd = 3, col = "red")

  box(bty = "l")
}
```
