## Supplementary material for "(De)composing sociality: disentangling individual-specific from dyad-specific propensities to interact": bamoso R package: read_write_model.R

```
#' write/read model to file
#'
#' @param model_object output of \code{\link{sociality_model}}
#' @param file_name the path and basename for the files (no file extension! this will be added automatically...)
#'
#' @details
#' \code{write_model_to_file} writes two files (one for the data list, one for the model environment).
#' \code{read_model_from_file} reads them both and combines them into a single object again.
#'
#' This is somewhat convoluted and if you know your way around cmdstanr environments you might as well interact with the model environment directly (e.g. \code{res$mod_res})
#'
#' @return writes to or reads from file
#' @export
#'
#' @examples
#' \dontrun{
#' set.seed(8)
#' d <- generate_data(n_ids = 7, n_beh = 1, behav_types = "count",
#'                    beh_intercepts = 0.2, indi_sd = 0.8, dyad_sd = 0.9)
#' res <- sociality_model(standat = d$standat, seed = 1, parallel_chains = 4,
#'                        refresh = 0, iter_sampling = 300, iter_warmup = 300)
#'
#' temploc <- file.path(tempdir(), "res")
#' write_model_to_file(model_object = res, file_name = temploc)
#' res2 <- read_model_from_file(temploc)
#'
#' # compare output
#' identical(res$mod_res$draws(), res2$mod_res$draws())
#'
#' set.seed(1)
#' pp_model(res)
#' set.seed(1)
#' pp_model(res2)
#' }
#'

write_model_to_file <- function(model_object, file_name) {

  nm1 <- normalizePath(paste0(file_name, "_model.rds"), mustWork = FALSE)
  nm2 <- normalizePath(paste0(file_name, "_data.rds"), mustWork = FALSE)
  model_object$mod_res$save_object(file = nm1)
  saveRDS(model_object$standat, file = nm2)

  if (interactive()) {
    cat("wrote two files:\n")
    cat(paste0("  - ", nm1, "\n"))
    cat(paste0("  - ", nm2, "\n"))
  }

}


#' @inherit write_model_to_file
#' @export
read_model_from_file <- function(file_name) {
  nm1 <- normalizePath(paste0(file_name, "_model.rds"))
  nm2 <- normalizePath(paste0(file_name, "_data.rds"))

  if (!file.exists(nm1)) stop("didn't find ", shQuote(nm1), call. = FALSE)
  if (!file.exists(nm2)) stop("didn't find ", shQuote(nm2), call. = FALSE)

  standat <- readRDS(nm2)
  mod_res <- readRDS(nm1)

  out <- list(standat = standat, mod_res = mod_res)
  class(out) <- "dyadicmodel"
  out
}
```
