## Supplementary material for "(De)composing sociality: disentangling individual-specific from dyad-specific propensities to interact": bamoso R package: ridge_plot.R

```
#' posteriors of sociality values
#' gregariousness or pairwise affinity
#'
#' @param mod_res model result from \code{\link{sociality_model}}
#' @param beh integer, indicates which behavior is to be returned. Only relevant
#'            if the model was fitted with correlations (and at least 2
#'            behaviors).
#' @param xlim numeric, x-axis limit (default is \code{c(-4, 4)})
#' @param greg logical, with default \code{TRUE}: plot gregariousness. If
#'             \code{FALSE}: plot pairwise affinity
#' @param sel_subset integer or character, select subset of individuals or
#'                   dyads to plot. If plotting dyads (\code{greg=FALSE}),
#'                   dyads are addressed by 'id1_@_id2'. If you supply single
#'                   individuals here, then all dyads for said individual(s)
#'                   are included as well.
#' @param dens_adjust numeric, adjust argument for \code{\link[stats]{density}}
#' @param vert_exp numeric, expansion factor for vertical stretching
#' @param add_median logical, add vertical lines at posterior medians
#' @param labcex numeric, cex factor for vertical axis labels (ids or dyads)
#' @param yaxs_exp numeric, expansion factor for upper limit of y-axis.
#'
#' @importFrom stats median mad median qlogis rt
#' @importFrom graphics polygon legend abline
#' @importFrom grDevices adjustcolor grey hcl.colors
#' @return a plot
#' @export
#'
#' @examples
#' \dontrun{
#' mat <- generate_data(n_ids = 6, n_beh = 1, behav_types = "count",
#'                      indi_sd = 2, beh_intercepts = 2, dyad_sd = 0.5)
#' mat <- mat$processed$interaction_matrices[[1]]
#' colnames(mat) <- letters[1:ncol(mat)]
#' sdat <- make_stan_data_from_matrices(mats = list(groom = mat),
#'                                      behav_types = "count",
#'                                      obseff = NULL)
#' res <- sociality_model(standat = sdat, seed = 123, parallel_chains = 4,
#'                        iter_sampling = 1000, iter_warmup = 1000,
#'                        refresh = 0, adapt_delta = 0.9)
#' ridge_plot(mod_res = res, greg = TRUE)
#' ridge_plot(mod_res = res, greg = FALSE, sel_subset = c("a", "b_@_d"))
#' }


ridge_plot <- function(mod_res,
                       beh = 1,
                       xlim = c(-4, 4),
                       sel_subset = NULL,
                       greg = TRUE,
                       dens_adjust = 1,
                       vert_exp = 2,
                       add_median = TRUE,
                       labcex = 0.5,
                       yaxs_exp = 1.2) {


  standat <- mod_res$standat
  mod_res <- mod_res$mod_res

  # extract draws for subsequent use
  if (greg) {
    alldraws <- mod_res$draws(variables = "indi_soc_vals", format = "draws_matrix")
  } else {
    alldraws <- mod_res$draws(variables = "dyad_soc_vals", format = "draws_matrix")
  }

  # select axis (if correlations are present)
  has_cors <- standat$n_cors > 0
  if (has_cors) {
    alldraws <- alldraws[, grepl(paste0(",", beh, "]"), colnames(alldraws), fixed = TRUE)]
  }

  # convert character ids back to numeric (indexes)
  if (greg) {
    if (!is.null(sel_subset)) {
      if (is.character(sel_subset) || is.factor(sel_subset)) {
        sel_subset <- sapply(as.character(sel_subset), function(x) {
          which(names(standat$id_codes) == x)
        })
      }
    }
  }
  if (!greg) {
    # sel_subset <- c("B", "C_@_D")
    if (!is.null(sel_subset)) {
      if (is.character(sel_subset) || is.factor(sel_subset)) {
        nm_mat <- apply(standat$dyads_navi, 1, function(x) names(standat$id_codes)[x])
        nm <- paste0(nm_mat[1, ], "_@_", nm_mat[2, ])

        sel1 <- which(apply(t(nm_mat), 1, function(x)any(sel_subset %in% x)))

        sel2 <- sapply(as.character(sel_subset), function(x) {
          which(nm == x)
        })

        sel_subset <- unique(c(sel1, unlist(sel2)))
      }
    }
  }


  if (greg) {
    n <- standat$n_ids
    if (is.null(sel_subset)) sel_subset <- seq_len(n)
    # xorder <- rep(0, n)

    # order according to posterior median
    xorder <- as.numeric(apply(alldraws, 2, median))
    names(xorder) <- standat$id_codes

    # actual plotting
    plot(0, 0, xlim = xlim, ylim = c(0.5, n * yaxs_exp), type = "n",
         xlab = "gregariousness", axes = FALSE, ylab = "")
    axis(1)

    cnt <- 0
    for (i in order(xorder)) {
      cnt <- cnt + 1
      if (has_cors) {
        p <- alldraws[, paste0("indi_soc_vals[", i, ",", beh, "]")]
      } else {
        p <- alldraws[, paste0("indi_soc_vals[", i, "]")]
      }

      p <- density(p, adjust = dens_adjust)
      px <- p$x
      py <- p$y
      py <- py * vert_exp
      if (i %in% sel_subset) {
        polygon(x = c(px, rev(px)), y = c(py + cnt, rep(cnt, length(py))),
                col = grey(0.2, 0.2), border = NA)
        points(px, py + cnt, col = grey(0, 0.8), type = "l")
        if (add_median) {
          xsel <- which.min(abs(px - xorder[i]))
          points(c(px[xsel], px[xsel]), c(cnt, cnt + py[xsel]), type = "l")
        }
        axis(2, at = cnt, labels = names(standat$id_codes)[i], lwd = 0,
             las = 1, cex.axis = labcex)
      }

    }
  }

  if (!greg) {
    n <- standat$n_dyads
    if (is.null(sel_subset)) sel_subset <- seq_len(n)
    # xorder <- rep(0, n)

    # order according to posterior median
    xorder <- as.numeric(apply(alldraws, 2, median))
    # nm <- combn(names(standat$id_codes), 2)
    # nm <- paste0(nm[1, ], "_@_", nm[2, ])
    nm <- apply(standat$dyads_navi, 1, function(x) paste(names(standat$id_codes)[x], collapse = "_@_"))
    names(xorder) <- nm

    # actual plotting
    plot(0, 0, xlim = xlim, ylim = c(0.5, n * yaxs_exp), type = "n",
         xlab = "pairwise affinity", axes = FALSE, ylab = "")
    axis(1)

    cnt <- 0
    for (i in order(xorder)) {
      cnt <- cnt + 1
      if (has_cors) {
        p <- alldraws[, paste0("dyad_soc_vals[", i, ",", beh, "]")]
      } else {
        p <- alldraws[, paste0("dyad_soc_vals[", i, "]")]
      }


      p <- density(p, adjust = dens_adjust)
      px <- p$x
      py <- p$y
      py <- py * vert_exp
      if (i %in% sel_subset) {
        polygon(c(px, rev(px)), c(py + cnt, rep(cnt, length(py))),
                col = grey(0.2, 0.2), border = NA)
        points(px, py + cnt, col = grey(0, 0.8), type = "l")
        if (add_median) {
          xsel <- which.min(abs(px - xorder[i]))
          points(c(px[xsel], px[xsel]), c(cnt, cnt + py[xsel]), type = "l")
        }
        axis(2, at = cnt, labels = nm[i], lwd = 0, las = 1, cex.axis = labcex)
      }

    }
  }

  invisible(round(xorder, 4))
}
```
