## Supplementary material for "(De)composing sociality: disentangling individual-specific from dyad-specific propensities to interact": bamoso R package: sim_data_whitehead2015.R

```
#' simulate data following Whitehead and James 2015
#'
#' @param n_ind numeric, default is 20: number of individuals
#' @param affi_prob numeric (default is 0.09): probability rho of a
#'                  pair being designated a pair of affiliates
#' @param greg_max numeric (default is 2): max value for individual
#'                 gregariousness ranging from 1 to greg_max
#' @param n_periods numeric (default is 50): number of sampling periods
#' @param switch_prb numeric (default is 0.1): probability of each each
#'                   individual to leave or enter from one sampling period
#'                   to the next
#' @param asso_prob_max numeric (default is 0.6): scaling factor that set
#'                      max value for association probability
#' @param sigma_fac numeric (default is 0.9): effect of sex (homophily)
#' @param phi_fac numeric (default is 2): effect of affiliation (actually
#'                effect of associating given a pair is affiliated via
#'                \code{affi_prob})
#' @param ignore_temporal logical (default is \code{FALSE}): if \code{TRUE}
#'                        then all individuals were present in all sampling
#'                        periods (overrides \code{switch_prb})
#'
#' @details some modifications(?) to deal with special cases that were not
#'          mentioned in the manuscript:
#'
#'          (1) The code makes sure that in each period there are at least 2
#'          individuals present.
#'
#'          (2) If at the end there are cases where an individual was never
#'          observed, these individuals are removed.
#'
#' @return a list
#'
#' @references
#'
#' \insertRef{whitehead2015}{bamoso}
#'
#' @export
#'

# test values
# sigma_fac = 0.9
# phi_fac = 2
# n_ind = 7
# affi_prob = 0.2
# greg_max = 2
# n_periods = 3
# switch_prb = 0.1
# asso_prob_max = 0.6

sim_data_whitehead2015 <- function(n_ind = 20,
                                   affi_prob = 0.09,
                                   greg_max = 2,
                                   n_periods = 50,
                                   switch_prb = 0.1,
                                   sigma_fac = 0.9,
                                   phi_fac = 2,
                                   asso_prob_max = 0.6,
                                   ignore_temporal = FALSE) {
  # initial setup
  n_dyads <- n_ind * (n_ind - 1) * 0.5

  # dyads
  dyads <- t(combn(seq_len(n_ind), 2))
  colnames(dyads) <- c("i1", "i2")

  # was a dyad affiliates?
  dyads <- cbind(dyads,
                 is_affi = sample(c(0, 1),
                                  size = n_dyads,
                                  replace = TRUE,
                                  prob = c(1 - affi_prob, affi_prob)))

  # individual gregariousness (g_i and g_j) and sex
  indis <- cbind(i = seq_len(n_ind),
                 greg = runif(n_ind, 1, greg_max),
                 sex = sample(c(0, 1), n_ind, TRUE),     # 50/50
                 present = sample(c(0, 1), n_ind, TRUE)) # .5 in initial period
  # make sure that at least 2 individuals were present
  done <- sum(indis[, "present"]) >= 2
  while (!done) {
    indis[, "present"] <- sample(c(0, 1), n_ind, TRUE)
    done <- sum(indis[, "present"]) >= 2
  }

  # add to dyadic data
  dyads <- cbind(dyads,
                 i1_sex = indis[dyads[, 1], "sex"],
                 i2_sex = indis[dyads[, 2], "sex"])
  dyads <- cbind(dyads, same_sex = dyads[, "i1_sex"] == dyads[, "i2_sex"])

  dyads <- cbind(dyads,
                 i1_greg = indis[dyads[, "i1"], "greg"],
                 i2_greg = indis[dyads[, "i2"], "greg"])

  dyads <- cbind(dyads,
                 i1_present = indis[dyads[, "i1"], "present"],
                 i2_present = indis[dyads[, "i2"], "present"])

  # temporal component
  pmat <- matrix(ncol = n_periods, nrow = n_ind, 0)

  if (ignore_temporal) {
    pmat[, ] <- 1
  } else {
    pmat[, 1] <- indis[, "present"]
    for (i in 2:n_periods) {
      cur <- pmat[, i - 1]
      # make sure that always at least 2 individuals are present
      newp <- pmat[, i]
      done <- sum(newp) >= 2
      while (!done) {
        for (k in seq_len(n_ind)) {
          if (sample(c(TRUE, FALSE), 1, prob = c(switch_prb, 1 - switch_prb))) {
            newp[k] <- ifelse(cur[k] == 1, 0, 1)
            newp[k] <- ifelse(cur[k] == 0, 1, 0)
          } else {
            newp[k] <- cur[k]
          }
        }
        pmat[, i] <- newp

        done <- sum(newp) >= 2
        if (!done) newp <- pmat[, i]
      }
    }
  }

  assolist <- vector(mode = "list", length = n_periods)

  for (i in 1:n_periods) {
    # update presence
    dyads[, "i1_present"] <- pmat[, i][dyads[, "i1"]]
    dyads[, "i2_present"] <- pmat[, i][dyads[, "i2"]]

    # make the probability
    part1 <- dyads[, "i1_greg"] *
      dyads[, "i2_greg"] *
      dyads[, "i1_present"] *
      dyads[, "i2_present"]
    part2 <- (1 + sigma_fac * dyads[, "same_sex"]) *
      (1 + phi_fac * dyads[, "is_affi"])
    tot_prob <- part1 * part2
    # scale to a_max
    tot_prob <- (tot_prob / max(tot_prob)) * asso_prob_max
    # sample dyads
    s <- rbinom(nrow(dyads), 1, prob = tot_prob)
    if (sum(s) >= 1) {
      # generate output
      temp <- matrix(ncol = n_ind, nrow = sum(s), 0)

      temp[cbind(seq_len(sum(s)), dyads[as.logical(s), "i1"])] <- 1
      temp[cbind(seq_len(sum(s)), dyads[as.logical(s), "i2"])] <- 1

      assolist[[i]] <- cbind(period = i, temp)
    }

  }

  assolist <- do.call("rbind", assolist)
  period_indicator <- assolist[, 1]
  assolist <- assolist[, -1]

  if (any(colSums(assolist) == 0)) {
    xcols <- which(colSums(assolist) == 0)
    assolist <- assolist[, -c(xcols)]
    pmat <- pmat[-c(xcols), ]
    rem <- which(dyads[, 1] %in% xcols | dyads[, 2] %in% xcols)
    dyads <- dyads[-c(rem), ]
  }

  list(assolist = assolist,
       period_indicator = period_indicator,
       pmat = pmat,
       dyads = dyads,
       indis = indis)
}
```
