## Supplementary material for "(De)composing sociality: disentangling individual-specific from dyad-specific propensities to interact": bamoso R package: sim_foo.R

```
#' run a single simulation on artificial data
#'
#' fit the model with simulated data and store results to file
#'
#' @param ... arguments for \code{\link{generate_data}}
#' @param indi_sd numeric, SD for gregariousness (defaults to
#'                \code{runif(1, 0.1, 2)})
#' @param dyad_sd numeric, SD for dyadic affinity (defaults to
#'                \code{runif(1, 0.1, 2)})
#' @param intercepts numeric of length two, determines the range for
#'                        intercepts. Defaults to \code{c(-2, 2)}. How many
#'                        intercepts are needed is determined inside the
#'                        function.
#' @param n_ids see \code{\link{generate_data}}
#' @param n_beh see \code{\link{generate_data}}
#' @param disp_pars_gamma see \code{\link{generate_data}}
#' @param disp_pars_beta see \code{\link{generate_data}}
#' @param outpath character, output path (defaults to R's \code{\link{tempdir}})
#' @param outfilename character, output file name (defaults to a random string)
#' @param silent logical, suppress all textual output
#' @param diagnostics logical, report problems with sampling
#' @param parallel_chains number of parallel processes
#' @param refresh refresh argument
#' @param iter_warmup iter_warmup argument
#' @param iter_sampling iter_sampling argument
#' @param testing_mode logical, default is \code{FALSE}. If \code{TRUE}: runs
#'          the model as prior simulation which illustrates how the function
#'          works while taking less time.
#'
#' @importFrom utils capture.output
#'
#' @return writes an rds file to the specified location, which contains a list
#'         with the data generation settings, generated data,
#'         fitted model's draws, and seed used during the fitting.
#'         Most importantly, the list contains a one-line data frame
#'         (\code{$results}) with the key results used for downstream
#'         analyses (see Details section).
#'
#' @details
#' The function starts by generating interaction data across variable settings
#' (see \code{\link{generate_data}}). Then it fits the model to these data.
#' From the model results it extracts the following quantities, which are in
#' the \code{$results} list item.
#'
#' * number of individuals
#' * number of behaviors
#' * true and estimated (posterior median) gregariousness SD
#' * true and estimated (posterior median) affinity SD
#' * correlation between true and estimated (posterior medians) of individual
#' gregariousness values
#' * correlation between true and estimated (posterior medians) of dyadic
#' affinity values
#' * number of posterior samples
#' * sampler diagnostics (divergent transitions etc)
#'
#'
#'
#' @export
#' @examples
#' \dontrun{
#' # path to results
#' # outpath <- "~/Desktop/bamosoeval" # probably doesn't exist on your computer
#' outpath <- tempdir() # default in function
#'
#' # use testing mode (faster, but only as demo)
#' for (i in 1:5) {
#'   sim_foo(silent = TRUE, outpath = outpath, testing_mode = TRUE)
#' }
#'
#' # analyse (read inidivual sim runs and collate to single data frame)
#' l <- list.files(outpath, pattern = ".rds", full.names = TRUE)
#' xdata <- sapply(l, function(x) {
#'   zz <- readRDS(x)
#'   zz$results
#' }, simplify = FALSE)
#' xdata <- do.call("rbind", xdata)
#'
#' # plot results
#' plot(xdata$indi_sd, xdata$indi_sd_est,
#'      xlab = 'true gregariousness SD', ylab = 'estimated SD')
#' abline(0, 1)
#' plot(xdata$intercept1, xdata$intercept1_est,
#'      xlab = 'true intercept (behavior 1)', ylab = 'estimated intercept (behavior 1)')
#' abline(0, 1)
#' }

# indi_sd = runif(1, 0.1, 2)
# dyad_sd = runif(1, 0.1, 2)
# n_beh = sample(3, 1)
# intercepts = runif(n_beh, -2, 1)
# n_ids = sample(5:30, 1)
# disp_pars_gamma = runif(n_beh, 1, 20)
# disp_pars_beta = runif(n_beh, 5, 30)
# outpath = NULL
# outfilename = NULL
# silent = FALSE
# diagnostics = TRUE
# parallel_chains = 4
# refresh = 200
# iter_warmup = 2000
# iter_sampling = 1000
# testing_mode = FALSE

sim_foo <- function(...,
                    n_ids = sample(5:30, 1),
                    indi_sd = runif(1, 0.1, 3),
                    dyad_sd = runif(1, 0.1, 3),
                    n_beh = sample(3, 1),
                    intercepts = runif(n_beh, -3, 1),
                    disp_pars_gamma = runif(n_beh, 1, 20),
                    disp_pars_beta = runif(n_beh, 5, 30),
                    outpath = NULL,
                    outfilename = NULL,
                    silent = TRUE,
                    diagnostics = TRUE,
                    parallel_chains = 1,
                    refresh = 0,
                    iter_warmup = 2000,
                    iter_sampling = 1000,
                    testing_mode = FALSE
                    ) {

  if (testing_mode) {
    if (interactive()) {
      message("you are in testing mode: the posteriors are not conditioned on the data")
    }
  }

  behav_types <- sample(c("count", "prop", "dur_gamma", "dur_beta"), n_beh, replace = TRUE)

  # occasionally, an empty matrix is produced (mostly in the gamma setting)
  good_to_go <- FALSE
  while (!good_to_go) {
    sdata <- generate_data(n_beh = n_beh,
                           n_ids = n_ids,
                           behav_types = behav_types,
                           beh_intercepts = intercepts,
                           indi_sd = indi_sd,
                           dyad_sd = dyad_sd,
                           disp_pars_gamma = disp_pars_gamma,
                           disp_pars_beta = disp_pars_beta,
                           # ...
                           )
    if (!sdata$input_data$empty) good_to_go <- TRUE
  }

  # ignore data when matrices with only 0s were generated
  # this check should be redundant, but better safe than sorry
  if (sdata$input_data$empty) {
    cat("empty matrix generated: skipped\n")
    return(NULL)
  }


  # fit the model
  seed <- sample(1e+7, 1)

  if (silent) {
    r <- sociality_model(standat = sdata$standat,
                         silent = TRUE,
                         show_messages = FALSE,
                         diagnostics = NULL,
                         parallel_chains = parallel_chains,
                         refresh = refresh,
                         iter_warmup = iter_warmup,
                         iter_sampling = iter_sampling,
                         adapt_delta = 0.98,
                         max_treedepth = 14,
                         seed = seed,
                         prior_sim = testing_mode
                         )
  } else {
    r <- sociality_model(standat = sdata$standat,
                         silent = FALSE,
                         show_messages = FALSE,
                         diagnostics = NULL,
                         parallel_chains = parallel_chains,
                         refresh = refresh,
                         iter_warmup = iter_warmup,
                         iter_sampling = iter_sampling,
                         adapt_delta = 0.98,
                         max_treedepth = 14,
                         seed = seed,
                         prior_sim = testing_mode
                         )
  }


  # extract results
  xres_draws <- r$mod_res$draws(format = "draws_matrix")
  # clean up draws (remove irrelevant pars)
  xres_draws <- xres_draws[, !grepl("log_lik", colnames(xres_draws))]
  xres_draws <- xres_draws[, !grepl("interactions_pred_cont", colnames(xres_draws))]
  xres_draws <- xres_draws[, !grepl("interactions_pred", colnames(xres_draws))]
  xres_draws <- xres_draws[, !grepl("scaled_indi_sums", colnames(xres_draws))]
  xres_draws <- xres_draws[, !grepl("dyad_soc_vals_z", colnames(xres_draws))]
  xres_draws <- xres_draws[, !grepl("indi_soc_vals_z", colnames(xres_draws))]
  colnames(xres_draws)

  indi_columns <- grepl("indi_soc_vals[", colnames(xres_draws), fixed = TRUE)
  cor_indi <- cor(sdata$input_data$indi_data$indi_soc_vals,
                  apply(xres_draws[, indi_columns], 2, median))
  dyad_columns <- grepl("dyad_soc_vals[", colnames(xres_draws), fixed = TRUE)
  cor_dyad <- cor(sdata$input_data$dyad_data$dyad_soc_vals,
                  apply(xres_draws[, dyad_columns], 2, median))

  d <- r$mod_res$diagnostic_summary(quiet = TRUE)


  if (is.null(outfilename)) {
    fn <- paste(sample(c(letters, 0:9), 10, TRUE), collapse = "")
    fn <- paste0(fn, ".rds")
  }

  # collect relevant info
  ## width of posteriors for soc components
  w1 <- quantile(c(xres_draws[, "indi_soc_sd"]), 0.0945) - quantile(c(xres_draws[, "indi_soc_sd"]), 0.055)
  w2 <- quantile(c(xres_draws[, "dyad_soc_sd"]), 0.0945) - quantile(c(xres_draws[, "dyad_soc_sd"]), 0.055)


  xres_results <- data.frame(label = fn,
                             n_ids = sdata$standat$n_ids,
                             n_beh = n_beh,
                             indi_sd = sdata$input_data$indi_sd,
                             indi_sd_est = median(xres_draws[, "indi_soc_sd"]),
                             width_indisd = w1,
                             dyad_sd = sdata$input_data$dyad_sd,
                             dyad_sd_est = median(xres_draws[, "dyad_soc_sd"]),
                             width_dyadsd = w2,
                             intercept1 = intercepts[1],
                             intercept2 = intercepts[2],
                             intercept3 = intercepts[3],
                             intercept1_est = median(xres_draws[, "beh_intercepts[1]"]),
                             intercept2_est = ifelse(n_beh >= 2, median(xres_draws[, "beh_intercepts[2]"]), NA),
                             intercept3_est = ifelse(n_beh >= 3, median(xres_draws[, "beh_intercepts[3]"]), NA),
                             cor_indi = cor_indi,
                             cor_dyad = cor_dyad,
                             tot_dur = round(r$mod_res$time()$total, 2),
                             n_samples = nrow(xres_draws),
                             divergent = sum(d$num_divergent),
                             treedepth = sum(d$num_max_treedepth),
                             ebfmi = sum(d$ebfmi < 0.2),
                             seed = seed,
                             behav_types = paste(sort(behav_types), collapse = ";")
  )

  if (is.null(outpath)) {
    op <- normalizePath(tempdir())
  } else {
    op <- normalizePath(outpath)
  }
  outloc <- file.path(op, fn)
  res <- list(sdata = sdata, # xres = xres_summary,
              results = xres_results,
              diagnostics = d,
              draws = xres_draws,
              seed = seed)
  suppressWarnings(saveRDS(res, file = outloc))

  dres <- "all good"
  if (any(d$num_divergent > 0)) dres <- "sampling problems"
  if (any(d$num_max_treedepth > 0)) dres <- "sampling problems"
  if (any(d$ebfmi < 0.2)) dres <- "sampling problems"
  if (!silent) {
    if (diagnostics) cat(dres, "\n")
  }

  # return NULL or file path?
  if (silent) {
    return(invisible(outloc))
  }
  outloc
}
```
