## Supplementary material for "(De)composing sociality: disentangling individual-specific from dyad-specific propensities to interact": bamoso R package: sociality_model.R

```
#' fit dyadic relationship model to one or more interaction matrices
#'
#' @param standat stan data list (typically the result of a call
#'          to \code{\link{make_stan_data_from_matrices}} or
#'          \code{\link{make_stan_data_from_association}})
#' @param sans_dyadic logical (default is \code{FALSE}). If \code{TRUE}
#'          fit the simple model without the dyadic components.
#' @param prior_sim logical, run only prior sims (default
#'          is \code{FALSE})
#' @param silent logical, try to suppress *all* output (default
#'          is \code{FALSE})
#' @param ... further arguments for \code{\link[cmdstanr]{sample}} (typically
#'          \code{chains}, \code{parallel_chains}, \code{refresh},
#'          \code{iter_warmup}, \code{iter_sampling}, \code{seed},
#'          \code{adapt_delta}, and/or \code{step_size})
#'
#'
#' @return a list (of class \code{dyadicmodel}) which contains the standata
#'         list as first item and the CmdStanFit as second item
#' @export
#'
#' @examples
#'
#' \dontrun{
#' data("grooming")
#' mat <- grooming$ass$groom
#' obseff <- grooming$ass$obseff
#' standat <- make_stan_data_from_matrices(mats = list(groom = mat),
#'                                         behav_types = "count",
#'                                         obseff = list(obseff))
#' res <- sociality_model(standat = standat, parallel_chains = 4,
#'                        adapt_delta = 0.9, seed = 123)
#' summary(res)
#' }
#'
#'
#' \dontrun{
#' x <- generate_data(n_ids = 16, n_beh = 1, indi_sd = 1,
#'                    dyad_sd = 1, beh_intercepts = 1)
#' standat <- x$standat
#' res <- sociality_model(standat = standat, parallel_chains = 4)
#' }
#'
#' \dontrun{
#' # with correlated axes
#' m1 <- matrix(c(0.3, -0.5, -0.5, 0.5), ncol = 2)
#' m2 <- matrix(c(0.5, 0.5, 0.5, 0.5), ncol = 2)
#' x <- generate_data(n_ids = 17, n_beh = 2,
#'                    behav_types = c("count", "prop"),
#'                    indi_sd = m1,
#'                    dyad_sd = m2,
#'                    beh_intercepts = c(1.4, -0.7), exact = TRUE)
#'
#' standat <- x$standat
#' standat$n_cors
#' res <- sociality_model(standat = standat, parallel_chains = 4)
#' res$mod_res$summary("cors_indi")
#' res$mod_res$summary("cors_dyad")
#' }
#'
#' x <- generate_data(n_ids = 5, indi_covariate_slope = 0.7, beh_intercepts = 2)

sociality_model <- function(standat,
                            sans_dyadic = FALSE,
                            prior_sim = FALSE,
                            silent = FALSE,
                            ...) {

  # determine which model...
  modeltype <- "simple" # default
  standat$prior_only <- as.integer(prior_sim)

  # detect predictors if present
  flag_indi_cat_pred <- "indi_cat_pred" %in% names(standat)
  flag_indi_covariate_pred <- "indi_covariate_pred" %in% names(standat)
  flag_dyad_cat_pred <- "dyad_cat_pred" %in% names(standat)
  flag_dyad_covariate_pred <- "dyad_covariate_pred" %in% names(standat)
  # flag_indi_cat_pred <- as.integer(!isTRUE(standat$indi_cat_pred == 0))
  # flag_indi_covariate_pred <- as.integer(!isTRUE(standat$indi_covariate_pred == 0))
  # flag_dyad_cat_pred <- as.integer(!isTRUE(standat$dyad_cat_pred == 0))
  # flag_dyad_covariate_pred <- as.integer(!isTRUE(standat$dyad_covariate_pred == 0))

  if (sum(flag_indi_cat_pred, flag_indi_covariate_pred,
          flag_dyad_cat_pred, flag_dyad_covariate_pred) > 0) {
    modeltype <- "with_preds"

    # modify flags in standat
    standat$do_indi_cat <- flag_indi_cat_pred
    standat$do_indi_covariate <- flag_indi_covariate_pred
    standat$do_dyad_cat <- flag_dyad_cat_pred
    standat$do_dyad_covariate <- flag_dyad_covariate_pred

    # reset empty data vector to 'correct length'
    if (!flag_indi_cat_pred) {
      standat$indi_cat_pred <- rep(0, standat$n_ids)
    }
    if (!flag_indi_covariate_pred) {
      standat$indi_covariate_pred <- rep(0, standat$n_ids)
    }
    if (!flag_dyad_cat_pred) {
      standat$dyad_cat_pred <- rep(0, standat$n_dyads)
    }
    if (!flag_dyad_covariate_pred) {
      standat$dyad_covariate_pred <- rep(0, standat$n_dyads)
    }
  }

  if (standat$n_cor > 0) {
    modeltype <- "cor_mod"
  }

  if (sans_dyadic) {
    modeltype <- "sans_dyadic"
  }

  if (!silent) {
    if (interactive() && modeltype != "simple") {
      cat("fitting the following non-standard model:", shQuote(modeltype), "\n")
    }
  }

  if (silent) {
    swallow1 <- capture.output(suppressMessages(mod <- get_model(type = modeltype)))
    swallow2 <- capture.output(suppressMessages(res <- mod$sample(data = standat, ...)))
  } else {
    mod <- get_model(type = modeltype)
    res <- mod$sample(data = standat, ...)
  }

  out <- list(standat = standat, mod_res = res, modeltype = modeltype)
  class(out) <- "dyadicmodel"
  out
}
```
