## Supplementary material for "(De)composing sociality: disentangling individual-specific from dyad-specific propensities to interact": bamoso R package: sociality_plot.R

```
#' plot posteriors of individual and dyadic sociality SDs (or correlations)
#'
#' @param mod_res model result from \code{\link{sociality_model}}
#' @param xlim limits for horizontal axis
#' @param labs named list with labels for the individual- and
#'             dyad-specific component
#' @param do_legend logical, should the legend be plotted
#'                  (default is \code{TRUE})
#' @param do_dyadic logical, should the posterior for the dyad-level
#'                  estimate be plotted (default is \code{TRUE})
#' @param do_indi logical, should the posterior for the individual-level
#'                estimate be plotted (default is \code{TRUE})
#' @param which_beh integer of length one. If supplied and if the underlying
#'                  model was fitted with multiple correlated behaviors,
#'                  this argument indicates which behavior is to be plotted.
#' @param which_cor integer of length two. If supplied and if the underlying
#'                  model was fitted with multiple correlated behaviors,
#'                  then this argument indicates the *pair* of behaviors
#'                  for which the correlation posteriors should be displayed.
#' @param xcols vector of length two, with the colors to use default are two
#'              Zissou colors (blueish and yellow/gold)
#' @param add_prior logical, default is \code{FALSE}. Should the prior be added
#'                  to the plot. Careful: for now the prior is hardcoded in
#'                  this function and doesn't take its value from
#'                  \code{mod_res}! (its value is \code{student_t(3, 0, 1)}).
#'
#' @return a plot
#' @export
#'
#' @importFrom brms rstudent_t
#' @importFrom grDevices hcl.colors
#' @importFrom graphics legend polygon
#' @examples
#' \dontrun{
#' # with correlated axes
#' m <- matrix(c(0.3, 0.7, 0.3, -0.5,
#'               0.7, 1.8, 0.5, -0.2,
#'               0.3, 0.5, 1.5, 0.5,
#'               -0.5, -0.2, 0.5, 1.1), ncol = 4)
#' set.seed(1)
#' x <- generate_data(n_ids = 17, n_beh = 4,
#'                    behav_types = c("count", "prop", "count", "prop"),
#'                    indi_sd = m,
#'                    dyad_sd = m,
#'                    beh_intercepts = c(1.4, -0.7, -1, 0.2))
#' res <- sociality_model(standat = x$standat, parallel_chains = 4, seed = 42)
#' sociality_plot(res, which_cor = c(2,3), xlim = c(-1, 1)) # true 0.7 for both
#' }
#'

sociality_plot <- function(mod_res,
                           xlim = c(0, 4),
                           labs = list(indi = "gregariousness",
                                       dyad = "affinity"),
                           do_legend = TRUE,
                           do_dyadic = TRUE,
                           do_indi = TRUE,
                           which_beh = NULL,
                           which_cor = NULL,
                           xcols = NULL,
                           add_prior = FALSE
                           ) {

  if (is.null(xcols)) xcols <- c("#3B99B1B3", "#EACB2BB3")
  # hcl.colors(3, palette = "zissou1", alpha = 0.7)[1:2]


  if (!is.null(which_beh) && !is.null(which_cor)) {
    stop("can only supply which_beh *OR* which_cor", call. = FALSE)
  }

  standat <- mod_res$standat
  bnames <- names(standat$beh_names)

  mod_res <- mod_res$mod_res


  is_a_cor_plot <- FALSE

  if (!is.null(which_cor)) {
    which_cor <- sort(unique(which_cor))
    if (standat$n_cors == 0) {
      stop ("can't plot correlation because only one axis fitted", call. = FALSE)
    }
    if (length(which_cor) != 2) {
      stop ("need to select two behaviors for correlation", call. = FALSE)
    }
    if (max(which_cor) > standat$n_beh) {
      stop ("can't find the correlation", call. = FALSE)
    }
    m <- matrix(0, ncol = standat$n_beh, nrow = standat$n_beh)
    m[upper.tri(m)] <- seq_len(sum(upper.tri(m)))
    index <- m[which_cor[1], which_cor[2]]
    p1 <- density(as.numeric(mod_res$draws(paste0("cors_indi[", index, "]"))))
    p2 <- density(as.numeric(mod_res$draws(paste0("cors_dyad[", index, "]"))))
    is_a_cor_plot <- TRUE
    if (identical(xlim, c(0, 4))) xlim <- c(-1, 1)
    corname <- paste(bnames[which_cor], collapse = " * ")
  } else {
    if (!is.null(which_beh)) {
      if (length(which_beh) != 1) {
        stop ("need to select *one* behaviour", call. = FALSE)
      }
      if (standat$n_cors == 0) {
        stop ("can't select behaviour because only one axis fitted", call. = FALSE)
      }
      p1 <- density(as.numeric(mod_res$draws(paste0("indi_soc_sd[", which_beh, "]"))))
      p2 <- density(as.numeric(mod_res$draws(paste0("dyad_soc_sd[", which_beh, "]"))))
    } else {
      p1 <- density(as.numeric(mod_res$draws("indi_soc_sd")))
      p2 <- density(as.numeric(mod_res$draws("dyad_soc_sd")))
    }
  }

  xxlab <- "estimated SD"
  if (is_a_cor_plot) xxlab <- "estimated correlation"

  plot(0, 0, xlim = xlim, ylim = c(0, max(p1$y, p2$y) * 1.05),
       type = "n", xaxs = "i", yaxs = "i", xlab = xxlab,
       ylab = "density", axes = FALSE, main = "")
  if (do_legend) {
    tit <- ""
    if (!is.null(which_beh)) {
      tit <- bnames[which_beh]
    }
    if (is_a_cor_plot) tit <- corname

    # legend with or without prior?
    leg_labs <- c(labs$indi, labs$dyad)
    leg_cols <- xcols[1:2]
    leg_pch <- c(15, 15)
    leg_lty <- c(FALSE, FALSE)
    if (add_prior) {
      leg_labs <- c(leg_labs, "prior")
      leg_cols <- c(leg_cols, "grey")
      leg_pch <- c(leg_pch, NA)
      leg_lty <- c(leg_lty, 1)
    }

    legend("topright",
           legend = leg_labs,
           col = leg_cols,
           pch = leg_pch,
           lty = leg_lty,
           bty = "n",
           cex = 0.7,
           title = tit)
  }

  if (do_indi) polygon(p1, border = NA, col = xcols[1])
  if (do_dyadic) polygon(p2, border = NA, col = xcols[2])

  axis(1)

  if (add_prior) {
    if (is.null(which_cor)) {
      pr <- brms::rstudent_t(4000, 3, 0, 1)
      pr <- density(pr, adjust = 2)
      polygon(pr, border = "grey")
    }
    if (!is.null(which_cor)) {
      aux <- sample(lkjpriors[, "2"], 1000) # that's the current prior in interactions_cor
      polygon(density(aux, adjust = 1.2), border = 'grey')
    }

  }

  box(bty = "l")
}
```
