## Supplementary material for "(De)composing sociality: disentangling individual-specific from dyad-specific propensities to interact": bamoso R package: summary.dyadicmodel.R

```
#' summarize dyadicmodel object
#'
#' @param object an object of class \code{"dyadicmodel"},
#'               usually the result of a call to \code{\link{sociality_model}}
#' @param ... further arguments passed to or from other methods (ignored)
#' @importFrom stats median quantile
#' @author Christof Neumann
#'
#' @export

summary.dyadicmodel <- function(object, ...) {
  nchains <- length(object$mod_res$metadata()$id)
  sperchain <- object$mod_res$metadata()$iter_sampling

  has_correlations <- object$standat$n_cors > 0

  sans_dyadic <- FALSE
  if (!has_correlations) {
    if ("dyad_soc_sd" %in% object$mod_res$metadata()$model_params) {
      d <- object$mod_res$draws(c("indi_soc_sd", "dyad_soc_sd"),
                                format = "draws_matrix")
      qs_i <- sprintf("%.2f", quantile(d[, "indi_soc_sd"], c(0.055, 0.5, 0.945)))
      qs_d <- sprintf("%.2f", quantile(d[, "dyad_soc_sd"], c(0.055, 0.5, 0.945)))
    } else {
      sans_dyadic <- TRUE
      d <- object$mod_res$draws(c("indi_soc_sd"),
                                format = "draws_matrix")
      qs_i <- sprintf("%.2f", quantile(d[, "indi_soc_sd"], c(0.055, 0.5, 0.945)))
      # qs_d <- sprintf("%.2f", quantile(d[, "dyad_soc_sd"], c(0.055, 0.5, 0.945)))
    }


  }
  if (has_correlations) {
    d <- object$mod_res$draws(c("indi_soc_sd", "dyad_soc_sd"),
                              format = "draws_matrix")

    qs_i <- apply(d[, grepl("indi_soc_sd", colnames(d))], 2, quantile, c(0.055, 0.5, 0.945))
    qs_i <- apply(qs_i, 1, function(x) sprintf("%.2f", x))
    qs_d <- apply(d[, grepl("dyad_soc_sd", colnames(d))], 2, quantile, c(0.055, 0.5, 0.945))
    qs_d <- apply(qs_d, 1, function(x) sprintf("%.2f", x))

    # and correlations
    d <- object$mod_res$draws(c("cors_indi", "cors_dyad"),
                              format = "draws_matrix")

    qs_i_cors <- apply(d[, grepl("cors_indi", colnames(d)), drop = FALSE], 2, quantile, c(0.055, 0.5, 0.945))
    qs_i_cors <- apply(qs_i_cors, 1, function(x) sprintf("%.2f", x))
    qs_d_cors <- apply(d[, grepl("cors_dyad", colnames(d))], 2, quantile, c(0.055, 0.5, 0.945))
    qs_d_cors <- apply(qs_d_cors, 1, function(x) sprintf("%.2f", x))
    if (is.null(nrow(qs_i_cors))) {
      qs_i_cors <- matrix(qs_i_cors, ncol = 3)
    }
    if (is.null(nrow(qs_d_cors))) {
      qs_d_cors <- matrix(qs_d_cors, ncol = 3)
    }
  }


  s <- paste0("Model of dyadic interactions from ", object$standat$n_ids,
              " individuals (", object$standat$n_dyads, " dyads)")
  cat(s, "\n")
  if ("removed_dyads" %in% names(object$standat)) {
    cat("(removed", nrow(object$standat$removed_dyads), "dyads with NA values)\n")
  }

  cat("number of post-warmup samples:", nchains * sperchain, "\n")

  if (!has_correlations) {
    cat("--------------------\n")
    cat("sociality estimates:\n")
    cat("  individual gregariousness SD (median [89% CI]):",
        qs_i[2], paste0("[", qs_i[1], " - ", qs_i[3], "]\n"))

    if (sans_dyadic) {
      cat("  ~~no pairwise affinity estimated~~...\n        ")
    } else {
      cat("  pairwise affinity SD (median [89% CI]):        ",
          qs_d[2], paste0("[", qs_d[1], " - ", qs_d[3], "]\n"))
    }

    cat("--------------------\n")
  }
  if (has_correlations) {
    cat("--------------------\n")
    cat("sociality estimates (by behavior):\n")
    for (i in seq_len(object$standat$n_beh)) {
      s <- paste0("(", i, "): ",
                  shQuote(names(object$standat$beh_names)[i]), " (type: ",
                  shQuote(names(object$standat$behav_types)[i]), "):")
      cat(s, "\n")
      cat("     individual gregariousness SD (median [89% CI]):",
          qs_i[i, 2], paste0("[", qs_i[i, 1], " - ", qs_i[i, 3], "]\n"))
      cat("     pairwise affinity SD (median [89% CI]):        ",
          qs_d[i, 2], paste0("[", qs_d[i, 1], " - ", qs_d[i, 3], "]\n"))
    }
    xindex <- which(upper.tri(matrix(ncol = object$standat$n_beh, nrow = object$standat$n_beh)), arr.ind = TRUE)
    xnames <- names(object$standat$beh_names)
    xnames <- apply(xindex, 1, function(x) paste(xnames[x[1]], xnames[x[2]], sep = " * "))
    cat("--------------------\n")
    cat("correlations (individual gregariousness):\n")
    for (i in seq_len(nrow(qs_i_cors))) {
      cat("  ", xnames[i], "(median [89% CI]):",
          qs_i_cors[i, 2], paste0("[", qs_i_cors[i, 1], " - ", qs_i_cors[i, 3], "]\n"))
    }
    cat("--------------------\n")
    cat("correlations (dyadic affinity):\n")
    for (i in seq_len(nrow(qs_d_cors))) {
      cat("  ", xnames[i], "(median [89% CI]):",
          qs_d_cors[i, 2], paste0("[", qs_d_cors[i, 1], " - ", qs_d_cors[i, 3], "]\n"))
    }
    cat("--------------------\n")
  }


  cat(object$standat$n_beh, "behavior(s) included:\n")
  for (i in seq_len(object$standat$n_beh)) {
    s <- paste0("(", i, "): ",
                shQuote(names(object$standat$beh_names)[i]), " (type: ",
                shQuote(names(object$standat$behav_types)[i]), ")")
    cat(s, "\n")
  }

  cat("--------------------\n")
  diagnostics <- object$mod_res$diagnostic_summary(quiet = TRUE)
  if (all(diagnostics$num_divergent == 0) &&
      all(diagnostics$num_max_treedepth == 0) &&
      all(diagnostics$ebfmi > 0.2)) {
    diags <- TRUE
    msg <- "no obvious sampling issues detected"
  } else {
    diags <- FALSE
    msg <- "sampling issues detected (see below)"
  }
  if (diags) cat(msg, "\n")
  if (!diags) {
    cat(msg, "\n")
    if (interactive()) {
      print(diagnostics)
    }
  }
  cat("\n")

}
```
