## Supplementary material for "(De)composing sociality: disentangling individual-specific from dyad-specific propensities to interact": bamoso R package: zzz.R

```
utils::globalVariables("lkjpriors")

#' grooming matrices and observation effort for four macaque species
#'
#' @format a list with four items. Each item is a list itself with one grooming
#'         matrix and a vector or matrix for observation effort.
#' @source
#' \itemize{
#'  \item{\emph{Macaca assamensis}:} {Oliver Schülke and Julia Ostner}
#'  \item{\emph{M. fuscata}:} {Andrew MacIntosh}
#'  \item{\emph{M. nigra}:} {Julie Duboscq}
#'  \item{\emph{M. sylvanus}:} {Julia Fischer}
#' }
"grooming"

#' socprog example: raw association data and metrics calculated within SOCPROG.
#'
#' @references
#'
#' \insertRef{whitehead2008}{bamoso}
#'
#' \insertRef{whitehead2015}{bamoso}
#'
#' @format a list with two items. First item are the raw associations,
#'         second are the different metrics returned by SOCPROG.
"socprogexample"

#' data for workflow example
#'
#'
#' @format a list with named items.
#' @examples
#' set.seed(755907)
#' workflow_example <- generate_data(n_ids = 25,
#'                                   n_beh = 1,
#'                                   behav_types = "count",
#'                                   indi_sd = 0.9,
#'                                   dyad_sd = 2.3,
#'                                   beh_intercepts = -1.5)
#' indivals <- workflow_example$input_data$indi_soc_vals
#' workflow_example$response <- rpois(n = workflow_example$standat$n_ids,
#'                                    lambda = exp(-0.5 + 1.2 * indivals))
#'
"workflow_example"

#' data for kangaroo reproductive success
#' @details
#' \code{kangaroos_rawdata} contains a subset 47 individuals of the raw data
#' taken from the primary sources (see references). It is formatted as a list.
#' The first item contains as a list the raw association data (one matrix
#' per year). The second item lists the fitness outcomes and values for
#' control variables for each individual in each year.
#'
#' \code{kangaroos_subset} contains a list derived from the association data
#' and the fitness outcome taken from \code{kangaroos_rawdata}. It also contains
#' the values of the variables included in the model that were not directly
#' quantified from the association data (parity status, residual group size,
#' number of sightings). It is formatted such that it can be passed to Stan.
#'
#' @aliases kangaroos_rawdata
#'
#' @references
#' \insertRef{menz2020}{bamoso}
#'
#' @source
#' \insertRef{menz2020data}{bamoso}
#'
#' @format a list with named items to be passed to Stan.
"kangaroos_subset"

#' data for shark associations
#' @details
#' \code{sharks} contains association data on 83 individually identified
#' blacktip reef sharks (\emph{Carcharhinus melanopterus}).
#' This is a subset of the original data, which was constrained to
#' only contain individuals for which relatedness data is available.
#'
#'
#' @references
#' \insertRef{mourier2021}{bamoso}
#'
#' @source
#' \insertRef{mourier2020}{bamoso}
#'
#' @format a list with named items.
"sharks"

#' crested macaque data
#' @details
#' \code{nigra2} contains grooming and proximity data for 19 female crested
#' macaques (\emph{Macaca nigra}).
#'
#'
#' @format a list with named items.
#' @examples
#' \dontrun{
#' s <- make_stan_data_from_matrices(mats = list(groom = nigra2$groom,
#'                                               prox = nigra2$prox),
#'                                   behav_types = c("count", "prop"),
#'                                   obseff = list(nigra2$obseff_groom,
#'                                                 nigra2$obseff_prox),
#'                                   correlations = TRUE)
#'
#' r <- sociality_model(s, parallel_chains = 4, seed = 17, adapt_delta = 0.9)
#' summary(r)
#' model_summary(r)
#' }
#'
"nigra2"


#' utility to convert observation effort that is vector-based or NULL to list of matrices
#'
#' @param obseff list with vectors of observation effort or \code{NULL}
#' @param mats list with matrices of dyadic interactions
#'
#' @return a list
#' @examples
#' obseff <- list(c(a = 2.3, b = 4.3, c = NA, d = 12),
#'                c(a = 2.3, b = 4.3, c = 222, d = 12))
#' bamoso:::obseff_to_matrix(obseff = obseff)
#'
obseff_to_matrix <- function(obseff, mats = NULL) {
  if (length(mats) > 1 & (is.vector(obseff, mode = "numeric") || is.vector(obseff, mode = "integer"))) {
    aux <- outer(obseff, obseff, "+")
    res <- list()
    for (i in seq_along(mats)) {
      res[[length(res) + 1]] <- aux
    }
    return(res)
  }


  if (is.null(obseff)) {
    res <- lapply(mats, function(x) {
      x[, ] <- 1
      diag(x) <- NA
      x[lower.tri(x)] <- 0
      x
    })

    for (i in seq_len(length(res))) {
      res[[i]][is.na(mats[[i]])] <- NA
    }

    return(res)
  }

  if (all(unlist(lapply(obseff, is.vector)))) {
    (res <- lapply(obseff, function(x)outer(x, x, "+")))
    res <- lapply(res, function(x) {
      diag(x) <- NA
      x[lower.tri(x)] <- 0
      x
    })
    return(res)
  }

  if (all(unlist(lapply(obseff, is.matrix)))) {
    return(obseff)
  }

  stop("can't decide what to do with the supplied observation effort (either provide matrices, or vectors, or keep it at NULL", call. = FALSE)

}


#' generate multivariate normally distributed numbers
#'
#' (based on MASS::mvrorm and lme4::sdcor2cov)
#'
#' @param n number of samples to generate
#' @param mu vector with means
#' @param Sigma square matrix with SDs on diagonal and cors on off-diagonal
#' @param empirical,tol see help of \code{MASS::mvrnorm()} (\code{?MASS::mvrnorm})
#'
#' @source Code is reused from \code{MASS} and \code{lme4} packages.
#'
#' @details
#' \code{Sigma} is a SD/correlation matrix (unlike mvrnorm, which uses variance/covariance)
#'
#'
#' @return a matrix
#' @examples
#' smat <- matrix(c(1.3, -0.2, -0.2, 0.5), ncol = 2)
#' res <- bamoso:::rnorm_multi(n = 20, mu = c(-0.5, 1.7), Sigma = smat)
#' apply(res, 2, mean)
#' apply(res, 2, sd)
#' cor(res)
#'
#' # univariate with SD=0.6:
#' res <- bamoso:::rnorm_multi(n = 20, mu = c(-0.5), Sigma = matrix(0.6))
#' apply(res, 2, mean)
#' apply(res, 2, sd)
#' cor(res)

rnorm_multi <- function(n, mu = 0, Sigma = matrix(1), empirical = TRUE, tol = 1e-06) {

  # lme4::sdcor2cov
  xsd <- diag(Sigma)
  diag(Sigma) <- 1
  Sigma <- Sigma * outer(xsd, xsd)

  # MASS::mvrnorm
  p <- length(mu)
  if (!all(dim(Sigma) == c(p, p))) {
    stop("incompatible arguments")
  }
  eS <- eigen(Sigma, symmetric = TRUE)
  ev <- eS$values
  if (!all(ev >= -tol * abs(ev[1L]))) {
    stop("'Sigma' is not positive definite")
  }

  X <- matrix(rnorm(p * n), n)
  if (empirical) {
    X <- scale(X, TRUE, FALSE)
    X <- X %*% svd(X, nu = 0)$v
    X <- scale(X, FALSE, TRUE)
  }
  X <- drop(mu) + eS$vectors %*% diag(sqrt(pmax(ev, 0)), p) %*% t(X)
  t(X)
}


#' link function for probabilities with observation effort
#' @param x linear predictor (unconstrained)
#' @param obseff positive observation effort (offset)
lin2prob <- function(x, obseff) {
  1 - exp(-exp(x) * obseff)
}

#' link function for probabilities with observation effort
#' @param x probability predictor (constrained)
#' @param obseff positive observation effort (offset)
prob2lin <- function(x, obseff) {
  log((-log(1 - x))/obseff)
}
```
