## Supplementary material for "(De)composing sociality: disentangling individual-specific from dyad-specific propensities to interact": bamoso R package: README.md

<!-- README.md is generated from README.Rmd. Please edit that file -->

# `bamoso`: Bayesian modelling of animal social interactions

<img src="man/figures/README-eq.png" width="100%" />

supported data types for observed interactions:

- frequency (Poisson)

- discrete proportion (Binomial)

- continuous proportion (Beta)

- duration (Gamma)

- binary (Bernoulli) (experimental)

- duration with zeros (mixture Bernoulli/Gamma) (experimental)

## Installation

You need [`cmdstanr`](https://mc-stan.org/cmdstanr/) in order to install
and run `bamoso`. This in turn requires a working C++ toolchain first.
Check out the [getting
started-guide](https://mc-stan.org/cmdstanr/articles/cmdstanr.html) from
`cmdstanr` to see how to get everything set up properly. Also [this
document](https://mc-stan.org/docs/cmdstan-guide/cmdstan-installation.html#cpp-toolchain)
might be helpful.

Then install `cmdstanr`.

``` r
install.packages("cmdstanr", repos = c("https://mc-stan.org/r-packages/", getOption("repos")))
```

And you also need the `remotes` packages, which is easy to install from
CRAN:

``` r
install.packages("remotes")
```

Then check whether things are set up correctly:

``` r
library(cmdstanr)
check_cmdstan_toolchain(fix = TRUE)
```

You may need to follow any instructions that the output of that call
gives. If you haven’t used `cmdstanr` before, it’s likely that you at
least need to install `cmdstan` itself:

``` r
cmdstanr::install_cmdstan()
```

If this gives positive feedback, install `bamoso`:

``` r
library(remotes)
remotes::install_github("gobbios/bamoso", dependencies = TRUE, build_vignettes = FALSE)
```

If you want to install (recompile) the intro vignette, use:

``` r
install.packages("tinytex")
library(remotes)
remotes::install_github("gobbios/bamoso", dependencies = TRUE, build_vignettes = TRUE)
```

This will take a little more time to finish, but if successful gives you
access to the vignette:

``` r
vignette("intro", package = "bamoso")
```

## Example

<div class="figure" style="text-align: center">

<img src="man/figures/README-groomdatasetplot-1.png" alt="\label{fig:syl_net}Example data set. Grooming frequencies on the left and dyadic observation hours on the right." width="60%" />
<p class="caption">

Example data set. Grooming frequencies on the left and dyadic
observation hours on the right.
</p>

</div>

``` r
library(bamoso)
```

``` r
data("grooming")
groom_mat <- grooming$syl$groom
obseff_mat <- grooming$syl$obseff
```

``` r
standat <- make_stan_data_from_matrices(mats = list(groom = groom_mat),
                                        behav_types = "count",
                                        obseff = list(obseff_mat))
```

``` r
res <- sociality_model(standat)
```

``` r
summary(res)
#> Model of dyadic interactions from 19 individuals (171 dyads) 
#> (removed 0 dyads with NA values)
#> number of post-warmup samples: 4000 
#> --------------------
#> sociality estimates:
#>   individual gregariousness SD (median [89% CI]): 0.99 [0.55 - 1.50]
#>   pairwise affinity SD (median [89% CI]):         1.84 [1.56 - 2.18]
#> --------------------
#> 1 behavior(s) included:
#> (1): 'groom' (type: 'count') 
#> --------------------
#> no obvious sampling issues detected
```

``` r
pp_model(res, xlab = "grooming bouts", ylab = "(predicted) frequency",
         xbreaks = 20)
pp_model_stat(res, stat = "max", xlab = "maximum number of grooming bouts",
              main = "")
```

<div class="figure" style="text-align: center">

<img src="man/figures/README-ppcheck-1.png" alt="\label{fig:syl_net_pp}Posterior predictive checks for Barbary macaque grooming data. On the left: bars represent observed grooming frequencies. The circles represent the median frequencies predicted by the model and the vertical bars the 89\% interval around these predictions. One the right: the histogram is the posterior distribution of the maximum number of grooming bouts predicted by the model across all posterior samples. The red line indicates the observed value in the data." width="80%" />
<p class="caption">

Posterior predictive checks for Barbary macaque grooming data. On the
left: bars represent observed grooming frequencies. The circles
represent the median frequencies predicted by the model and the vertical
bars the 89% interval around these predictions. One the right: the
histogram is the posterior distribution of the maximum number of
grooming bouts predicted by the model across all posterior samples. The
red line indicates the observed value in the data.
</p>

</div>

``` r
sociality_plot(res, add_prior = TRUE)
```

<div class="figure" style="text-align: center">

<img src="man/figures/README-socplot-1.png" alt="\label{fig:syl_net_soc}Posteriors for the two sociality components. Variation among dyads is larger than variation among individuals." width="50%" />
<p class="caption">

Posteriors for the two sociality components. Variation among dyads is
larger than variation among individuals.
</p>

</div>

``` r
ridge_plot(res, greg = TRUE)
ridge_plot(res, greg = FALSE, sel_subset = 10:20, vert_exp = 15)
```

<div class="figure" style="text-align: center">

<img src="man/figures/README-ridgeplot-1.png" alt="\label{fig:syl_net_ridge}Posteriors for the two sociality components. The affinity distributions on the right only reflect a subset of 10 dyads." width="80%" />
<p class="caption">

Posteriors for the two sociality components. The affinity distributions
on the right only reflect a subset of 10 dyads.
</p>

</div>

## Acknowledgments

This work was supported by Deutsche Forschungsgemeinschaft, Grant/Award
Number: 254142454 / GRK 2070.

<img src="man/figures/RTG-LogoCYMK300.png" width="50%" style="display: block; margin: auto;" />
