## Supplementary material for "(De)composing sociality: disentangling individual-specific from dyad-specific propensities to interact": bamoso R package: testthat.R

```
# This file is part of the standard setup for testthat.
# It is recommended that you do not modify it.
#
# Where should you do additional test configuration?
# Learn more about the roles of various files in:
# * https://r-pkgs.org/testing-design.html#sec-tests-files-overview
# * https://testthat.r-lib.org/articles/special-files.html

library(testthat)
library(bamoso)

test_check("bamoso")
```
