## Supplementary material for "(De)composing sociality: disentangling individual-specific from dyad-specific propensities to interact": bamoso R package: test-check_data.R

```
x <- generate_data(n_beh = 2, n_ids = 16, beh_intercepts = runif(2, -1, 1),
                   behav_types = sample(c("count", "prop", "dur_gamma", "dur_beta"), 2))
mats <- x$processed$interaction_matrices
mats[[2]] <- mats[[2]][, -1]

test_that("check_data throws error if dimension mismatch", {
  expect_error(check_data(mats = mats))
})

# generate a falsely ordered obseff
m <- mats[[1]]
colnames(m) <- rownames(m) <- letters[seq_len(nrow(m))]
o <- m
rownames(o) <- sample(rownames(o))
test_that("check_data throws message if names mismatch", {
  expect_message(check_data(mats = list(m), behav_types = "prop", obseff = list(o)),
                 regexp = "name mismatch found between or within observation effort data (not good)",
                 fixed = TRUE)
})


# recognize individuals with all NA values
test_that("check_data throws error if all interactions are NA for an individual", {
  x <- generate_data(n_beh = 2, n_ids = 16, beh_intercepts = runif(2, -1, 1),
                     behav_types = sample(c("count", "prop", "dur_gamma", "dur_beta"), 2))
  mats <- x$processed$interaction_matrices[[1]]
  mats[, 4] <- NA
  mats[4, ] <- NA
  expect_message(check_data(mats = list(mats)),
                 regexp = "found *individuals* in interaction data for which *all* dyadic behavior values are NA",
                 fixed = TRUE)
})
```
