## Supplementary material for "(De)composing sociality: disentangling individual-specific from dyad-specific propensities to interact": bamoso R package: test-cor_plot.R

```
test_that("cor_plot doesn't crash", {
  capture.output(suppressMessages(model <- fit_example_model(1)))
  expect_null(cor_plot(model, asp = 1, xlab = "A", ylab = "B"))
  expect_null(cor_plot(model, greg = FALSE, xlim = c(-3, 3), asp = 1))
})
```
