## Supplementary material for "(De)composing sociality: disentangling individual-specific from dyad-specific propensities to interact": bamoso R package: test-extract_samples.R

```
good_to_go <- FALSE
ni <- sample(6:12, 1)
while (!good_to_go) {
  x <- generate_data(n_beh = 2, n_ids = ni, dyad_sd = 1, indi_sd = 1, beh_intercepts = runif(2, -1, 1),
                     behav_types = sample(c("count", "prop", "dur_gamma", "dur_beta"), 2))
  if (!x$input_data$empty) good_to_go <- TRUE
}

standat <- x$standat
suppressMessages(z <- capture.output(r <- sociality_model(standat, parallel_chains = 4, adapt_delta = 0.8, show_exceptions = FALSE, show_messages = FALSE, refresh = 0)))


test_that("extract_samples returns the correct amount of columns", {
  e <- extract_samples(r, what = "indi_vals")
  expect_true(ncol(e) == ni)
  e <- extract_samples(r, what = "dyad_vals")
  expect_true(ncol(e) == (ni * (ni - 1)/2))
  e <- extract_samples(r, what = "beh_intercepts")
  expect_true(ncol(e) == 2)
  e <- extract_samples(r, what = c("indi_sd", "dyad_sd"))
  expect_true(ncol(e) == 2)
})


# correlation model

good_to_go <- FALSE
ni <- sample(4:6, 1)
while (!good_to_go) {
  x <- generate_data(n_beh = 4, n_ids = ni, dyad_sd = 1, indi_sd = 1, beh_intercepts = runif(4, -1, 1),
                     behav_types = c("count", "count", "count", "count"))
  if (!x$input_data$empty) good_to_go <- TRUE
}


standat <- make_stan_data_from_matrices(list(A = x$processed$interaction_matrices[[1]], B = x$processed$interaction_matrices[[2]],
                                             C = x$processed$interaction_matrices[[3]], D = x$processed$interaction_matrices[[4]]),
                             correlations = TRUE)

suppressMessages(z <- capture.output(r <- sociality_model(standat, parallel_chains = 4, adapt_delta = 0.8, show_exceptions = FALSE, show_messages = FALSE, refresh = 0)))


test_that("extract_samples returns the correct amount of columns even with cors", {
  e <- extract_samples(r, what = "indi_cors")
  expect_true(ncol(e) == 6)
  e <- extract_samples(r, what = "dyad_cors")
  expect_true(ncol(e) == 6)

})
```
