## Supplementary material for "(De)composing sociality: disentangling individual-specific from dyad-specific propensities to interact": bamoso R package: test-fit_example_model.R

```
test_that("example models fit...", {
  capture.output(suppressMessages(r1 <- fit_example_model(1)))
  capture.output(suppressMessages(r2 <- fit_example_model(2)))
  capture.output(suppressMessages(r3 <- fit_example_model(3)))
  capture.output(suppressMessages(r4 <- fit_example_model(4)))
  capture.output(suppressMessages(r5 <- fit_example_model(5)))
  capture.output(suppressMessages(r6 <- fit_example_model(6)))

  expect_s3_class(r1, "dyadicmodel")
  expect_s3_class(r2, "dyadicmodel")
  expect_s3_class(r3, "dyadicmodel")
  expect_s3_class(r4, "dyadicmodel")
  expect_s3_class(r5, "dyadicmodel")
  expect_s3_class(r6, "dyadicmodel")
})
```
