## Supplementary material for "(De)composing sociality: disentangling individual-specific from dyad-specific propensities to interact": bamoso R package: test-gamma0.R

```
# single run for testing, extend to check parameter recovery during development
# devtools::load_all()
n <- 2

test_pars <- data.frame(r = seq_len(n),
                        do1 = c(FALSE, TRUE), # TRUE
                        # do_mix = sample(c(TRUE, FALSE), n, TRUE),
                        n_ind = sample(c(16, 20, 24), n, TRUE), # 8
                        int_gamma = runif(n, -2, 2),
                        shape_gamma = runif(n, 1, 20),
                        int_bern = runif(n, -3, 1), # that's the range
                        int_pois = runif(n, -2, 2),
                        indi_sd = runif(n, 0.8, 1.5),
                        dyad_sd = runif(n, 0.8, 1.5)
)
table(test_pars$n_ind, test_pars$do1)
est_pars <- test_pars - test_pars
est_pars$r <- NULL
est_pars$do1 <- test_pars$do1
est_pars$do_mix <- test_pars$do_mix
est_pars$n_ind <- test_pars$n_ind
est_pars$n_div <- 0
est_pars$n_dep <- 0
est_pars$n_ebfmi <- 0
est_pars$int_pois[est_pars$do1] <- NA
est_pars$done <- FALSE
est_pars$bern_prior <- NA

i=1
for (i in sample(seq_len(n))) {
  if (est_pars$done[i]) next
  if (test_pars$do1[i]) {
    xx <- generate_data(n_ids = test_pars$n_ind[i], n_beh = 1, behav_types = c("dur_gamma0"),
                        indi_sd = test_pars$indi_sd[i], dyad_sd = test_pars$dyad_sd[i],
                        beh_intercepts = test_pars$int_bern[i],
                        beh_intercepts_add = test_pars$int_gamma[i], count_obseff = c(0.1, 11),
                        exact = TRUE, disp_pars_gamma = c(test_pars$shape_gamma[i]))
  } else {
    xx <- generate_data(n_ids = test_pars$n_ind[i], n_beh = 2, behav_types = c("dur_gamma0", "count"),
                        indi_sd = test_pars$indi_sd[i], dyad_sd = test_pars$dyad_sd[i],
                        beh_intercepts = c(test_pars$int_bern[i], test_pars$int_pois[i]),
                        beh_intercepts_add = c(test_pars$int_gamma[i], 0), count_obseff = c(0.1, 11),
                        exact = TRUE, disp_pars_gamma = c(test_pars$shape_gamma[i], 0))

  }


  # mm <- cmdstan_model("inst/extdata/interaction_model.stan")
  d <- xx$standat
  est_pars$bern_prior[i] <- d$prior_matrix[1, 1]
  xres <- sociality_model(standat = d, parallel_chains = 4, show_exceptions = F, refresh = 0, silent = TRUE)
  # r <- mm$sample(d, parallel_chains = 4,  show_exceptions = F, refresh = 0)
  r <- xres$mod_res
  di <- r$diagnostic_summary(quiet = TRUE)
  est_pars$n_div[i] <- sum(di$num_divergent)
  est_pars$n_dep[i] <- sum(di$num_max_treedepth)
  est_pars$n_ebfmi[i] <- sum(di$ebfmi < 0.3)

  # digress, look at plots:
  if (FALSE) {
    edata <- extract_samples(xres, c("indi_vals", "dyad_vals"))

    zz <- xx$input_data$dyad_data
    rownames(zz) <- paste0(zz$id1, "_@_", zz$id2)
    r <- sample(nrow(edata), 1)
    plot(zz$dyad_soc_vals, edata[r, rownames(zz)])
    cor(zz$dyad_soc_vals, edata[r, rownames(zz)])

    zz <- xx$input_data$indi_data
    r <- sample(nrow(edata), 1)
    rownames(zz) <- paste0(zz$id)
    plot(zz$indi_soc_vals, edata[r, rownames(zz)])
    cor(zz$indi_soc_vals, edata[r, rownames(zz)])

  }


  temp <- as.numeric(d$interactions_cont[, 1] > 0)
  qlogis(mean(temp))
  zz <- data.frame(r$summary(c("beh_intercepts", "beh_intercepts_add", "shapes_gamma", "indi_soc_sd", "dyad_soc_sd")))
  zz

  postpairs <- r$draws(c("beh_intercepts[1]", "shapes_gamma_raw[1]", "beh_intercepts_add[1]"), format = "draws_matrix")
  colnames(postpairs) <- c("int_bern", 'shape_raw', "int_gamma")
  pairs(postpairs)
  est_pars$int_bern[i] <- zz$median[zz$variable == "beh_intercepts[1]"]
  est_pars$shape_gamma[i] <- zz$median[zz$variable == "shapes_gamma[1]"]
  est_pars$int_gamma[i] <- zz$median[zz$variable == "beh_intercepts_add[1]"]
  if (!test_pars$do1[i]) est_pars$int_pois[i] <- zz$median[zz$variable == "beh_intercepts[2]"]
  est_pars$indi_sd[i] <- zz$median[zz$variable == "indi_soc_sd"]
  est_pars$dyad_sd[i] <- zz$median[zz$variable == "dyad_soc_sd"]
  est_pars$done[i] <- TRUE
  cat(sum(est_pars$done), "\n")
}


if (FALSE) {
  do_leg <- function() legend("topleft",
                              legend = levels(as.factor(ori$n_ind)),
                              col = hcl.colors(n = nlevels(as.factor(ori$n_ind)), palette = "viridis"),
                              pch = 16)

  ori <- test_pars[est_pars$done, ]
  est <- est_pars[est_pars$done, ]

  ori <- ori[est$n_div <= 10, ]
  est <- est[est$n_div <= 10, ]

  cols <- hcl.colors(n = nlevels(as.factor(ori$n_ind)), palette = "viridis")[as.factor(ori$n_ind)]

  plot(est$bern_prior, est$int_bern, pch = c(16, 1)[est$do1 + 1], col = cols)
  abline(0, 1)

  plot(ori$indi_sd, est$indi_sd, pch = c(16, 1)[est$do1 + 1], col = cols)
  abline(0, 1)
  do_leg()
  plot(ori$dyad_sd, est$dyad_sd, pch = c(16, 1)[est$do1 + 1], col = cols)
  abline(0, 1)
  do_leg()

  plot(ori$int_gamma, est$int_gamma, pch = c(16, 1)[est$do1 + 1], col = cols)
  do_leg()
  abline(0, 1)

  plot(ori$int_bern, est$int_bern, pch = c(16, 1)[est$do1 + 1], col = cols)
  abline(0, 1)
  do_leg()

  plot(ori$int_pois, est$int_pois, pch = c(16, 1)[est$do1 + 1], col = cols)
  abline(0, 1)
  do_leg()

  plot(ori$shape_gamma, est$shape_gamma, pch = c(16, 1)[est$do1 + 1], col = cols)
  abline(0, 1)
  do_leg()

  # plot(est$int_gamma, est$shape_gamma, pch = c(16, 1)[est$do1 + 1], col = cols)
  # plot(est$int_bern, est$shape_gamma, pch = c(16, 1)[est$do1 + 1], col = cols)

}
```
