## Supplementary material for "(De)composing sociality: disentangling individual-specific from dyad-specific propensities to interact": bamoso R package: test-gamma.R

```
# single run for testing, extend to check parameter recovery during development

n <- 1 #

test_pars <- data.frame(r = seq_len(n),
                        n_ind = sample(c(8, 16, 24, 32), n, TRUE),
                        int_gamma = runif(n, -2, 2),
                        shape_gamma = runif(n, 1, 10),
                        int_pois = runif(n, -2, 2),
                        indi_sd = runif(n, 0.2, 1.2),
                        dyad_sd = runif(n, 0.2, 1.2)
)
est_pars <- test_pars - test_pars
est_pars$r <- NULL
est_pars$do1 <- test_pars$do1
est_pars$n_ind <- test_pars$n_ind
est_pars$n_div <- 0
est_pars$n_dep <- 0
est_pars$n_ebfmi <- 0
est_pars$done <- FALSE

i=2
for (i in sample(seq_len(n))) {
  if (est_pars$done[i]) next

  xx <- generate_data(n_ids = test_pars$n_ind[i], n_beh = 2, behav_types = c("dur_gamma", "count"),
                      indi_sd = test_pars$indi_sd[i], dyad_sd = test_pars$dyad_sd[i],
                      beh_intercepts = c(test_pars$int_gamma[i], test_pars$int_pois[i]),
                      beh_intercepts_add = c(NA, NA),
                      exact = TRUE, disp_pars_gamma = c(test_pars$shape_gamma[i], 0))


  # mm <- cmdstan_model("inst/extdata/interaction_model.stan")
  d <- xx$standat
  d$prior_only <- 0
  # r <- mm$sample(d, parallel_chains = 4,  show_exceptions = F, refresh = 0)
  xres <- sociality_model(standat = d, parallel_chains = 4,  show_exceptions = F, refresh = 0)
  r <- xres$mod_res

  di <- r$diagnostic_summary(quiet = TRUE)
  est_pars$n_div[i] <- sum(di$num_divergent)
  est_pars$n_dep[i] <- sum(di$num_max_treedepth)
  est_pars$n_ebfmi[i] <- sum(di$ebfmi < 0.3)

  zz <- data.frame(r$summary(c("beh_intercepts", "beh_intercepts_add", "shapes_gamma", "indi_soc_sd", "dyad_soc_sd")))
  zz

  est_pars$shape_gamma[i] <- zz$median[zz$variable == "shapes_gamma[1]"]
  est_pars$int_gamma[i] <- zz$median[zz$variable == "beh_intercepts[1]"]
  est_pars$int_pois[i] <- zz$median[zz$variable == "beh_intercepts[2]"]
  est_pars$indi_sd[i] <- zz$median[zz$variable == "indi_soc_sd"]
  est_pars$dyad_sd[i] <- zz$median[zz$variable == "dyad_soc_sd"]
  est_pars$done[i] <- TRUE
  cat(i, "\n")
}


if (FALSE) {
  do_leg <- function() legend("topleft",
                              legend = levels(as.factor(ori$n_ind)),
                              col = hcl.colors(n = nlevels(as.factor(ori$n_ind)), palette = "viridis"),
                              pch = 16)

  ori <- test_pars[est_pars$done, ]
  est <- est_pars[est_pars$done, ]

  ori <- ori[est$n_div <= 10, ]
  est <- est[est$n_div <= 10, ]

  cols <- hcl.colors(n = nlevels(as.factor(ori$n_ind)), palette = "viridis")[as.factor(ori$n_ind)]

  plot(ori$indi_sd, est$indi_sd, pch = 16, col = cols)
  abline(0, 1)
  do_leg()
  plot(ori$dyad_sd, est$dyad_sd, pch = 16, col = cols)
  abline(0, 1)
  do_leg()

  plot(ori$int_gamma, est$int_gamma, pch = c(16), col = cols)
  do_leg()
  abline(0, 1)


  plot(ori$int_pois, est$int_pois, pch = c(16), col = cols)
  abline(0, 1)
  do_leg()

  plot(ori$shape_gamma, est$shape_gamma, pch = c(16), col = cols)
  abline(0, 1)
  do_leg()

}
```
