## Supplementary material for "(De)composing sociality: disentangling individual-specific from dyad-specific propensities to interact": bamoso R package: test-generate_data.R

```
# try to consistently produce an empty matrix which should trigger a warning (if interactive)

test_that("warning/message is generated if any matrix is empty", {
  if (interactive()) {
    expect_warning(generate_data(n_beh = 4, n_ids = 6, dyad_sd = 1, indi_sd = 1, beh_intercepts = runif(4, -100, -90),
                                 behav_types = c("count", "prop", "dur_gamma", "dur_beta")))
  } else {
    expect_message(generate_data(n_beh = 4, n_ids = 6, dyad_sd = 1, indi_sd = 1, beh_intercepts = runif(4, -100, -90),
                                 behav_types = c("count", "prop", "dur_gamma", "dur_beta")))
  }

})


# make sure data are generated such that make_prior doesn't return Inf
has_inf <- data.frame(run = 1:100, count = NA, prop = NA, gamma = NA, beta = NA)
for (i in 1:100) {
  x <- suppressWarnings(generate_data(n_beh = 4, n_ids = 12, dyad_sd = 2, indi_sd = 1, beh_intercepts = runif(4, 0, 10),
                behav_types = c("count", "prop", "dur_gamma", "dur_beta")))
  if (!x$input_data$empty) {
    aux <- x$standat$prior_matrix[, 1]
    has_inf$count[i] <- is.infinite(aux[1])
    has_inf$prop[i] <- is.infinite(aux[2])
    has_inf$gamma[i] <- is.infinite(aux[3])
    has_inf$beta[i] <- is.infinite(aux[4])
  }
}

has_inf <- na.omit(has_inf)

test_that("no infinite values for priors", {
  expect_true(all(!has_inf$count))
  expect_true(all(!has_inf$prop))
  expect_true(all(!has_inf$gamma))
  expect_true(all(!has_inf$beta))
})


# error is thrown if dimensions of intercepts and SDs and correlations don't match
test_that("dimension mismatches are detected", {
  expect_error(generate_data(n_beh = 2, dyad_sd = 2, indi_sd = 1, beh_intercepts = c(3)))
  m1 <- matrix(c(0.3, -0.5, -0.5, 0.5), ncol = 2)
  m2 <- matrix(c(0.5, 0.5, 0.5, 0.5), ncol = 2)
  expect_error(generate_data(n_beh = 3, dyad_sd = m1, indi_sd = m2, beh_intercepts = c(3, 0)))
  expect_error(generate_data(n_beh = 2, dyad_sd = 1, indi_sd = m2, beh_intercepts = c(3, 0)))
  expect_error(generate_data(n_beh = 1, dyad_sd = m1, indi_sd = m2, beh_intercepts = c(3, 0)))

})


test_that("stan data from simulated data contains the correct list elements", {
  x <- suppressWarnings(generate_data(n_beh = 2, n_ids = 12, dyad_sd = 1, indi_sd = 1, beh_intercepts = runif(2, 0, 10),
                                      behav_types = c("count", "prop")))
  standat <- x$standat
  expect_true("id1" %in% names(standat))
  expect_true("id2" %in% names(standat))
  expect_true("interactions" %in% names(standat))
  expect_true("interactions_cont" %in% names(standat))
  expect_true("n_beh" %in% names(standat))
  expect_true("n_ids" %in% names(standat))
  expect_true("n_dyads" %in% names(standat))
  expect_true("dyads_navi" %in% names(standat))
  expect_true("gamma_shape_pos" %in% names(standat))
  expect_true("gamma_shape_n" %in% names(standat))
  expect_true("beta_shape_pos" %in% names(standat))
  expect_true("beta_shape_n" %in% names(standat))
  expect_true("obseff" %in% names(standat))
  expect_true("obseff_int" %in% names(standat))
  expect_true("prior_matrix" %in% names(standat))
  expect_true("prior_matrix2" %in% names(standat))
  expect_true("id_codes" %in% names(standat))
  expect_true("beh_names" %in% names(standat))
  expect_true("behav_types" %in% names(standat))
})
```
