## Supplementary material for "(De)composing sociality: disentangling individual-specific from dyad-specific propensities to interact": bamoso R package: test-get_model.R

```
library(bamoso)


# test model type from simulated data (use simple or with_preds) --------

x <- generate_data(n_ids = 5, behav_types = "prop", beh_intercepts = 0.7)
standat <- x$standat
# if predictor data is absent, fit the simple model
suppressMessages(r1 <- sociality_model(standat, chains = 1, refresh = 0, show_exceptions = FALSE))

# if predictor data is present, fit the with_preds model
x <- generate_data(n_ids = 5, behav_types = "prop", beh_intercepts = 0.7, indi_cat_slope = 1)
standat <- x$standat
suppressMessages(r2 <- sociality_model(standat, chains = 1, refresh = 0, show_exceptions = FALSE))

# if no predictor data is present, fit the simple model
standat$indi_cat_pred <- NULL
suppressMessages(r3 <- sociality_model(standat, chains = 1, refresh = 0, show_exceptions = FALSE))


test_that("model uses predictor data appropriately", {
  expect_true(r1$modeltype == "simple")
  expect_true(r2$modeltype == "with_preds")
  expect_true(r3$modeltype == "simple")
})


test_that("get_model returns a CmdStanModel", {
  m <- get_model(type = "simple")
  expect_s3_class(m, "CmdStanModel")
  m <- get_model(type = "sans_dyadic")
  expect_s3_class(m, "CmdStanModel")
  m <- get_model(type = "cor_mod")
  expect_s3_class(m, "CmdStanModel")
  m <- get_model(type = "with_preds")
  expect_s3_class(m, "CmdStanModel")
})


x <- generate_data(n_ids = 12, n_beh = 2, behav_types = c("prop", "count"), indi_sd = 1, dyad_sd = 1, indi_covariate_slope = 0.2, indi_cat_slope = -0.5, dyad_covariate_slope = 0, dyad_cat_slope = 0, beh_intercepts = c(0, 0), prop_trials = 500, count_obseff = 10)
b <- list(b1 = x$processed$interaction_matrices[[1]], b2 = x$processed$interaction_matrices[[2]])
o <- x$processed$obseff_matrices

# basic model
d1 <- make_stan_data_from_matrices(mats = b, behav_types = c("prop", "count"), obseff = o)
f1 <- sociality_model(d1, parallel_chains = 4, iter_warmup = 500, iter_sampling = 200, show_exceptions = FALSE, show_messages = FALSE, diagnostics = NULL)
xx <- capture.output(expect_null(suppressMessages(summary(f1))))

# corr model
d2 <- make_stan_data_from_matrices(mats = b, behav_types = c("prop", "count"), obseff = o, correlations = TRUE)
f2 <- sociality_model(d2, parallel_chains = 4, iter_warmup = 500, iter_sampling = 200, show_exceptions = FALSE, show_messages = FALSE, diagnostics = NULL)
xx <- capture.output(expect_null(suppressMessages(summary(f2))))

# sans dyadic model
d3 <- make_stan_data_from_matrices(mats = b, behav_types = c("prop", "count"), obseff = o)
f3 <- sociality_model(d3, parallel_chains = 4, iter_warmup = 500, iter_sampling = 200, show_exceptions = FALSE, show_messages = FALSE, diagnostics = NULL, sans_dyadic = TRUE)
xx <- capture.output(expect_null(suppressMessages(summary(f3))))

# covar model
d4 <- make_stan_data_from_matrices(mats = b, behav_types = c("prop", "count"), obseff = o, indi_cat_pred = x$input_data$indi_data$feature_cat)
f4 <- sociality_model(d4, parallel_chains = 4, iter_warmup = 500, iter_sampling = 200, show_exceptions = FALSE, show_messages = FALSE, diagnostics = NULL)
xx <- capture.output(expect_null(suppressMessages(summary(f4))))
# f4$mod_res$summary("indi_cat_eff")
```
