## Supplementary material for "(De)composing sociality: disentangling individual-specific from dyad-specific propensities to interact": bamoso R package: test-make_stan_data_from_association.R

```
x <- sim_data_whitehead2015(n_ind = 12, n_periods = 50, ignore_temporal = TRUE)
assos <- x$assolist
# head(assos)

standat <- make_stan_data_from_association(assos)

test_that("make_stan_data_from_association does not produce NA values", {
  expect_true(all(unlist(lapply(standat, function(x) !any(is.na(x))))))
})

test_that("make_stan_data_from_association contains the correct list elements", {
  expect_true("id1" %in% names(standat))
  expect_true("id2" %in% names(standat))
  expect_true("interactions" %in% names(standat))
  expect_true("interactions_cont" %in% names(standat))
  expect_true("n_beh" %in% names(standat))
  expect_true("n_ids" %in% names(standat))
  expect_true("n_dyads" %in% names(standat))
  expect_true("dyads_navi" %in% names(standat))
  expect_true("gamma_shape_pos" %in% names(standat))
  expect_true("gamma_shape_n" %in% names(standat))
  expect_true("beta_shape_pos" %in% names(standat))
  expect_true("beta_shape_n" %in% names(standat))
  expect_true("obseff" %in% names(standat))
  expect_true("obseff_int" %in% names(standat))
  expect_true("prior_matrix" %in% names(standat))
  expect_true("prior_matrix2" %in% names(standat))
  expect_true("id_codes" %in% names(standat))
  expect_true("beh_names" %in% names(standat))
  expect_true("behav_types" %in% names(standat))
})
```
