## Supplementary material for "(De)composing sociality: disentangling individual-specific from dyad-specific propensities to interact": bamoso R package: test-make_stan_data_from_matrices.R

```
# occasionally, an empty matrix is produced (mostly in the gamma setting)
good_to_go <- FALSE
while (!good_to_go) {
  x <- generate_data(n_beh = 2, dyad_sd = 1, indi_sd = 1, beh_intercepts = runif(2, -1, 1),
                     behav_types = sample(c("count", "prop", "dur_gamma", "dur_beta"), 2))
  if (!x$input_data$empty) good_to_go <- TRUE
}

x$input_data$empty
x$input_data$behav_type
bmats <- list(b1 = x$processed$interaction_matrices[[1]],
              b2 = x$processed$interaction_matrices[[2]])


standat <- make_stan_data_from_matrices(mats = bmats,
                                        behav_types = x$input_data$behav_type)

test_that("make_stan_data_from_matrices does not produce NA values", {
  expect_true(all(unlist(lapply(standat, function(x) !any(is.na(x))))))
})


test_that("make_stan_data_from_matrices removes NA dyads appropriately", {
  m <- x$processed$interaction_matrices[[1]]
  standat <- make_stan_data_from_matrices(mats = list(m))
  nd <- standat$n_dyads
  rem <- sample(ncol(m), 3)

  m[rem[1], rem[2]] <- NA
  m[rem[2], rem[1]] <- NA
  standat <- make_stan_data_from_matrices(mats = list(m))
  expect_true(standat$n_dyads == nd - 1)
  expect_true(nrow(standat$dyads_navi) == nd - 1)
  expect_true(nrow(standat$dyads_navi) == nrow(standat$interactions))
  expect_true(nrow(standat$dyads_navi) == nrow(standat$interactions_cont))
  expect_true(nrow(standat$dyads_navi) == nrow(standat$obseff))
  expect_true(nrow(standat$dyads_navi) == nrow(standat$obseff_int))

  m[rem[1], rem[3]] <- NA
  m[rem[3], rem[1]] <- NA
  standat <- make_stan_data_from_matrices(mats = list(m))
  expect_true(standat$n_dyads == nd - 2)
  expect_true(nrow(standat$dyads_navi) == nd - 2)
  expect_true(nrow(standat$dyads_navi) == nrow(standat$interactions))
  expect_true(nrow(standat$dyads_navi) == nrow(standat$interactions_cont))
  expect_true(nrow(standat$dyads_navi) == nrow(standat$obseff))
  expect_true(nrow(standat$dyads_navi) == nrow(standat$obseff_int))

})


test_that("make_stan_data_from_matrices contains the correct list elements", {
  expect_true("id1" %in% names(standat))
  expect_true("id2" %in% names(standat))
  expect_true("interactions" %in% names(standat))
  expect_true("interactions_cont" %in% names(standat))
  expect_true("n_beh" %in% names(standat))
  expect_true("n_ids" %in% names(standat))
  expect_true("n_dyads" %in% names(standat))
  expect_true("dyads_navi" %in% names(standat))
  expect_true("gamma_shape_pos" %in% names(standat))
  expect_true("gamma_shape_n" %in% names(standat))
  expect_true("beta_shape_pos" %in% names(standat))
  expect_true("beta_shape_n" %in% names(standat))
  expect_true("obseff" %in% names(standat))
  expect_true("obseff_int" %in% names(standat))
  expect_true("prior_matrix" %in% names(standat))
  expect_true("prior_matrix2" %in% names(standat))
  expect_true("id_codes" %in% names(standat))
  expect_true("beh_names" %in% names(standat))
  expect_true("behav_types" %in% names(standat))
})


# dimensions match
nrow_interactions <- nrow(standat$interactions)
nrow_interactions_cont <- nrow(standat$interactions_cont)


## misc
test_that("make_stan_data_from_matrices contains the correct list elements", {
  expect_error(generate_data(n_beh = 1, behav_types = "dur_gamma0", count_obseff = c(-0.5, 4)))
})


## handling 0s ----
test_that("warning when all ints are 0 in dur_gamma0", {
  x <- suppressWarnings(generate_data(n_ids = 6, n_beh = 1, behav_types = "dur_gamma0", count_obseff = c(0.5, 4)))
  m <- x$processed$interaction_matrices[[1]]
  o <- x$processed$obseff_matrices[[1]]
  m <- m - m # make all zero
  expect_warning(make_stan_data_from_matrices(mats = list(m), behav_types = "dur_gamma0", obseff = list(o)))
})

test_that("warning when no 0 in dur_gamma0", {
  x <- suppressWarnings(generate_data(n_ids = 6, n_beh = 1, behav_types = "dur_gamma0", count_obseff = c(0.5, 4)))
  m <- x$processed$interaction_matrices[[1]]
  o <- x$processed$obseff_matrices[[1]]
  m[upper.tri(m)] <- m[upper.tri(m)] + 0.00001
  expect_warning(make_stan_data_from_matrices(mats = list(m), behav_types = "dur_gamma0", obseff = list(o)))
})

test_that("warning when all ints are 0 in binary", {
  x <- suppressWarnings(generate_data(n_ids = 6, n_beh = 1, behav_types = "binary", count_obseff = c(0.5, 4)))
  m <- x$processed$interaction_matrices[[1]]
  o <- x$processed$obseff_matrices[[1]]
  m <- m - m # make all zero
  expect_warning(make_stan_data_from_matrices(mats = list(m), behav_types = "binary", obseff = list(o)))
})

test_that("warning when no 0 in binary", {
  x <- suppressWarnings(generate_data(n_ids = 6, n_beh = 1, behav_types = "binary", count_obseff = c(0.5, 4)))
  m <- x$processed$interaction_matrices[[1]]
  o <- x$processed$obseff_matrices[[1]]
  m[upper.tri(m)] <- 1
  expect_warning(make_stan_data_from_matrices(mats = list(m), behav_types = "binary", obseff = list(o)))
})
```
