## Supplementary material for "(De)composing sociality: disentangling individual-specific from dyad-specific propensities to interact": bamoso R package: test-model-related.R

```
# occasionally, an empty matrix is produced (mostly in the gamma setting)
good_to_go <- FALSE
while (!good_to_go) {
  x <- generate_data(n_beh = 4, n_ids = 16, dyad_sd = 1, indi_sd = 1, beh_intercepts = runif(4, -1, 1),
                     behav_types = sample(c("count", "prop", "dur_gamma", "dur_beta"), 4))
  if (!x$input_data$empty) good_to_go <- TRUE
  # x$standat$indi_cat_pred <- 0
  # x$standat$indi_covariate_pred <- 0
  # x$standat$dyad_cat_pred <- 0
  # x$standat$dyad_covariate_pred <- 0

}

standat <- x$standat
suppressMessages(z <- capture.output(r <- sociality_model(standat,
                                                          parallel_chains = 4,
                                                          show_exceptions = FALSE,
                                                          adapt_delta = 0.9,
                                                          show_messages = FALSE,
                                                          refresh = 0)))

summary(r)


z <- model_summary(r, raw = T)
z <- model_summary(r)


# plots ------------
b_discrete <- which(x$input_data$behav_type %in% c("prop", "count"))[1]
b_cont <- which(x$input_data$behav_type %in% c("dur_gamma", "dur_beta"))[1]

pp_model_stat(mod_res = r, xvar = b_discrete, stat = "mean")
pp_model_stat(mod_res = r, xvar = b_discrete, stat = "range_width")
pp_model_stat(mod_res = r, xvar = b_discrete, stat = "iqr")
pp_model_stat(mod_res = r, xvar = b_discrete, stat = "max")
pp_model_stat(mod_res = r, xvar = b_cont, stat = "mean")
pp_model_stat(mod_res = r, xvar = b_cont, stat = "range_width")
pp_model_stat(mod_res = r, xvar = b_cont, stat = "iqr")
pp_model_stat(mod_res = r, xvar = b_cont, stat = "max")

sociality_plot(mod_res = r)


test_that("pp functions return invisible stuff", {
  expect_invisible(pp_model(mod_res = r, xvar = b_discrete, xbreaks = 100))
  expect_invisible(pp_model(mod_res = r, xvar = b_discrete, selected_id = "1"))
  expect_invisible(pp_model_dens(mod_res = r, xvar = b_cont, selected_id = "3"))
  expect_invisible(pp_model_dens(mod_res = r, xvar = b_cont, selected_id = "5"))
  expect_invisible(pp_model_dens(mod_res = r, xvar = b_cont))
  expect_invisible(ridge_plot(mod_res = r))
  expect_invisible(ridge_plot(mod_res = r, greg = FALSE, sel_subset = "6"))
})
```
