## Supplementary material for "(De)composing sociality: disentangling individual-specific from dyad-specific propensities to interact": bamoso R package: test-model_summary.R

```
# test that model_summary produces sensible results given different combinations of behaviors and correlations

test_that("model_summary works", {
  cors_indi <- matrix(c(1, 0.5, 0.5, -0.2, 0.5, 1, 0.5, 0.5, 0.5, 0.5, 1, 0.5, -0.2, 0.5, 0.5, 1), ncol = 4)
  cors_dyad <- matrix(c(1, 0.5, 0.5, -0.2, 0.5, 1, 0.5, 0.5, 0.5, 0.5, 1, 0.5, -0.2, 0.5, 0.5, 1), ncol = 4)


  upto=2
  for (upto in 2:4) {
    x <- generate_data(n_ids = 4,
                       n_beh = upto,
                       beh_intercepts = c(0.5, 0.5, 0.5, 0.5)[1:upto],
                       behav_types = c("count", "count", "count", "count")[1:upto],
                       indi_sd = cors_indi[1:upto, 1:upto],
                       dyad_sd = cors_dyad[1:upto, 1:upto]
    )

    s <- x$standat
    r <- suppressMessages(suppressWarnings(sociality_model(s, parallel_chains = 4, seed = 1, adapt_delta = 0.85, show_exceptions = FALSE, refresh = 0,
                                          iter_warmup = 500, iter_sampling = 100, show_messages = FALSE)))
    res1 <- model_summary(r, raw = TRUE)
    res2 <- model_summary(r, raw = FALSE)

    expect_true(all(unlist(strsplit(res2$beh, " * ", fixed = TRUE)) %in% names(s$beh_names)))
    expect_true(all(res2$stanvarname %in% res1$variable))
    expect_true("cor" %in% res2$categ)
    expect_true(all(!is.na(res2$label)))
    expect_true(all(!is.na(res2$categ)))
    expect_true(all(!is.na(res2$beh)))
    expect_true(all(!is.na(res2$stanvarname)))

    s$n_cors <- 0
    r <- suppressMessages(suppressWarnings(sociality_model(s, parallel_chains = 4, seed = 1, adapt_delta = 0.85, show_exceptions = FALSE, refresh = 0,
                                          iter_warmup = 500, iter_sampling = 100, show_messages = FALSE)))
    res1 <- model_summary(r, raw = TRUE)
    res2 <- model_summary(r, raw = FALSE)

    expect_true(all(is.na(res2$beh)))
    expect_true(all(res2$stanvarname %in% res1$variable))
    expect_false("cor" %in% res2$categ)
    expect_true(all(!is.na(res2$label)))
    expect_true(all(!is.na(res2$categ)))
    expect_true(all(!is.na(res2$stanvarname)))
  }

})
```
