## Supplementary material for "(De)composing sociality: disentangling individual-specific from dyad-specific propensities to interact": bamoso R package: test-pp_binary.R

```
xx <- generate_data(n_ids = 15, n_beh = 2,
                    behav_types = c("count", "binary"),
                    indi_sd = 1, dyad_sd = 2,
                    beh_intercepts = c(-1.4, -0.5))

xres <- sociality_model(standat = xx$standat, parallel_chains = 4,
                        chains = 4, iter_sampling = 350, refresh = 0,
                        show_exceptions = FALSE, adapt_delta = 0.95,
                        silent = TRUE, seed = 47)


test_that("pp_binary works", {
  ## error if behavior is not dur_gamma0 or binary
  expect_error(pp_binary(mod_res = xres, xvar = 1, all_ids = TRUE))
  ## error if more than one id
  expect_error(pp_binary(mod_res = xres, xvar = 2, selected_id = c(1, 4)))

  expect_error(pp_binary(mod_res = xres, xvar = 2, selected_id = "haha"))

  zz <- pp_binary(mod_res = xres, xvar = 2, all_ids = TRUE, n_draws = 7, selected_id = c(1))
  expect_true(is.list(zz))
  expect_true(length(zz$observed) == ncol(zz$fromdraws))

  zz <- pp_binary(mod_res = xres, xvar = 2, all_ids = FALSE, n_draws = 7)
  expect_true(is.list(zz))
  expect_true(length(zz$observed) == 1)
  expect_true(ncol(zz$fromdraws) == 1)

})
```
