## Supplementary material for "(De)composing sociality: disentangling individual-specific from dyad-specific propensities to interact": bamoso R package: test-sociality_model.R

```
test_that("silencing works", {
  s1 <- generate_data(10, behav_types = "count", beh_intercepts = 1.5,
                      indi_sd = 1, dyad_sd = 0.7)$standat
  expect_message(sociality_model(s1, chains = 1, adapt_delta = 0.5, refresh = 0))
  expect_no_message(sociality_model(s1, chains = 1, adapt_delta = 0.5, silent = TRUE))

  s2 <- generate_data(10, behav_types = "count", beh_intercepts = 1.5,
                      indi_sd = 1, dyad_sd = 0.7, indi_cat_slope = -0.2)$standat
  expect_message(sociality_model(s2, chains = 1, adapt_delta = 0.5, refresh = 0 ))
  expect_no_message(sociality_model(s2, chains = 1, adapt_delta = 0.5, silent = TRUE))
})
```
