## Supplementary material for "(De)composing sociality: disentangling individual-specific from dyad-specific propensities to interact": bamoso R package: test-sociality_plot.R

```
test_that("sociality plots", {
  capture.output(suppressMessages(r1 <- fit_example_model(1)))
  capture.output(suppressMessages(r5 <- fit_example_model(5)))
  capture.output(suppressMessages(r6 <- fit_example_model(6)))

  expect_error(sociality_plot(r5, which_cor = c(1, 2)),
               regexp = "only one axis fitted")
  expect_error(sociality_plot(r6, which_cor = c(1, 2)),
               regexp = "only one axis fitted")

  expect_error(sociality_plot(r1, which_cor = c(1, 6)),
               regexp = "can't find the correlation")
  expect_error(sociality_plot(r1, which_cor = c(2)),
               regexp = "need to select two behaviors")

  expect_error(sociality_plot(r1, which_cor = c(2), which_beh = 1),
               regexp = "can only supply") # can only supply which_beh *OR* which_cor


  sociality_plot(r1, which_cor = c(1, 2))
  sociality_plot(r1, which_cor = c(1, 2), add_prior = TRUE)
  expect_no_error(sociality_plot(r1, which_cor = c(1, 2)))
  expect_no_error(sociality_plot(r1, which_cor = c(1, 2), add_prior = TRUE))

  expect_error(sociality_plot(r1, which_beh = c(1, 2)),
               regexp = "need to select") # need to select *one* behaviour

  expect_error(sociality_plot(r6, which_beh = c(1)),
               regexp = "can't select behaviour because only one axis fitted")

  sociality_plot(r1, which_beh = c(2), add_prior = TRUE)
  expect_no_error(sociality_plot(r1, which_beh = c(2), add_prior = TRUE))

  sociality_plot(r1, add_prior = TRUE)
  expect_no_error(sociality_plot(r1, add_prior = TRUE))

  sociality_plot(r6)
  expect_no_error(sociality_plot(r6))


})
```
