## Supplementary material for "(De)composing sociality: disentangling individual-specific from dyad-specific propensities to interact": bamoso R package: test-standata_consistency.R

```
# stan data is generated currently in three functions:
# - generate_data
# - make_stan_data_from_associations
# - make_stan_data_from_matrices

# testing here whether the necessary elements are produced in each function


# no confounders -----------------------
behav_types <- sample(c("count", "prop", "dur_gamma", "dur_beta"), 2)
x <- generate_data(n_beh = 2, n_ids = 6, beh_intercepts = runif(2, -1, 1),
                   behav_types = behav_types)
mats <- x$processed$interaction_matrices

s1 <- x$standat
s2 <- make_stan_data_from_matrices(mats = mats, behav_types = behav_types)

res <- data.frame(list_item = names(s1), is_there = FALSE, is_equal = NA)

for (i in 1:nrow(res)) {
  if (res$list_item[i] %in% names(s2)) {
    res$is_there[i] <- TRUE
    res$is_equal[i] <- identical(s1[[res$list_item[i]]], s2[[res$list_item[i]]])
  }
}

test_that("data items are present in both simulated output and make_stan_data_from_matrices", {
  expect_true(all(res$is_there))
  expect_true(all(res$is_equal))
})


# and again with covariates ----
behav_types <- sample(c("count", "prop", "dur_gamma", "dur_beta"), 2)
x <- generate_data(n_beh = 2, n_ids = 6, beh_intercepts = runif(2, -1, 1),
                   behav_types = behav_types, indi_covariate_slope = 0, dyad_covariate_slope = 0, indi_cat_slope = 0, dyad_cat_slope = 0)
mats <- x$processed$interaction_matrices
s1 <- x$standat

s2 <- make_stan_data_from_matrices(mats = mats, behav_types = behav_types,
                                   indi_cat_pred = x$input_data$indi_data$feature_cat,
                                   indi_covariate_pred = x$input_data$indi_data$feature_cont,
                                   dyad_cat_pred = x$input_data$dyad_data$feature_cat,
                                   dyad_covariate_pred = x$input_data$dyad_data$feature_cont
)

# do some rounding to avoid false positives
s1$interactions_cont <- round(s1$interactions_cont, 15)
s2$interactions_cont <- round(s2$interactions_cont, 15)

res <- data.frame(list_item = names(s1), is_there = FALSE, is_equal = NA)

for (i in 1:nrow(res)) {
  if (res$list_item[i] %in% names(s2)) {
    res$is_there[i] <- TRUE
    res$is_equal[i] <- identical(s1[[res$list_item[i]]], s2[[res$list_item[i]]])
  }
}

test_that("data items are present and identical in both simulated output and make_stan_data_from_matrices", {
  expect_true(all(res$is_there))
  expect_true(all(res$is_equal))
})


# association data... -------
x <- sim_data_whitehead2015(n_ind = 5, n_periods = 9, ignore_temporal = TRUE)
assos <- x$assolist
s3 <- make_stan_data_from_association(assos)

res <- data.frame(list_item = names(s1), is_there = FALSE)
for (i in 1:nrow(res)) {
  if (res$list_item[i] %in% names(s3)) {
    res$is_there[i] <- TRUE
  }
}

test_that("data items are present in both simulated output and make_stan_data_from_association", {
  expect_true(all(res$is_there))
})


# with individual-level confounder -----------------------
behav_types <- sample(c("count", "prop", "dur_gamma", "dur_beta"), 2)
x <- generate_data(n_beh = 2, n_ids = 6, beh_intercepts = runif(2, -1, 1),
                   behav_types = behav_types, indi_cat_slope = 1)
mats <- x$processed$interaction_matrices

s1 <- x$standat
s2 <- make_stan_data_from_matrices(mats = mats, behav_types = behav_types, indi_cat_pred = x$input_data$indi_data$feature_cat)

res <- data.frame(list_item = names(s1), is_there = FALSE, is_equal = NA)

for (i in 1:nrow(res)) {
    res$is_there[i] <- TRUE
    res$is_equal[i] <- identical(s1[[res$list_item[i]]], s2[[res$list_item[i]]])
}


test_that("data items are present in both simulated output and make_stan_data_from_matrices", {
  expect_true(all(res$is_there))
})

x <- sim_data_whitehead2015(n_ind = 5, n_periods = 9, ignore_temporal = TRUE)
assos <- x$assolist
s3 <- make_stan_data_from_association(assos, indi_cat_pred = sample(c(0, 1), ncol(assos), TRUE))

res <- data.frame(list_item = names(s1), is_there = FALSE)
for (i in 1:nrow(res)) {
  if (res$list_item[i] %in% names(s3)) {
    res$is_there[i] <- TRUE
  }
}

test_that("data items are present in both simulated output and make_stan_data_from_association", {
  expect_true(all(res$is_there))
})


# simple model fit comparison ----
behav_types <- sample(c("count", "prop"), 2, TRUE)
x <- generate_data(n_beh = 2, n_ids = 5, beh_intercepts = runif(2, -1, 1),
                   behav_types = behav_types)

mats <- x$processed$interaction_matrices
s1 <- x$standat
s2 <- make_stan_data_from_matrices(mats = mats, behav_types = behav_types)
names(s1);names(s2)

suppressMessages(r1 <- sociality_model(s1, chains = 1, iter_sampling = 200, seed = 1, refresh = 0))
suppressMessages(r2 <- sociality_model(s2, chains = 1, iter_sampling = 200, seed = 1, refresh = 0))

r1 <- round(c(r1$mod_res$draws("indi_soc_sd", format = "draws_matrix")), 4)
r2 <- round(c(r2$mod_res$draws("indi_soc_sd", format = "draws_matrix")), 4)

test_that("model output is the same regardless of standat origin", {
  expect_true(all(r1 == r2))
})

#  model fit comparison with predictors ----

behav_types <- sample(c("count", "prop"), 2, TRUE)
x <- generate_data(n_beh = 2, n_ids = 5, beh_intercepts = runif(2, -1, 1),
                   behav_types = behav_types, indi_covariate_slope = 0.1, dyad_cat_slope = -0.2)

mats <- x$processed$interaction_matrices
s1 <- x$standat
s2 <- make_stan_data_from_matrices(mats = mats, behav_types = behav_types,
                                   indi_covariate_pred = x$input_data$indi_data$feature_cont,
                                   dyad_cat_pred = x$input_data$dyad_data$feature_cat)
names(s1);names(s2)

suppressMessages(r1 <- sociality_model(s1, chains = 1, iter_sampling = 200, seed = 1, refresh = 0))
suppressMessages(r2 <- sociality_model(s2, chains = 1, iter_sampling = 200, seed = 1, refresh = 0))

r1 <- round(c(r1$mod_res$draws("indi_soc_sd", format = "draws_matrix")), 4)
r2 <- round(c(r2$mod_res$draws("indi_soc_sd", format = "draws_matrix")), 4)

test_that("model output is the same regardless of standat origin", {
  expect_true(all(r1 == r2))
})
```
