## Supplementary material for "(De)composing sociality: disentangling individual-specific from dyad-specific propensities to interact": bamoso R package: covariates.Rmd

````
---
title: "\\texttt{bamoso}: models with covariates"
author: "Christof Neumann"
date: "`r Sys.Date()` (built with v. `r packageVersion('bamoso')`)"
output: 
  bookdown::pdf_document2:
    # toc: true
    toc_depth: 2
    number_sections: false
vignette: >
  %\VignetteIndexEntry{models with covariates}
  %\VignetteEncoding{UTF-8}
  %\VignetteEngine{knitr::rmarkdown}
editor_options: 
  chunk_output_type: console
link-citations: yes
linkcolor: red
geometry: margin = 0.7in
bibliography: ../inst/REFERENCES.bib
---

```{r, include = FALSE}
knitr::opts_chunk$set(
  collapse = TRUE,
  comment = "#>"
)
# csl: animal-behaviour.csl
knitr::opts_chunk$set(echo = TRUE)
options(width = 120)
def.chunk.hook  <- knitr::knit_hooks$get("chunk")
knitr::knit_hooks$set(chunk = function(x, options) {
  x <- def.chunk.hook(x, options)
  ifelse(options$size != "normalsize", paste0("\\", options$size,"\n\n", x, "\n\n \\normalsize"), x)
})
knitr::opts_chunk$set(size = 'footnotesize', fig.align = 'center')
# compile time
compile_start <- Sys.time()

```


\vspace{2ex}

This document (and package) is work in progress.
Installation instructions can be found at https://github.com/gobbios/bamoso/.
If you find bugs, mistakes or have suggestions, I'd be very happy to hear about them at https://github.com/gobbios/bamoso/issues.

```{r setup, include=FALSE}
library(bamoso)
```

# Incorporating covariates and categorical predictors

Here we illustrate how the basic sociality model can be extended. 
We start with the same premise as before:
Interactions arise because of variation in the individual gregariousness and dyadic affinity.
But now let's consider the possibility that dyadic affinities are stronger between related individuals.
Further, let's consider the possibility that there are sex differences in individual gregariousness.
Both seem like reasonable and plausible features of a social system.
The slight twist is that we do not talk about potential sex differences in *interaction rates*.
Instead we talk about potential sex differences in *propensities* to interact.
Of course the same applies to relatedness and affinity: it's about propensities to interact, not actual interactions.
Figure \ref{fig:schemesociality} visualizes the idea.

```{r, echo=FALSE}
cap <- "A simple flow chart illustrating the idea how covariates might affect individual and dyadic propensities to interact."
```

```{r schemesociality, echo=FALSE, out.width="60%", fig.cap=cap}
knitr::include_graphics("_images/scheme_sociality.png")
```


We can also formalize this.
And that's the advantage of taking the generative approach.
In the framework of simple network metrics (I'm thinking of SRI or HWI or DSI) this is just not possible and we rely on ad-hoc solutions with potentially sever shortcomings (for some background see @hart2022): SRI, DSI and friends are non-parametric indices and therefore can't be incorporated in generative simulations.
We can do better, I like to think [@mcelreath2020].
So here is an attempt to a formal generative model:

\begin{equation}
\label{eq:sharks}
\begin{aligned}
\textrm{interactions} & = b_0 + \sqrt{0.5} \times (G_i + G_j) + A_{ij} \\
A_{ij} &= a_{ij} + b_r \times \textrm{relatedness}\\
G_{i} &= g_{i} + b_s \times \textrm{sex}\\
g &\sim \textrm{Normal}(0, \sigma^2_g) \\
r &\sim \textrm{Normal}(0, \sigma^2_r) \\
\sigma^2_g &\sim \textrm{Exponential}(1) \\
\sigma^2_r &\sim \textrm{Exponential}(1) \\
b_s &\sim \textrm{Normal(0, 0.5)} \\
b_r &\sim \textrm{Normal(0, 0.5)}
\end{aligned}
\end{equation}

In this equation we just add slope parameters ($b_r$ and $b_s$) for the effects of relatedness and sex. 
Note how $b_r$ only affects affinity and $b_s$ only affects gregariousness (figure \ref{fig:schemesociality}).
Everything else is the same as before.
In the future I might also translate this model into a DAG, which might also prove helpful, but not just now (and please don't consider figure \ref{fig:schemesociality} as a formal DAG).

Before we get to a worked example: this is still a fairly simple model, and as such it can be fitted with the `bamoso` package.
Naturally, we can envision models much more complex than this. 
At this point it becomes hard to provide a universal syntax to fit such models without actually doing some hand-coding in Stan.
But for now, let's stay with this scenario where we have a dyadic predictor and a individual predictor.

## An example with sharks

The example data set that we use concerns reef sharks (*Carcharhinus melanopterus*) and the data come from @mourier2021 [@mourier2020].
The only reason for using this data set is that it is the first that I found with the following features:

- raw interaction or association data

- a dyad-level predictor (here relatedness)

- an individual-level predictor (here sex) 

The purpose of this example is primarily to illustrate the feasibility of integrating covariates.
I won't discuss the biological relevance of any results.
So let's load the data:


```{r}
library(bamoso)
data("sharks")
sdat <- make_stan_data_from_association(asso_table = sharks$associations, 
                                        indi_cat_pred = sharks$is_female, 
                                        dyad_covariate_pred = sharks$relatedness)
```

And fit the model (which takes about two minutes on my laptop):

```{r sharkmodelfit, cache=TRUE, include=FALSE}
res <- sociality_model(sdat, parallel_chains = 4, seed = 2)
```

```{r, eval = FALSE}
res <- sociality_model(sdat, parallel_chains = 4, seed = 2)
```


Let's start by looking at some diagnostics plots (figure \ref{fig:ppsharks}).

```{r ppsharks, echo=2:55, fig.width=8, fig.height=4.7, out.width="80%", fig.align='center', fig.cap="Posterior predictive checks for the shark model."}
par(family = "serif", mgp = c(1.2, 0.2, 0), tcl = 0, mar = c(2.5, 2.5, 0.5, 0.5), mfrow = c(2, 2))

pp_model(res, xlab = "associated", ylab = "")
pp_model(res, selected_id = "Op18", xlab = "associated", ylab = "")
title(main = "ID: Op18", line = -1, cex.main = 1)
pp_model(res, selected_id = "She5", xlab = "associated", ylab = "")
title(main = "ID: She5", line = -1, cex.main = 1)
pp_model_stat(res, stat = "max", main = "", xlab = "max seen associated")
```


Now let's extract the posteriors of the $b_s$ and $b_r$ parameters.
This is done by accessing the results object's internal components but will in the future be possible via `extract_samples()`.

```{r}
br <- c(res$mod_res$draws("dyad_covariate_eff"))
bs <- c(res$mod_res$draws("indi_cat_eff"))
```

Now let's plot the posteriors:

```{r, echo=FALSE}
cap <- "Association between relatedness and propensities to interact with specific partners (dyadic affinity) and association between sex and propensity to interact with anyone (gregariousness). Negative values for the sex parameters indicate the females have lower values than males."
```

```{r sharkeffects, echo=3:55, fig.width=8, fig.height=2.4, out.width="80%", fig.align='center', fig.cap=cap}
par(family = "serif", mgp = c(1.2, 0.2, 0), tcl = -0.1, mar = c(2.5, 2.5, 1, 0.5), mfrow = c(1, 2), 
    cex.axis = 0.8, las = 1, yaxs = "i", bty = "l")

hist(br, main = "relatedness effect", xlab = "estimate", ylim = c(0, 1000))
abline(v = 0, col = "red")
hist(bs, main = "sex effect", xlab = "estimate", ylim = c(0, 1000))
abline(v = 0, col = "red")
```

It appears that most posterior mass for the relatedness falls in the positive range (posterior median: $`r sprintf("%.2f", median(br))`$, proportion of samples larger than 0: $`r sprintf("%.2f", mean(br > 0))`$, figure \ref{fig:sharkeffects}). 
It also appears that the sex effect is pretty reliably negative, i.e. females have lower gregariousness values than males (posterior median: $`r sprintf("%.2f", median(bs))`$, proportion of samples smaller than 0: $`r sprintf("%.2f", mean(bs < 0))`$).


\clearpage

```{r, echo=FALSE, eval = TRUE}
# this is forced to be evaluated!
stime <- "unknown"
if (identical(Sys.getenv("NOT_CRAN"), "true")) {
  stime <- round(as.numeric(difftime(time2 = compile_start, time1 = Sys.time(), units = "mins")), 1)
}
```

This document took `r stime` minutes to compile.

\clearpage

# References
````
