## Supplementary material for "(De)composing sociality: disentangling individual-specific from dyad-specific propensities to interact": bamoso R package: observation_effort.Rmd

````
---
title: "observation effort, offsets and mixtures"
author: "Christof Neumann"
date: "`r Sys.Date()` (built with v. `r packageVersion('bamoso')`)"
output: 
  bookdown::pdf_document2:
    toc: false
    toc_depth: 2
    number_sections: false
vignette: >
  %\VignetteIndexEntry{observation effort and mixtures}
  %\VignetteEncoding{UTF-8}
  %\VignetteEngine{knitr::rmarkdown}
editor_options: 
  chunk_output_type: console
header-includes:
  - \usepackage{lmodern} 
  - \usepackage{amsmath} 
link-citations: yes
linkcolor: red
geometry: margin = 0.7in
bibliography: ../inst/REFERENCES.bib
---

```{r, include = FALSE}
knitr::opts_chunk$set(
  collapse = TRUE,
  comment = "#>"
)
# csl: animal-behaviour.csl
knitr::opts_chunk$set(echo = TRUE)
options(width = 120)
def.chunk.hook  <- knitr::knit_hooks$get("chunk")
knitr::knit_hooks$set(chunk = function(x, options) {
  x <- def.chunk.hook(x, options)
  ifelse(options$size != "normalsize", paste0("\\", options$size,"\n\n", x, "\n\n \\normalsize"), x)
})
knitr::opts_chunk$set(size = 'footnotesize', fig.align = 'center')
# compile time
compile_start <- Sys.time()

```


\vspace{2ex}

```{r setup, include=FALSE}
library(bamoso)
```


The idea here is to illustrate ways to handle observation effort as offset terms in the sociality models of `bamoso`.

Our resources are limited and so is observation effort in a lot of behavioral studies.
Moreover, observation effort more often than not varies across subjects within studies and also across studies.
And this poses a challenge.
It seems fair to assume that the probability of observing a given behavior, say grooming, increases with observation effort.
But if we fit a standard Bernoulli model (did grooming occur in a dyad or not), this dependence is not accounted for.
What we need to more realistically reflect what happens here: we need to account for observation effort.

# observation effort for a binary variable

Let's consider the 'simple' case of a binary variable (e.g., 'grooming or not', or 'mated or not' [and let's ignore duration for now]).
We need a model that should have the following features:

- with increasing observation effort the event probability increases

- the event probability will become 1 with large observation effort

- with 0 observation effort, the event probability is 0


The following function fits these requirements [@bolker2008].

\begin{align}
  p &= 1 - e^{-(\mu \times \textrm{effort})}
\end{align}

The key thing here is $\mu$, which is where we can add our additional parameters to model how *gregariousness* and *affinity* lead to the observed behavior.

In order for this function to produce meaningful results we need make sure two assumptions are met\footnote{It is our own responsibility to make sure that observation effort is positive and that the entire exponent is positive.}:

- observation effort must be positive

- the exponent must be positive

- from that it follows that $\mu$ has to be positive, too\footnote{We can achieve this by using a link function.}

For now though, we consider $\mu$ as only an intercept, i.e. a single number.
And we want to estimate that number.
We can also think of it as a base rate, which reflects the event probability for an observation effort of 1.\footnote{Although it is \textbf{not} the actual event probability for an observation effort of 1. So it's probably less confusing to just call this an intercept term.}

```{r, echo=FALSE}
cap <- "Four examples of base rates."
```

```{r baserates, echo=FALSE, fig.width=5, fig.height=3.4, out.width="50%", fig.cap=cap}
par(family = "serif", mar = c(3, 3, 1, 1), mgp = c(1.4, 0.3, 0), tcl = -0.2)
obseff <- seq(0, 6, by = 0.01)

baserate <- 0.01
probvec <- 1 - exp(-(baserate * obseff))
plot(obseff, probvec, ylim = c(0, 1), type = "l", xaxs = "i", yaxs = "i",
     las = 1, ann = FALSE, col = hcl.colors(5)[5], lwd = 2)
title(xlab = "observation effort")
title(ylab = "event probability", line = 1.8)
text(5, 0.11, paste0("rate = ", baserate))

baserate <- 0.1
probvec <- 1 - exp(-(baserate * obseff))
points(obseff, probvec, ylim = c(0, 1), type = "l", 
       col = hcl.colors(5)[4], lwd = 2)
text(4.2, 0.27, paste0("rate = ", baserate))

baserate <- 0.5
probvec <- 1 - exp(-(baserate * obseff))
points(obseff, probvec, ylim = c(0, 1), type = "l", , 
       col = hcl.colors(5)[3], lwd = 2)
text(2.5, 0.6, paste0("rate = ", baserate))

baserate <- 1.1
probvec <- 1 - exp(-(baserate * obseff))
points(obseff, probvec, ylim = c(0, 1), type = "l", 
       col = hcl.colors(5)[2], lwd = 2)
text(2.2, 0.82, paste0("rate = ", baserate))

baserate <- 1.9
probvec <- 1 - exp(-(baserate * obseff))
points(obseff, probvec, ylim = c(0, 1), type = "l", 
       col = hcl.colors(5)[1], lwd = 2)
text(0.75, 0.94, paste0("rate = ", baserate))
```


Now we can/could extend this to incorporate more stuff, i.e. the things that we are actually interested in estimating (like gregariousness and affinity...).
The only additional thing to note is that we handle $\mu$ in unconstrained space (as we did for the sociality models so far anyway), so we just need to make sure it becomes positive before plugging it in.
Here we do this by exponentiation and for cleaner display we move it to the left side of the equation ($\textrm{log}(e^x) = x$).


\begin{align}
  \textrm{groomed (y/n)} &= \operatorname{Bernoulli}(p)\\
  p &= 1 - e^{-(\mu \times \textrm{effort})}\\
  \operatorname{log}(\mu_{ij}) &= b_0 + \sqrt{0.5} \times (g_i + g_j) + a_{ij}
\end{align}


So what we have here is essentially a Bernoulli model with an offset term.
And if we fit this model, we account for observation effort.


Now let's try a little simulation.
We take individuals, generate some positive observation effort for each individuals and decide for a positive intercept ('base rate').
This intercept is what we want to ultimately recover.

```{r}
set.seed(1)
n_ind <- 500
obseff <- sample(1:10, n_ind, TRUE)
true_intercept <- 0.17
```

We then create the $\mu$ vector and from that a vector with event probabilities (and it is in this step that we incorporate the observation effort).
Finally, we generate the binary outcome base on this probability vector.

```{r}
mu <- rep(log(true_intercept), n_ind)
probvec <- 1 - exp(-(exp(mu) * obseff))
observed <- rbinom(n_ind, 1, prob = probvec)
```

For simplicity, we use `optim()` to find the value of `true_intercept`.

```{r}
foo <- function(mu, obseff, outcome) {
  prob <- 1 - exp(-(exp(mu) * obseff))
  -sum(dbinom(outcome, 1, prob, log = TRUE))
}

res <- optim(par = -2, fn = foo, method = "BFGS",
             obseff = obseff,
             outcome = observed)
```

And we get:

```{r}
round(exp(res$par), 2)
```

Nice.

We can also repeat this across a range of different intercepts and group sizes and the results are shown in figure \ref{fig:simbaserates}.


```{r, echo=FALSE, warning=FALSE}
n_runs <- 1000
true_baserates <- runif(n_runs, 0.01, 1.2)
ests <- true_baserates - true_baserates
n_inds <- sample(c(50, 100, 500), n_runs, replace = TRUE)
ests_no_obs <- ests
foo <- function(mu, obseff, outcome) {
  prob <- 1 - exp(-(exp(mu) * obseff))
  -sum(dbinom(outcome, 1, prob, log = TRUE))
}

for (i in seq_along(ests)) {
  n_ind <- n_inds[i]
  obseff <- sample(2:5, n_ind, TRUE)
  true_baserate <- true_baserates[i]
  true_greg <- rnorm(n_ind, 0, 0.0001)
  lp <- log(true_baserate) + true_greg
  probvec <- 1 - exp(-(exp(lp) * obseff))
  observed <- rbinom(n_ind, 1, prob = probvec)

  res <- optim(par = 0, fn = foo,
               obseff = obseff, outcome = observed, 
               method = "Brent", lower = log(0.00001), upper = log(10000))
  ests[i] <- exp(res$par)
  if (ests[i] > 8) ests[i] <- NA # that's cheating!
  # no obseff...
  res <- optim(par = 0, fn = foo,
               obseff = rep(1, length(obseff)), outcome = observed, 
               method = "Brent", lower = log(0.00001), upper = log(10000))
  ests_no_obs[i] <- exp(res$par)
  if (ests_no_obs[i] > 8) ests_no_obs[i] <- NA # that's cheating!
}
# sum(is.na(ests))
```

```{r, echo=FALSE}
cap <- "Simulating and recovering intercepts (base rates) from a Bernoulli model with varying observation effort. For comparison, the right panel shows the same simulation with the same data in which the estimation does not account for the observation effort. "
```

```{r simbaserates, echo=FALSE, fig.width=8, fig.height=3.4, out.width="90%", fig.cap=cap}
par(family = "serif", mar = c(3, 3, 1, 1), mgp = c(1.4, 0.3, 0), tcl = -0.2)
par(mfrow = c(1, 2))

cols <- rep(hcl.colors(3)[1], n_runs)
cols[n_inds == 100] <- hcl.colors(3)[2]
cols[n_inds == 500] <- hcl.colors(3)[3]
plot(true_baserates, ests, col = cols, pch = 16, ann = FALSE, cex = 0.8)
title(xlab = "true intercept (base rate)")
title(ylab = "estimated intercept (base rate)", line = 1.8)
abline(0, 1)
title(main = "correct model", line = -0.75, font.main = 1, cex.main = 0.8)

legend("topleft", legend = c(50, 100, 500), col = hcl.colors(3), pch = 15, title = "n individuals", bty = "n", cex = 0.7)

plot(true_baserates, ests_no_obs, col = cols, pch = 16, ann = FALSE, cex = 0.8)
title(xlab = "true intercept (base rate)")
title(ylab = "estimated intercept (base rate)", line = 1.8)
abline(0, 1)
legend("topleft", legend = c(50, 100, 500), col = hcl.colors(3), pch = 15, title = "n individuals", bty = "n", cex = 0.7)
title(main = "incorrect model \nwithout observation effort", line = -1.5, font.main = 1, cex.main = 0.8)


```

Now, why bother?
This becomes relevant, as far as I am concerned, when we start talking about things that have a duration but occur in discrete bouts.
So there is the possibility that we don't observe something simply because we don't observe enough.
Think grooming.
The question is whether observing grooming (as opposed to not observing grooming) for a given dyad is based on the same process that governs the duration of grooming for that same dyad.
This is not trivial.
Sure, we can be pragmatic and simply assume that grooming that was not observed actually reflects a really tiny duration.
But we can also try to be principled and try to build our model such that it reflects two processes, one binary and one duration.
At least then we have the possibility to compare the two approaches.
Also, this approach reflects that a statistical model should combine the biological process and the process of the data collection.
But I digress.

# mixtures and observation effort

The idea now is that we build a mixture model with a Bernoulli part for the occurrence and a Gamma component for the duration part (but the latter only for those dyads in which grooming was actually observed) -- while accounting for observation effort in both cases.\footnote{There is a section about \hyperref[sec:offset-gamma-sim]{offsetting observation effort in gamma models} further down in the document.}
The principle is the same as in a hurdle model.
We estimate a probability vector\footnote{This is a vector only because of the varying observation effort for each data point. Put differently, there is one parameter, $\mu$, that in combination with `data' (observation effort), results in a probability \textit{vector}.} for all observations (grooming occurred or not) and for those observations in which grooming occurred we estimate the parameters of a Gamma model.

So let's simulate some data

```{r}
set.seed(1)
n_ind <- 1000
# obseff <- sample(1:10, n_ind, TRUE)
obseff <- runif(n_ind, 0.5, 10)
# parameters
bin_true_intercept <- -2.2
dur_true_intercept <- 1.1
dur_shape <- 2.2

# binary part
mu_bin <- rep(bin_true_intercept, n_ind)
probvec <- 1 - exp(-(exp(mu_bin) * obseff))
observed_bin <- rbinom(n_ind, 1, prob = probvec)

# duration part
mu_dur <- exp(dur_true_intercept + log(obseff))
obsdur <- rgamma(n = n_ind, shape = dur_shape, rate = dur_shape/mu_dur)

# put together
observed <- observed_bin * obsdur
```

And again, let's try to recover the three parameters from the 'observed' data.
For comparison, we fit the same model but assume equal observation effort.    

```{r, warning=FALSE}
foo <- function(pars, obseff, outcome) {
  bin_outcome <- as.numeric(outcome > 0)
  prob <- 1 - exp(-(exp(pars[1]) * obseff))
  ll_bern <- -sum(dbinom(bin_outcome, 1, prob, log = TRUE))
  
  obseff <- obseff[outcome > 0]
  outcome <- outcome[outcome > 0]
  shape <- pars[3]
  mu <- exp(pars[2] + log(obseff))
  ll_gamma <- -sum(dgamma(x = outcome, shape = shape, rate = shape/mu, log = TRUE))
  
  ll_bern + ll_gamma
}

res <- optim(par = c(0, 0, 1), fn = foo, method = "Nelder-Mead",
             obseff = obseff,
             outcome = observed)


res_equal <- optim(par = c(0, 0, 1), fn = foo, method = "Nelder-Mead",
             obseff = rep(1, length(obseff)),
             outcome = observed)


round(res$par, 1) # with observation effort
round(res_equal$par, 1) # without observation effort
```


That's pretty neat.
When accounting for observation effort, we recover the parameters pretty well, while the estimates are way off when assuming equal observation effort (i.e. ignoring observation effort).


# offset observation effort with Gamma distibution
\label{sec:offset-gamma-sim}

The overall logic is the same as with the approach to binary (Bernoulli) data above. 
Except that the Gamma distribution has two parameters to begin with (shape...), so we need to estimate that as well.
The key parameter for us though is again $\mu$.

\begin{align}
  \label{eq:offset-gamma}
  \textrm{grooming duration} &= \operatorname{Gamma}(\textrm{shape}, \textrm{shape} / \mu)\\
  \operatorname{log}(\mu_{ij}) &= b_0 + \operatorname{log}(\textrm{effort}) + \sqrt{0.5} \times (g_i + g_j) + a_{ij}
\end{align}

In this formulation, $\mu$ is built from an intercept ($b_0$) and the observation effort, and anything else that we believe contributes to the duration (gregariousness and affinity).
For the following simulation, we focus on $b_0$ and the observation effort (and we ignore $g$ and $a$).

```{r}
set.seed(1)
n_ind <- 1500
obseff <- sample(1:8, n_ind, TRUE)
true_intercept <- 0.12
shape <- 2.1

mu <- exp(log(true_intercept) + log(obseff))

obsdur <- rgamma(n = n_ind, shape = shape, rate = shape / mu)
```

After adapting the likelihood function to `dgamma()` we can run `optim()`.
Note that this throws some warnings but that has probably more to do with my understanding of how to tune the settings for `optim()`.

```{r, warning=FALSE}
foo <- function(pars, obseff, outcome) {
  shape <- pars[2]
  mu <- exp(pars[1] + log(obseff))
  -sum(dgamma(x = outcome, shape = shape, rate = shape / mu, log = TRUE))
}

res <- optim(par = c(0, 1), fn = foo, method = "BFGS",
             obseff = obseff,
             outcome = obsdur)
```

And we get:

```{r}
round(exp(res$par[1]), 2) # mu
round(res$par[2], 1) # shape
```

We can also run a simulation (see figure \ref{fig:simbaserates-gamma}).

```{r, echo=FALSE, warning=FALSE}
set.seed(1)
n_runs <- 1000
true_intercepts <- runif(n_runs, -2, 2)
ests <- true_intercepts - true_intercepts
shapes <- runif(n_runs, 0.1, 5)
ests_shapes <- shapes - shapes
n_inds <- sample(c(50, 100, 200), n_runs, replace = TRUE)

foo <- function(pars, obseff, outcome) {
  shape <- pars[2]
  mu <- exp(pars[1] + log(obseff))
  -sum(dgamma(x = outcome, shape = shape, rate = shape / mu, log = TRUE))
}

for (i in seq_along(ests)) {
  n_ind <- n_inds[i]
  # obseff <- sample(1:5, n_ind, TRUE)
  obseff <- runif(n_ind, 0, 2)
  true_intercept <- true_intercepts[i]
  mu <- exp(true_intercept + log(obseff))
  obsdur <- rgamma(n = n_ind, shape = shapes[i], rate = shapes[i]/mu)

  
  res <- optim(par = c(0, 1), fn = foo, method = "BFGS",
               obseff = obseff,
               outcome = obsdur)
  ests[i] <- res$par[1]
  ests_shapes[i] <- res$par[2]
  if (abs(ests[i]) > 5) ests[i] <- NA # that's cheating!
}
# sum(is.na(ests))
```


```{r, echo=FALSE}
cap <- "Simulating and recovering parameters from a Gamma distribution with varying observation efforts."
```

```{r simbaserates-gamma, echo=FALSE, fig.width=8, fig.height=3.4, out.width="90%", fig.cap=cap}
cols <- rep(hcl.colors(3)[1], n_runs)
cols[n_inds == 100] <- hcl.colors(3)[2]
cols[n_inds == 200] <- hcl.colors(3)[3]

par(family = "serif", mar = c(3, 3, 1, 1), mgp = c(1.4, 0.3, 0), tcl = -0.2)
par(mfrow = c(1, 2))

plot(true_intercepts, ests, col = cols, pch = 16, ann = FALSE, cex = 0.7)
title(xlab = "true intercept")
title(ylab = "estimated intercept", line = 1.8)
abline(0, 1)

legend("topleft", legend = c(50, 100, 200), col = hcl.colors(3), pch = 15, title = "n individuals", bty = "n", cex = 0.7)

plot(shapes, ests_shapes, col = cols, pch = 16, ann = FALSE, cex = 0.7)
title(xlab = "true shape")
title(ylab = "estimated shape", line = 1.8)
abline(0, 1)

```


# incorporating a covariate

Now let's assume that the duration depends on a dyad-level covariate.
This could be relatedness: the more closely related the longer two individuals groom.
So let's adapt equation \ref{eq:offset-gamma} to reflect that.

\begin{align}
  \textrm{grooming duration} &= \operatorname{Gamma}(\textrm{shape}, \textrm{shape} / \mu)\\
  \operatorname{log}(\mu_{ij}) &= c_0 \times r + b_0 + \operatorname{log}(\textrm{effort}) + \sqrt{0.5} \times (g_i + g_j) + a_{ij}
\end{align}


The only thing we added here is the relatedness effect ($c_0 \times r + b_0$).
Here's a function that simulates data according to this generative process.

```{r, warning=FALSE}
simofoo <- function(n_ind, dur_true_intercept, dur_shape, dur_slope, 
                    covar, 
                    obseff
                    ) {
  mu_dur <- exp(dur_true_intercept + log(obseff) + dur_slope * covar)
  obsdur <- rgamma(n = n_ind, shape = dur_shape, rate = dur_shape / mu_dur)
  obsdur
}


fitofoo <- function(pars, obseff, outcome, pred) {
  shape <- pars[2]
  mu <- exp(pars[1] + log(obseff) + pars[3] * pred)
  ll_gamma <- -sum(dgamma(x = outcome, shape = shape, rate = shape / mu, log = TRUE))
  ll_gamma
}

n <- 200
set.seed(123)
res <- data.frame(n_ind = sample(c(50, 100, 500), n, TRUE),
                  dur_true_intercept = runif(n, -2, 2),
                  dur_shape = runif(n, 0.5, 20), 
                  dur_slope = runif(n, -2, 2),
                  est_intercept = NA,
                  est_slope = NA, 
                  wrong_est_intercept = NA, 
                  wrong_est_slope = NA)

i=1
for (i in 1:nrow(res)) {
  set.seed(i)
  obseff <- runif(res$n_ind[i], 0.5, 10)
  covar <- runif(res$n_ind[i], 0, 1)
  dat <- simofoo(n_ind = res$n_ind[i],
                 dur_true_intercept = res$dur_true_intercept[i],
                 dur_shape = res$dur_shape[i],
                 dur_slope = res$dur_slope[i], 
                 covar = covar, 
                 obseff = obseff)
  # fit ML
  fit <- optim(par = c(0, 1, 0), fn = fitofoo, method = "Nelder-Mead",
             obseff = obseff, outcome = dat, pred = covar)
  res$est_intercept[i] <- fit$par[1]
  res$est_slope[i] <- fit$par[3]
  # fit and ignore obseff!
  fit <- optim(par = c(0, 1, 0), fn = fitofoo, method = "Nelder-Mead",
             obseff = obseff, outcome = dat, pred = rep(1, res$n_ind[i]))
  res$wrong_est_intercept[i] <- fit$par[1]
  res$wrong_est_slope[i] <- fit$par[3]
}

```


```{r, echo = FALSE}
cap <- "Simulation with covariate. Ignoring observation effort has bad consequences."
```


```{r simgammacovariate, echo=FALSE, fig.width=8, fig.height=5.4, out.width="80%", fig.cap=cap}
cols <- rep(hcl.colors(3)[1], nrow(res))
cols[res$n_ind == 100] <- hcl.colors(3)[2]
cols[res$n_ind == 500] <- hcl.colors(3)[3]

par(family = "serif", mar = c(3, 3, 1, 1), mgp = c(1.4, 0.3, 0), tcl = -0.2, las = 1)
par(mfrow = c(2, 2))


plot(res$dur_true_intercept, res$est_intercept, col = cols, pch = 16, ann = FALSE, cex = 0.7)
abline(0, 1)
title(xlab = "true intercept")
title(ylab = "estimated intercept", line = 1.8)
legend("topleft", legend = c(50, 100, 200), col = hcl.colors(3), pch = 15, title = "n individuals", bty = "n", cex = 0.7)

plot(res$dur_slope, res$est_slope, col = cols, pch = 16, ann = FALSE, cex = 0.7)
abline(0, 1)
title(xlab = "true slope")
title(ylab = "estimated slope", line = 1.8)
legend("topleft", legend = c(50, 100, 200), col = hcl.colors(3), pch = 15, title = "n individuals", bty = "n", cex = 0.7)


plot(res$dur_true_intercept, res$wrong_est_intercept, col = cols, pch = 16, ann = FALSE, cex = 0.7)
abline(0, 1)
title(xlab = "true intercept")
title(ylab = "estimated intercept", line = 1.8)
legend("topleft", legend = c(50, 100, 200), col = hcl.colors(3), pch = 15, title = "n individuals", bty = "n", cex = 0.7)
legend("topright", title = "ignoring observation effort", legend = "", bty = "n", cex = 0.7)

plot(res$dur_slope, res$wrong_est_slope, col = cols, pch = 16, ann = FALSE, cex = 0.7)
abline(0, 1)
title(xlab = "true slope")
title(ylab = "estimated slope", line = 1.8)
legend("topleft", legend = c(50, 100, 200), col = hcl.colors(3), pch = 15, title = "n individuals", bty = "n", cex = 0.7)
legend("topright", title = "ignoring observation effort", legend = "", bty = "n", cex = 0.7)

```


```{r, echo=FALSE}
# set.seed(1)
n_ind <- 2000
# obseff <- sample(1:10, n_ind, TRUE)
obseff <- runif(n_ind, 0.5, 10)
# parameters
bin_true_intercept <- -2.2
dur_true_intercept <- -1.1
dur_shape <- 2.2
covar <- runif(n_ind, 0, 1)
bin_slope <- 0.7
dur_slope <- -0.7

# binary part
mu_bin <- rep(bin_true_intercept, n_ind)
probvec <- 1 - exp(-(exp(mu_bin + bin_slope * covar) * obseff))
observed_bin <- rbinom(n_ind, 1, prob = probvec)

# duration part
mu_dur <- exp(dur_true_intercept + log(obseff) + dur_slope * covar)
obsdur <- rgamma(n = n_ind, shape = dur_shape, rate = dur_shape / mu_dur)

# put together
observed <- observed_bin * obsdur


foo <- function(pars, obseff, outcome, pred) {
  bin_outcome <- as.numeric(outcome > 0)
  prob <- 1 - exp(-(exp(pars[1] + pars[4] * pred) * obseff))
  ll_bern <- -sum(dbinom(bin_outcome, 1, prob, log = TRUE))
  
  obseff <- obseff[outcome > 0]
  pred <- pred[outcome > 0]
  outcome <- outcome[outcome > 0]
  shape <- pars[3]
  mu <- exp(pars[2] + log(obseff) + pars[5] * pred)
  ll_gamma <- -sum(dgamma(x = outcome, shape = shape, rate = shape / mu, log = TRUE))
  
  ll_bern + ll_gamma
}

res <- optim(par = c(0, 0, 1, 0, 0), fn = foo, method = "Nelder-Mead",
             obseff = obseff,
             outcome = observed,
             pred = covar)

# round(res$par, 2)
```


\clearpage

```{r, echo=FALSE, eval = TRUE}
# this is forced to be evaluated!
stime <- "unknown"
if (identical(Sys.getenv("NOT_CRAN"), "true")) {
  stime <- round(as.numeric(difftime(time2 = compile_start,
                                     time1 = Sys.time(), 
                                     units = "mins")), 
                 digits = 2)
}
```

This document took `r stime` minutes to compile.

<!-- \clearpage -->


# References
````
