## Supplementary figures and images for "(De)composing sociality: disentangling individual-specific from dyad-specific propensities to interact"

### README-eq.png

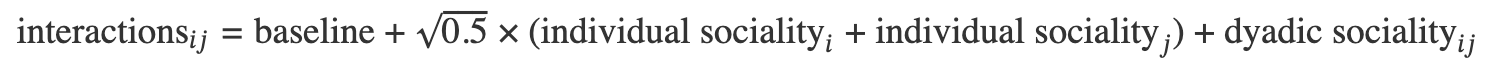

### README-groomdatasetplot-1.png

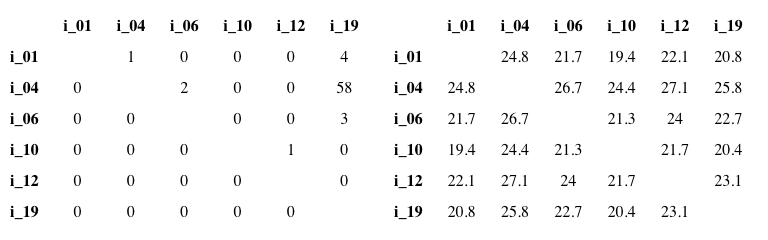

### README-ppcheck-1.png

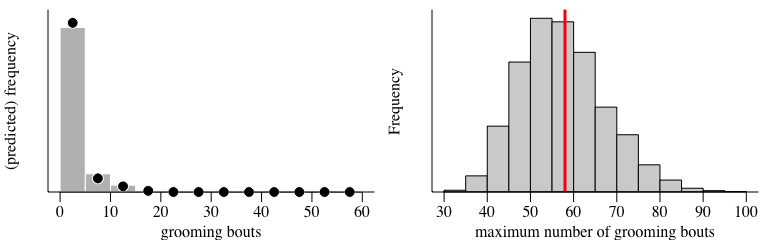

### README-ridgeplot-1.png

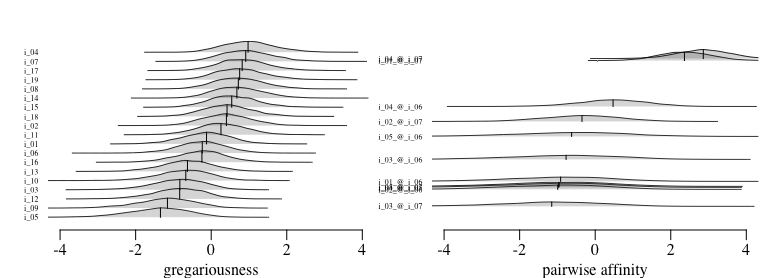

### README-socplot-1.png

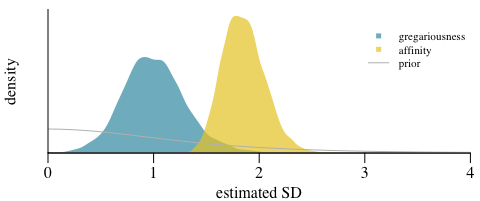

### RTG-LogoCYMK300.png

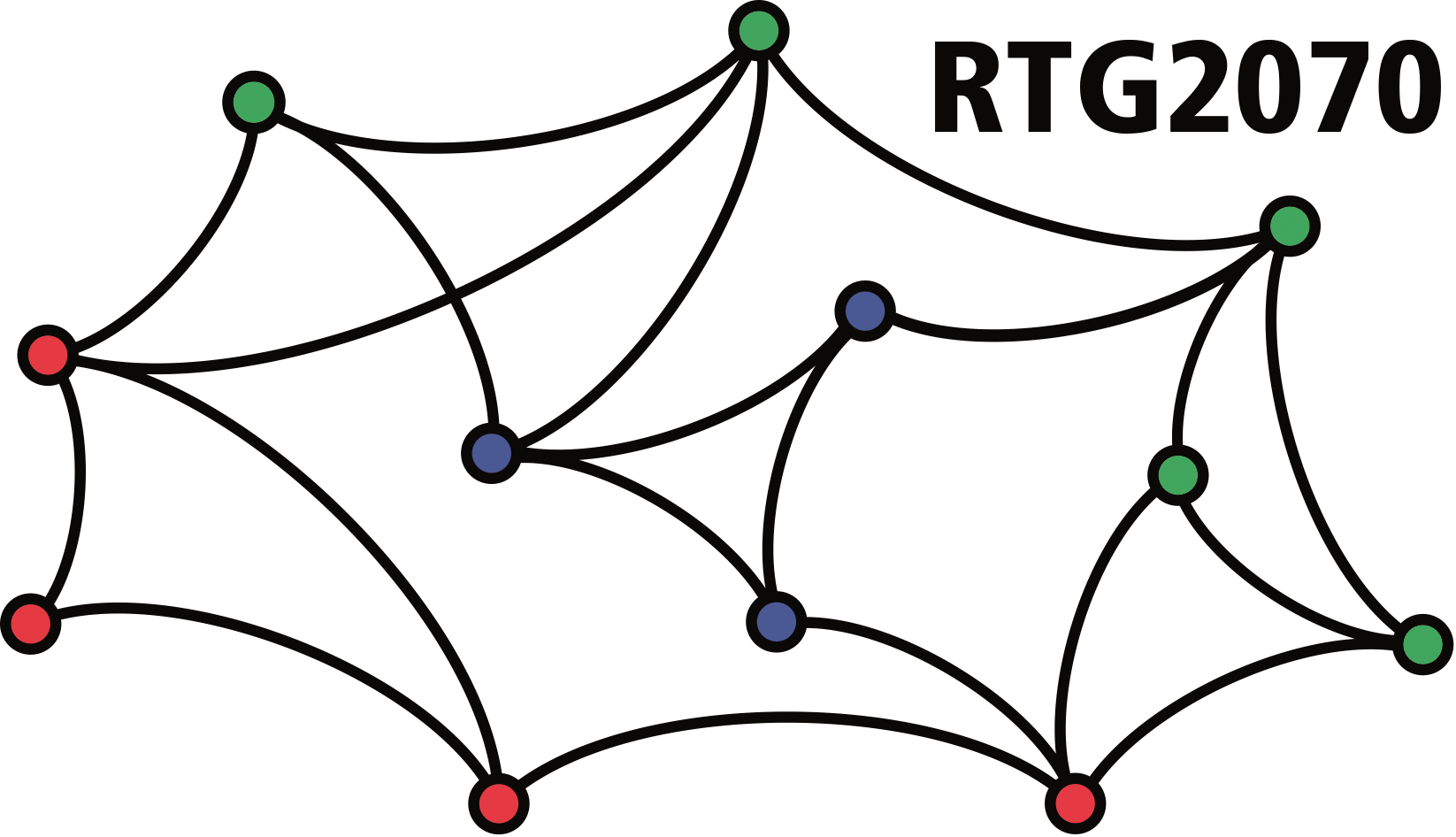

### scheme_sociality.png

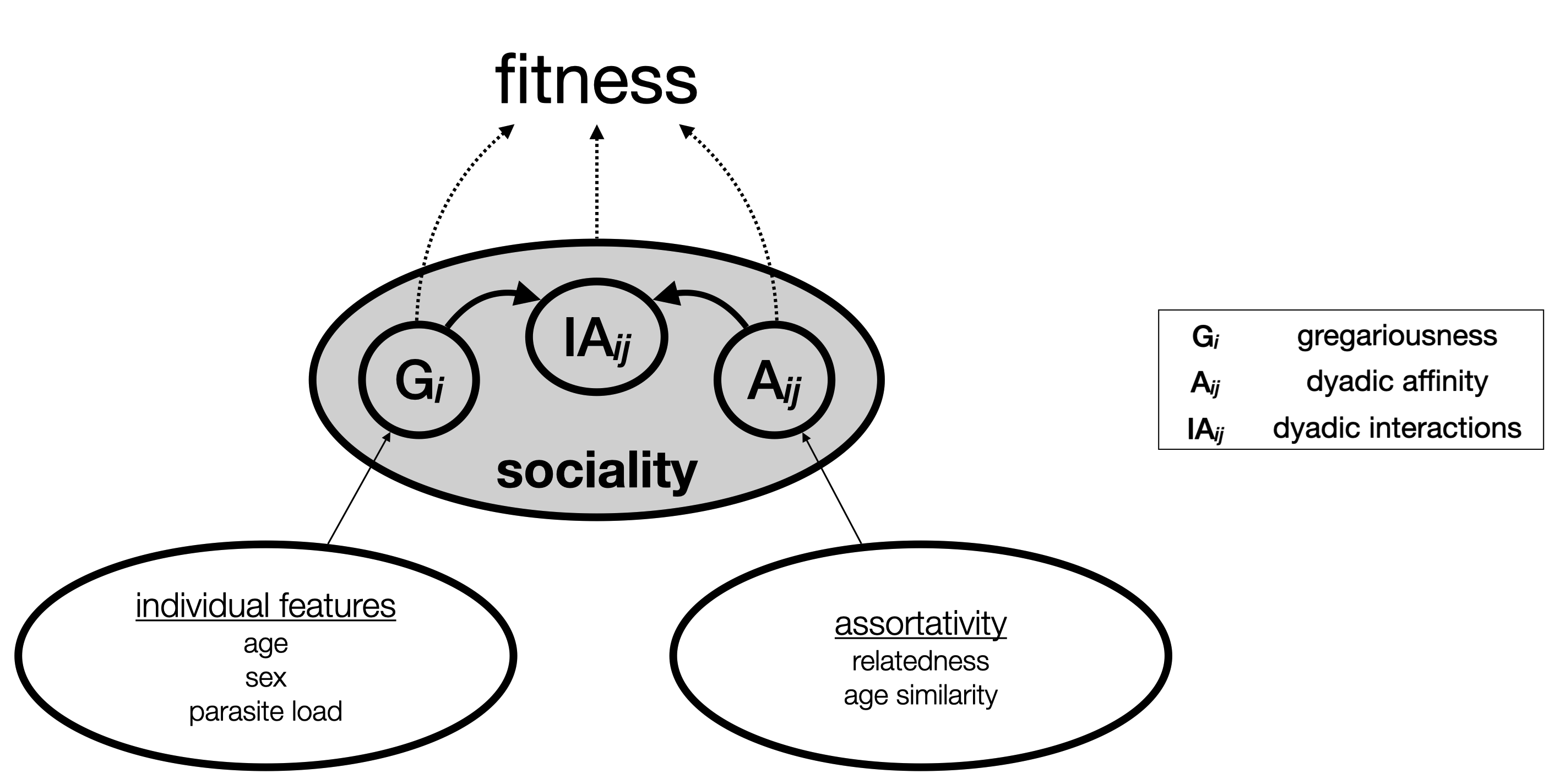
