## Supplementary material for "(De)composing sociality: disentangling individual-specific from dyad-specific propensities to interact": readme

Successful installation of the `bamoso` package is a requirement for running the code in the replication scripts (`replication_code_files.zip`).

# Requirements

* `R` (v. > 4.1)

* `cmdstanr` (see instructions below)

* a number of other R packages (`brms`, `posterior`, `knitr`, `igraph`, amongst others): these will be installed automatically when installing the `bamoso` package (see below).

# Content of replication scripts

* `repl_01_sim_example.R`: simulated example (figure 3 in main text)

* `repl_02_workflow_example.R`: workflow illustration (figure 4 in main text)

* `repl_03_macaques.R`: empirical example 1 (figure 5a-d in main text)

* `repl_04_kangaroos.R`: empirical example 2 (figure 5e-h in main text)

# Installation

You need [`cmdstanr`](https://mc-stan.org/cmdstanr/) in order to install and run `bamoso`. 
This in turn requires a working C++ toolchain first. 
Check out the [getting started-guide](https://mc-stan.org/cmdstanr/articles/cmdstanr.html) from `cmdstanr` to see how to get everything set up properly. 
Also [this document](https://mc-stan.org/docs/cmdstan-guide/cmdstan-installation.html#cpp-toolchain) might be helpful.

Then install `cmdstanr`.

```
install.packages("cmdstanr", repos = c("https://mc-stan.org/r-packages/", getOption("repos")))
```

And you also need the `remotes` packages, which is easy to install from CRAN:

```
install.packages("remotes")
```

Then check whether things are set up correctly:

```
library(cmdstanr)
check_cmdstan_toolchain(fix = TRUE)
```

You may need to follow any instructions that the output of that call gives.
If you haven't used `cmdstanr` before, it's likely that you at least need to install `cmdstan` itself:

```
cmdstanr::install_cmdstan()
```

If this gives positive feedback, install `bamoso`:

```
library(remotes)
remotes::install_local("bamoso_0.5.1.tar.gz", dependencies = TRUE, build_vignettes = FALSE)
```

If you want to install (recompile) the intro vignette, use:

```
install.packages("tinytex")
library(remotes)
remotes::install_local("bamoso_0.5.1.tar.gz", dependencies = TRUE, build_vignettes = TRUE)
```

This will take a little more time to finish, but if successful gives you access to the vignette.

Note that the intro vignette is also included in the supplements as a standalone file (`tutorial_bamoso.pdf`).

To access the vignettes from R (if you used `build_vignettes = TRUE` before):

```
vignette("intro", package = "bamoso")
vignette("covariates", package = "bamoso")
```


## Installation from GitHub

```
library(remotes)
remotes::install_github("gobbios/bamoso", dependencies = TRUE, build_vignettes = FALSE)
```

If you want to install (recompile) the intro vignette, use:

```
install.packages("tinytex")
library(remotes)
remotes::install_github("gobbios/bamoso", dependencies = TRUE, build_vignettes = TRUE)
```

This will take a little more time to finish, but if successful gives you access to the vignette.
