## Supplementary material for "(De)composing sociality: disentangling individual-specific from dyad-specific propensities to interact": replication R code files: repl_01_sim_example.R

```
# simulated example

# generate data ------------------
library(bamoso)

# new example for revision
set.seed(123)
d1 <- generate_data(n_ids = 15, n_beh = 2, behav_types = c("count", "dur_beta"),
                    indi_sd = 1.7, dyad_sd = 0.3, beh_intercepts = c(-1.7, -1.7))

set.seed(5333997)
d2 <- generate_data(n_ids = 15, n_beh = 2, behav_types = c("count", "dur_beta"),
                    indi_sd = 0.3, dyad_sd = 1.7, beh_intercepts = c(-1.7, -1.7))


# fit models -------------------
r1 <- sociality_model(d1$standat, silent = TRUE, parallel_chains = 4, seed = 1)
summary(r1)

r2 <- sociality_model(d2$standat, silent = TRUE, parallel_chains = 4, seed = 42,
                      adapt_delta = 0.99)
summary(r2)


# figure 3 ------------------------

par(mfrow = c(5, 2))
set.seed(123)
par(family = "serif", mgp = c(1.5, 0.5, 0), las = 1, mar = c(3, 3, 1, 1), tcl = 0)

## pp checks for behavior 1 (count (poisson))
pp_model(mod_res = r1, xvar = 1, xlim = c(0, 20), xbreaks = seq(-0.5, 20.5, by = 1),
         xlab = "interactions", ylab = "frequency")
legend("topright", legend = c("observed data", "posterior samples"),
       pch = c(15, 16), col = c("grey", "black"), lty = c(NA, 1),
       bty = "n", cex = 0.7)

pp_model(mod_res = r2, xvar = 1, xlim = c(0, 20), xbreaks = seq(-0.5, 50.5, by = 1),
         xlab = "interactions", ylab = "frequency")
legend("topright", legend = c("observed data", "posterior samples"),
       pch = c(15, 16), col = c("grey", "black"), lty = c(NA, 1),
       bty = "n", cex = 0.7)

## pp checks for behavior 2 (continuous proportion (beta))
pp_model_dens(mod_res = r1, xvar = 2, xlim = c(0, 1),
              xlab = "interaction duration", ylab = "density")
legend("topright", legend = c("observed data", "posterior samples"),
       pch = c(NA, NA), col = c("red", "black"), lty = c(1, 1),
       bty = "n", cex = 0.7)

pp_model_dens(mod_res = r2, xvar = 2, xlim = c(0, 1),
              xlab = "interaction duration", ylab = "density")
legend("topright", legend = c("observed data", "posterior samples"),
       pch = c(NA, NA), col = c("red", "black"), lty = c(1, 1),
       bty = "n", cex = 0.7)

# posteriors for gregariousness and affinity
sociality_plot(r1, xlim = c(0, 3), add_prior = TRUE)
abline(v = d1$input_data$indi_sd, col = "#3B99B1", lwd = 2)
abline(v = d1$input_data$dyad_sd, col = "#EACB2B", lwd = 2)

sociality_plot(r2, xlim = c(0, 3), add_prior = TRUE)
abline(v = d2$input_data$indi_sd, col = "#3B99B1", lwd = 2)
abline(v = d2$input_data$dyad_sd, col = "#EACB2B", lwd = 2)


## correlations between true individual gregariousness values and estimated values
p <- r1$mod_res$summary(variables = "indi_soc_vals")
plot(d1$input_data$indi_data$indi_soc_vals, p$median, xlim = c(-6, 6), ylim = c(-6, 6), bty = "l",
     lwd = 0.5, cex = 1.3, xlab = "true gregariousness", ylab = "estm'd gregariousness")
abline(0, 1)

xx <- paste0(
  paste("true:", sprintf("%0.2f", d1$input_data$indi_sd)), "\n",
  paste("estimated:", sprintf("%0.2f", r1$mod_res$summary(variables = "indi_soc_sd")["median"]))
)
text(-6, 6, labels = xx, adj = c(0, 1), cex = 0.8)


p <- r2$mod_res$summary(variables = "indi_soc_vals")
plot(d2$input_data$indi_data$indi_soc_vals, p$median, xlim = c(-6, 6), ylim = c(-6, 6), bty = "l",
     lwd = 0.5, cex = 1.3, xlab = "true gregariousness", ylab = "estm'd gregariousness")
abline(0, 1)
xx <- paste0(
  paste("true:", sprintf("%0.2f", d2$input_data$indi_sd)), "\n",
  paste("estimated:", sprintf("%0.2f", r2$mod_res$summary(variables = "indi_soc_sd")["median"]))
)
text(-6, 6, labels = xx, adj = c(0, 1), cex = 0.8)


## correlations between true dyadic affinity values and estimated values
p <- r1$mod_res$summary(variables = "dyad_soc_vals")
plot(d1$input_data$dyad_data$dyad_soc_vals, p$median, xlim = c(-6, 6), ylim = c(-6, 6),
     lwd = 0.5, cex = 1.3, xlab = "true affinity", ylab = "estimated affinity", bty = "l")
abline(0, 1)
xx <- paste0(
  paste("true:", sprintf("%0.2f", d1$input_data$dyad_sd)), "\n",
  paste("estimated:", sprintf("%0.2f", r1$mod_res$summary(variables = "dyad_soc_sd")["median"]))
)
text(-6, 6, labels = xx, adj = c(0, 1), cex = 0.8)

p <- r2$mod_res$summary(variables = "dyad_soc_vals")
plot(d2$input_data$dyad_data$dyad_soc_vals, p$median, xlim = c(-6, 6), ylim = c(-6, 6),
     lwd = 0.5, cex = 1.3, xlab = "true affinity", ylab = "estimated affinity", bty = "l")
abline(0, 1)
xx <- paste0(
  paste("true:", sprintf("%0.2f", d2$input_data$dyad_sd)), "\n",
  paste("estimated:", sprintf("%0.2f", r2$mod_res$summary(variables = "dyad_soc_sd")["median"]))
)
text(-6, 6, labels = xx, adj = c(0, 1), cex = 0.8)
```
