## Supplementary material for "(De)composing sociality: disentangling individual-specific from dyad-specific propensities to interact": replication R code files: repl_02_workflow_example.R

```
# workflow example

# preparatory steps
library(bamoso)
data("workflow_example", package = "bamoso")
standat <- workflow_example$standat
standat$y <- workflow_example$response
cstanmod <- cmdstan_model(system.file("extdata/workflow_example.stan", package = "bamoso"), compile = TRUE)

# fit model
toy_mod_model <- cstanmod$sample(data = standat, seed = 123, parallel_chains = 4, refresh = 0, init = 0)


# figure 4 ----

indi_soc <- workflow_example$input_data$indi_soc_vals

par(family = "serif", mgp = c(1.5, 0.5, 0), las = 1, mar = c(3, 3, 1, 1), tcl = 0)
par(mfrow = c(2, 2))

## true relationship (panel a)
plot(indi_soc, workflow_example$response, xlab = "true gregariousness",
     ylab = "outcome", ylim = c(0, 10), xlim = c(-2, 2))

s <- seq(min(indi_soc), max(indi_soc), length.out = 101)
points(s, exp(-0.5 + 1.2 * s), type = "l") # plot the TRUE model (not the estimated one)


## model estimates (panel b)
set.seed(1)
sel <- sample(4000, 50) # 50 draws

ivals <- toy_mod_model$draws("indi_soc_vals", format = "draws_matrix")[sel, ]
slopes <- toy_mod_model$draws("slope", format = "draws_matrix")[sel, ]
intercepts <- toy_mod_model$draws("intercept", format = "draws_matrix")[sel, ]

p1 <- apply(ivals, 2, quantile, 0.25)
p2 <- apply(ivals, 2, quantile, 0.75)
p_est <- apply(ivals, 2, quantile, 0.5)
yvals <- workflow_example$response + runif(workflow_example$standat$n_ids, -0.2, 0.2)
xlim <- c(-2, 2) # range(p1, p2)
plot(0, 0, xlim = xlim, ylim = c(0, 10), type = "n",
     xlab = "estimated gregariousness", ylab = "outcome", las = 1)
# uncertainty in gregariousness
segments(p1, yvals, p2, yvals, lwd = 0.5)

# predictions (50 draws)
xseq <- seq(from = min(p_est), to = max(p_est), length.out = 101)
xx <- sapply(seq_along(sel), function(X)points(xseq, exp(intercepts[X] + slopes[X] * xseq),
                                               type = "l", lwd = 0.2, col = "grey"))

points(p_est, yvals, pch = 21, bg = hcl.colors(5, "zissou")[3], col = "black", cex = 1.5)
p <- data.frame(toy_mod_model$summary(variables = c("intercept", "slope")))
rownames(p) <- p$variable
points(xseq, exp(p["intercept", "median"] + p["slope", "median"] * xseq),
       type = "l", lwd = 3, col = "#F5191C")


# reviewer suggests: add model without error propagation
# tempres <- glm(workflow_example$response ~ p_est, family = "poisson")
# points(xseq, exp(coef(tempres)[1] + xseq * coef(tempres)[2]), type = "l", lwd = 1, lty = 3, col = "#F5191C")

# pp checks model response (panel c)
x <- hist(workflow_example$response, plot = FALSE, breaks = seq(-0.5, 9.5, 1))
p <- toy_mod_model$draws("y_rep", format = "draws_matrix")

pd <- as.matrix(t(apply(p, 1, function(X) table(cut(X, breaks = x$breaks, include.lowest = TRUE)))))
pd <- apply(pd[sel, ], 2, quantile, probs = c(0.055, 0.5, 0.945))
sel <- which(colSums(rbind(x$counts, pd)) > 0)
maxval <- max(x$counts, pd)
xlim <- range(x$breaks)
plot(0, 0, type = "n", xlim = xlim, ylim = c(0, maxval * 1.15), yaxs = "i",
     axes = FALSE, las = 1, bty = "n", xlab = "outcome", ylab = "frequency", tcl = 0)
axis(1, at = 0:9, tcl = 0)
for (i in sel) {
  rect(xleft = x$breaks[i], ybottom = 0, xright = x$breaks[i + 1], ytop = x$counts[i],
       col = "grey", border = "white")
  points(x$mids[i], pd[2, i], pch = 21, cex = 1.5, xpd = TRUE,
         col = "white", bg = "black")
  segments(x0 = x$mids[i], y0 = pd[1, i], x1 = x$mids[i], y1 = pd[3, i])
}
box(bty = "l")


# model predictions: interactions (panel d)
x <- hist(workflow_example$standat$interactions[, 1], plot = FALSE, breaks = seq(-0.5, 139.5, 2))
p <- toy_mod_model$draws("interactions_pred", format = "draws_matrix")
pd <- as.matrix(t(apply(p, 1, function(X) table(cut(X, breaks = x$breaks, include.lowest = TRUE)))))
pd <- apply(pd[sel, ], 2, quantile, probs = c(0.055, 0.5, 0.945))
sel <- which(colSums(rbind(x$counts, pd)) > 0)
maxval <- max(x$counts, pd)
xlim <- range(x$breaks)
plot(0, 0, type = "n", xlim = xlim, ylim = c(0, maxval * 1.15), yaxs = "i",
     axes = FALSE, las = 1, bty = "n", xlab = "number of interactions", ylab = "frequency", tcl = 0)
axis(1, tcl = 0)
for (i in sel) {
  rect(xleft = x$breaks[i], ybottom = 0, xright = x$breaks[i + 1], ytop = x$counts[i],
       col = "grey", border = "white")
  points(x$mids[i], pd[2, i], pch = 21, cex = 1, xpd = TRUE,
         col = "white", bg = "black")
  segments(x0 = x$mids[i], y0 = pd[1, i], x1 = x$mids[i], y1 = pd[3, i])
}
box(bty = "l")
```
