## Supplementary material for "(De)composing sociality: disentangling individual-specific from dyad-specific propensities to interact": replication R code files: repl_03_macaques.R

```
# replicate macaque example

library(bamoso)
data(grooming)

# assamensis
x <- grooming$ass
sdat_ass <- make_stan_data_from_matrices(mats = list(groom = x$groom),
                                         behav_types = "count",
                                         obseff = list(x$obseff))
res_ass <- sociality_model(standat = sdat_ass, seed = 12754, parallel_chains = 4,
                           silent = TRUE, adapt_delta = 0.99, max_treedepth = 12)
# fuscata
x <- grooming$fus
sdat_fus <- make_stan_data_from_matrices(mats = list(groom = x$groom),
                                         behav_types = "count",
                                         obseff = list(x$obseff))
res_fus <- sociality_model(standat = sdat_fus, seed = 85679, parallel_chains = 4,
                           silent = TRUE, adapt_delta = 0.99, max_treedepth = 12)
# nigra
x <- grooming$nig
sdat_nig <- make_stan_data_from_matrices(mats = list(groom = x$groom),
                                         behav_types = "dur_beta",
                                         obseff = list(x$obseff))
res_nig <- sociality_model(standat = sdat_nig, seed = 765149, parallel_chains = 4,
                           silent = TRUE, adapt_delta = 0.99, max_treedepth = 12)
# sylvanus
x <- grooming$syl
sdat_syl <- make_stan_data_from_matrices(mats = list(groom = x$groom),
                                         behav_types = "count",
                                         obseff = list(x$obseff))
res_syl <- sociality_model(standat = sdat_syl, seed = 27374, parallel_chains = 4,
                           silent = TRUE, adapt_delta = 0.99, max_treedepth = 12)


# figure 5a-d --------------------
par(mfrow = c(2, 2), family = "serif", mgp = c(1.5, 0.5, 0))
sociality_plot(res_ass, add_prior = TRUE)
title(main = "assamensis", font.main = 4)
sociality_plot(res_fus, add_prior = TRUE)
title(main = "fuscata", font.main = 4)
sociality_plot(res_nig, add_prior = TRUE)
title(main = "nigra", font.main = 4)
sociality_plot(res_syl, add_prior = TRUE)
title(main = "sylvanus", font.main = 4)
```
