## Supplementary material for "(De)composing sociality: disentangling individual-specific from dyad-specific propensities to interact": replication R code files: repl_04_kangaroos.R

```
# The two Stan functions written to accomplish the calculation of strength
#   and individual cluster coefficients are written in a fairly inefficient
#   style, and hence slow.

# It is likely that they could be written much more efficiently.

# Model fitting with the subset of 47 individuals takes around 20 minutes on a
#   fairly current laptop computer, while for the complete data set, it is
#   likely model fitting would take several hours.


library(bamoso)

data("kangaroos_subset")
standat <- kangaroos_subset

# get and compile model
mod <- cmdstan_model(stan_file = system.file("extdata/bayesian_kangaroos.stan",
                                             package = "bamoso"))

# fit model ----------------------

# model fit for MS:
# r <- mod$sample(data = standat, parallel_chains = 6, chains = 6,
#                 show_exceptions = FALSE, iter_warmup = 1000, iter_sampling = 500,
#                 seed = 1, refresh = 10)

# model fit for replication (less iterations) (about 10 minutes on my computer)
r <- mod$sample(data = standat, parallel_chains = 4, chains = 4, show_exceptions = FALSE,
                iter_warmup = 200, iter_sampling = 150, seed = 1, refresh = 10)

# store model results if needed
# r$save_object(file = "results/kangaroo_wean_model.rds")

# or reload from stored file
# r <- readRDS(file = "results/kangaroo_wean_model.rds")


# figure 5e-h ----
par(family = "serif", mgp = c(1.5, 0.5, 0), las = 1, mar = c(3, 3, 1, 1), tcl = 0)
par(mfrow = c(2, 2))

xdrawsn <- length(r$draws("intercept", format = "draws_matrix")[, 1])

# pp checks (panel a) ----
seentogether <- standat$seen_together[standat$time_period == 1]
x <- suppressWarnings(hist(seentogether, plot = FALSE,
                           breaks = seq(-0.5, max(seentogether) + 0.5, 1)))
p <- r$draws(variables = "seen_together_rep", format = "draws_matrix")
p <- p[, standat$time_period == 1]
pd <- as.matrix(t(apply(p, 1, function(X) table(cut(X, breaks = x$breaks, include.lowest = TRUE)))))
sel_draws <- sample(nrow(r$draws("intercept", format = "draws_matrix")), 100)
pd <- apply(pd[sel_draws, ], 2, quantile, probs = c(0.055, 0.5, 0.945))
sel <- which(colSums(rbind(x$counts, pd)) > 0)
maxval <- max(x$counts, pd)
xlim <- range(x$breaks)
plot(0, 0, type = "n", xlim = xlim, ylim = c(0, maxval * 1.15), yaxs = "i",
     axes = FALSE, las = 1, bty = "n",
     xlab = "two individuals seen together", ylab = "frequency", tcl = 0)
axis(1, tcl = 0)
axis(2, las = 1)
for (i in sel) {
  rect(xleft = x$breaks[i], ybottom = 0, xright = x$breaks[i + 1],
       ytop = x$counts[i], col = "grey", border = "white")
  points(x$mids[i], pd[2, i], pch = 21, cex = 1.5, xpd = TRUE, col = "white",
         bg = "black")
  segments(x0 = x$mids[i], y0 = pd[1, i], x1 = x$mids[i], y1 = pd[3, i])
}
box(bty = "l")

# pp checks (panel b) ----
weaned <- standat$model_y_wean[standat$model_period == 1]
x <- suppressWarnings(hist(weaned, plot = FALSE, breaks = seq(-0.5, 1.5, 1)))
p <- r$draws(variables = "model_y_rep", format = "draws_matrix")
p <- p[, standat$model_period == 1]
pd <- as.matrix(t(apply(p, 1, function(X) {
  table(cut(X, breaks = x$breaks, include.lowest = TRUE))
})))
sel_draws <- sample(nrow(r$draws("intercept", format = "draws_matrix")), 100)
pd <- apply(pd[sel_draws, ], 2, quantile, probs = c(0.055, 0.5, 0.945))
sel <- which(colSums(rbind(x$counts, pd)) > 0)
maxval <- max(x$counts, pd)
xlim <- range(x$breaks)
plot(0, 0, type = "n", xlim = xlim, ylim = c(0, maxval * 1.15), yaxs = "i",
     axes = FALSE, las = 1, bty = "n",
     xlab = "weaned offspring", ylab = "frequency", tcl = 0)
axis(1, at = c(0, 1), labels = c("no", "yes"), tcl = 0)
axis(2, las = 1)
xlocs <- list(c(-0.25, 0.25), c(0.75, 1.25))
for (i in sel) {
  rect(xleft = xlocs[[i]][1], ybottom = 0, xright = xlocs[[i]][2], ytop = x$counts[i],
       col = "grey", border = "white")
  points(x$mids[i], pd[2, i], pch = 21, cex = 1.5, xpd = TRUE, col = "white",
         bg = "black")
  segments(x0 = x$mids[i], y0 = pd[1, i], x1 = x$mids[i], y1 = pd[3, i])
}
box(bty = "l")

## components (panel c) ----
sociality_plot(list(mod_res = r), xlim = c(0, 2.5), add_prior = TRUE)


## model fit (panel d) ----
xstrength <- r$draws("strength_for_pp", format = "draws_matrix")
xstrength <- apply(xstrength, 2, quantile, probs = c(0.055, 0.5, 0.945))

plot(xstrength[2, ], standat$model_y, xlim = c(-11, 11), las = 1,
     xlab = "estimated strength (posterior median)", ylab = "weaning probability",
     pch = 16, col = grey(0, 0.5), cex = 1.5, xpd = TRUE)


xvals <- seq(min(xstrength[2, ]), max(xstrength[2, ]), length.out = 51)
xcoefs <- c(r$summary("intercept")$mean,
            0,
            r$summary("slope_strength")$mean,
            0
)

# CI
pdata <- sapply(sample(xdrawsn, 500), function(X){
  lp <- r$draws("intercept")[X] + 0 +
    (r$draws("slope_strength")[X] + 0) * xvals
  plogis(lp)
})

pd <- apply(pdata, 1, quantile, probs = c(0.055, 0.5, 0.945))
polygon(c(xvals, rev(xvals)), c(pd[1, ], rev(pd[3, ])), border = NA,
        col = adjustcolor("#EACB2B", 0.5))

# global fit
points(xvals, plogis(xcoefs[1] + xcoefs[2] + (xcoefs[3] + xcoefs[4]) * xvals),
       type = "l", lwd = 3, col = "black")

# yearly fit
for (i in 1:4) {
  period <- i
  per <- which(standat$model_period == period)
  weaned <- standat$model_y[per]

  xvals <- seq(min(xstrength[2, per]), max(xstrength[2, per]), length.out = 11)
  xcoefs <- c(r$summary("intercept")$mean,
              r$summary(paste0("model_period_intercepts[", period, "]"))$mean,
              r$summary("slope_strength")$mean,
              r$summary(paste0("model_period_strength_slope[", period, "]"))$mean)
  points(xvals, plogis(xcoefs[1] + xcoefs[2] + (xcoefs[3] + xcoefs[4]) * xvals),
         type = "l", lwd = 1, col = "black", lty = 3)

}
```
