## Supplementary material for "(De)composing sociality: disentangling individual-specific from dyad-specific propensities to interact": package tutorial

### intro to `bamoso`

Christof Neumann

2025-05-21 (built with v. 0.5.1)

**Simple model with `brms`** **20**

**References** **24**

This document (and package) is work in progress. Installation instructions can be found at https://github.com/gobbios/bamoso/. If you find mistakes or have suggestions, I'd be very happy to hear about them at https://github.com/gobbios/bamoso/issues.

### Terminology

When **interactions** are mentioned in this document we refer to frequencies and durations of dyadic interactions.

**Components** and **axes** are the overarching terms to refer to **affinity** and **gregariousness**. E.g. 'Affinity is the dyadic component.'

### Quick start

This first example illustrates how to apply the model to pretty much the simplest possible data set. We will skip over a lot of the details, and just show the general workflow.

#### Preparing the data

Here we use grooming data from 19 Barbary macaques (*Macaca sylvanus*) recorded with focal animal sampling. These grooming data represent frequencies of grooming bouts (so we need a count model eventually) and are undirected. We also have information about observation effort, i.e. the time any two individuals *could* have interacted. The data are included in the `bamoso` package and are illustrated in figure 1 (only six individuals shown there). The data are stored initially in matrix form.

| | **i_01** | **i_04** | **i_06** | **i_10** | **i_12** | **i_19** |
|---|---|---|---|---|---|---|
| **i_01** | | 1 | 0 | 0 | 0 | 4 |
| **i_04** | 0 | | 2 | 0 | 0 | 58 |
| **i_06** | 0 | 0 | | 0 | 0 | 3 |
| **i_10** | 0 | 0 | 0 | | 1 | 0 |
| **i_12** | 0 | 0 | 0 | 0 | | 0 |
| **i_19** | 0 | 0 | 0 | 0 | 0 | |

| | **i_01** | **i_04** | **i_06** | **i_10** | **i_12** | **i_19** |
|---|---|---|---|---|---|---|
| **i_01** | | 24.8 | 21.7 | 19.4 | 22.1 | 20.8 |
| **i_04** | 24.8 | | 26.7 | 24.4 | 27.1 | 25.8 |
| **i_06** | 21.7 | 26.7 | | 21.3 | 24 | 22.7 |
| **i_10** | 19.4 | 24.4 | 21.3 | | 21.7 | 20.4 |
| **i_12** | 22.1 | 27.1 | 24 | 21.7 | | 23.1 |
| **i_19** | 20.8 | 25.8 | 22.7 | 20.4 | 23.1 | |

Figure 1: Example data set. Grooming frequencies on the left and dyadic observation hours on the right.

```
library(bamoso)
```

```
data("grooming")
groom_mat <- grooming$syl$groom
obseff_mat <- grooming$syl$obseff
```

Now we need to convert this data set into a format that is suitable to be passed to Stan. There would be a few things to note here but we postpone this to the more extensive example below. Here there are no problems with the data and the conversion is straightforward. We pass a named list with the interaction matrix (`mats = list(groom = groom_mat)`). We indicate that the data represent frequencies or counts (`behav_types = "count"`). And we pass the observation effort as list as well (`obseff = list(obseff_mat)`).

```
standat <- make_stan_data_from_matrices(mats = list(groom = groom_mat),
                                        behav_types = "count",
                                        obseff = list(obseff_mat))
```

#### Fitting the model

Now we can fit the model, using the `sociality_model()` function. This function requires only the converted data set as argument. If you run this, you might get a warning about a few divergent transitions. This is a mild problem here, and we postpone showing how to fix this to further below in the more extensive simulated example.

```
res <- sociality_model(standat)
```

Once the model is fitted, we can obtain a quick summary. If there were sampling issues (like divergent transitions) the summary will report this as well.

```
summary(res)
#> Model of dyadic interactions from 19 individuals (171 dyads)
#> (removed 0 dyads with NA values)
#> number of post-warmup samples: 4000
#> --------------------
#> sociality estimates:
#>   individual gregariousness SD (median [89% CI]): 0.99 [0.55 - 1.50]
#>   pairwise affinity SD (median [89% CI]):         1.84 [1.56 - 2.18]
#> --------------------
#> 1 behavior(s) included:
#> (1): 'groom' (type: 'count')
```

```
#> --------------------
#> no obvious sampling issues detected
```

In this document, sampling went without troubles.

Now we can look into model performance by doing posterior predictive checks. This just means that we look at how well (or badly) the model is able to predict interaction patterns that resemble what we actually observed. And by 'interaction pattern' we simply mean the distribution of grooming bouts across all dyads here. There are different ways of going about that, but this one is easy and if there are severe problems, we will be able to detect them here.

```
pp_model(res, xlab = "grooming bouts", ylab = "(predicted) frequency",
         xbreaks = 20)
pp_model_stat(res, stat = "max", xlab = "maximum number of grooming bouts",
              main = "")
```

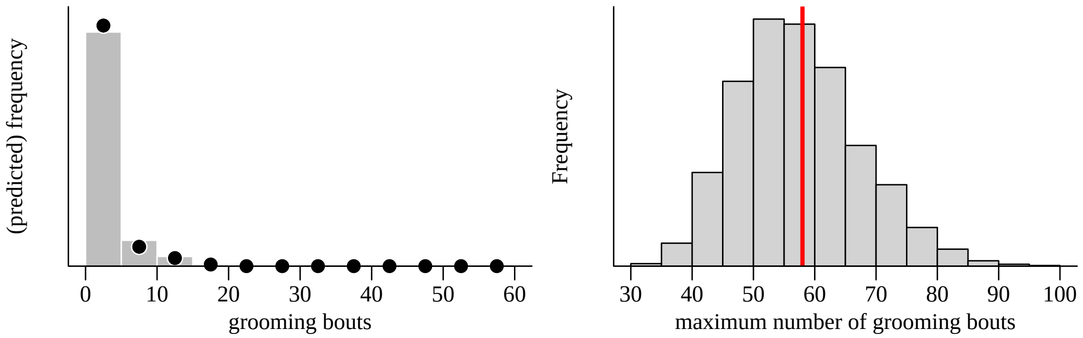

Figure 2: Posterior predictive checks for Barbary macaque grooming data. On the left: bars represent observed grooming frequencies. The circles represent the median frequencies predicted by the model and the vertical bars the 89% interval around these predictions. One the right: the histogram is the posterior distribution of the maximum number of grooming bouts predicted by the model across all posterior samples. The red line indicates the observed value in the data.

The two example checks in figure 2 indicate no obvious problems.

#### Visualising the results

Now we can look at the actual results graphically. We start by looking at the posteriors for the spread (i.e., the SD) of the individual and the dyadic component.

```
sociality_plot(res)
```

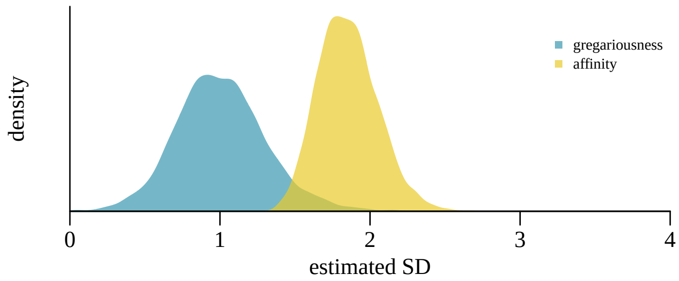

Figure 3: Posteriors for the two sociality components. Variation among dyads is larger than variation among individuals.

We can also display the individual-level gregariousness posteriors using `ridge_plot()` (see figure 4). In principle, this is also possible for the dyadic affinity values, but this becomes cluttered quickly if there are too many dyads to display. In the example in figure 4 only a subset of dyads is therefore displayed.

```
ridge_plot(res, greg = TRUE)
ridge_plot(res, greg = FALSE, sel_subset = 10:20, vert_exp = 15)
```

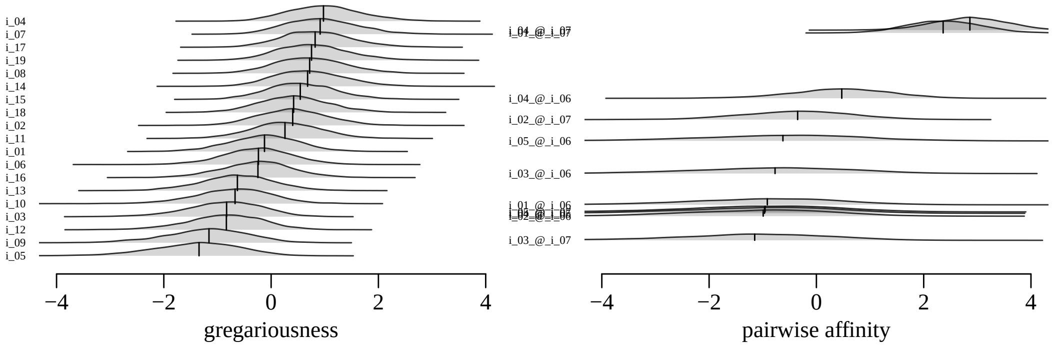

Figure 4: Posteriors for the two sociality components. The affinity distributions on the right only reflect a subset of 10 dyads.

#### Numerical results

To extract numerical results, there are two functions: `model_summary()` and `extract_samples()`. `model_summary()` provides a summary of the posteriors for all the parameters of interest and is simply a data frame.

```
xsum <- model_summary(res)
head(xsum)
```

```
#>     label     categ beh       stanvarname   mean median    sd   5.5%  94.5% rhat ess_bulk ess_tail
#> 1 greg_sd        sd  NA       indi_soc_sd  1.005  0.990 0.301  0.552  1.501    1      642      839
#> 2 affi_sd        sd  NA       dyad_soc_sd  1.853  1.840 0.199  1.559  2.183    1     1294     2461
#> 3   groom intercept  NA beh_intercepts[1] -3.895 -3.887 0.410 -4.561 -3.262    1     1890     2586
#> 4    i_01 indi_vals  NA  indi_soc_vals[1] -0.141 -0.119 0.653 -1.185  0.868    1     2528     2774
#> 5    i_02 indi_vals  NA  indi_soc_vals[2]  0.407  0.398 0.652 -0.589  1.467    1     1783     2646
#> 6    i_03 indi_vals  NA  indi_soc_vals[3] -0.870 -0.827 0.710 -2.070  0.187    1     2612     3137
```

We only see the first six rows, but the structure hopefully becomes clear. Each row represents one parameter. The two top rows represent the estimated variation for the gregariousness and dyadic affinity components (in fact we estimated *standard deviation*). This is followed by the estimates for the behavior intercept(s) (there is only one here because we only modeled one behavior). Next are the individual gregariousness values, followed by the estimates for all the dyads. Alongside the point estimates for the posteriors (mean and median), we get measures of uncertainty around these point estimates (SD, quantiles). And finally, there are some diagnostic values from Stan (rhat, ess_bulk and ess_tail).

In order to get the full posteriors, we use `extract_samples()`. This requires specifying the parameter(s) for which we want to return the posteriors. For example, getting the individual gregariousness values requires `what = "indi_vals"`.[1]

```
e <- extract_samples(res, what = "indi_vals")
head(round(e[, 1:5], 3))
#>        i_01   i_02   i_03  i_04   i_05
#> [1,] -0.196  0.711 -1.019 0.819 -0.875
#> [2,]  0.162  0.629 -0.401 2.189 -2.953
#> [3,]  0.824  0.752 -1.656 1.158 -0.630
#> [4,] -1.030  0.424 -1.508 0.761 -1.365
#> [5,] -0.487 -0.075 -0.453 0.446 -1.998
#> [6,]  0.339  0.613 -1.220 0.531 -1.353
```

#### Summary

This was a brief run through a simple use case of estimating individual-level and dyad-level propensities to interact. It illustrated the key functions of the `bamoso` package, mostly with their default settings.

[1]see the help file (`?extract_samples`) for more information.

### An example with simulated data

We now turn to an example that uses simulated interaction data. To generate these data we use a function included in the package that served as backbone for the evaluations in the manuscript. The key settings for data generation are the arguments for the magnitude of individual- and dyad-level variation (`indi_sd=` and `dyad_sd=`). Both values were chosen to be fairly large. We simulate two behaviors, one a count behavior and the second a proportion. Both also have their own observation effort specified, varying between 5 and 50 ‘hours’ for the count behavior, and between 5 and 100 ‘trials’ for the proportion[2]. The behavior intercepts are both set to -3, which corresponds to a fairly low baseline for each behavior[3]. Recall that these intercepts represent the expected interactions for an average dyad composed of two average individuals.

```
set.seed(1)
d <- generate_data(n_ids = 14, n_beh = 2,
                   behav_types = c("count", "prop"),
                   indi_sd = 1.2, dyad_sd = 0.8,
                   beh_intercepts = c(-1, -1),
                   prop_trials = c(5, 100),
                   count_obseff = c(5, 20))
```

The output of `generate_data()` is already pre-processed to simplify the quantitative evaluations presented in the manuscript. In order to make this example somewhat more realistic, we extract data from it that reflects of what one would typically start with, i.e. interaction matrices and dyadic observation effort (as in the quick start example above). We also label the individuals with letters, which is helpful down the line to keep track of them[4]

```
m1 <- d$processed$interaction_matrices[[1]]
m2 <- d$processed$interaction_matrices[[2]]
colnames(m1) <- rownames(m1) <- LETTERS[1:14]
colnames(m2) <- rownames(m2) <- LETTERS[1:14]

o1 <- m1 - m1
o1[d$standat$dyads_navi] <- d$input_data$obseff[, 1]
o2 <- m2 - m2
o2[d$standat$dyads_navi] <- d$input_data$obseff[, 2]
```

**interactions 1**

| | G | H | I | J | K | L |
|---|---|---|---|---|---|---|
| **G** | | 18 | 96 | 1 | 20 | 9 |
| **H** | 0 | | 16 | 4 | 55 | 3 |
| **I** | 0 | 0 | | 35 | 27 | 7 |
| **J** | 0 | 0 | 0 | | 8 | 0 |
| **K** | 0 | 0 | 0 | 0 | | 6 |
| **L** | 0 | 0 | 0 | 0 | 0 | |

**interactions 2**

| | G | H | I | J | K | L |
|---|---|---|---|---|---|---|
| **G** | | 24 | 40 | 7 | 36 | 4 |
| **H** | 0 | | 34 | 4 | 68 | 18 |
| **I** | 0 | 0 | | 34 | 60 | 36 |
| **J** | 0 | 0 | 0 | | 31 | 0 |
| **K** | 0 | 0 | 0 | 0 | | 29 |
| **L** | 0 | 0 | 0 | 0 | 0 | |

**effort 1**

| | G | H | I | J | K | L |
|---|---|---|---|---|---|---|
| **G** | | 11.3 | 17.6 | 6.2 | 14.7 | 19.6 |
| **H** | 0 | | 16.1 | 12.9 | 12.2 | 10.3 |
| **I** | 0 | 0 | | 19.4 | 12.4 | 10.9 |
| **J** | 0 | 0 | 0 | | 10.7 | 19.3 |
| **K** | 0 | 0 | 0 | 0 | | 6.6 |
| **L** | 0 | 0 | 0 | 0 | 0 | |

**effort 2**

| | G | H | I | J | K | L |
|---|---|---|---|---|---|---|
| **G** | | 42 | 50 | 53 | 69 | 10 |
| **H** | 0 | | 59 | 26 | 90 | 88 |
| **I** | 0 | 0 | | 51 | 89 | 58 |
| **J** | 0 | 0 | 0 | | 95 | 15 |
| **K** | 0 | 0 | 0 | 0 | | 79 |
| **L** | 0 | 0 | 0 | 0 | 0 | |

Figure 5: Interaction matrices of two behaviors (top row), and two observation effort matrices (bottom row). Only a subset of six individuals is displayed.

So we end up with two interaction matrices, and with two matrices for observation effort. For the first behavior, the interactions are counts while the observation effort is continuous (think observation time) (left column in figure 5). For the second behavior, the interactions are also counts, but the observation effort is also recorded as counts, which results in a proportion (right column in figure 5).

[2]Note that the proportion is actually also coded as a count, i.e. ‘number of successes’, which in combination with the ‘trials’ provided results in a proportion.

[3]For a count variable an intercept of $-3$ corresponds to a rate of 0.05 events per 1 unit of observation effort ($exp(-3) = 0.0498$). For the proportion a baseline of $-3$ corresponds to probability of success in a single ‘trial’ of 0.05 ($\text{logit}^{-1}(-3) = 0.0474$)

[4]Stan only works with numeric data so there is no way in Stan to track individuals by their id. Instead, we keep track of ids indirectly, by storing them in the Stan data with a little trick (named vectors).

#### The first behavior

Although we generated two behaviors above, we start with one. The first step converts the dyadic data (and observation effort) into a format that can be passed to Stan. If we supply interaction data and observation effort, we need to make sure that the two matrices match, i.e. they need to have the same dimensions and they need to be ordered in the same way. The easiest way of achieving this is to use column and row names. Currently, there is a rudimentary function `check_data()` to check for consistency in the data, which should flag some (but likely not all) problems in the data.

```
check_data(mats = list(groom = m1),
           behav_types = c("count"),
           obseff = list(o1))
#> dimensions of interaction matrices match (good)
#> dimension of observation effort data match (good)
#> all observation effort data have matching column and row names (good)
#> all interaction matrices have matching column and row names (good)
#> found 14 individuals in all data sources (good)
```

Now we prepare the data. We require two lists, one for the interaction matrix and one for the observation effort. Ideally, we name the interaction matrix when creating the list (`mats = list(groom = m1)`), so that later in any output we can find back the corresponding values by referencing the name(s) supplied here (rather than by their cryptic Stan variable names). If the interaction matrix is not named, the function will make up names (like `behav_A`). The observation effort list does not need to be named.

As far as specifying the `behav_types=` goes: This determines which statistical model will be applied to a given interaction matrix. Table 1 lists the possibilities.

```
sdat1 <- make_stan_data_from_matrices(mats = list(groom = m1),
                                      behav_types = c("count"),
                                      obseff = list(o1))
```

So now we can fit the model. As mentioned in the first example, the only required argument for `sociality_model()` is the prepared data (`sdat1` here). Additional arguments pertain to setting parameters for `cmdstanr`'s `$sample()` method, which is what is running under the hood. In this example, I actually already used one such argument, `seed=16`, which should result in a warning message about divergent transitions.[5]

```
fit1 <- sociality_model(standat = sdat1, seed = 16)
```

Divergent transitions occur quite regularly, but usually can be resolved by setting a higher value to `adapt_delta` (away from its default 0.8). So let's do this, and in the process also change a few more settings, just to illustrate how this works.

```
fit1 <- sociality_model(standat = sdat1,
                        seed = 16,
                        adapt_delta = 0.9,
                        parallel_chains = 4,
                        iter_warmup = 1200)
```

So here we indicate that we want to run four chains in parallel and also increase the number of warm-up iterations. We keep the same random seed. Now the model should sample fine without warnings. Check out the documentation for `sociality_model()` to see other frequently helpful options.

The resulting object is a list with two items, which combines the Stan data used to fit the model and the `cmdstanr` model environment. If you are familiar with `cmdstanr` models, you can directly access the model environment via

[5]The seed argument just fixes the generation of random numbers to a replicable state. In this particular example I chose it such that we actually do get divergent transitions, but it doesn't *cause* the divergent transitions. Try it out with different values and you will see that sometimes divergent transitions occur and sometimes they don't.

Table 1: Interaction types and matching observation effort specification supported in `basomo`.

| name in `basomo` | interactions | observation effort |
|---|---|---|
| `count` | frequencies, counts (integers) | positive continuous |
| `prop` | counts (integers), 'successes' | counts (integers), 'trials' |
| `dur_beta` | continuous proportion (between 0 and 1) | NA |
| `dur_gamma` | positive continuous | positive continuous |
| `dur_gamma0` (experimental) | positive continuous including 0 | positive continuous |
| `binary` (experimental) | 0 or 1 | positive continuous |

```
fit1$mod_res
```

and take it from there. If you are unfamiliar with such objects, the `basomo` package provides a few helper functions to visualize and summarize the results.

Before looking at the results of interest in detail, we look at the model summary.

```
summary(fit1)
#> Model of dyadic interactions from 14 individuals (91 dyads)
#> (removed 0 dyads with NA values)
#> number of post-warmup samples: 4000
#> --------------------
#> sociality estimates:
#>   individual gregariousness SD (median [89% CI]): 1.06 [0.75 - 1.50]
#>   pairwise affinity SD (median [89% CI]):         0.80 [0.67 - 0.96]
#> --------------------
#> 1 behavior(s) included:
#> (1): 'groom' (type: 'count')
#> --------------------
#> sampling issues detected (see below)
```

This just gives an overview over the estimated model, and if there were issues with fitting, these will be reported here too.

#### Model checks

Next, we go to some visual model checks. Because the underlying observed data are discrete (counts in this first example), we use histograms provided by `pp_model()` to look at predictions from the model.[6] One thing to be careful about for these kinds of plot is the width of bins for the histogram. Here I chose 10 bins, ranging from 0 to 100 (143 being the value for the dyad that interacted the most). Actually, I set the bins as a sequence from $-0.5$ to 99.5 in steps of 10 (`bins <- seq(-0.5, 99.5, by = 10)`) to be sure that we cover all values from 0 through 9 in the first bin, from 10 to 19 in the second and so forth.

```
bins <- seq(-0.5, 199.5, by = 10)
pp_model(mod_res = fit1, xbreaks = bins,
         xlab = "count of behaviour 1", ylab = "frequency")
axis(2, las = 1, cex.axis = 0.8)
```

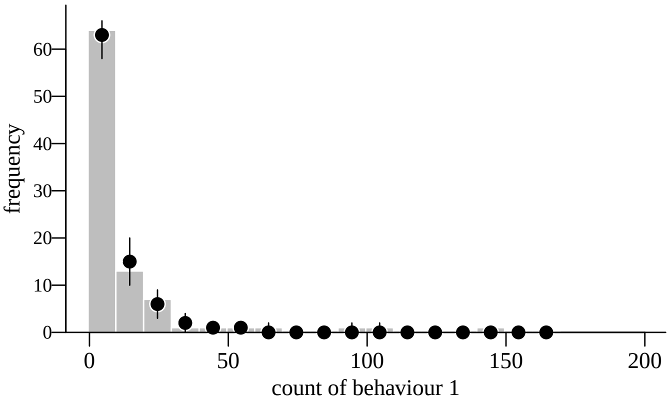

Figure 6: Posterior predictive checks for the entire group.

Figure 6 shows the observed data (for all dyads) as histogram. For example, there are slightly less than 70 dyads with at most 10 interactions (65 actually). The added circles represent the median predicted frequencies for each bin from all model draws (4000 using the default settings). Ideally, these median predictions match the observed frequencies closely, and here they do reasonably well. If the model had difficulties to fit the observed data, we would see predictions that are far from the actually observed data.

One thing to note here is that the model predicts values that were never actually observed. For example, no dyad was observed interacting between 40 and 50 times (no bar in this bin). The model however predicts such values to occur

[6]for continuous data, use density plots from 'pp_model_dens()'.

sometimes (median prediction for that bin is 1 interaction). This is not problematic per se and we will come back to this below.

In addition the plot shows how far the predictions spread around the medians (aka uncertainty). The vertical lines represent 89% credible intervals. These intervals were also calculated from all model draws. We would like to see that these intervals are not too wide. Here they are not, but if they were this just illustrates that our model has not enough information to really make accurate predictions (or that we really have a bad model fit).

One other thing to keep in mind when looking at these predictions is that the assessment of how good the model fit is depends a lot on the resolution, i.e. the bin width. In figure 6 we used bins of width 10, i.e. we counted dyads that interacted between 0 and 9 times, between 10 and 19 times and so forth. For figure 7a we use a finer resolution[7]. Here we see some (slight) discrepancies between observed and predicted values. For example, we observed 4 dyads with 4 interactions, but our model predicted a median of 7 dyads. We also observed 3 dyads with 7 interactions, but the model predicted a median of 4 dyads. In this particular example, I would not consider this problematic though because in absolute terms these cases of under- or over-estimation are of small magnitude (we are not predicting 200 dyads when we observed 3).

At the other extreme end, we could use a very small bin width (figure 7b). Here the median predictions match the observed data almost perfectly. I would not use this plot to conclude that my model fits the data well, but rather I put it here to illustrate the consequences of choosing the bin width.

One last thing these two extreme cases illustrate is how the amount of uncertainty around the median predictions varies with the chosen bin width. For the smaller bin width (figure 7a), the intervals are fairly wide, while the intervals for the very crude bins (figure 7b) are so small that they are not even visible in the figure.

Which bin width to choose is a matter of taste, mostly (I suppose). Bins that are too large won't be useful. The goal is to a get a grasp on how well the model performs. My personal rule of thumb is to use between 10 to 20 bins (equally distributed along the range of data).

```
bins <- seq(-0.5, 199.5, by = 1)
pp_model(mod_res = fit1, xlab = "count of behaviour 1", ylab = "frequency",
         xbreaks = bins, xlim = c(0, 20))
axis(2, las = 1, cex.axis = 0.8)

bins <- seq(-0.5, 199.5, by = 50)
pp_model(mod_res = fit1, xlab = "count of behaviour 1", ylab = "frequency",
         xbreaks = bins, xlim = c(0, 200))
axis(2, las = 1, cex.axis = 0.8)
```

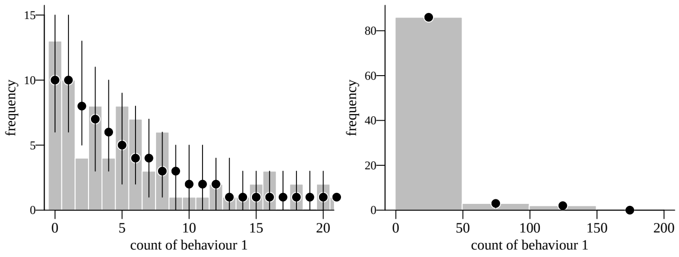

Figure 7: Posterior predictive checks for the entire group with different resolutions along the x-axis (compare to figure 6). Note that the first plot is cut at 20 along the x-axis for visual purposes.

To dig into model performance even further, we can stratify the predictions by individual. In other words, we can select a specific individual and look at observed and predicted interactions for all dyads that this specific individual was part of. We use the same function (`pp_model()`), but supply the argument `selected_id=`. Figure 8 shows the results for the first three individuals (A through C).

The building parts of these plots are the same as above. There are at least two things to note here. First, the uncertainties around the predictions might, on a first glance, appear larger than in the group level plots above. But note the range along the vertical axis: we are talking about much smaller number of dyads (each plot only reflects the portion of dyads the selected individual is part of), so in absolute terms these uncertainties are very similar to the

[7] Note that we want to explicitely separate the occurences of 0 and 1 interactions, which is why we set the break points for the bin widths to start at -0.5.

ones in the group level plots. Second, recall that the model also predicts interaction numbers that were not actually observed. Individual E for example, was part of a dyad that interacted 15 times, which is an extreme value for that particular individual[8]. Now, the model predicts also values of 150 up to 200 interactions for that dyad. The key point here is that this not problematic, but just illustrates that the model is not predicting *perfectly* but in any case does a good job at predicting that this dyad interacts substantially more than the other dyads of individual E.

```
bins <- seq(-0.5, 199.5, by = 2)
pp_model(mod_res = fit1, selected_id = "A", xbreaks = bins, xlim = c(-1, 50),
         xlab = "count of behaviour 1", ylab = "frequency")
legend("top", "A", bty = "n")
axis(2, las = 1)

bins <- seq(-0.5, 199.5, by = 5)
pp_model(mod_res = fit1, selected_id = "B", xbreaks = bins, xlim = c(-1, 200),
         xlab = "count of behaviour 1", ylab = "frequency")
legend("top", "B", bty = "n")
axis(2, las = 1)

bins <- seq(-0.5, 199.5, by = 10)
pp_model(mod_res = fit1, selected_id = "E", xbreaks = bins, xlim = c(-1, 200),
         xlab = "count of behaviour 1", ylab = "frequency")
legend("top", "E", bty = "n")
axis(2, las = 1)
```

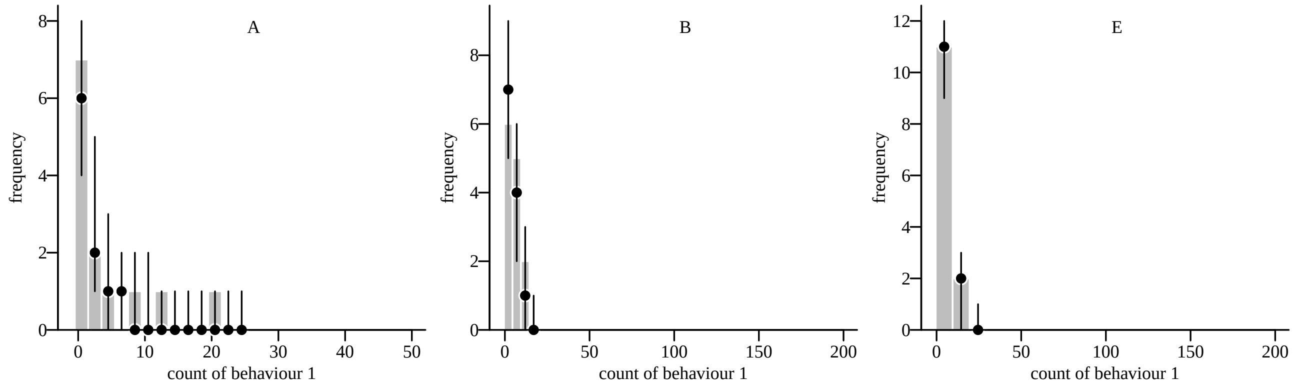

Figure 8: Posterior predictive checks for selected individuals.

Finally, there are even more ways to illustrate how well or badly the model predicts our observations. We could for example look at the maximum value of interactions predicted. A good model would on average predict maximum interactions that are close on average to the observed data. So we extract for each model draw the number of interactions for the dyad that interacted the most. Figure 9a shows a histogram of these (4,000 by default) maximum values. The red line represents what we observed in the actual data. The conclusion here is that our model performs well with respect to predicting interaction patterns that are consistent with our observed data. Figure 9b shows the same approach for the inter-quartile range.

At the moment, there are only a handful of these summary statistics defined in `pp_model_stat()` (`"mean"`, `"min"`, `"max"`, `"iqr"`, and `"range_width"`).

```
pp_model_stat(mod_res = fit1, stat = "max", main = "", xlab = "max interactions")
pp_model_stat(mod_res = fit1, stat = "iqr", main = "", xlab = "IQR of interactions")
```

If you want to dig deeper into to the concept of visual model checks, Gabry et al. (2019) is a good starting point.

#### Visual model results

Visualization of the key results works the same way as in the first example. `sociality_plot()` produces a plot of the posteriors of the variation (the estimated SD, actually) of the two sociality components (figure 10). There are number of arguments to the function in addition to the one required (`mod_res = fit1`), and below I specify them at their

[8] It could be that this dyad has a very strong affinity, or both dyad members were extremely gregarious, or we simply observed this dyad for a long time.

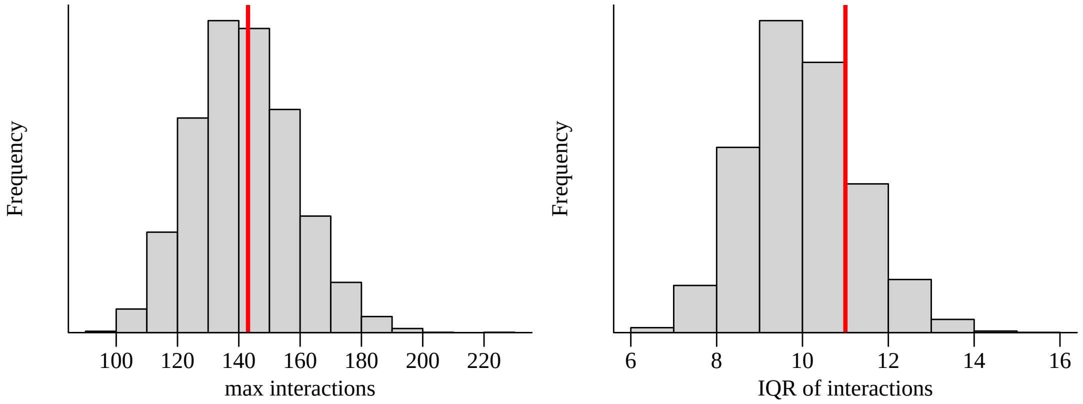

Figure 9: Posterior predictive checks for the entire group using summary statistics.

defaults. Their meaning should be self-explanatory. Since this model is based on simulated data, we can also add the ground truth for both parameters, which are the vertical lines at 1.2 and 0.8.

```
sociality_plot(mod_res = fit1, do_legend = TRUE, do_dyadic = TRUE, do_indi = TRUE)
abline(v = d$input_data$indi_sd) # value used during data generation
```

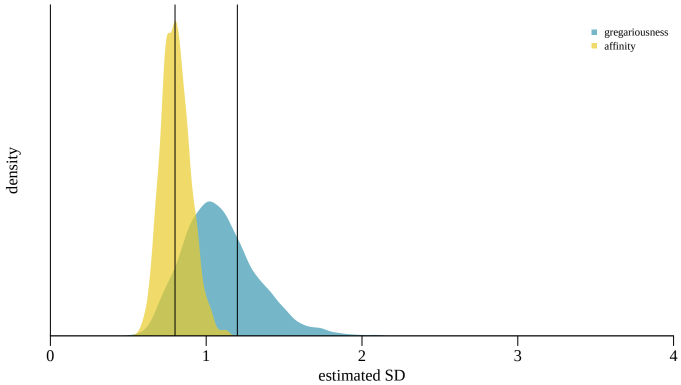

Figure 10: Individual and dyadic variation components.

We can also plot the posteriors of the estimated individual gregariousness and dyadic affinity values with `ridge_plot()` (figure 11). Note that these plots become cluttered quickly depending on the number of individuals in the data set, and this is especially true for plotting dyadic estimates.

```
ridge_plot(fit1)
ridge_plot(fit1, sel_subset = c("A", "C"))
ridge_plot(fit1, greg = FALSE, sel_subset = c("F_@_J", "M_@_N", "D_@_L"), vert_exp = 10)
ridge_plot(fit1, greg = FALSE, sel_subset = c("F", "A_@_B", "D_@_L"), vert_exp = 10)
```

#### Numeric model results

There are basically two ways of getting numerical output of the model results. `model_summary()` returns summaries of the posteriors, and `extract_samples()` extracts the entire posteriors of the primary parameters.

`model_summary()` as used below returns a data frame with point estimates (mean and median) and some measures of uncertainty and sampling diagnostics. The returned parameters are grouped in the `$categ` column as either `sd` (standard deviations of gregariousness and pairwise affinity), `intercept` (intercepts for all behaviors), `indi_vals` (gregariousness for each individuals) and `dyad_vals` (pairwise affinity for each dyad), which allows filtering. The only relevant argument to the function is needed for specifying the quantiles to be returned for each posterior (the default is an 89% interval: `probs = c(0.055, 0.945)`).

```
modsum <- model_summary(fit1, probs = c(0.1, 0.9))
head(modsum)
head(modsum[modsum$categ == "dyad_vals", ])
```

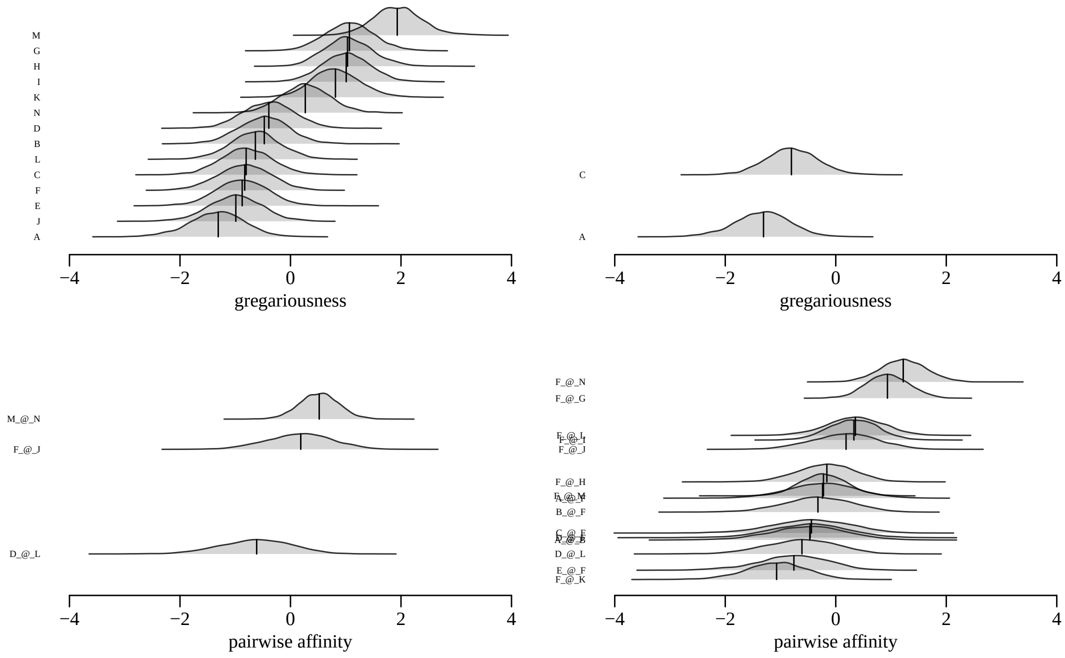

Figure 11: Posteriors for individual gregariousness and dyadic affinity.

```
#>     label     categ beh       stanvarname   mean median     sd    10%     90% rhat ess_bulk ess_tail
#> 1 greg_sd        sd  NA       indi_soc_sd  1.090  1.065 0.2367  0.807  1.4084    1     1080     1488
#> 2 affi_sd        sd  NA       dyad_soc_sd  0.806  0.801 0.0934  0.691  0.9308    1     1366     2359
#> 3   groom intercept  NA beh_intercepts[1] -0.894 -0.882 0.4192 -1.429 -0.3785    1      677     1037
#> 4       A indi_vals  NA  indi_soc_vals[1] -1.329 -1.310 0.5026 -1.962 -0.7021    1     1297     1709
#> 5       B indi_vals  NA  indi_soc_vals[2] -0.486 -0.476 0.4638 -1.085  0.0834    1     1230     1842
#> 6       C indi_vals  NA  indi_soc_vals[3] -0.795 -0.803 0.4693 -1.394 -0.1962    1     1074     2084
```

```
#>    label     categ beh      stanvarname   mean median    sd    10%    90% rhat ess_bulk ess_tail
#> 18 A_@_B dyad_vals  NA dyad_soc_vals[1] -0.497 -0.472 0.698 -1.399 0.3822    1     4077     2968
#> 19 A_@_C dyad_vals  NA dyad_soc_vals[2] -0.359 -0.347 0.703 -1.238 0.5084    1     5069     3021
#> 20 B_@_C dyad_vals  NA dyad_soc_vals[3] -0.646 -0.625 0.593 -1.407 0.0968    1     3354     2971
#> 21 A_@_D dyad_vals  NA dyad_soc_vals[4] -0.183 -0.184 0.615 -0.971 0.5941    1     4478     3199
#> 22 B_@_D dyad_vals  NA dyad_soc_vals[5]  0.155  0.158 0.513 -0.490 0.8058    1     2903     2402
#> 23 C_@_D dyad_vals  NA dyad_soc_vals[6]  0.814  0.812 0.470  0.221 1.4204    1     2790     2736
```

With `extract_samples()`, we can get the full posteriors for each parameter. Here, the output is a matrix, where each row represents one posterior sample (so the total number of rows is by default 4,000, i.e. 4 chains each with 1,000 samples). The columns represent the parameters and they are grouped in the same way as in `model_summary()`, i.e. the two standard deviations for gregariousness and affinity, followed by the intercepts for each behavior, followed by the values for each individual and dyad.

```
post <- extract_samples(fit1)
#> model doesn't contain correlations (removing 'indi_cors' and/or 'dyad_cors' from argument 'what = ')
head(post)[, 1:12]
```

```
#>      greg_sd affi_sd groom     A     B     C      D       E     F   G    H    I
#> [1,]    1.21    0.72 -1.15 -1.91 -0.16 -0.73 -0.023 -0.9157 -0.72 1.4 1.26 1.69
#> [2,]    1.07    0.80 -1.30 -1.62 -0.85 -0.38 -0.183 -0.6776 -1.13 1.9 1.43 1.56
#> [3,]    1.06    0.80 -1.36 -1.55 -0.87 -0.67 -0.081 -0.5961 -0.93 1.8 1.37 1.60
#> [4,]    1.40    0.85 -1.35 -0.93 -0.25 -0.45  0.177 -0.5515 -1.54 1.8 1.11 2.32
#> [5,]    1.10    0.75 -0.99 -1.35 -1.19 -0.30 -0.610 -1.1980 -0.88 1.9 1.96 0.92
#> [6,]    0.81    0.78 -1.13 -1.15 -0.14 -0.99 -0.321 -0.0035 -0.99 1.0 0.98 1.26
```

There is one relevant argument for `extract_samples()` is `what=`, and this determines which parameters are returned. The default is to extract all parameters, but this can be changed for example to return only the dyadic affinity values. For the remaining options, see the function's help file (`?extract_samples`).

```
post <- extract_samples(fit1, what = "dyad_vals")
head(post)[, 1:12]
```

```
#>      A_@_B  A_@_C   B_@_C  A_@_D  B_@_D C_@_D A_@_E B_@_E  C_@_E D_@_E  A_@_F   B_@_F
#> [1,] -1.17 -1.437 -0.3985 -0.740 -0.283  0.44  0.14 -0.13 -0.406 0.787  0.162 -0.0069
#> [2,] -0.69  0.099  0.2093  0.498  0.175  1.10 -0.51  0.38 -0.022 0.808  0.398 -0.1372
#> [3,] -0.85  0.399  0.0904  0.474  0.397  1.18 -0.81  0.42 -0.110 0.665  0.574 -0.0796
#> [4,] -0.23 -0.626 -0.7864 -0.611  0.113  0.36 -0.84  0.46 -0.751 0.210  0.035  0.3673
#> [5,] -0.22 -0.514 -0.0057  0.086  0.977  1.08 -0.10  0.90 -0.549 1.147 -0.032  0.4062
#> [6,] -1.00 -0.190 -0.8781 -0.437  0.024 -0.02 -1.00 -0.12 -0.069 0.065 -0.388 -0.4321
```

#### The second behavior

Now we turn to the second behavior, which represents a proportion. Here, the interaction matrix also contains integers, and crucially, the corresponding observation effort is also recorded as integer values. In other words, it's only through the combination of interactions and observation effort that we can actually talk about proportions.

Apart from that, everything works the same as in the example(s) above. We prepare the data and fit the model.

```
sdat2 <- make_stan_data_from_matrices(mats = list(prox = m2),
                                      behav_types = c("prop"),
                                      obseff = list(o2))
```

```
fit2 <- sociality_model(standat = sdat2, seed = 8)
```

We now also run posterior predictive checks, but this time we look at densities rather than at histograms (figure 12). This makes sense if the quantity we want to display can be seen as continuous.[9]

The function we use is `pp_model_dens()`. The underlying idea is the same as in `pp_model()`: we plot the density of the observed data (red line in the plot) and combine this with densities from a small number of draws representing predictions from the model (the default is `n_draws=20`). A good model leads to predictions that correspond approximately to the observed data.

```
pp_model_dens(mod_res = fit2, xlab = "count of behaviour 2", ylab = "frequency")
axis(2, las = 1, cex.axis = 0.8)
```

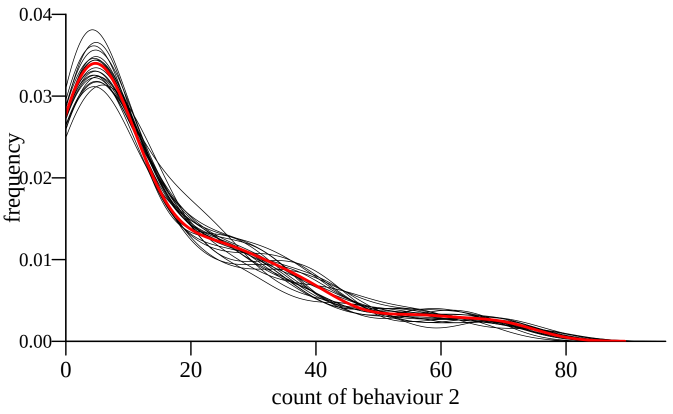

Figure 12: Posterior predictive checks for a behavior representing a proportion.

We can also produce a plot of the two sociality components using `sociality_plot()` (figure 13).

```
sociality_plot(mod_res = fit2)
abline(v = 1.5)
```

#### Combining two behaviors

It becomes really interesting when we model two behaviors simultaneously. In other words, we want to estimate the underlying sociality components from two or more interaction matrices. There is the implicit assumption that such underlying sociality components exist in the first place.

So when preparing the data, we just need to make sure that we combine the two behaviors in question:

```
sdat3 <- make_stan_data_from_matrices(mats = list(groom = m1, prox = m2),
                                      behav_types = c("count", "prop"),
                                      obseff = list(o1, o2))
```

[9]Technically, these data are still discrete and not continuous, but we proceed anyway for illustration purposes.

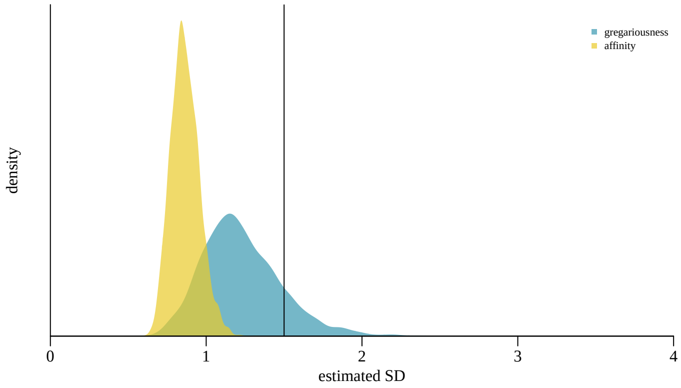

Figure 13: Individual and dyadic variation components.

The fitting of the model then works the same way as before.

```
fit3 <- sociality_model(standat = sdat3,
                        refresh = 0, parallel_chains = 4, seed = 3)
```

```
summary(fit3)
#> Model of dyadic interactions from 14 individuals (91 dyads)
#> (removed 0 dyads with NA values)
#> number of post-warmup samples: 4000
#> --------------------
#> sociality estimates:
#>   individual gregariousness SD (median [89% CI]): 1.18 [0.86 - 1.68]
#>   pairwise affinity SD (median [89% CI]):         0.82 [0.71 - 0.97]
#> --------------------
#> 2 behavior(s) included:
#> (1): 'groom' (type: 'count')
#> (2): 'prox' (type: 'prop')
#> --------------------
#> no obvious sampling issues detected
```

The most important thing to remember when working with more than one behavior is that we have to declare for which behavior we want to make predictions when using `pp_model()` and `pp_model_dens()` (figure 14).

```
pp_model(mod_res = fit3, xvar = "groom",
         xlab = "grooming frequency", ylab = "frequency")
pp_model_dens(mod_res = fit3, xvar = "prox",
         xlab = "prox frequency", ylab = "frequency")
```

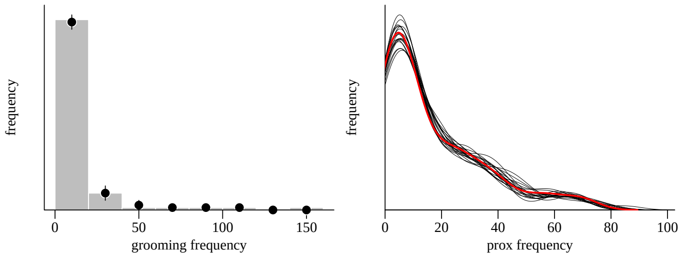

Figure 14: Posterior predictive checks for two behaviors.

### Extending the model

#### More than one behaviour with correlated axes

As mentioned above, when we fit the model to data we make the implicit (or hopefully explicit) assumption that there is exactly one, not two or three, underlying gregariousness axes (and of course exactly one affinity axis (not two or three or . . . )). But this is just an assumption. We might as well make the assumption that more than one gregariousness axis exists and that they are correlated. Imagine two behaviors. Here we could estimate two gregariousness axes, one for each behavior. Plus, we can actually investigate whether these multiple axes are correlated with each other.

Let's look at an example with three behaviors, so we have three gregariousness axes and three affinity axes. Since there are three behaviors there are also three correlations between them (G1 with G2, G1 with G3 and G2 with G3 (and the same for affinity)). We simulate the data so we can actually be sure what the true correlations are. Without going into too much detail: we simulate interactions such that the correlations are $-0.4$, 0.4, and 0.6 for gregariousness. We also set correlations for affinity. These values are handed over to `generate_data` via SD/correlation matrices, which also encode the ground truth SD values in the diagonal.

```
corgreg12 <- -0.4
corgreg13 <- 0.4
corgreg23 <- 0.6

gregmat <- matrix(c(0.9, corgreg12, corgreg13,
                    corgreg12, 1.5, corgreg23,
                    corgreg13, corgreg23, 1.2),
                  ncol = 3, byrow = TRUE)

coraffi12 <- -0.6
coraffi13 <- 0.3
coraffi23 <- -0.4
affimat <- matrix(c(0.5, coraffi12, coraffi13,
                    coraffi12, 1.2, coraffi23,
                    coraffi13, coraffi23, 1.2),
                  ncol = 3, byrow = TRUE)

set.seed(1)
ex <- generate_data(n_ids = 18, n_beh = 3,
                    behav_types = c("count", "prop", "prop"),
                    indi_sd = gregmat,
                    dyad_sd = affimat,
                    beh_intercepts = c(1.4, -0.7, -1.7), exact = TRUE)
```

The `generate_data()` function actually prepares the data directly, but we take the detour and create everything by hand. Here are the interactions matrices

```
b1 <- ex$processed$interaction_matrices[[1]]
b2 <- ex$processed$interaction_matrices[[2]]
b3 <- ex$processed$interaction_matrices[[3]]
```

and the observation effort matrices[10].

```
o1 <- ex$processed$obseff_matrices[[1]]
o2 <- ex$processed$obseff_matrices[[2]]
o3 <- ex$processed$obseff_matrices[[3]]
```

Now we can prepare the data for usage with `sociality_model()`. In fact, we want to fit two models. One should mirror the true structure of the data-generating process and one should be oblivious and keep assuming single gregariousness and affinity axes.

```
s1 <- make_stan_data_from_matrices(mats = list(b1 = b1, b2 = b2, b3 = b3),
                                   behav_types = c("count", "prop", "prop"),
                                   obseff = list(o1, o2, o3))
```

And to indicate that we want to fit separate axes and correlations we set `correlation = TRUE`:

```
s2 <- make_stan_data_from_matrices(mats = list(b1 = b1, b2 = b2, b3 = b3),
                                   behav_types = c("count", "prop", "prop"),
                                   obseff = list(o1, o2, o3),
                                   correlations = TRUE)
```

[10]The matrices with the observation effort are somewhat simple looking, because during the data generation we assume equal observation effort over all dyads

With these two data objects we can now fit the two models.

```
# incorrect model
r1 <- sociality_model(s1, parallel_chains = 4, refresh = 0,
                      show_exceptions = FALSE, seed = 42)
```

```
# correct model
r2 <- sociality_model(s2, parallel_chains = 4, refresh = 0,
                      show_exceptions = FALSE, seed = 42)
```

```
summary(r1)
#> Model of dyadic interactions from 18 individuals (153 dyads)
#> (removed 0 dyads with NA values)
#> number of post-warmup samples: 4000
#> --------------------
#> sociality estimates:
#>   individual gregariousness SD (median [89% CI]): 0.80 [0.62 - 1.08]
#>   pairwise affinity SD (median [89% CI]):         0.54 [0.49 - 0.61]
#> --------------------
#> 3 behavior(s) included:
#> (1): 'b1' (type: 'count')
#> (2): 'b2' (type: 'prop')
#> (3): 'b3' (type: 'prop')
#> --------------------
#> no obvious sampling issues detected
```

```
summary(r2)
#> Model of dyadic interactions from 18 individuals (153 dyads)
#> (removed 0 dyads with NA values)
#> number of post-warmup samples: 4000
#> --------------------
#> sociality estimates (by behavior):
#> (1): 'b1' (type: 'count'):
#>      individual gregariousness SD (median [89% CI]): 0.81 [0.62 - 1.09]
#>      pairwise affinity SD (median [89% CI]):         0.58 [0.48 - 0.67]
#> (2): 'b2' (type: 'prop'):
#>      individual gregariousness SD (median [89% CI]): 1.38 [1.07 - 1.82]
#>      pairwise affinity SD (median [89% CI]):         1.19 [1.07 - 1.34]
#> (3): 'b3' (type: 'prop'):
#>      individual gregariousness SD (median [89% CI]): 1.21 [0.90 - 1.63]
#>      pairwise affinity SD (median [89% CI]):         1.28 [1.15 - 1.44]
#> --------------------
#> correlations (individual gregariousness):
#>    b1 * b2 (median [89% CI]): -0.47 [-0.72 - -0.15]
#>    b1 * b3 (median [89% CI]): 0.33 [-0.01 - 0.60]
#>    b2 * b3 (median [89% CI]): 0.40 [0.06 - 0.66]
#> --------------------
#> correlations (dyadic affinity):
#>    b1 * b2 (median [89% CI]): -0.52 [-0.65 - -0.36]
#>    b1 * b3 (median [89% CI]): 0.22 [0.03 - 0.40]
#>    b2 * b3 (median [89% CI]): -0.30 [-0.43 - -0.16]
#> --------------------
#> 3 behavior(s) included:
#> (1): 'b1' (type: 'count')
#> (2): 'b2' (type: 'prop')
#> (3): 'b3' (type: 'prop')
#> --------------------
#> no obvious sampling issues detected
```

Those results for the 'correct' model look alright (the estimated correlations are close to the true values and so are the SDs and intercepts). In contrast, the incorrect model doesn't even get the SDs right (and how could it, the model only estimates one individual and one dyadic axis, no three). But how do we know which model to use? In reality we don't know the true model. The obvious choice would be to use some kind of model comparison, which should at least let us drop the worst models. But that's yet to be implemented.

For now, let's look at posterior predictive checks. And there it becomes clear that there is a problem with the second model (figures 15 and 16), while the first model seems to do a reasonable job.

And finally, let's look at some results plots of the model with the correlations.

```
par(mfcol = c(1, 3), family = "serif", mar = c(3, 2, 1, 1), xaxs = "i")
sociality_plot(r2, which_cor = c(1, 2), xlim = c(-1, 1))
abline(v = rep(gregmat[1, 2], 2), col = c("darkgrey", "#3B99B1B3"), lwd = c(3.5, 2))
```

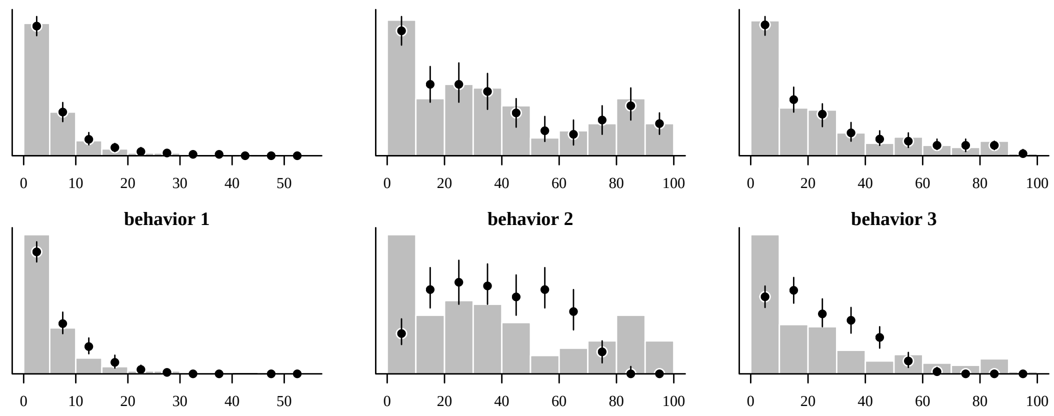

Figure 15: Top row: correct model, bottom row: incorrect model. Histogram is the observed data.

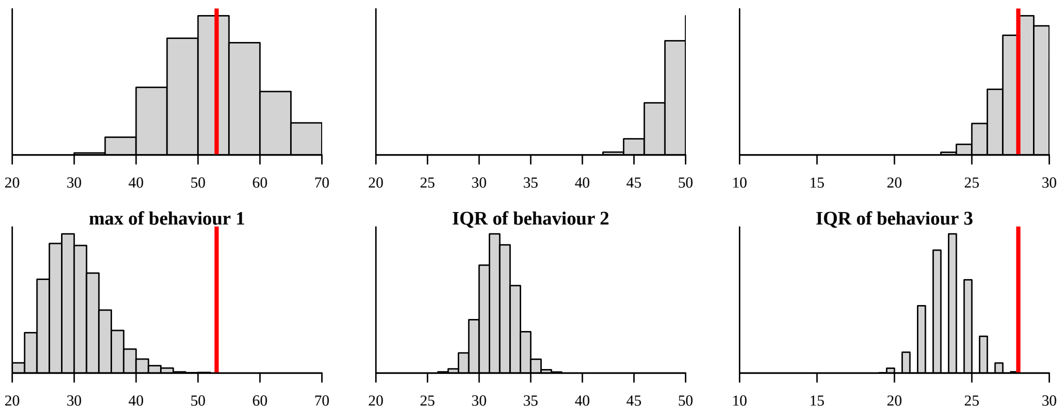

Figure 16: Top row: correct model, bottom row: incorrect model. Red is the observed value.

```
abline(v = rep(affimat[1, 2], 2), col = c("darkgrey", "#EACB2BB3"), lwd = c(3.5, 2))

sociality_plot(r2, which_cor = c(1, 3), xlim = c(-1, 1))
abline(v = rep(gregmat[1, 3], 2), col = c("darkgrey", "#3B99B1B3"), lwd = c(3.5, 2))
abline(v = rep(affimat[1, 3], 2), col = c("darkgrey", "#EACB2BB3"), lwd = c(3.5, 2))

sociality_plot(r2, which_cor = c(2, 3), xlim = c(-1, 1))
abline(v = rep(gregmat[2, 3], 2), col = c("darkgrey", "#3B99B1B3"), lwd = c(3.5, 2))
abline(v = rep(affimat[2, 3], 2), col = c("darkgrey", "#EACB2BB3"), lwd = c(3.5, 2))
```

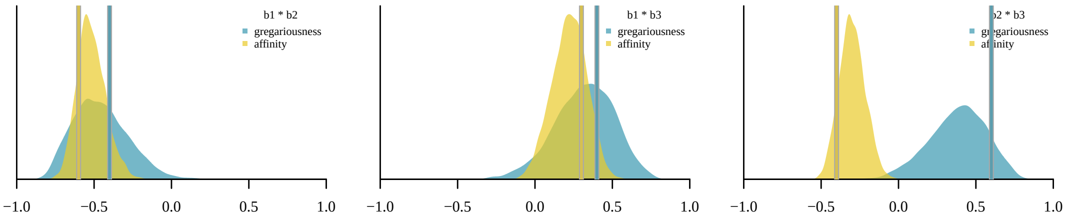

Figure 17: Posteriors of estimated correlations. Vertical lines indicate true values (input values for simulated data).

```
par(mfcol = c(1, 3), family = "serif", mar = c(3, 2, 1, 1), xaxs = "i")
sociality_plot(r2, which_beh = 1, xlim = c(0, 3))
abline(v = rep(gregmat[1, 1], 2), col = c("darkgrey", "#3B99B1B3"), lwd = c(3.5, 2))
abline(v = rep(affimat[1, 1], 2), col = c("darkgrey", "#EACB2BB3"), lwd = c(3.5, 2))

sociality_plot(r2, which_beh = 2, xlim = c(0, 3))
abline(v = rep(gregmat[2, 2], 2), col = c("darkgrey", "#3B99B1B3"), lwd = c(3.5, 2))
abline(v = rep(affimat[2, 2], 2), col = c("darkgrey", "#EACB2BB3"), lwd = c(3.5, 2))

sociality_plot(r2, which_beh = 3, xlim = c(0, 3))
abline(v = rep(gregmat[3, 3], 2), col = c("darkgrey", "#3B99B1B3"), lwd = c(3.5, 2))
abline(v = rep(affimat[3, 3], 2), col = c("darkgrey", "#EACB2BB3"), lwd = c(3.5, 2))
```

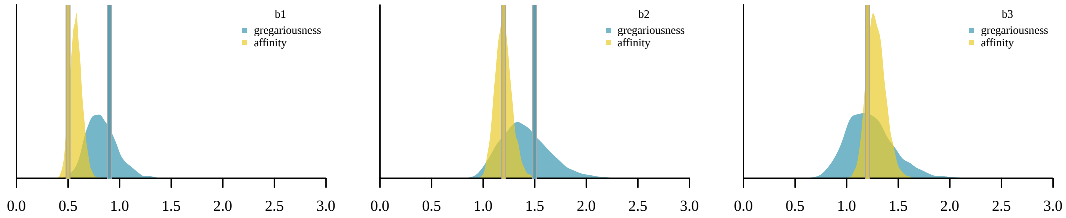

Figure 18: Posteriors of estimated differentiation in gregariousness and affinity. Vertical lines indicate true values (input values for simulated data).

#### Correlation example with empirical data

TBD.

more to be written

#### Prior predictive simulations

For now, the priors for most parameters in the model are hardcoded in the Stan model code.[11] This is likely to change in the future, but for now the priors for the standard deviations of the individual gregariousness and the dyadic affinity seem to work reasonably well across a range of situations.

The simulations in this section just demonstrate what happens if we use only the priors to estimate the parameters of interest. Actually, we still need the data because we require the observation effort (if present) and the interactions (to determine the priors for the behavioral intercepts).

```
set.seed(123)
x <- generate_data(n_ids = 33, n_beh = 1, behav_types = "count",
                   indi_sd = 0.3, dyad_sd = 0.3,
                   beh_intercepts = 0.2)

mats <- list(x$processed$interaction_matrices[[1]])
obseff <- list(x$processed$obseff_matrices[[1]])
dat <- make_stan_data_from_matrices(mats = mats, obseff = obseff,
                                    behav_types = "count")

# prior only ('null model')
rnull <- sociality_model(dat, prior_sim = TRUE,
                         parallel_chains = 4, refresh = 0,
                         iter_sampling = 247, seed = 42)
# actual model
r <- sociality_model(dat, prior_sim = FALSE,
                     parallel_chains = 4, refresh = 0,
                     iter_sampling = 247, seed = 42)
```

In the prior only model, we expect to see an intercept of 0.2 (as we set in the data generation step). The estimates for both variance components should be around 0.5 (which is the mean of the prior distribution (exponential(2))). So that works quite well.

```
rnull$mod_res$summary(c("beh_intercepts", "indi_soc_sd", "dyad_soc_sd"))
```

When we condition on the actual data, we still see the intercept of around 0.2 . But this time we recover the actual parameter values (0.3 and 0.3) (reasonably well, (I'm very generous with rounding here)). If you are not impressed, simulate again with a larger number of individuals.

```
r$mod_res$summary(c("beh_intercepts", "indi_soc_sd", "dyad_soc_sd"))
```

We can also visualize these simulations. We do so by using the prior-only model to make predictions about the behavior. These predictions only 'know' two things from the observed data: the mean of the interaction numbers and the observation effort (if any). And now we predict interactions. The plot below shows this in a quick and dirty way[12]

```
pp_model_dens(rnull, xlim = c(0, 20), ylim = c(0, 0.5), n_draws = 20, xadjust = 2)
pp_model_dens(r1, ylim = c(0, 0.5), xlim = c(0, 50), n_draws = 100, xadjust = 2)
```

[11] The exceptions are the priors for the behavioural intercepts, which are derived from the observed interactions.
[12] Confusingly, it also displays the observed data in red..., this is gonna be turned off sometime soon.

### Intuitions

In this section, I just want to demonstrate a few intuitions I have/had and see how they play out.

#### Uncertainty and observation effort

to be written

#### Correlated axes 1

We assume that when we have two or more behaviors, that they have one underlying affinity and gregariousness axis. This should be essentially the same as assuming that we have one axis per behavior and that these axes perfectly correlated with each other. So if we generate a data set of interactions of two behaviors and we generate them with two axes that strongly correlate, I would expect no major difference in inference between a model that incorporate the correlations and one that doesn't.

```
post_withcor <- extract_samples(r_withcor, what = c("indi_sd", "dyad_sd"))
post_nocor <- extract_samples(r_nocor, what = c("indi_sd", "dyad_sd"))
apply(post_withcor, 2, median)
apply(post_nocor, 2, median)
```

So that didn't work out as I expected... What I overlooked (duh) was that this intuition can only be correct if the SD of the two affinity axes is identical (and not only the correlation large). So if we look at the results of the models this becomes apparent quickly.

```
par(family = "serif", mgp = c(1.5, 0.5, 0), las = 1, mar = c(3, 3, 1, 1), tcl = 0)
par(mfrow = c(2, 3))

p <- model_summary(r_withcor)
pa <- p[p$categ == "dyad_vals" & p$beh == "behav_A", "median"]
pb <- p[p$categ == "dyad_vals" & p$beh == "behav_B", "median"]

p <- model_summary(r_nocor)
ps <- p[p$categ == "dyad_vals", "median"]

plot(x$input_data$dyad_soc_vals[, 1], pa, asp = 1, xlab = "true values (behavior 1)",
     ylab = "estimated (cor model, behav 1)",
     xlim = c(-4, 4), ylim = c(-4, 4), yaxs = "i", xaxs = "i")
abline(0, 1, lty = 2)
plot(x$input_data$dyad_soc_vals[, 1], pb, asp = 1, xlab = "true values (behavior 1)",
     ylab = "estimated (cor model, behav 2)",
     xlim = c(-4, 4), ylim = c(-4, 4), yaxs = "i", xaxs = "i")
abline(0, 1, lty = 2)
plot(x$input_data$dyad_soc_vals[, 1], ps, asp = 1, xlab = "true values (behavior 1)",
     ylab = "estimated (model without cor)",
     xlim = c(-4, 4), ylim = c(-4, 4), yaxs = "i", xaxs = "i")
abline(0, 1, lty = 2)

plot(x$input_data$dyad_soc_vals[, 2], pa, asp = 1, xlab = "true values (behavior 2)",
     ylab = "estimated (cor model, behav 1)",
     xlim = c(-4, 4), ylim = c(-4, 4), yaxs = "i", xaxs = "i")
abline(0, 1, lty = 2)
plot(x$input_data$dyad_soc_vals[, 2], pb, asp = 1, xlab = "true values (behavior 2)",
     ylab = "estimated (cor model, behav 2)",
     xlim = c(-4, 4), ylim = c(-4, 4), yaxs = "i", xaxs = "i")
abline(0, 1, lty = 2)
plot(x$input_data$dyad_soc_vals[, 2], ps, asp = 1, xlab = "true values (behavior 2)",
     ylab = "estimated (model without cor)",
     xlim = c(-4, 4), ylim = c(-4, 4), yaxs = "i", xaxs = "i")
abline(0, 1, lty = 2)
```

#### Correlated axes 2

```
set.seed(123)
cors_indi <- matrix(c(1.3, 0.7, 0.7, 1.3), ncol = 2)
cors_dyad <- matrix(c(0.3, 0.3, 0.3, 0.3), ncol = 2)
x <- generate_data(n_ids = 12, n_beh = 2,
                   beh_intercepts = c(0.5, 0.5), behav_types = c("count", "count"),
```

```
                   indi_sd = cors_indi, dyad_sd = cors_dyad)
s <- x$standat
r <- sociality_model(s, parallel_chains = 4, seed = 1, adapt_delta = 0.85,
                     show_exceptions = FALSE, refresh = 0,
                     iter_warmup = 500, iter_sampling = 1000)
summary(r)
s1 <- extract_samples(r, "indi_vals", axis = 1)
s2 <- extract_samples(r, "indi_vals", axis = 2)

plot(0, 0, type = "n", xlim = c(-5, 5), ylim = c(-5, 5), asp = 1)

for (i in ix) {
  points(s1[, i], s2[, i], pch = ".", cex = 0.5, col = grey(0.1, 0.2))
}
for (i in ix) {
  zz <- MASS::kde2d(s1[, i], s2[, i], n = 51)
  pdata <- contourLines(zz, levels = quantile(zz$z, 0.95))
  polygon(pdata[[1]]$x, pdata[[1]]$y, border = "red", lwd = 1.6)
}
points(colMeans(s1), colMeans(s2), pch = 4, col = "red", lwd = 3)
```

#### Simple model with brms

The simplest implementation of the model, i.e. with interactions from one behavior, can also be fit with brms (Bürkner 2017) but requires some post fitting adjustment. In order to bring the individual level parameters (gregariousness SD and individual gregariousness values) to the same scale, we need to multiply by $\sqrt{0.5}$. The resulting brms estimates correspond fairly closely to the ones from bamoso (see figures 19 and 20).

```
set.seed(50)

true_indi_sd <- 0.8
true_dyad_sd <- 1.3
true_intercept <- -0.2

x <- generate_data(n_ids = 20,
                   n_beh = 1,
                   beh_intercepts = true_intercept,
                   behav_types = c(approach = "count"),
                   count_obseff = 10,
                   indi_sd = true_indi_sd,
                   dyad_sd = true_dyad_sd)

# interaction matrices
imats <- list(approach = x$processed$interaction_matrices[[1]])
# observation effort matrices
omats <- list(x$processed$obseff_matrices[[1]])
# data object
standat <- make_stan_data_from_matrices(mats = imats,
                                        behav_types = c("count"),
                                        obseff = omats)

# data object for brms call (a data.frame)
dyads <- x$input_data$dyad_data[, c("dyad", "id1", "id2")]
dyads$approach <- imats[[1]][as.matrix(dyads[, 2:3])]
dyads$obseff <- omats[[1]][as.matrix(dyads[, 2:3])]
```

```
plot(0, 0, xlim = c(-1, 2), xaxs = "i", ylim = c(0, 6), yaxs = "i", type = "n",
     xlab = "estimate", ylab = "density", bty = "l")
cols <- adjustcolor(hcl.colors(3, palette = "zissou"), 0.5)

legend(1.5, 6, xpd = TRUE, bty = "n", pt.cex = 1.5, cex = 0.7,
       legend = c("true", "bamoso", "brms", "individual", "dyadic", "baseline"),
       lty = c(1, 3, 2, NA, NA, NA),
       pch = c(NA, NA, NA, 15, 15, 15),
       col = c("black", "black", "black", cols[1], cols[2], cols[3])
)

res_brms <- rstan::extract(bres$fit, "sd_mmid1id2__Intercept")$sd_mmid1id2__Intercept
```

```
res_brms <- res_brms * sqrt(0.5)
res_bam <- extract_samples(bamres, "indi_sd")[, 1]
polygon(density(res_brms), col = cols[1])
polygon(density(res_bam), col = cols[1])
abline(v = c(true_indi_sd, median(res_brms), median(res_bam)),
       lty = c(1, 2, 3), col = cols[1], lwd = 2)

res_brms <- rstan::extract(bres$fit, "sd_dyad__Intercept")$sd_dyad__Intercept
res_bam <- extract_samples(bamres, "dyad_sd")[, 1]
polygon(density(res_brms), col = cols[2])
polygon(density(res_bam), col = cols[2])
abline(v = c(true_dyad_sd, median(res_brms), median(res_bam)),
       lty = c(1, 2, 3), col = cols[2], lwd = 2)

res_brms <- rstan::extract(bres$fit, "b_Intercept")$b_Intercept
res_bam <- extract_samples(bamres, "beh_intercepts")[, 1]
polygon(density(res_brms), col = cols[3])
polygon(density(res_bam), col = cols[3])
abline(v = c(true_intercept, median(res_brms), median(res_bam)),
       lty = c(1, 2, 3), col = cols[3], lwd = 2)
```

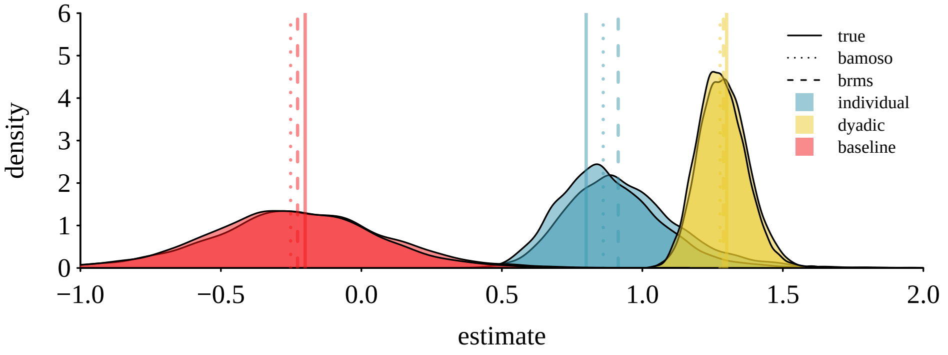

Figure 19: Fitting the simplest version of the sociality model is also possible using the multi-membership syntax in extttbrms. The resulting posteriors of the key parameters are very similar to those obtained from extttbamoso package.

```
# gregariousness (needs adjustment)
x <- ranef(bres)$mmid1id2[, , 1][, "Estimate"]
x <- x * sqrt(0.5)
y <- colMeans(extract_samples(bamres, "indi_vals"))
plot(x, y, xlab = "brms estimates (adjusted)", ylab = "bamoso estimates")
abline(0, 1)

# affinity (no adjustment)
x <- ranef(bres)$dyad[, , 1][, "Estimate"]
y <- colMeans(extract_samples(bamres, "dyad_vals"))
plot(x, y, xlab = "brms estimates", ylab = "bamoso estimates")
abline(0, 1)
```

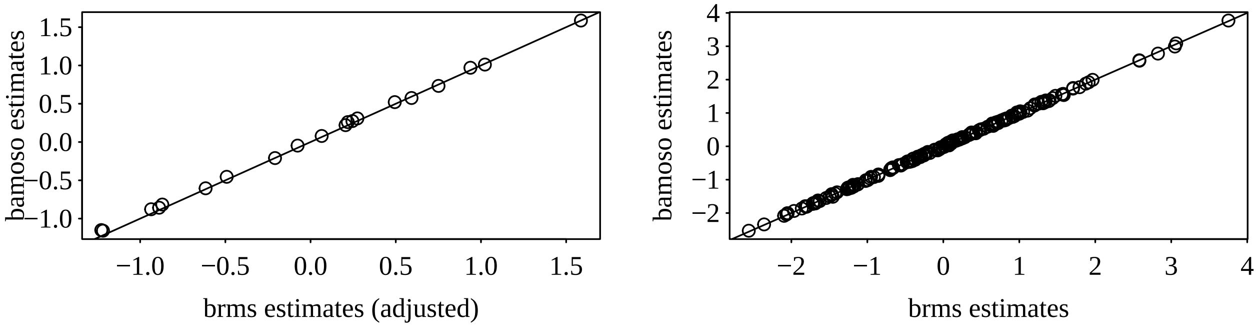

Figure 20: The estimated individual level gregariousness values and the dyadic affinity values also correspond very well between the extttbrms and extttbamoso implementations.

This document took 1.5 minutes to compile.
